# Chemoenzymatic Synthesis of 6-Sulfo Sialyl Lewis^x^ Containing Glycans to Probe the Receptor Specificity of MERS Coronavirus

**DOI:** 10.64898/2026.08.06.743223

**Authors:** Yunfei Wu, Anne L.M. Kimpel, Jacobus P. van Trijp, Elif Uslu, Gaёl M. Vos, Luca Unione, Robert P. de Vries, Geert-Jan Boons

## Abstract

The initial attachment of Middle East Respiratory Syndrome Coronavirus (MERS-CoV) to host cell sialosides is critical for infection, yet its precise receptor specificity remains poorly understood. Here, we describe a chemoenzymatic methodology to synthesize a comprehensive panel of 6-sulfo sialyl Lewis^x^ (6-sulfo-SLe^x^) containing glycans. Our approach entails the enzymatic assembly of an oligo-lactosamine chain modified at specific positions with *N*-trifluoroacetyl-glucosamine (GlcNTFA) moieties. Mild base treatment removes the TFA group to yield glucosamine, which effectively blocks enzymatic fucosylation. By leveraging this approach alongside the unique substrate selectivity of GlcNAc-6-O-sulfotransferases 2 (CHST-2), we achieved the selective preparation of fucosylated 6-sulfo-SLe^x^ glycans. Microarray screening of these printed glycans revealed that a 6-sulfo-SLe^x^ derivative presented on an extended LacNAc chain is the preferred host receptor for MERS-CoV. Conjugation of this lead compound to a polyglycerol-based dendrimer generated a multivalent inhibitor that potently blocks hemagglutination of human red blood cells by the MERS-CoV spike protein *N*-terminal domain (NTD). Furthermore, computational modeling demonstrated that the fucose moiety does not directly contact the viral spike protein. Instead, it pre-organizes the ligand into a favorable conformation, enabling a critical salt bridge between the glycan’s sulfate group and the guanidinium side chain of viral residue Arg307.

## Introduction

Bats and dromedary camels are the natural reservoir of Middle East Respiratory Syndrome Coronavirus (MERS-CoV).^[1–7]^ Occasionally this virus infects humans causing acute respiratory distress and renal failure resulting in a fatality rate of 36%.^[8]^ Coronaviruses are expected to cause future zoonotic events because of the large reservoir in bats and birds. The spillover potential of these viruses is supported by the fact that seven different coronaviruses of animal origin have entered the human population.^[9]^ Understanding the receptor binding properties of coronaviruses is critical to develop surveillance, treatment and vaccine approaches to curb transmission.^[10–11]^

The spike protein (S) of MERS-CoV is composed of two subunits, S1 and S2, in which S1 harbors an N-terminal domain (NTD) and a C-terminal domain (CTD).^[12–17]^ The CTD can bind to dipeptidyl-peptidase 4 (DPP4) which is the proteinaceous entry receptor for this virus. The NTD can bind sialosides, which is important for initial cell attachment.^[18]^ The latter is supported by the observation that treatment of human airway cells by a neuraminidase to remove cell surface sialosides inhibits cell entry. Glycan array screening has indicated that NTD-Fc of MERS-CoV preferentially binds glycans having an α2,3-linked *N*-acetyl neuraminic acid (Neu5Ac) including sialyl Lewis^x^ (SLe^x^) and 6-sulfo-SLe^x^ (6-sulfo-SLe^x^).^[18]^ The latter compound gave the highest response on the glycan array. The importance of the sulfate and fucoside of 6-sulfo-SLe^x^ for binding could, however, not clearly be established due to a lack of structurally diverse glycans on the employed array.^[18–19]^ A cryo-EM structure of the spike of MERS-CoV with SLe^x^ demonstrated that Gln36, Phe39, Phe101, Ile132 and R307 of MERS-CoV can make critical interactions with Neu5Ac.^[20]^ Sequence alignments of MERS-CoV isolates showed that residues that contact the sialoside or participate in the formation of the binding groove are conserved. Mutations of amino acids of the sialic acid binding site resulted in loss of infectivity, highlighting the importance of sialoside binding for infectivity.

6-Sulfo-SLe^x^ (Neu5Acα2,3Galβ1,4(Fucα1,3)GlcNAc(6-SO_3_^−^)β-OR; Figure 1a) is a terminal epitope found on several classes of glycoconjugates including *O*- and *N*-glycans. It is a ligand for L-selectin, human sialic acid-binding immunoglobulin-like lectin 9 (Siglec 9) and various influenza viruses.^[21–26]^ Membrane bound mucins of the periciliary layer of airway surfaces abundantly express keratan sulfate (KS),^[27]^ which can harbor terminal epitopes such as SLe^x^ and 6-sulfo-SLe^x^. KS is composed of poly-*N*-acetyl-lactosamine (poly-LacNAc) chains that are modified by sulfation at C-6 positions of *N*-acetyl-glucosamine (GlcNAc) and galactose (Gal) residues. The terminal LacNAc unit can be 2,3- or 2,6-sialylated. Furthermore, KS-II, which is attached to *O*-linked glycans *via* a core-2 structure, is further decorated by 1,3-linked fucosides at GlcNAc to give Le^x^ and SLe^x^ epitopes and their sulfated counterparts.^[28]^

**Figure 1.**
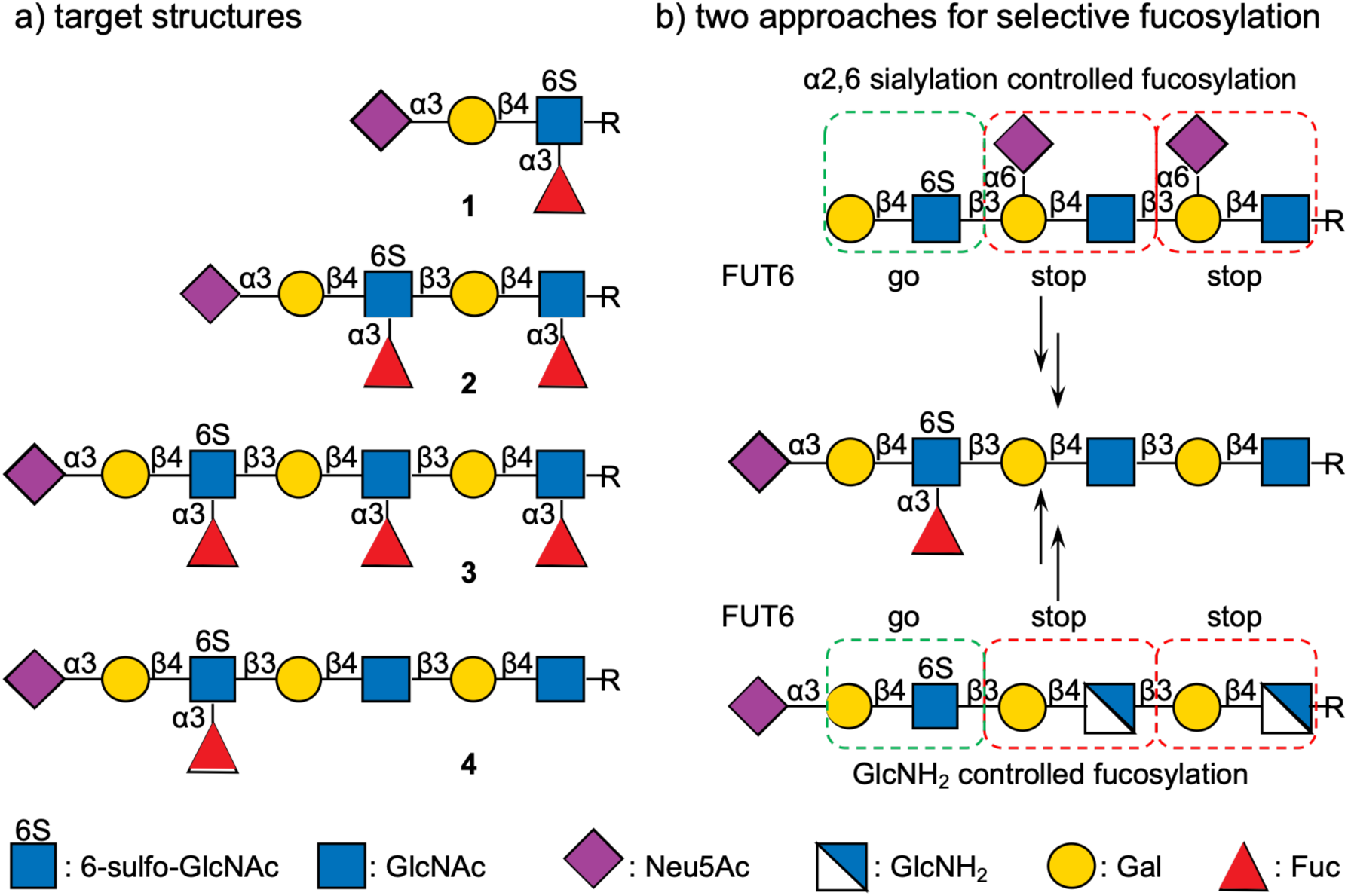
a) 6-Sulfo sialyl Lewis^x^ is a ligand of MERS-CoV NTD; b) 2 Synthetic approaches of terminal 6-sulfo sialyl Lewis^x^. R = O(CH_2_)_5_NHCbz or O(CH_2_)_5_NH_2_ or O(CH_2_)_5_NHCO_2_CH_2_BCN.

We were compelled to further investigate the glycan binding properties of the NTD of MERS-CoV to establish the importance of the sulfate and fucoside of 6-sulfo-SLe^x^ and the presentation of this epitope on a poly-LacNAc chain as in KS-II. We set out to synthesize compounds **1**-**4** (Figure 1a) for binding studies. The synthesis of compound **4** was challenging because it requires the selective introduction of a fucoside at the terminal LacNAc moiety to generate 6-sulfo-SLe^x^ without modifying the other LacNAc units. Although chemical and chemoenzymatic strategies have been reported for 6-sulfo-SLe^x^,^[21, 29–31^] these strategies do not allow for the precise installation of fucosides such as in compound **4**. We explored various strategies to prepare compounds such as **4** and established an efficient strategy utilizing an oligo-lactosamine chain containing site-specific *N*-trifluoroacetyl-glucosamine (GlcNTFA) modifications. Mild basic deprotection of the TFA groups generates glucosamine residues that blocks subsequent unwanted enzymatic fucosylation. By combining this masking technique with the terminal-specific activity of GlcNAc-6-O-sulfotransferase 2 (CHST-2), compound **4** could readily be prepared. Glycans **1**–**4** and a series of reference compounds were printed as a microarray to evaluate the binding specificity of the MERS-CoV NTD. Notably, 6-sulfo-SLe^x^ presented on an extended oligo-LacNAc backbone bound with significantly higher avidity than the 6-sulfo-SLe^x^ epitope alone. Computational modeling revealed that the sulfate participates in a direct, highly favorable electrostatic interaction with an arginine residue adjacent to the sialic acid-binding pocket. Conversely, the fucose moiety does not engage in direct protein contacts; instead, it pre-organizes the ligand into an optimal conformation for receptor binding. Finally, conjugation of glycan **4** to a polyglycerol-based dendrimer generated a multivalent derivative that potently inhibited MERS-CoV NTD attachment to host cells, demonstrating potential for downstream therapeutic development.

## Results and Discussion

### Controlling α1,3-Fucosylation to Selective Install a 6-Sulfo-SLe^x^ Epitope on PolyLacNAc Chains

We explored several chemoenzymatic strategies to prepare poly-LacNAc chains having a 6-sulfo-SLe^x^ epitope such as in compound **4** (Figure 1). The biosynthesis of 6-sulfo-SLe^x^ involves the sulfation of the C-6 hydroxyl of a terminal GlcNAc residue by GlcNAc-6-*O*-sulfotransferases 2 or 6 (CHST2 or 6)^[32]^ in the presence of 3′-phosphoadenosine-5′-phosphosulfate (PAPS),^[33–34]^ which is followed by β1,4-galactosylation by β(1,4)-galactosyl transferases 4 (B4GalT4) and further 2,3-sialylation by α2,3-sialyltransferase 4 (ST3Gal4). The resulting 6-sulfo-sialyl-LacNAc moiety is then α1,3-fucosylated by an appropriate fucosyl transferase to form 6-sulfo-SLe^x^. The challenge of the enzymatic synthesis of 6-sulfo-SLe^x^ containing poly-LacNAc derivatives, such as **4**, is the regioselective installation of an α1,3- fucoside. To synthesize glycan **4**, we exploiting that glucosamine (GlcNH_2_) resists modification by fucosyl transferases.^[35–36]^ Such a residue can be chemo-enzymatically installed into a poly-LacNAc chain by employing UDP-GlcNTFA and an appropriate GlcNAc transferases followed by hydrolysis of the TFA moiety.^[37]^ At an appropriate stage in the synthesis, the GlcNH_2_ residues can be selectively acetylated to provide natural GlcNAc-containing glycans.

The synthesis of **4** started from *N*-acetyl-lactosamine modified by an anomeric benzyloxycarbonyl (Cbz)-protected aminopentyl spacer that was extended by an *N*-trifluoracetyl lactosamine moiety by the consecutive action of Hpβ3GlcNAcT in the presence of UDP-GlcNTFA and B4GalT1 in combination with UDP-Gal to give tetrasaccharide **5**. The latter compound was enzymatically extended by a 1,3-linked GlcNAc residue using Hpβ3GlcNAcT and UDP-GlcNAc to yield pentasaccharide **6** (Scheme 1). The terminal GlcNAc moiety of **6** was selectively sulfated by CHST2 in the presence of PAPS to provide compound **7**. In this transformation, we exploited that CHTS2 only modifies terminal GlcNAc residues.^[38]^ Control of the pH during this transformation was important to avoid loss of the TFA group. Terminal 6-sulfo-GlcNAc of compound **7** was further extended by the action of B4GalT4 in the presence of UDP-Gal to give compound **8**, which was modified by an α2,3-linked sialoside using ST3Gal4 and CMP-Neu5Ac resulting in the formation of **9**. Hydrolysis of TFA moiety of **9** was achieved by treatment with aqueous NH_4_OH producing **10** without affecting the sulfate ester. As anticipated,^[35]^ the GlcNH_2_ moieties at the reducing end and central position of compound **10** resisted fucosylation by FUT6 resulting in the selective modification of the GlcNAc moiety to form 6-sulfo-SLe^x^ containing derivative **11**. Finally, the free amines of the latter compound were acetylated with AcOSu in the presence of DIPEA without affecting the sulfate producing target compound **4**, which was purified by size exclusion column chromatography over Bio-Gel P6 using 50 mM NH_4_HCO_3_ as eluent. Detailed NMR analysis confirmed the structural integrity of the compounds (see supporting information). The 1D ^1^H NMR and 2D ^1^H−^13^C HSQC spectra made it possible to assign all proton and carbon signals. The H-3 of the fucosylated GlcNAc6S moiety had substantially shifted from δ3.74 to δ3.91 and the corresponding nearby H-2 also shifted from δ3.84 to δ4.00 which confirmed the regioselectivity of the α1,3-fucosylation of GlcNAc6S.^[38–39]^

**Scheme 1.**
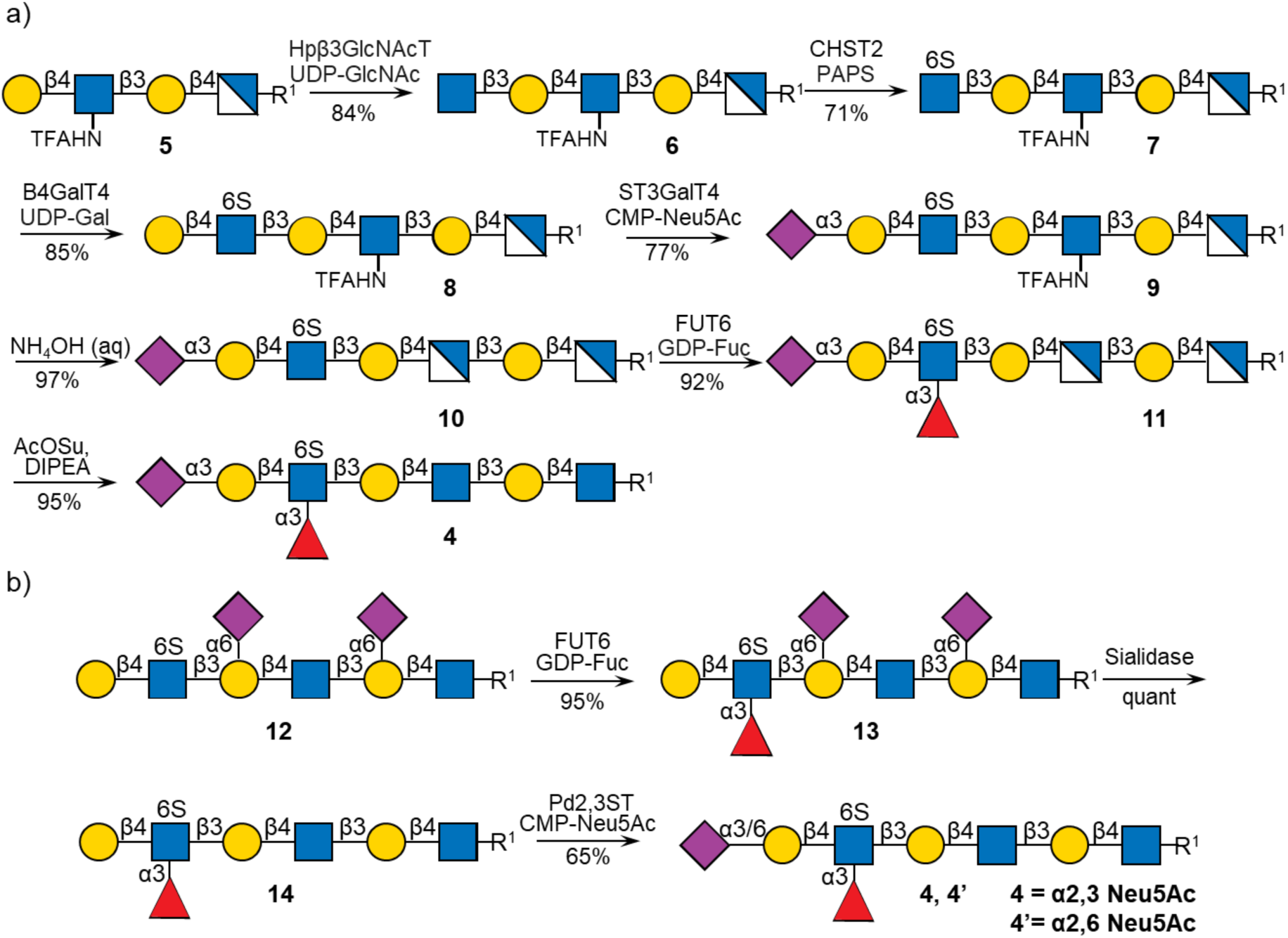
a) Enzymatic assembly line for site-specific fucosylation UDP-GlcNTFA control. b) Enzymatic assembly site-specific fucosylation by 2,6 sialylation control. **4** is major α2,3 Neu5Ac product, **4’** is minor α2,6 Neu5Ac side product. R^1^= O(CH_2_)_5_NHCbz.

We also examined an earlier developed strategy to prepare selectively fucosylated glycan **4** by employing 2,6-sialosides at the reducing and central LacNAc moiety to selectively block fucosylation (Scheme 1b).^[39]^ The bacterial α2,6-sialyltransferase from *Photobacterium damselae* (Pd2,6ST)^[40–41]^ can sialylate both terminal and internal galactosides and these residues can block specific sites of fucosylation. Subsequent removed of the sialosides by a sialidase was anticipated to provide entry into 6-sulfo-SLe^x^ containing glycans.^[39, 41^] To implement this strategy, it was essential to generate first a terminal 6-sulfo-Le^x^ containing compound followed by sialidase treatment and then introduction of a terminal α2,3-sialoside using a β-galactoside α2,3-sialyltransferase.^[42]^ Thus, compound **12**, containing two internal LacNAc moieties masked by α2,6-sialosides, was prepared by our previously developed methodology.^[39]^ This compound could selectively be fucosylated at the terminal 6-sulfo-LacNAc moiety by FUT6 in the presence of GDP-Fuc, resulting in the formation of **13** (Scheme 1b). The latter compound was treated with the sialidase of *C. perfringens* to remove the 2,6-linked sialosides to give **14**. 2,3-Sialylation of glycan **14** bearing a terminal 6-sulfo-Le^x^ moiety was challenging. As expected, the human sialyltransferase, ST3Gal4, was unable to install an α2,3-sialoside (Scheme S3). Neither the bacterial sialyltransferase PmST3 nor CSTI facilitated efficient α2,3-sialylation to form the terminal 6-sulfo-SLe^x^ epitope.^[43–44]^ Owing to its high reactivity and ease of expression, we explored the use of PmST1 M144D, which is a mutant α2,3-sialyltransferase from *Pasteurella multocida* that has been employed in the synthesis of sialyl Le^x^ (SLe^x^) and sialyl 6-sulfo LacNAc.^[35, 39, 45]^ Treatment of compound **14** with an excess of CMP-Neu5Ac resulted in a mixture of the α2,3-sialoside **4** and the α2,6-sialoside **4’** in a 1:1.7 ratio. It was difficult to separate these two positional isomers (Scheme S3). The lack of regioselectivity is likely due to the multifunctional enzymatic reactivity of PmST1 M144D, which despite the mutation conferring predominant α2,3-sialyltransferase activity, still retains some α2,6-sialyltransferase, *trans*-sialidase, and sialidase activity.^[46]^ The fucoside of 6-sulfo-Le^x^ of compound **14** probably impedes α2,3-sialyltransferase activity of PmST1 M144D, requiring prolonged reaction times and excess of CMP-Neu5Ac. It is likely that the sialidase activity gradually degrades the α2,3-sialoside of **4** while the α2,6-sialoside of **4’** remains intact resulting in accumulation of the latter compound. Pd2,3ST, which is a multifunctional sialyl transferase identified from the *P. dagmatis* genome, also exhibits α2,3-sialyltransferase, α2,3-sialidase, and α2,3-*trans*-sialidase activity.^[47]^ While Pd2,3ST shows sialidase and *trans*-sialidase activities at low pH (pH 4.5), these activities diminish at neutral conditions, while its transferase function remains intact. Compound **14** was treated with Pd2,3ST and CMP-Neu5Ac (10 eq) in a 50 mM MOPS buffer at pH 7.2 resulting in the formation of a mixture of **4** and **4’** in a more favorable ratio of 10:1. The mixture of products was first subjected to size-exclusion column chromatography followed HPLC using a HILIC column providing compound **4** in a yield of 65% at 2 mg scale.

It is unclear which α1,3-fucosyltransferase (FUT3–7, FUT9–11) can readily form a 6-sulfo-SLe^x^ epitope.^[48–51]^ To address this question, sialyl 6-sulfo LacNAc derivatives (**15**-**17**) were incubated with the human fucosyltransferases FUT5, FUT6, and FUT9 to attempt to synthesize 6-sulfo-SLe^x^ containing glycans (Scheme 2). Treatment of the compounds with FUT5, FUT6, or FUT9 and GDP-Fuc (1.5 eq per LacNAc) resulted in fully fucosylated 6-sulfo-SLe^x^ derivatives **1**-**3**, along with varying amounts of partially fucosylated side products. Notably, FUT6 provided the highest yield of fucosylated glycans (**1** 96%, **2** 94%, **3** 68%) and was superior to the use of FUT5 and FUT9. To assess the substrate specificities of the FUTs, GDP-fucose was added in a rate-controlled manner. When compound **17** was incubated with FUT6 and GDP-Fuc (0.9 equiv), a mixture of fucosylated mono-LacNAc moieties was obtained. Similarly, when compound **17** was incubated with FUT5 or FUT9 and GDP-Fuc (0.9 equiv), internal LacNAc mono-fucosylated **S1** was produced as the major compound in the mixture (Scheme S1a). It has been reported that the selectivity of FUT9 is reversed to the terminal LacNAc moiety upon removal of the terminal sialic acid.^[52–53]^ However, FUT9 showed only a modest distal preference for its non-sialylated substrate (Scheme S1b), resulting in a mixture of internally fucosylated LacNAc **S3** and terminally fucosylated LacNAc **14**. Although fucosyltransferases FUT3, FUT4 and FUT7 are also capable of installing α1,3-fucosides, they were not readily accessible and were therefore excluded from evaluation.

**Scheme 2.**
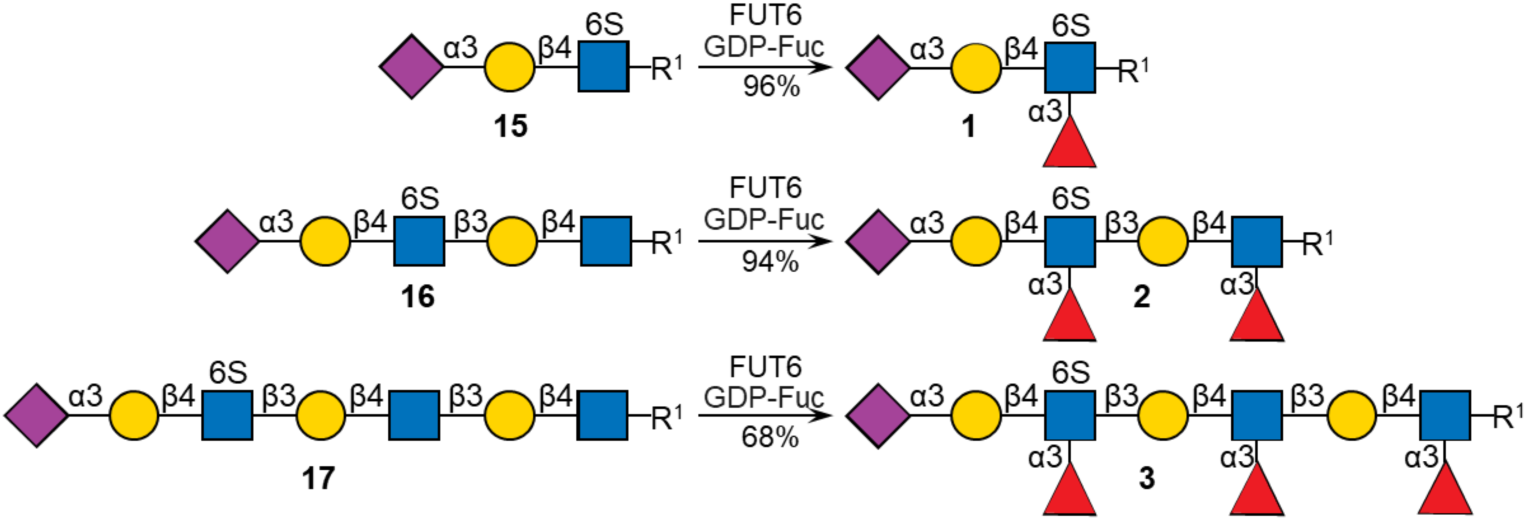
Enzymatic assembly line for multiple fucosylation. R^1^= O(CH_2_)_5_NHCbz.

### Glycan Array Screening to Examine the Binding Specificity of MERS-CoV NTD

We developed a glycan microarray to investigate the glycan binding properties of MERS-CoV NTD. Thus, the Cbz protecting group of the anomeric linkers of compounds **1**-**4** was removed by hydrogenation over Pd(OH)_2_, and the resulting products purified by P6 size-exclusion chromatography using 50 mM ammonium bicarbonate as the eluent yielding the corresponding aminopentenyl derivatives **G**, **N**, **X**, and **Y**, respectively (Figure 2 and Scheme S4). The newly synthesized glycans and several reference compounds, including different linear glycans presenting various relevant epitopes (Figure 2), were printed onto amine-reactive *N*-hydroxysuccinimide (NHS)-activated glass slides.^[39]^ In addition to MERS-CoV NTD, the resulting glycan microarray was also used to probe the binding selectivity of sialic acid-binding immunoglobulin-type lectin-9 (Siglec-9), which is known to preferentially bind to sialoglycans including SLe^x^ and 6-sulfo-SLe^x^.^[54–55]^

**Figure 2.**
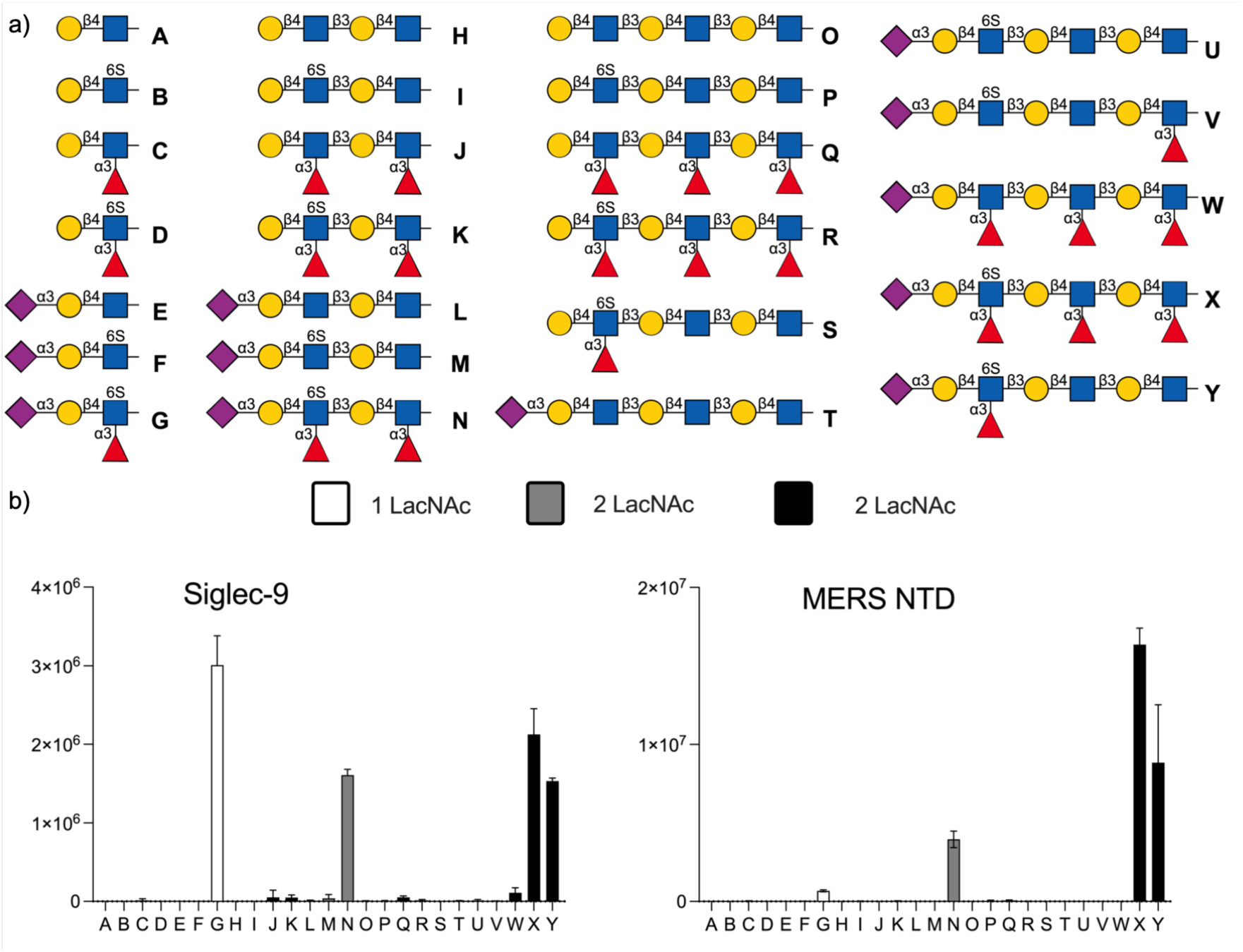
Probing receptor properties of Siglec-9 and MERS-NTD by microarray technology.

Multivalency is important for the interaction of glycans with Siglec’s and viruses and is an important determinant of avidity and selectivity.^[56–57]^ Therefore, Siglec-9 and MERS-CoV NTD were expressed as Fc-fusion proteins that were employed to be displayed on pA-LS nanoparticles.^[58]^ The pA-LS assembling nanoparticle platform is based of Lumazine Synthase (LS) from *Aquifex aeolicus* fused to domain B of protein A (pA) from *Staphylococcus aureus*. LS is a protein that self-assembles into 60-mer icosahedral virus-like nanoparticles and pA binds with high-affinity to the Fc region of Fc-tagged proteins. Accordingly, Fc-tagged Siglec-9 and MERS-CoV NTD proteins were incubated with pA-LS particles for display on the nanoparticle surface, which were employed to probe glycan binding selectivities using glycan binding array technology. Detection was accomplished by anti-strep antibody.

Siglec-9 preferentially bound to glycans having a 6-sulfo-SLe^x^ epitope and in this case the length of the LacNAc chain did not impact binding (**G** *vs.* **N** *vs.* **X**, **Y**). MERS-CoV NTD (Figure 2) also displayed a preference for glycans having a 6-sulfo-SLe^x^ epitope but unlike Siglec-9, the binding greatly depended on the length of the LacNAc chain and glycans having a longer backbone gave much greater responsiveness (**G**, **N** *vs.* **X**, **Y**). The sulfate (**X** *vs.* **W**) and the fucoside of 6-sulfo-SLe^x^ are also important for recognition (**X**, **Y** *vs.* **U**, **V**). However, an absence of fucosides at the internal or reducing LacNAc moieties (**V** *vs.* **X**, **Y**) had little impact on MERS-CoV NTD binding. We also investigated the binding of Siglec-9 and MERS-CoV NTD to *O*-glycans and found that core 2 structures having a 6-sulfo-SLe^x^ epitope were well recognized by both proteins (Figure S2). Some binding was observed for glycopeptides lacking the fucoside of 6-sulfo-SLe^x^.

Furthermore, several H5N1 influenza virus HAs reported to recognize the 6-sulfo-SLe^x^ epitope, including A/Vietnam/1194/04, A/Anhui/05, A/European polecat/NL/1/22 and A/Texas/37/24, were selected for evaluation. No marked selectivity toward glycan backbone length was observed (Figure S3).

### Inhibition of MERS-CoV by 6-sulfo-SLe^x^ Conjugated to Polyglycerol-Based Dendrimers

The recognition of glycans of host cells by a pathogen is pivotal for many viral infections. Although the binding affinity between a glycan receptor and glycan binding protein of a virus is generally low, a multivalently display of glycan receptors can result in high avidity of binding^[59–60]^ to prevent binding of a pathogen to glycans of host cells.^[61–63]^ Thus, we were compelled to synthesize multivalent 6-sulfo-SLe^x^ derivatives to investigate inhibitory activities of binding of MERS-CoV NTD to cells (Figure 3). Thus, 6-sulfo-SLe^x^ containing glycans **G**, **X**, **Y**, together with lactosamine and 3sialyl lactosamine as control, were modified by bicyclo[6.1.0]nonyne (BCN)^[64]^ by reactions with *N*-hydroxysuccinimide (NHS)-BCN to give derivatives **19**-**23**. The latter compounds were conjugated to azide-functionalized polyglycerol-based dendrimers *via* strain-promoted alkyne–azide cycloaddition (SPAAC) to give multivalent derivatives **24**-**28**.^[65–68]^ The conjugates were analyzed by NMR to determine total amount and molar quantity of glycan (see SI). The concentration of glycan was determined by proton NMR using (3-(trimethylsilyl)-2,2,3,3-tetradeuteropropionic acid (TMSP-d_4_) as internal standard. The known concentration of TMSP-d_4_ was compared by integration of the trimethylsilyl peaks at δ 0.0 to the characteristic peaks of the equatorial H3 of the sialic acid of 6-sulfo-SLe^x^-containing dendrimers at δ 2.7, as well as the anomeric protons of the negative control disaccharides **19** (lactose), at δ 4.5. The average glycan loading on the dendrimers was calculated based on the mass ratio of the conjugated glycan to the total weigh, determined gravimetrically of the glycan-dendrimer conjugates. This analysis revealed an average functionalization degree of approximately 50% for each dendrimer.

**Figure 3.**
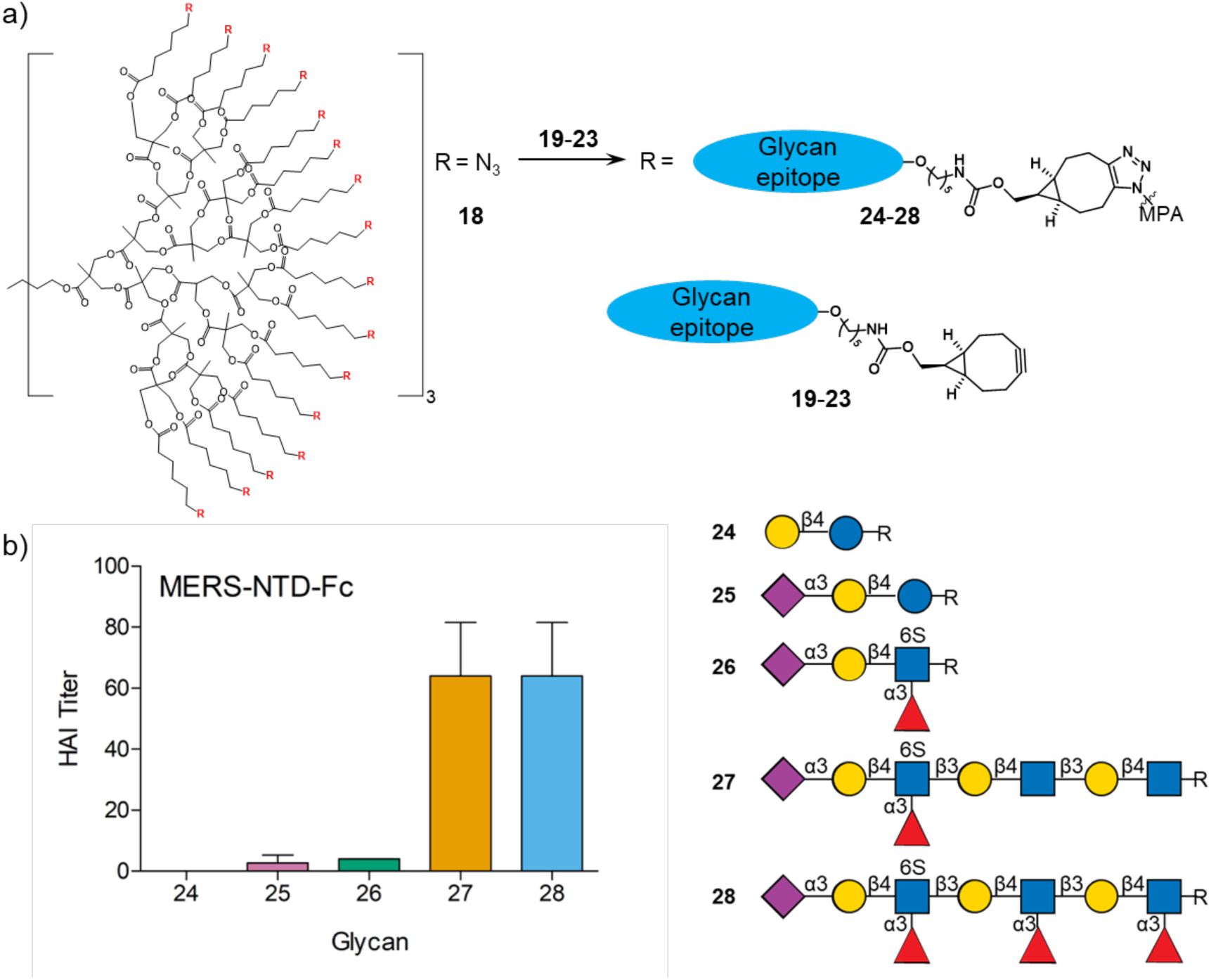
Inhibition of MERS-CoV NTD binding to human erythrocytes by 6-sulfo-SLe^x^-conjugated polyglycerol nanoparticles. For **18**, R= N_3_. For **24**-**28**, R= the clicked linked glycans.

The inhibitory potencies of glycodendrimers **24**-**28** were evaluated using a hemagglutination inhibition (HAI) assay with human erythrocytes and virus-like nanoparticles displaying the MERS-CoV N-terminal domain (NTD). This assay exploits the capacity of viral proteins to cross-link red blood cells (RBCs) *via* surface glycan interactions, a process that can be disrupted by soluble competitive inhibitors targeting the viral lectin domain.^[69]^ The concentration of protein A-coated lipid nanoparticles (pA-LNPs) functionalized with MERS-CoV-NTD-Fc was adjusted to a standardized agglutination titer of 8. To ensure an accurate comparison across the structurally distinct architectures, all dendrimers were normalized to the total molar quantity of glycan present on the dendritic scaffold, as quantified by ^1^H NMR spectroscopy (see above).

Serial dilutions of glycodendrimers **24**-**28** were pre-incubated with the NTD-displaying nanoparticles prior to the addition of erythrocytes to determine the minimum inhibitory concentration (MIC), defined as the lowest ligand concentration required to completely suppress agglutination. Dose-dependent hemagglutination inhibition was observed for the 6-sulfo-SLex-functionalized dendrimers **26**-**28**, whereas the control scaffolds **24** and **25** showed no inhibitory activity. Glycodendrimers **27** and **28** emerged as potent inhibitors, achieving a maximal HAI titer of 64, which corresponds to a glycan monomer concentration of 312 nM. These findings reinforce the microarray data indicating that an extended LacNAc backbone substantially enhances 6-sulfo-SLex recognition by the MERS-CoV NTD, establishing glycodendrimers **27** and **28** as promising lead candidates for therapeutic intervention.

### In Silico Studies of Binding of MERS-CoV Spike to Sulfo-SLe^x^

Docking studies were performed of the complex of MERS-CoV spike protein and glycan **4** that presents 6-sulfo-SLe^x^ on an extended LacNAc chain. The resulting models showed a very similar pose of the sulfated ligand to that observed in cryo-EM structure of MERS-CoV spike in complex with SLe^x^.^[20]^ The sialic acid residue interacts within a groove of the *N*-terminal domain of the spike protein located ∼50 Å away from the proteinaceous receptor (DPP4) binding site. The two major pendant groups of the Neu5Ac residue are accommodated into two hydrophobic depressions on the protein surface (Figure 4). Specifically, the *N*-acetyl moiety establishes methyl-π interactions with the aromatic side chains of Phe39, Phe101, and His91, reinforced by the hydrophobic interaction with Ile132. The orientation of acetamide within the hydrophobic pocket is guided by hydrogen bonds with Gln36 and Ile132 backbone residues. In addition, the glycerol side chain of sialic acid makes hydrogen bonds with His91 and Ala92. Finally, the carboxylate group of the Neu5Ac residue makes a bidentate hydrogen bond interaction with the side chains of Ser133 and Ser135 (Figure 4a). This observation is in line with a reported structure of the hemagglutinin proteins of A/H3N2 influenza viruses (NL03, NL09, and Sing16) in which hydroxyls of two contiguous serine residues chelate the sialoside through its carboxylate.^[70]^

**Figure 4.**
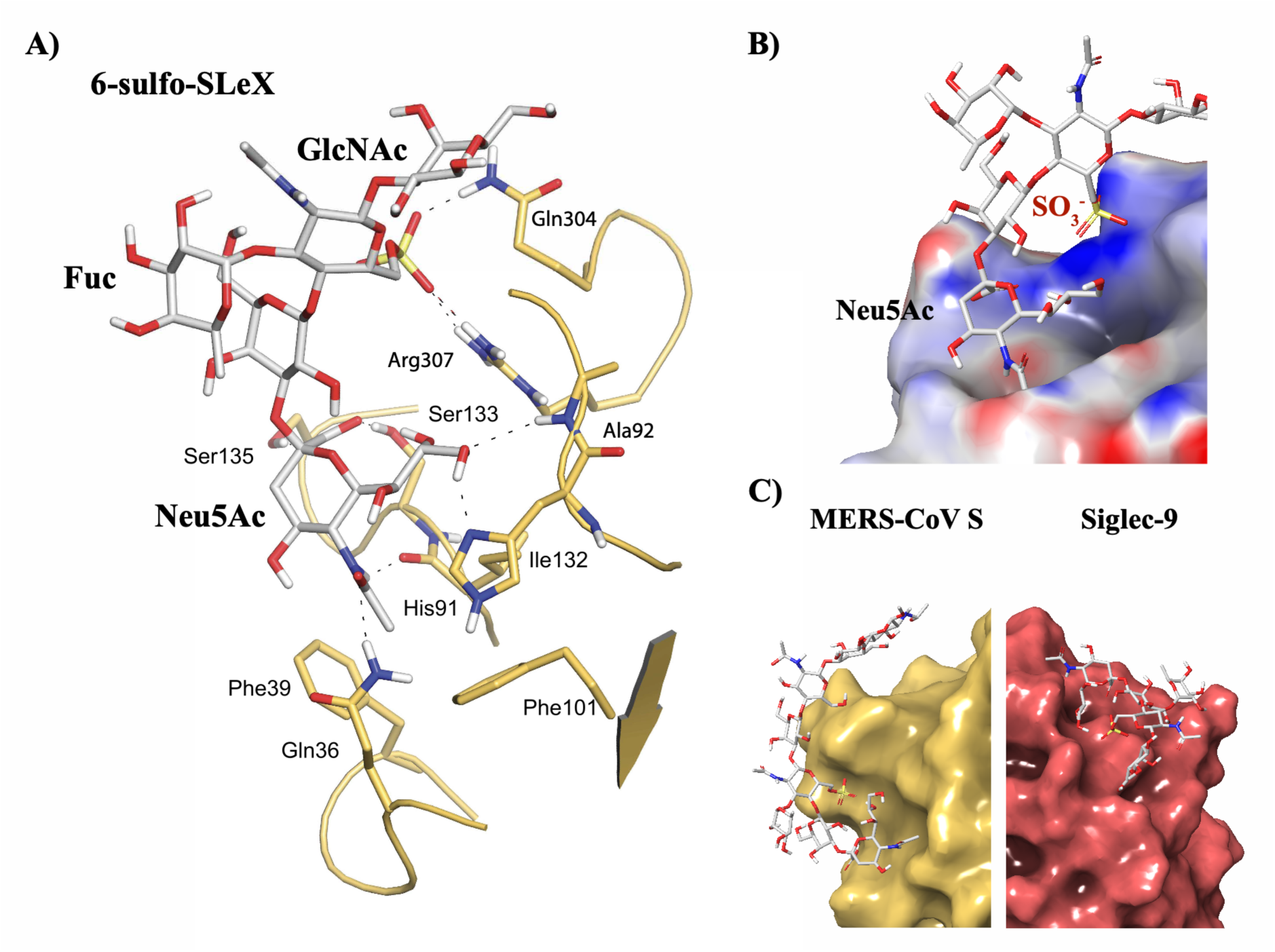
Structural basis of MERS-CoV S selectivity for 6-sulfo-SLe^x^. A) Intermolecular interaction network as derived from docking studies. The protein is rendered as a ribbon diagram. Selected side chains of interacting protein residues and the ligand are shown as sticks. B) Molecular surface representation of the ligand-binding site colored by electrostatic potential. Blue reports for positively charged patch and red for negatively charged patch on the protein surface. C) Comparison of the deep MERS-CoV S and shallow Siglec-9 carbohydrate binding site.

The array experiments indicate a preference of MERS-CoV NTD for fucosylated epitopes. Yet the *in-silico* derived model shows that while the galactoside and *N*-acetylglucosamine of the 6-sulfo-SLe^x^ moiety of **4** are positioned within contact distance of the protein surface, the fucoside points in the opposite direction and does not make direct contact with the protein. In complex glycans, branching residues can restrict the conformational flexibility of an oligosaccharide which can modulate thermodynamic and kinetic parameters of binding.^[71]^ It is known that Le^x^-type structures adopt a rather rigid closed conformation in solution stabilized by hydrophobic interactions between the methyl group of the fucoside and the β-face of the galactoside, aided by a non-conventional CH···O hydrogen bond between the H-5 of the fucose and the O-5 of the galactoside.^[72–77]^ This well-defined hairpin structure limits the flexibility around the Galβ1-4GlcNAc glycosidic linkage. In case of the 6-sulfo-SLe^x^ in complex with the MERS-CoV S protein, the pre-organized conformation of the ligand places the sulfate group at contact distance of a positively charged patch proximal to the sialic acid binding site (Figure 4b). The sulfate establishes a salt bridge with the Arg307 side chain guanidium, further reinforced by a hydrogen bond with Gln304 (Figure 4a).

The microarray results indicate that Siglec-9 recognize sulfated SLe^x^ epitopes largely independent of the underlying saccharide moiety, whereas MERS NTD showed a strong preference for compounds having the sulfated SLe^x^ epitope presented on an extended LacNAc chain (compound **Y** in Figure 2). The *in-silico* modeling indicates, however, that only the terminal tetrasaccharide makes direct contacts with MERS NTD, while the underlying poly-LacNAc chain remains solvent-exposed. Comparison of the glycan binding site architecture of Siglec-9 and MERS NTD reveals key structural differences that most likely provide a rationale for the distinct glycan recognition profiles (Figure 4c). Siglec-9 features a shallow, solvent-exposed binding pocket that can accommodate sulfo-SLe^x^ independent from the further oligosaccharide chain. In contrast, the carbohydrate-binding site of MERS-CoV spike is positioned approximately 50 Å below the exposed proteinaceous receptor-binding site for DPP4. Accordingly, extended glycans facilitate effective presentation of the sulfated SLe^x^ epitope for receptor recognition.

## Conclusions

Sialosides play an important role in the initial attachment of MERS-CoV to host cells. Previous studies have indicated that α2,3-linked sialosides, including sialyl Lewis^x^ (SLe^x^) and 6-sulfo-SLe^x^ (6-sulfo-SLe^x^), are recognized by MERS-CoV NTD.^[18, 78]^ The importance of the sulfate and fucoside of 6-sulfo-SLe^x^ for binding was, however, not systematically studied.^[18–19]^ We performed an initial screen of a previously developed glycan microarray and found that MERS-CoV NTD has a preference for 6-sulfo-SLe^x^ presented on an extended LacNAc moiety (compound **3**). Such structural elements are part of keratan sulfates, which are proteoglycans found on respiratory tissue.^[27]^ To further explore the structure-binding relationship of MERS-CoV NTD, we explored various chemoenzymatic approaches to prepare 6-sulfo-SLe^x^ containing derivatives presented on an oligo-LacNAc chain (*e.g.* compound **4**). To prepare such compounds, various approaches were explored for the regioselective installation of fucosides on an oligo-LacNAc chain. In the first strategy, 2,6-linked sialosides, which can readily be installed by the bacterial α2,6-sialyltransferases Pd2,6ST, were employed to block certain LacNAc moieties from fucosylation.^[39]^ Next, the 2,6-sialosides were removed by a sialidase to give a compound having a terminal 6-sulfo-Le^x^ moiety, which was subjected to various sialyl transferases. However, in each case a mixture of 2,3- and 2,6-linked sialosides was obtained. The targeted compounds could, however, be prepared in a regioselective manner by exploiting that glucosamine (GlcNH_2_) is resistant to modification by fucosyl transferases. Such a residue can be installed using UDP-GlcNTFA followed by hydrolysis of the TFA moieties.^[37]^ In this approach, the sulfate and sialoside are introduced before fucosylation thereby avoiding the formation of a mixture of 2,3- and 2,6-linked sialosides. These resulting compounds and several reference derivatives were printed as a glycan micro-array to explore the receptor specificity of MERS-CoV NTD. It was found it prefers compounds having 6- sulfo-SLe^x^ presented on an extended LacNAc chain. The microarray screening also indicated that the sulfate and fucoside of 6-sulfo-SLe^x^ are important for binding. Computational studies revealed that the fucoside of 6-sulfo-SLe^x^ does not directly interact with the protein but pre-organizes it in a conformation that allows a salt bridge between the sulfate of 6-sulfo-SLe^x^ and the guanidinium side chain of Arg307 of the spike, which is further reinforced by a hydrogen bond with Gln304. Several 6-sulfo-SLe^x^ derivatives were attached to a polyglycerol-based dendrimer, and it was found that a compound having 6-sulfo-SLe^x^ presented on an extended LacNAc chain can potently inhibit hemagglutination of human red blood cells by MERS-CoV NTD. These results underscore the importance of presentation of 6-sulfo-SLe^x^ on an extended LacNAc chain and provide a lead compound for the development of an anti-MERS-CoV agent.

### Experimental Section

Experimental details can be found in the Supporting Information. The Supporting Information (PDF) contains synthetic protocols, compound characterization, experimental procedure for enzyme and protein expression, microarray screening, nanoparticles hemagglutination inhibition assay, computational approaches, copies of NMR spectra, LC-MS data and HPLC traces.

## Acknowledgements

The research was supported by the European Research Council (grants nr. 101020769 to G.J.B.). L.U. is supported by the European Research Council ERC-2023-STG (project 101117639-Glyco13Cell) and by the Agencia Estatal de Investigación of Spain MCIN/AEI (project PID2022-142639OA-I00). We thank Bart Straten (Utrecht University) for providing a starting material and useful discussions.

## Conflict of Interest

The authors declare no conflict of interest.

## Data Availability Statement

The data that support the findings of this study are available in the Supporting Information of this article.

## Notes

### Competing Interest Statement

The authors have declared no competing interest.

## References

1 A. M. Zaki, S. van Boheemen, T. M. Bestebroer, A. D. Osterhaus, and R. A. Fouchier “Isolation of a Novel Coronavirus from a Man with Pneumonia in Saudi Arabia,” N. Engl. J. Med. 367 (2012): 1814–1820. 10.1056/NEJMoa1211721

2 R. J. de Groot, S. C. Baker, R. S. Baric, C. S. Brown, C. Drosten, L. Enjuanes, R. A. Fouchier, M. Galiano, A. E. Gorbalenya, Z. A. Memish, S. Perlman, L. L. Poon, E. J. Snijder, G. M. Stephens, P. C. Woo, A. M. Zaki, M. Zambon, and J. Ziebuhr “Middle East Respiratory Syndrome Coronavirus (MERS-CoV): Announcement of the Coronavirus Study Group,” J. Virol. 87 (2013): 7790–7792. 10.1128/JVI.01244-13

3 B. L. Haagmans, S. H. Al Dhahiry, C. B. Reusken, V. S. Raj, M. Galiano, R. Myers, G. J. Godeke, M. Jonges, E. Farag, A. Diab, H. Ghobashy, F. Alhajri, M. Al-Thani, S. A. Al-Marri, H. E. Al Romaihi, A. Al Khal, A. Bermingham, A. D. Osterhaus, M. M. AlHajri, and M. P. Koopmans “Middle East Respiratory Syndrome Coronavirus in Dromedary Camels: An Outbreak Investigation,” Lancet Infect. Dis. 14 (2014): 140–145. 10.1016/S1473-3099(13)70690-X

4 V. D. Menachery, B. L. Yount, Jr., K. Debbink, S. Agnihothram, L. E. Gralinski, J. A. Plante, R. L. Graham, T. Scobey, X. Y. Ge, E. F. Donaldson, S. H. Randell, A. Lanzavecchia, W. A. Marasco, Z. L. Shi, and R. S. Baric “A Sars-Like Cluster of Circulating Bat Coronaviruses Shows Potential for Human Emergence,” Nat. Med. 21 (2015): 1508–1513. 10.1038/nm.3985

5 V. D. Menachery, B. L. Yount, Jr., A. C. Sims, K. Debbink, S. S. Agnihothram, L. E. Gralinski, R. L. Graham, T. Scobey, J. A. Plante, S. R. Royal, J. Swanstrom, T. P. Sheahan, R. J. Pickles, D. Corti, S. H. Randell, A. Lanzavecchia, W. A. Marasco, and R. S. Baric “SARS-Like WIV1-CoV Poised for Human Emergence,” Proc. Natl. Acad. Sci. U. S. A. 113 (2016): 3048–3053. 10.1073/pnas.1517719113

6 J. S. Sabir, T. T. Lam, M. M. Ahmed, L. Li, Y. Shen, S. E. Abo-Aba, M. I. Qureshi, M. Abu-Zeid, Y. Zhang, M. A. Khiyami, N. S. Alharbi, N. H. Hajrah, M. J. Sabir, M. H. Mutwakil, S. A. Kabli, F. A. Alsulaimany, A. Y. Obaid, B. Zhou, D. K. Smith, E. C. Holmes, H. Zhu, and Y. Guan “Co-Circulation of Three Camel Coronavirus Species and Recombination of MERS-CoVs in Saudi Arabia,” Science 351 (2016): 81–84. 10.1126/science.aac8608

7 B. Hu, L. P. Zeng, X. L. Yang, X. Y. Ge, W. Zhang, B. Li, J. Z. Xie, X. R. Shen, Y. Z. Zhang, N. Wang, D. S. Luo, X. S. Zheng, M. N. Wang, P. Daszak, L. F. Wang, J. Cui, and Z. L. Shi “Discovery of a Rich Gene Pool of Bat SARS-Related Coronaviruses Provides New Insights into the Origin of SARS Coronavirus,” PLoS Pathog. 13 (2017): e1006698. 10.1371/journal.ppat.1006698

8 World Health Organization, https://www.emro.who.int/health-topics/mers-cov/mers-outbreaks.html, 2025.

9 Q. Li, T. Shah, B. Wang, L. Qu, R. Wang, Y. Hou, Z. Baloch, and X. Xia “Cross-Species Transmission, Evolution and Zoonotic Potential of Coronaviruses,” Front. Cell. Infect. Microbiol. 12 (2022): 1081370. 10.3389/fcimb.2022.1081370

10 Y. Zhou, S. Jiang, and L. Du “Prospects for a Mers-Cov Spike Vaccine,” Expert Rev. Vaccines 17 (2018): 677–686. 10.1080/14760584.2018.1506702

11 J. Xu, W. Jia, P. Wang, S. Zhang, X. Shi, X. Wang, and L. Zhang “Antibodies and Vaccines against Middle East Respiratory Syndrome Coronavirus,” Emerg. Microbes Infect. 8 (2019): 841–856. 10.1080/22221751.2019.1624482

12 A. C. Walls, M. A. Tortorici, B. Frenz, J. Snijder, W. Li, F. A. Rey, F. DiMaio, B. J. Bosch, and D. Veesler “Glycan Shield and Epitope Masking of a Coronavirus Spike Protein Observed by Cryo-Electron Microscopy,” Nat. Struct. Mol. Biol. 23 (2016): 899–905. 10.1038/nsmb.3293

13 A. C. Walls, M. A. Tortorici, B. J. Bosch, B. Frenz, P. J. M. Rottier, F. DiMaio, F. A. Rey, and D. Veesler “Cryo-Electron Microscopy Structure of a Coronavirus Spike Glycoprotein Trimer,” Nature 531 (2016): 114–117. 10.1038/nature16988

14 Y. Yuan, D. Cao, Y. Zhang, J. Ma, J. Qi, Q. Wang, G. Lu, Y. Wu, J. Yan, Y. Shi, X. Zhang, and G. F. Gao “Cryo-EM Structures of MERS-CoV and SARS-CoV Spike Glycoproteins Reveal the Dynamic Receptor Binding Domains,” Nat. Commun. 8 (2017): 15092. 10.1038/ncomms15092

15 J. Shang, Y. Zheng, Y. Yang, C. Liu, Q. Geng, C. Luo, W. Zhang, and F. Li “Cryo-Em Structure of Infectious Bronchitis Coronavirus Spike Protein Reveals Structural and Functional Evolution of Coronavirus Spike Proteins,” PLoS Pathog. 14 (2018): e1007009. 10.1371/journal.ppat.1007009

16 J. Shang, Y. Zheng, Y. Yang, C. Liu, Q. Geng, W. Tai, L. Du, Y. Zhou, W. Zhang, and F. Li “Cryo-Electron Microscopy Structure of Porcine Deltacoronavirus Spike Protein in the Prefusion State,” J. Virol. 92 (2018): e01556–01517. 10.1128/JVI.01556-17

17 X. Xiong, M. A. Tortorici, J. Snijder, C. Yoshioka, A. C. Walls, W. Li, A. T. McGuire, F. A. Rey, B. J. Bosch, and D. Veesler “Glycan Shield and Fusion Activation of a Deltacoronavirus Spike Glycoprotein Fine-Tuned for Enteric Infections,” J. Virol. 92 (2018): e01628–01617. 10.1128/JVI.01628-17

18 W. Li, R. J. G. Hulswit, I. Widjaja, V. S. Raj, R. McBride, W. Peng, W. Widagdo, M. A. Tortorici, B. van Dieren, Y. Lang, J. W. M. van Lent, J. C. Paulson, C. A. M. de Haan, R. J. de Groot, F. J. M. van Kuppeveld, B. L. Haagmans, and B. J. Bosch “Identification of Sialic Acid-Binding Function for the Middle East Respiratory Syndrome Coronavirus Spike Glycoprotein,” Proc. Natl. Acad. Sci. U. S. A. 114 (2017): E8508–E8517. 10.1073/pnas.1712592114

19 M. A. Tortorici, A. C. Walls, Y. Lang, C. Wang, Z. Li, D. Koerhuis, G. J. Boons, B. J. Bosch, F. A. Rey, R. J. de Groot, and D. Veesler “Structural Basis for Human Coronavirus Attachment to Sialic Acid Receptors,” Nat. Struct. Mol. Biol. 26 (2019): 481–489. 10.1038/s41594-019-0233-y

20 Y. J. Park, A. C. Walls, Z. Wang, M. M. Sauer, W. Li, M. A. Tortorici, B. J. Bosch, F. DiMaio, and D. Veesler “Structures of MERS-CoV Spike Glycoprotein in Complex with Sialoside Attachment Receptors,” Nat. Struct. Mol. Biol. 26 (2019): 1151–1157. 10.1038/s41594-019-0334-7

21 M. R. Pratt, and C. R. Bertozzi “Syntheses of 6-Sulfo Sialyl Lewis X Glycans Corresponding to the L-Selectin Ligand “Sulfoadhesin”,” Org. Lett. 6 (2004): 2345–2348. 10.1021/ol0493195

22 A. Gambaryan, A. Tuzikov, G. Pazynina, N. Bovin, A. Balish, and A. Klimov “Evolution of the Receptor Binding Phenotype of Influenza A (H5) Viruses,” Virology 344 (2006): 432–438. 10.1016/j.virol.2005.08.035

23 A. S. Gambaryan, A. B. Tuzikov, G. V. Pazynina, J. A. Desheva, N. V. Bovin, M. N. Matrosovich, and A. I. Klimov “6-Sulfo Sialyl Lewis X Is the Common Receptor Determinant Recognized by H5, H6, H7 and H9 Influenza Viruses of Terrestrial Poultry,” Virol. J. 5 (2008): 85. 10.1186/1743-422X-5-85

24 K. Miyazaki, K. Sakuma, Y. I. Kawamura, M. Izawa, K. Ohmori, M. Mitsuki, T. Yamaji, Y. Hashimoto, A. Suzuki, Y. Saito, T. Dohi, and R. Kannagi “Colonic Epithelial Cells Express Specific Ligands for Mucosal Macrophage Immunosuppressive Receptors Siglec-7 and -9,” J. Immunol. 188 (2012): 4690–4700. 10.4049/jimmunol.1100605

25 M. A. Stanczak, S. S. Siddiqui, M. P. Trefny, D. S. Thommen, K. F. Boligan, S. von Gunten, A. Tzankov, L. Tietze, D. Lardinois, V. Heinzelmann-Schwarz, M. von Bergwelt-Baildon, W. Zhang, H. J. Lenz, Y. Han, C. I. Amos, M. Syedbasha, A. Egli, F. Stenner, D. E. Speiser, A. Varki, A. Zippelius, and H. Laubli “Self-Associated Molecular Patterns Mediate Cancer Immune Evasion by Engaging Siglecs on T Cells,” J. Clin. Invest. 128 (2018): 4912–4923. 10.1172/JCI120612

26 S. Duan, and J. C. Paulson “Siglecs as Immune Cell Checkpoints in Disease,” Annu. Rev. Immunol. 38 (2020): 365–395. 10.1146/annurev-immunol-102419-035900

27 J. Carpenter, and M. Kesimer “Membrane-Bound Mucins of the Airway Mucosal Surfaces Are Densely Decorated with Keratan Sulfate: Revisiting Their Role in the Lung’s Innate Defense,” Glycobiology 31 (2021): 436–443. 10.1093/glycob/cwaa089

28 M. Ohmae, Y. Yamazaki, K. Sezukuri, and J. Takada “Keratan Sulfate, a “Unique” Sulfo-Sugar: Structures, Functions, and Synthesis,” Trends Glycosci. Glycotechnol. 31 (2019): E129–E136. 10.4052/tigg.1830.1E

29 S. Komba, C. Galustian, H. Ishida, T. Feizi, R. Kannagi, and M. Kiso “The First Total Synthesis of 6-Sulfo-De-*N-*Acetylsialyl Lewis(X) Ganglioside: A Superior Ligand for Human L-Selectin,” Angew. Chem. Int. Ed. 38 (1999): 1131–1133. 10.1002/(SICI)1521-3773(19990419)38:8<1131::AID-ANIE1131>3.0.CO;2-B

30 A. K. Misra, Y. Ding, J. B. Lowe, and O. Hindsgaul “A Concise Synthesis of the 6-*O*- and 6’-*O*-Sulfated Analogues of the Sialyl Lewis X Tetrasaccharide,” Bioorg. Med. Chem. Lett. 10 (2000): 1505–1509. 10.1016/s0960-894x(00)00207-9

31 A. Santra, H. Yu, N. Tasnima, M. M. Muthana, Y. Li, J. Zeng, N. J. Kenyond, A. Y. Louie, and X. Chen “Systematic Chemoenzymatic Synthesis of *O*-Sulfated Sialyl Lewis X Antigens,” Chem. Sci. 7 (2016): 2827–2831. 10.1039/C5SC04104J

32 S. Ma, J. Zhang, F. Wei, X. Tian, Y. Tian, and L. Wen “De Novo Chemoenzymatic Assembly of Complex Sulfated *N*-Glycans to Comprehensively Profile the Ligand Binding of Human Siglecs,” J. Am. Chem. Soc. 147 (2025): 35042–35054. 10.1021/jacs.5c11949

33 H. Kawashima, B. Petryniak, N. Hiraoka, J. Mitoma, V. Huckaby, J. Nakayama, K. Uchimura, K. Kadomatsu, T. Muramatsu, J. B. Lowe, and M. Fukuda “*N*-Acetylglucosamine-6-*O*-Sulfotransferases 1 and 2 Cooperatively Control Lymphocyte Homing through L-Selectin Ligand Biosynthesis in High Endothelial Venules,” Nat. Immunol. 6 (2005): 1096–1104. 10.1038/ni1259

34 K. Uchimura, J. M. Gauguet, M. S. Singer, D. Tsay, R. Kannagi, T. Muramatsu, U. H. von Andrian, and S. D. Rosen “A Major Class of L-Selectin Ligands Is Eliminated in Mice Deficient in Two Sulfotransferases Expressed in High Endothelial Venules,” Nat. Immunol. 6 (2005): 1105–1113. 10.1038/ni1258

35 I. A. Gagarinov, T. Li, N. Wei, J. Sastre Toraño, R. P. de Vries, M. A. Wolfert, and G. J. Boons “Protecting-Group-Controlled Enzymatic Glycosylation of Oligo-*N*- Acetyllactosamine Derivatives,” Angew. Chem. Int. Ed. 58 (2019): 10547–10552. 10.1002/anie.201903140

36 H. K. Tseng, H. K. Wang, C. Y. Wu, C. K. Ni, and C. C. Lin “Exploring Regioselective Fucosylation Catalyzed by Bacterial Glycosyltransferases through Substrate Promiscuity and Acceptor-Mediated Glycosylation,” ACS Catal. 13 (2023): 10661–10671. 10.1021/acscatal.3c01563

37 G. M. Vos, PhD thesis, Utrecht University (Utrecht University Repository), 2023.

38 Y. Wu, G. M. Vos, C. Huang, D. Chapla, A. L. M. Kimpel, K. W. Moremen, R. P. de Vries, and G. J. Boons “Exploiting Substrate Specificities of 6-*O*-Sulfotransferases to Enzymatically Synthesize Keratan Sulfate Oligosaccharides,” JACS Au 3 (2023): 3155–3164. 10.1021/jacsau.3c00488

39 Y. Wu, G. P. Bosman, D. Chapla, C. Huang, K. W. Moremen, R. P. de Vries, and G. J. Boons “A Biomimetic Synthetic Strategy Can Provide Keratan Sulfate I and II Oligosaccharides with Diverse Fucosylation and Sulfation Patterns,” J. Am. Chem. Soc. 146 (2024): 9230–9240. 10.1021/jacs.4c00363

40 H. Yu, S. Huang, H. Chokhawala, M. C. Sun, H. Zheng, and X. Chen “Highly Efficient Chemoenzymatic Synthesis of Naturally Occurring and Non-Natural alpha-2,6-Linked Sialosides: A *P. Damsela* alpha-2,6-Sialyltransferase with Extremely Flexible Donor-Substrate Specificity,” Angew. Chem. Int. Ed. 45 (2006): 3938–3944. 10.1002/anie.200600572

41 J. Ye, H. Xia, N. Sun, C. C. Liu, A. Sheng, L. Chi, X. W. Liu, G. Gu, S. Q. Wang, J. Zhao, P. Wang, M. Xiao, F. Wang, and H. Cao “Reprogramming the Enzymatic Assembly Line for Site-Specific Fucosylation,” Nat. Catal. 2 (2019): 514–522. 10.1038/s41929-019-0281-z

42 J. B. McArthur, H. Yu, N. Tasnima, C. M. Lee, A. J. Fisher, and X. Chen “Alpha2-6- Neosialidase: A Sialyltransferase Mutant as a Sialyl Linkage-Specific Sialidase,” ACS Chem. Biol. 13 (2018): 1228–1234. 10.1021/acschembio.8b00002

43 C. P. Chiu, L. L. Lairson, M. Gilbert, W. W. Wakarchuk, S. G. Withers, and N. C. Strynadka “Structural Analysis of the alpha-2,3-Sialyltransferase Cst-I from *Campylobacter jejuni* in Apo and Substrate-Analogue Bound Forms,” Biochemistry 46 (2007): 7196–7204. 10.1021/bi602543d

44 V. Thon, Y. Li, H. Yu, K. Lau, and X. Chen “PmST3 from *Pasteurella Multocida* Encoded by Pm1174 Gene Is a Monofunctional α2–3-Sialyltransferase,” Appl. Microbiol. Biotechnol. 94 (2011): 977–985. 10.1007/s00253-011-3676-6

45 G. Sugiarto, K. Lau, J. Qu, Y. Li, S. Lim, S. Mu, J. B. Ames, A. J. Fisher, and X. Chen “A Sialyltransferase Mutant with Decreased Donor Hydrolysis and Reduced Sialidase Activities for Directly Sialylating Lewis^x^,” ACS Chem. Biol. 7 (2012): 1232–1240. 10.1021/cb300125k

46 F. Wei, L. Zang, P. Zhang, J. Zhang, and L. Wen “Concise Chemoenzymatic Synthesis of *N*-Glycans,” Chem 10 (2024): 2844–2860. 10.1016/j.chempr.2024.05.006

47 K. Schmolzer, D. Ribitsch, T. Czabany, C. Luley-Goedl, D. Kokot, A. Lyskowski, S. Zitzenbacher, H. Schwab, and B. Nidetzky “Characterization of a Multifunctional Alpha2,3-Sialyltransferase from *Pasteurella Dagmatis*,” Glycobiology 23 (2013): 1293–1304. 10.1093/glycob/cwt066

48 P. Maly, A. Thall, B. Petryniak, C. E. Rogers, P. L. Smith, R. M. Marks, R. J. Kelly, K. M. Gersten, G. Cheng, T. L. Saunders, S. A. Camper, R. T. Camphausen, F. X. Sullivan, Y. Isogai, O. Hindsgaul, U. H. von Andrian, and J. B. Lowe “The alpha(1,3)Fucosyltransferase Fuc-TVII Controls Leukocyte Trafficking through an Essential Role in L-, E-, and P-Selectin Ligand Biosynthesis,” Cell 86 (1996): 643–653. 10.1016/s0092-8674(00)80137-3

49 K. G. Bowman, B. N. Cook, C. L. de Graffenried, and C. R. Bertozzi “Biosynthesis of L-Selectin Ligands: Sulfation of Sialyl Lewis X-Related Oligosaccharides by a Family of GlcNAc-6-Sulfotransferases,” Biochemistry 40 (2001): 5382–5391. 10.1021/bi001750o

50 J. W. Homeister, A. D. Thall, B. Petryniak, P. Malý, C. E. Rogers, P. L. Smith, R. J. Kelly, K. M. Gersten, S. W. Askari, G. Cheng, G. Smithson, R. M. Marks, A. K. Misra, O. Hindsgaul, U. H. von Andrian, and J. B. Lowe “The α(1,3)Fucosyltransferases FucT-IV and FucT-VII Exert Collaborative Control over Selectin-Dependent Leukocyte Recruitment and Lymphocyte Homing,” Immunity 15 (2001): 115–126. 10.1016/s1074-7613(01)00166-2

51 M. Trinchera, A. Aronica, and F. Dall’Olio “Selectin Ligands Sialyl-Lewis a and Sialyl-Lewis X in Gastrointestinal Cancers,” Biology 6 (2017): 16. 10.3390/biology6010016

52 S. Nishihara, H. Iwasaki, M. Kaneko, A. Tawada, M. Ito, and H. Narimatsu “Alpha1,3- Fucosyltransferase 9 (FUT9; Fuc-TIX) Preferentially Fucosylates the Distal GlcNAc Residue of Polylactosamine Chain While the Other Four Alpha1,3FUT Members Preferentially Fucosylate the Inner GlcNAc Residue,” FEBS Lett 462 (1999): 289–294. 10.1016/s0014-5793(99)01549-5

53 S. Toivonen, S. Nishihara, H. Narimatsu, O. Renkonen, and R. Renkonen “FUC-Tix: A Versatile Alpha1,3-Fucosyltransferase with a Distinct Acceptor- and Site-Specificity Profile,” Glycobiology 12 (2002): 361–368. 10.1093/glycob/12.6.361

54 H. Yu, A. Gonzalez-Gil, Y. Wei, S. M. Fernandes, R. N. Porell, K. Vajn, J. C. Paulson, C. M. Nycholat, and R. L. Schnaar “Siglec-8 and Siglec-9 Binding Specificities and Endogenous Airway Ligand Distributions and Properties,” Glycobiology 27 (2017): 657–668. 10.1093/glycob/cwx026

55 A. Gonzalez-Gil, and R. L. Schnaar “Siglec Ligands,” Cells 10 (2021): 1260. 10.3390/cells10051260

56 L. L. Kiessling, J. E. Gestwicki, and L. E. Strong “Synthetic Multivalent Ligands in the Exploration of Cell-Surface Interactions,” Curr. Opin. Chem. Biol. 4 (2000): 696–703. 10.1016/s1367-5931(00)00153-8

57 Y. Kim, J. Y. Hyun, and I. Shin “Multivalent Glycans for Biological and Biomedical Applications,” Chem. Soc. Rev. 50 (2021): 10567–10593. 10.1039/d0cs01606c

58 I. Tomris, A. L. M. Kimpel, R. Liang, R. van der Woude, G. P. H. Boons, Z. Li, and R. P. de Vries “The HcoV-HKU1 N-Terminal Domain Binds a Wide Range of 9-*O*- Acetylated Sialic Acids Presented on Different Glycan Cores,” ACS Infect. Dis. 10 (2024): 3880–3890. 10.1021/acsinfecdis.4c00488

59 B. E. Collins, and J. C. Paulson “Cell Surface Biology Mediated by Low Affinity Multivalent Protein-Glycan Interactions,” Curr. Opin. Chem. Biol. 8 (2004): 617–625. 10.1016/j.cbpa.2004.10.004

60 J. C. Paulson, O. Blixt, and B. E. Collins “Sweet Spots in Functional Glycomics,” Nat. Chem. Biol. 2 (2006): 238–248. 10.1038/nchembio785

61 A. Imberty, Y. M. Chabre, and R. Roy “Glycomimetics and Glycodendrimers as High Affinity Microbial Anti-Adhesins,” Chem. Eur. J. 14 (2008): 7490–7499. 10.1002/chem.200800700

62 A. Bernardi, J. Jimenez-Barbero, A. Casnati, C. De Castro, T. Darbre, F. Fieschi, J. Finne, H. Funken, K. E. Jaeger, M. Lahmann, T. K. Lindhorst, M. Marradi, P. Messner, A. Molinaro, P. V. Murphy, C. Nativi, S. Oscarson, S. Penades, F. Peri, R. J. Pieters, O. Renaudet, J. L. Reymond, B. Richichi, J. Rojo, F. Sansone, C. Schaffer, W. B. Turnbull, T. Velasco-Torrijos, S. Vidal, S. Vincent, T. Wennekes, H. Zuilhof, and A. Imberty “Multivalent Glycoconjugates as Anti-Pathogenic Agents,” Chem. Soc. Rev. 42 (2013): 4709–4727. 10.1039/c2cs35408j

63 S. Bhatia, M. Dimde, and R. Haag “Multivalent Glycoconjugates as Vaccines and Potential Drug Candidates,” Med. Chem. Commun. 5 (2014): 862–878. 10.1039/c4md00143e

64 J. Dommerholt, S. Schmidt, R. Temming, L. J. Hendriks, F. P. Rutjes, J. C. van Hest, D. J. Lefeber, P. Friedl, and F. L. van Delft “Readily Accessible Bicyclononynes for Bioorthogonal Labeling and Three-Dimensional Imaging of Living Cells,” Angew. Chem. Int. Ed. 49 (2010): 9422–9425. 10.1002/anie.201003761

65 M. Ogata, S. Umemura, N. Sugiyama, N. Kuwano, A. Koizumi, T. Sawada, M. Yanase, T. Takaha, J. I. Kadokawa, and T. Usui “Synthesis of Multivalent Sialyllactosamine-Carrying Glyco-Nanoparticles with High Affinity to the Human Influenza Virus Hemagglutinin,” Carbohydr. Polym. 153 (2016): 96–104. 10.1016/j.carbpol.2016.07.083

66 D. Lauster, M. Glanz, M. Bardua, K. Ludwig, M. Hellmund, U. Hoffmann, A. Hamann, C. Bottcher, R. Haag, C. P. R. Hackenberger, and A. Herrmann “Multivalent Peptide-Nanoparticle Conjugates for Influenza-Virus Inhibition,” Angew. Chem. Int. Ed. 56 (2017): 5931–5936. 10.1002/anie.201702005

67 K. Farabi, Y. Manabe, H. Ichikawa, S. Miyake, M. Tsutsui, K. Kabayama, T. Yamaji, K. Tanaka, S. C. Hung, and K. Fukase “Concise and Reliable Syntheses of Glycodendrimers Via Self-Activating Click Chemistry: A Robust Strategy for Mimicking Multivalent Glycan-Pathogen Interactions,” J. Org. Chem. 85 (2020): 16014–16023. 10.1021/acs.joc.0c01547

68 M. N. Stadtmueller, S. Bhatia, P. Kiran, M. Hilsch, V. Reiter-Scherer, L. Adam, B. Parshad, M. Budt, S. Klenk, K. Sellrie, D. Lauster, P. H. Seeberger, C. P. R. Hackenberger, A. Herrmann, R. Haag, and T. Wolff “Evaluation of Multivalent Sialylated Polyglycerols for Resistance Induction in and Broad Antiviral Activity against Influenza a Viruses,” J. Med. Chem. 64 (2021): 12774–12789. 10.1021/acs.jmedchem.1c00794

69 L. Coudeville, F. Bailleux, B. Riche, F. Megas, P. Andre, and R. Ecochard “Relationship between Haemagglutination-Inhibiting Antibody Titres and Clinical Protection against Influenza: Development and Application of a Bayesian Random-Effects Model,” BMC Med. Res. Methodol. 10 (2010): 18. 10.1186/1471-2288-10-18

70 L. Unione, A. N. A. Ammerlaan, G. P. Bosman, E. Uslu, R. Liang, F. Broszeit, R. van der Woude, Y. Liu, S. Ma, L. Liu, M. Gomez-Redondo, I. A. Bermejo, P. Valverde, T. Diercks, A. Arda, R. P. de Vries, and G. J. Boons “Probing Altered Receptor Specificities of Antigenically Drifting Human H3N2 Viruses by Chemoenzymatic Synthesis, NMR, and Modeling,” Nat. Commun. 15 (2024): 2979. 10.1038/s41467-024-47344-y

71 A. Gimeno, S. Delgado, P. Valverde, S. Bertuzzi, M. A. Berbis, J. Echavarren, A. Lacetera, S. Martin-Santamaria, A. Surolia, F. J. Canada, J. Jimenez-Barbero, and A. Arda “Minimizing the Entropy Penalty for Ligand Binding: Lessons from the Molecular Recognition of the Histo Blood-Group Antigens by Human Galectin-3,” Angew. Chem. Int. Ed. 58 (2019): 7268–7272. 10.1002/anie.201900723

72 M. Zierke, M. Smiesko, S. Rabbani, T. Aeschbacher, B. Cutting, F. H. Allain, M. Schubert, and B. Ernst “Stabilization of Branched Oligosaccharides: Lewis(X) Benefits from a Nonconventional C-H…O Hydrogen Bond,” J. Am. Chem. Soc. 135 (2013): 13464–13472. 10.1021/ja4054702

73 M. D. Battistel, H. F. Azurmendi, M. Frank, and D. I. Freedberg “Uncovering Nonconventional and Conventional Hydrogen Bonds in Oligosaccharides through NMR Experiments and Molecular Modeling: Application to Sialyl Lewis-X,” J. Am. Chem. Soc. 137 (2015): 13444–13447. 10.1021/jacs.5b03824

74 T. Aeschbacher, M. Zierke, M. Smiesko, M. Collot, J. M. Mallet, B. Ernst, F. H. Allain, and M. Schubert “A Secondary Structural Element in a Wide Range of Fucosylated Glycoepitopes,” Chem. Eur. J. 23 (2017): 11598–11610. 10.1002/chem.201701866

75 G. Fittolani, T. Tyrikos-Ergas, A. Poveda, Y. Yu, N. Yadav, P. H. Seeberger, J. Jimenez-Barbero, and M. Delbianco “Synthesis of a Glycan Hairpin,” Nat. Chem. 15 (2023): 1461–1469. 10.1038/s41557-023-01255-5

76 J. Kwon, A. Ruda, H. F. Azurmendi, J. Zarb, M. D. Battistel, L. Liao, A. Asnani, F. I. Auzanneau, G. Widmalm, and D. I. Freedberg “Glycan Stability and Flexibility: Thermodynamic and Kinetic Characterization of Nonconventional Hydrogen Bonding in Lewis Antigens,” J. Am. Chem. Soc. 145 (2023): 10022–10034. 10.1021/jacs.2c13104

77 K. Hollingsworth, A. Di Maio, S. J. Richards, J. B. Vendeville, D. E. Wheatley, C. E. Council, T. Keenan, H. Ledru, H. Chidwick, K. Huang, F. Parmeggiani, A. Marchesi, W. Chai, R. McBerney, T. P. Kaminski, M. R. Balmforth, A. Tamasanu, J. D. Finnigan, C. Young, S. L. Warriner, M. E. Webb, M. A. Fascione, S. Flitsch, M. C. Galan, T. Feizi, M. I. Gibson, Y. Liu, W. B. Turnbull, and B. Linclau “Synthesis and Screening of a Library of Lewis(X) Deoxyfluoro-Analogues Reveals Differential Recognition by Glycan-Binding Partners,” Nat. Commun. 15 (2024): 7925. 10.1038/s41467-024-51081-7

78 J. Yang, Y. Song, W. Jin, K. Xia, G. C. Burnett, W. Qiao, J. T. Bates, V. H. Pomin, C. Wang, M. Qiao, R. J. Linhardt, J. S. Dordick, and F. Zhang “Sulfated Glycans Inhibit the Interaction of MERS-CoV Receptor Binding Domain with Heparin,” Viruses 16 (2024): 237. 10.3390/v16020237

