## Supplementary material for "Chemoenzymatic Synthesis of 6-Sulfo Sialyl Lewis^x^ Containing Glycans to Probe the Receptor Specificity of MERS Coronavirus": SI

**Table of Contents** page

1) Supplementary Schemes - - - - - - - - - - - - - - - - - - - - - - - - - - - - - - - - - - - - - - - - - S2

**Scheme S1.** FUTs selectivity for mono fucosylation. - - - - - - - - - - - - - - - - - - - - - - - - - - S2

**Scheme S2.** Failure of the strategy firstly introduce fucose then remove α2,6 sialic acid. - - - - - S2

**Scheme S3.** Enzymatic selectivity test of different α2,3 sialyltransferases. - - - - - - - - - - - - - - - S2

**Scheme S4.** Generalized derivatization scheme. - - - - - - - - - - - - - - - - - - - - - - - - - - - - - - - S3

2) Materials and Methods - - - - - - - - - - - - - - - - - - - - - - - - - - - - - - - - - - - - - - - - - - - S4

3) Analytical Data - - - - - - - - - - - - - - - - - - - - - - - - - - - - - - - - - - - - - - - - - - - - - - -S12

**Figure S1.** a) 600 MHz 1D $^1$H NMR of **4** and **4'**. b) $^1$H NMR spectra of **4** and **4'**. - - - - - - - - - S13

4) Experimental Procedures and Analysis- - - - - - - - - - - - - - - - - - - - - - - - - - - - - - - - - S14

5) Microarray and Nanoparticles - - - - - - - - - - - - - - - - - - - - - - - - - - - - - - - - - - - - - -S55

**Figure S2.** Probing the binding properties of Siglec-9 and MERS-NTD-Fc to 6-sulfo-SLe$^x$ epitope presented on O-Glycan backbone. - - - - - - - - - - - - - - - - - - S57

**Figure S3.** Probing different backbone length of 6-sulfo-SLex epitope binding properties of Siglec-9, MERS-NTD and HAs of Recombinant H5N1 influenza viruses. - - - - S58

6) Computational Approach - - - - - - - - - - - - - - - - - - - - - - - - - - - - - - - - - - - - - - - - S60

**Figure S4.** A full picture of structural basis of MERS-CoV S selectivity for the 6-sulfo-SLe$^x$ receptor. - - - - - - - - - - - - - - - - - - - - - - - - - - - - - - - - - - - - - - S61

**Figure S5.** Conformation of the ligand bound in the MERS-CoV S protein binding pocket. - - S61

**Figure S6.** Calculated interatomic distances between the ligand and interacting protein residues. - - - - - - - - - - - - - - - - - - - - - - - - - - - - - - - - - - - - - - - - - - - - S62

**Figure S7.** Overlay of the docking-derived structure of the MERS-CoV S protein in complex with 6-sulfo-SLe$^x$ and the crystal structure of the viral protein in complex with SLe$^x$. - - - - - - - - - - - - - - - - - - - - - - - - - - - - - - - - - - - - - - - - - - - - - S62

7) References- - - - - - - - - - - - - - - - - - - - - - - - - - - - - - - - - - - - - - - - - - - - - - - - S63

8) NMR Spectra, LC-MS Data and HPLC Traces - - - - - - - - - - - - - - - - - - - - - - - - - - - S66

### 1) Supplementary Schemes

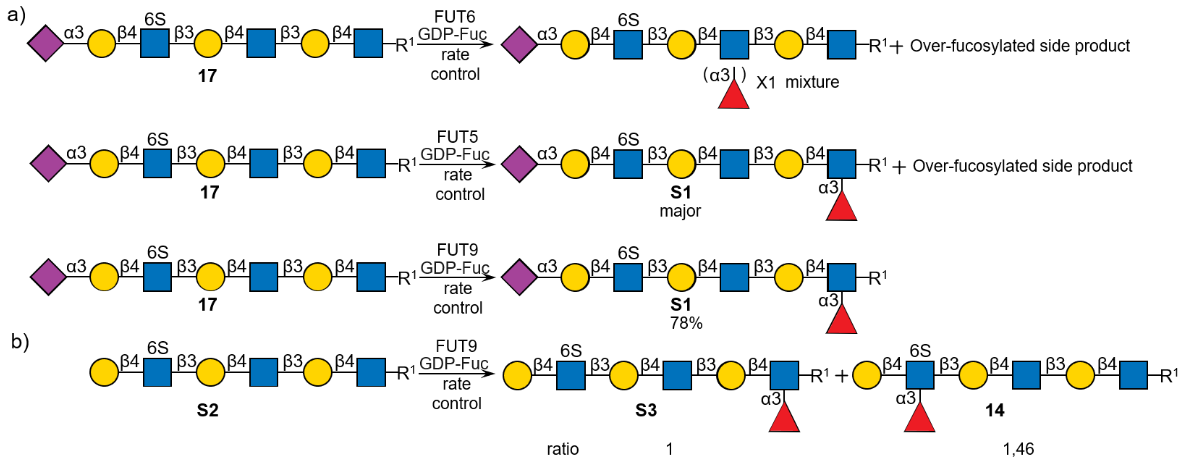

**Scheme S1.** FUTs selectivity for mono fucosylation. R[1]= $O(CH_2)_5NHCbz$.

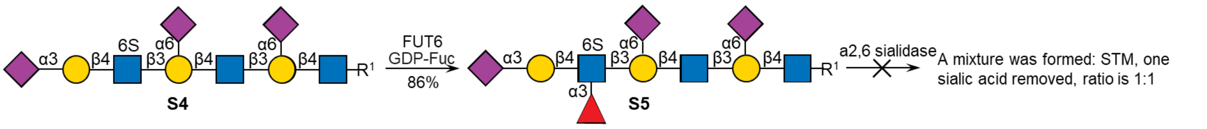

**Scheme S2.** Failure of the strategy firstly introduce fucose then remove α2,6 sialic acid. R[1]= $O(CH_2)_5NHCbz$.

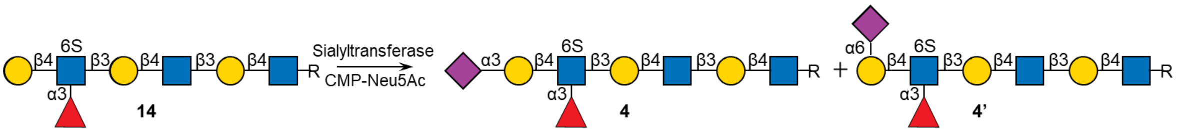

| Sialyltransferases | result |
|---|---|
| ST3Gal4 | no reaction |
| PmST3 | <10% |
| CSTI | <10% |
| PmST1 M144D | 1:1.7 Mixture of α2,3 sialic acid **4** and α2,6 sialic acid **4'** |
| Pd2,3 ST | **4** 65% **4'** trace (**4** : **4'** >10:1) |

**Scheme S3.** Enzymatic selectivity test of different α2,3 sialyltransferases. R= $O(CH_2)_5NHCbz$.

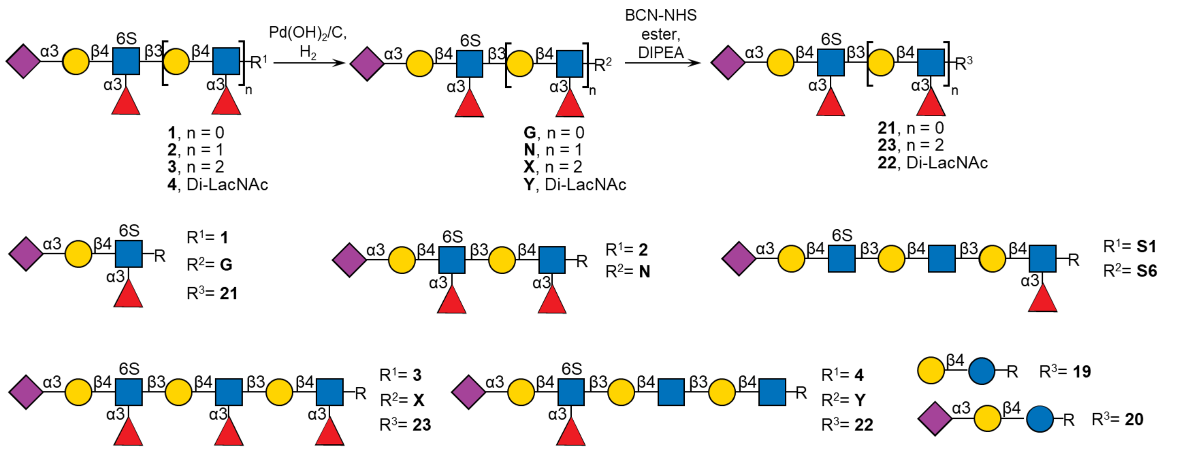

**Scheme S4.** Generalized derivatization scheme. $R^1$=O(CH$_2$)$_5$NHCbz, $R^2$=O(CH$_2$)$_5$NH$_2$, $R^3$=O(CH$_2$)$_5$NHCO$_2$CH$_2$BCN.

#### 2) Materials and Methods

Glycosyltransferases Pd2,6ST, Hpβ3GlcNAcT, Hpβ4GalT, β4GalT4, FUT5, FUT6, FUT9, PmST1 M144D, ST3Gal4, PmST3, CSTI, Pd2,3ST and α2,6 sialidase were expressed and purified according to reported protocols.[1-7] The synthesis of advanced intermediates **5**, **12**, **15**, **16**, **17**, and **S4** are described previously.[7-8] Reagents were purchased from Sigma-Aldrich. Neuraminidase (Sialidase) was obtained from Merck [Cat# 11585886001]. Uridine 5'-diphosphogalactose (UDP-Gal), uridine 5'-diphospho-N-acetyl-glucosamine (UDP-GlcNAc) and cytidine-5'-monophospho-N-acetylneuraminic acid (CMP-Neu5Ac) were obtained from Roche Diagnostics [UDP-Gal: Cat# 07703562103; UDP-GlcNAc: Cat# 06369855103; CMPNeu5Ac: Cat# 05974003103]. Adenosine 3′-phosphate 5′-phosphosulfate (PAPS) was obtained from Merck [Cat# 118410, Purity ≥ 80% by HPLC]. GDP-Fucose was prepared using L-fucokinase/GDP-fucose pyrophosphorylase.[9] UDP-GlcNTFA was prepared according to published protocol.[10] Progress of reactions was monitored by liquid chromatography mass spectrometry system (LCMS) from Shimadzu (system controller: SCL10A-VP; HPLC pumps: LC10AD-VP; injector: SIL10AD-VP) using a ZIC HILIC column (ZeQuant, PEEK coated guard HPLC column, 3.5 μm particle size, 20x 2.1 mm). The LC system was attached to a Bruker Daltonics micro TOF-Q mass spectrometer. Mass spectra were recorded on an Applied Biosystems SCIEX MALDI TOF/TOF 5800 mass spectrometer, a Shimadzu Biotech Axima-CFR MALDI-TOF, or a high-resolution Shimadzu LCMS-IT-TOF mass spectrometer. Reaction mixtures were purified using a size exclusion Biogel (P2) or Biogel (P6) resins from BioRad in Econo glass columns (0.7 x 30 cm / 1.5 x 30 cm / 1.5 x 50 cm/ 1.5 x 120 cm) coupled to a BioFrac fraction collector (BioRad). Carbohydrate-containing fractions were detected by thin layer chromatography and an appropriate staining reagent (15 mL AcOH and 3.5 mL p-Anisaldehyde in 350 mL EtOH and 50 mL $H_2SO_4$). If needed, further purification was performed by HPLC-MS using a ZIC HILIC column.

**Expression and purification of recombinant human glycosyltransferases and sulfotransferases**

Expression constructs were generated encoding the truncated catalytic domains of human glycosyltransferases (B4GALT4, FUT5, FUT6, FUT9 and ST3GAL4) and sulfotransferases CHST2 as $NH_2$-terminal fusion proteins in the pGEn2 expression vector essentially as described in prior studies.[8, 11-12] Briefly, the fusion protein coding regions were comprised of a 25-amino acid signal sequence, an $His_8$ tag, AviTag, the "superfolder" GFP coding region, the 7-amino acid recognition sequence of the tobacco etch virus (TEV) protease followed by the respective catalytic domain regions (for human CHST1 (Uniprot ID: O43916) and CHST2 (Uniprot ID: Q9Y4C5) catalytic domain region comprising of 388 and 454 amino acid residues, respectively). The recombinant human glycosyltransferases and sulfotransferases were expressed as a soluble secreted proteins by transient transfection of suspension culture HEK293-F cells (FreeStyle™ 293-F cells, Thermo Fisher Scientific, Waltham MA) and purified by $Ni^{2+}$-NTA chromatography as previously described.[11-12] Each protein was concentrated to approximately 3 mg/mL using an ultrafiltration pressure cell (Millipore, Billerica, MA) with a 10-kDa molecular mass cutoff membrane. The enzymes were further purified by gel filtration on a Superdex G-75 column (GE Healthcare) preconditioned with a buffer containing 20 mM HEPES, 150 mM NaCl, 0.05% sodium azide, pH 7.0. Peak fractions of recombinant human enzymes were pooled, respectively, concentrated at 1 mg/mL and buffer exchanged with 20 mM HEPES, 100 mM NaCl, 0.05% sodium azide, pH 7.0, 10% glycerol. The final protein preparations were aliquoted and stored at -80 °C until use.

**General protocols for enzymatic reactions**

**General procedure for the installation of β1,3 GlcNTFA using Hpβ3GlcNAcT**

Glycosyl acceptor (1.0 eq) and UDP-GlcNTFA (1.5 eq) were dissolved to a final acceptor concentration of 5 mM in a HEPES buffer (50 mM, pH 7.0) containing KCl (25 mM), $MgCl_2$ (2 mM) and DTT (1 mM) in 2 mL total volume. Calf intestine alkaline phosphotase (CIAP, 1% total volume, 1 kU/mL) and Hpβ3GlcNAcT (1% w/w relative to acceptor substrate) were added, and the reaction mixture was incubated overnight at 37 °C with gentle shaking. The progress of the reaction was monitored by LC-MS, after 18 h, another portion of Hpβ3GlcNAcT (1% w/w relative to acceptor substrate) was added until no starting material could be detected. The reaction mixture was centrifuged over a Nanosep Omega ultrafiltration device (10 kDa MWCO) to remove proteins and the filtrate was lyophilized. The residue was applicated to P2 size-exclusion column chromatography using Milli–Q water as eluent (1 mL/min), providing the desired product. High performance liquid chromatography (HPLC) using a HILIC column (see materials) was employed when impurities were detected.

**General procedure for the installation of β1,3 GlcNAc using Hpβ3GlcNAcT**

Glycosyl acceptor (1.0 eq) and UDP-GlcNAc (1.5 eq) were dissolved to provide a final acceptor concentration of 8 mM in a HEPES buffer (50 mM, pH 7.0) containing KCl (25 mM), $MgCl_2$ (2 mM) and DTT (1 mM) in 0.8 mL total volume. Calf intestine alkaline phosphotase (CIAP, 1% total volume, 1 kU/mL) and Hpβ3GlcNAcT (1% w/w relative to acceptor substrate) were added, and the reaction mixture was incubated overnight at 37 °C with gentle shaking. The progress of the reaction was monitored by LC-MS, after 18 h, another portion of Hpβ3GlcNAcT (1% w/w relative to acceptor substrate) was added until no starting material could be detected. The reaction mixture was centrifuged over a Nanosep Omega ultrafiltration device (10 kDa MWCO) to remove proteins and the filtrate was lyophilized. The residue was applicated to P6 size-exclusion column chromatography using Milli–Q water as eluent (1 mL/min), providing the desired product.

**General procedure for the installation of β1,4 Gal using B4GalT4**

Glycosyl acceptor (1.0 eq) and UDP-Gal (1.5 eq per Gal) were dissolved to provide a final acceptor concentration of 5 mM in a Tris buffer buffer (100 mM, pH 7.0) containing $MnCl_2$ (10 mM) and BSA (1% total volume) in 0.8 mL total volume. CIAP (1% volume total) and B4GalT4 (1% w/w relative to acceptor substrate) were added, and the reaction mixture was incubated 16h at 37 °C with gentle shaking. The progress of the reaction was monitored by LC-MS. The reaction mixture was centrifuged over a Nanosep Omega ultrafiltration device (10 kDa MWCO) to remove proteins and the filtrate was lyophilized. The residue was applicated to P6 size-exclusion column chromatography using Milli–Q water as eluent (1 mL/min), providing the desired product.

**General procedure for the installation of α2,3 Neu5Ac using ST3Gal4**

Glycosyl acceptor (1 eq) and CMP-Neu5Ac (1.5 eq) were dissolved in a Hepes buffer (50 mM, pH 7.2) containing BSA (1% total volume) in 0.4 mL total volume to an acceptor concentration of 10 mM. CIAP (1% volume total) and ST3Gal4 (1% w/w relative to acceptor substrate) were added, and the reaction mixture was incubated overnight at 37 °C with gentle agitation. The progress of the reaction was monitored by LC-MS, and when starting material was still detected, another portion of CMP-Neu5Ac (1.5 eq) was added and incubated overnight until no starting material was detected, which took around 36 h in total for completion. The reaction mixture was centrifuged over a Nanosep Omega ultrafiltration device (10 kDa MWCO) to remove proteins, and the filtrate was lyophilized. The residue was applicated to P6 size-exclusion column chromatography using Milli–Q water as eluent (1 mL/min), provided the desired product. High performance liquid chromatography (HPLC) using a HILIC column (see materials) was employed when the impurities were detected after size exclusion column chromatography.

**General procedure for the installation of α2,3 Neu5Ac using PmST1 M144D**

Glycosyl acceptor (1.0 eq) and CMP-Neu5Ac (1.5 eq) were dissolved at a final acceptor concentration of 6 mM in a MOPS buffer (50 mM, pH 7.2) containing BSA (1% total volume) in 0.4 mL total volume. CIAP (1% volume total) and PmST1 M144D (1% w/w relative to acceptor substrate) were added, and the reaction mixture was incubated overnight at 37 °C with gentle shaking. The progress of the reaction was monitored by LC-MS. Another portion of PmST1 M144D (1% w/w relative to acceptor substrate) and CIAP (1% volume total) and CMP-Neu5Ac (1.5 eq) was added and incubated overnight until no movement of the progress could be detected, which took around 48 h in total for completion. The reaction mixture was centrifuged over a Nanosep Omega ultrafiltration device (10 kDa MWCO) to remove proteins, and the filtrate was lyophilized. The residue was applicated to P6 size-exclusion column chromatography using $NH_4HCO_3$ buffer (50 mM) as eluent (1 mL/min), providing the desired product. High performance liquid chromatography (HPLC) using a HILIC column (see materials) was employed when the impurities were detected after size exclusion column chromatography.

**General procedure for the installation of α2,3 Neu5Ac using Pd2,3ST**

Glycosyl acceptor (1.0 eq) and CMP-Neu5Ac (10 eq) were dissolved at a final acceptor concentration of 2 mM in a MOPS buffer (50 mM, pH 7.2) containing BSA (1% total volume) in 0.6 mL total volume. CIAP (1% volume total) and Pd2,3ST (1% w/w relative to acceptor substrate) were added, and the reaction mixture was incubated overnight at 37 °C with gentle shaking. Then, another portion of CMP-Neu5Ac (10 eq) was added. The progress of the reaction was monitored by LC-MS. When there was no movement of the progress, which took around 36 h in total for completion, the reaction mixture was centrifuged over a Nanosep Omega ultrafiltration device (10 kDa MWCO) to remove proteins, and the filtrate was lyophilized. The residue was applicated to P6 size-exclusion column chromatography using $NH_4HCO_3$ buffer (50 mM) as eluent (1 mL/min), providing the desired product. High performance liquid chromatography (HPLC) using a HILIC column (see materials) was employed when the impurities were detected after size exclusion column chromatography.

**General procedure for the installation of α2,6 Neu5Ac using PT2,6ST**

Glycosyl acceptor (1.0 eq) and CMP-Neu5Ac (2.0 eq per Gal to be added) were dissolved at a final acceptor concentration of 5 mM in a MOPS buffer (50 mM, pH 7.2) containing BSA (1% volume total) in 0.4 mL total volume. CIAP (1% volume total) and PT2,6ST (1% w/w relative to acceptor substrate) were added, and the reaction mixture was incubated overnight at 37 °C with gentle shaking. The reaction mixture was centrifuged over a Nanosep Omega ultrafiltration device (10 kDa MWCO) to remove proteins, and the filtrate was lyophilized. The residue was applicated to P6 size-exclusion column chromatography using $NH_4HCO_3$ buffer (50 mM) as eluent (1 mL/min), providing the desired product. High performance liquid chromatography (HPLC) using HILIC column (see materials) was employed when the impurities were detected after size exclusion.

**General procedure for the installation of α1,3 Fuc using FUT5 in rate control**

Glycosyl acceptor (1 eq) was dissolved at a final acceptor concentration of 2 mM in a Tris buffered solution (50 mM, pH 7.3) containing $MnCl_2$ (10 mM) in 0.3 mL total volume. CIAP (1% total volume) and FUT5 (1% w/w) were added. Reaction progress was monitored by LC-MS. GDP-Fuc (0.9 eq to be added) was added in 3 portions (each portion 0.3 eq, added after GDP-Fuc could not be detected) and the reaction mixture was incubated at 37 °C with gentle shaking. After the major starting material was consumed, which took around 8 h in total for completion, the reaction mixture was centrifuged over a Nanosep Omega ultrafiltration device (10 kDa MWCO) to remove proteins, and the filtrate was lyophilized. The residue was applicated to P6 size-exclusion column chromatography using $NH_4HCO_3$ buffer (50 mM) as eluent (1 mL/min), providing the desired product. High performance liquid chromatography (HPLC) using HILIC column (see materials) was employed when the impurities were detected after size exclusion.

**General procedure for the installation of α1,3 Fuc using FUT6**

Glycosyl acceptor (1 eq) and GDP-Fuc (1.5 eq per GlcNAc to be added) were dissolved at a final acceptor concentration of 5 mM in a Tris buffered solution (50 mM, pH 7.3) containing $MnCl_2$ (10 mM). CIAP (1% total volume) and FUT6 (1% w/w) were added, and the reaction mixture was incubated overnight at 37 °C with gentle shaking. The reaction mixture was centrifuged over a Nanosep Omega ultrafiltration device (10 kDa MWCO) to remove proteins, and the filtrate was lyophilized. The residue was applicated to P2 or P6 size-exclusion column chromatography using $NH_4HCO_3$ buffer (50 mM) as eluent (1 mL/min), providing the desired product. High performance liquid chromatography (HPLC) using

HILIC column (see materials) was employed when the impurities were detected after size exclusion.

**General procedure for the installation of α1,3 Fuc using FUT6 in rate control**

Glycosyl acceptor (1 eq) was dissolved at a final acceptor concentration of 2 mM in a Tris buffered solution (50 mM, pH 7.3) containing $MnCl_2$ (10 mM) in 0.3 mL total volume. CIAP (1% total volume) and FUT6 (1% w/w) were added. Reaction progress was monitored by LC-MS. GDP-Fuc (0.9 eq to be added) was added in 3 portions (each portion 0.3 eq, added after GDP-Fuc could not be detected) and the reaction mixture was incubated at 37 °C with gentle shaking. After the major starting material was consumed, which took around 8 h in total for completion, the reaction mixture was centrifuged over a Nanosep Omega ultrafiltration device (10 kDa MWCO) to remove proteins, and the filtrate was lyophilized. The residue was applicated to P6 size-exclusion column chromatography using $NH_4HCO_3$ buffer (50 mM) as eluent (1 mL/min), providing a mixture of mono- or di- or tri-fucosylated side products which are too complicated to make further analysis.

**General procedure for the installation of α1,3 Fuc using FUT9 in rate control**

Glycosyl acceptor (1 eq) was dissolved at a final acceptor concentration of 2 mM in a Tris buffered solution (50 mM, pH 7.3) containing $MnCl_2$ (10 mM) in 0.3 mL total volume. CIAP (1% total volume) and FUT9 (1% w/w) were added. Reaction progress was monitored by LC-MS. GDP-Fuc (0.9 eq to be added) was added in 3 portions (each portion 0.3 eq, added after GDP-Fuc could not be detected) and the reaction mixture was incubated at 37 °C with gentle shaking. After the major starting material consumed, which took around 8 h in total for completion, the reaction mixture was centrifuged over a Nanosep Omega ultrafiltration device (10 kDa MWCO) to remove proteins, and the filtrate was lyophilized. The residue was applicated to P6 size-exclusion column chromatography using $NH_4HCO_3$ buffer (50 mM) as eluent (1 mL/min), providing the desired product. High performance liquid chromatography (HPLC) using a HILIC column (see materials) was employed when the impurities were detected after size exclusion.

**General procedure for the 6-O-sulfate installation of terminal GlcNAc using CHST2**

Glycosyl acceptor (1.0 eq) and PAPS (1.6 eq) were dissolved at a final acceptor concentration of 10 mM in a MOPS buffered solution (100 mM, pH 7.2) containing $MgCl_2$ (10 mM) in 0.5 mL total volume. CHST2 (10% w/w relative to acceptor substrate) was added, and the reaction mixture was incubated overnight at 37 °C with gentle shaking. The progress of the reaction was monitored by LC-MS. When there was no movement of the progress, which took around 48 h in total for completion, the reaction mixture was centrifuged over a Nanosep Omega ultrafiltration device (10 kDa MWCO) to remove proteins, and the filtrate was lyophilized. The residue was applicated to P6 size-exclusion column chromatography using Milli–Q water as eluent (1 mL/min), providing the desired product.

**General procedure for the removal of Neu5Ac using sialidase from *Clostridium perfringens* (*C. perfringens* neuraminidase)**

The sialoside (starting material) was dissolved with 0.4 mL PBS buffer (pH 6.5) in a 15 mL centrifuge tube. After the addition of appropriate amount (0.1 unit per mg substrate) of sialidase, the reaction mixture was incubated in an incubator shaker at 37 °C overnight with shaking at 300 rpm. The reaction mixture was centrifuged over a Nanosep Omega ultrafiltration device (10 kDa MWCO) to remove proteins, and the filtrate was lyophilized. The residue was applicated to P6 size-exclusion column chromatography using $NH_4HCO_3$ buffer (50 mM) as eluent (1 mL/min), providing the desired product.

**General procedure for Cbz deprotection from the linker with $Pd(OH)_2$ reduction**

Palladium hydroxide on carbon (Degussa type, 20%, 1.5 times the weight of starting material) was added to a solution of starting material in $H_2O$ (0.1% AcOH as additive) in 0.6 mL total volume. The mixture was placed under an atmosphere of hydrogen until ESI-LC-MS indicated completion of the reaction which usually takes overnight gentle shaking. The mixture was filtered through a spin filter, and the residue was washed with $H_2O$. The filtrate was lyophilized to give the final product. P6 size-exclusion column chromatography was used for purification using $NH_4HCO_3$ buffer (50 mM) as eluent (1 mL/min). Fractions containing compound were lyophilized to give the desired product.

**General procedure for synthesis of BCN linked glycans used in inhibitor experiments**

Cbz deprotected glycans were mixed with 1.1 eq of BCN-NHS (Sigma-Aldrich, Cat#: 744867, (1R,8S,9s)-Bicyclo[6.1.0]non-4-yn-9-ylmethyl N-succinimidyl carbonate, CAS: 1426827-79-3) and 2 eq of DIPEA in 200 µL DMSO. The reactions were stirred at room temperature and monitored by LC-MS. After the starting materials were consumed, the residue was applied to P6 size-exclusion column chromatography using $NH_4HCO_3$ buffer (50 mM) as eluent (1 mL/min), providing the desired product.

**General procedure SPAAC coupling to MPA polymers**

BCN-functionalized glycans **19-23** (1.5 eq) were dissolved in 200 µL DMSO. Commercially available MPA-OH **18** (1 eq, PFD-G4-TMP-AZIDE, Polymer factory Stockholm, Sweden) was added from a 10 mg/mL stock solution in DMSO. The reaction mixture was incubated in the incubation shaker at 37 °C overnight. Reaction mixture was diluted with $H_2O$ and lyophilized to dryness. Reaction mixture was redissolved in 500 µL $NH_4HCO_3$ buffer (50 mM) and loaded in a dialysis cassette with a 10 kDa cutoff. Sample was then dialysed for 24 h against $NH_4HCO_3$ buffer (50 mM), changing the buffer thrice daily. Followed by 24 h against DI water changing the buffer thrice daily. Sample was lyophilized to give the desired product **24-28**.

**Glycopolymer characterization**

Glycan loading was determined by the ratio of total weight to glycan weight, as determined by NMR using TMSP (3-(trimethylsilyl)propionic-2,2,3,3-d4 acid sodium salt, CAS: 24493-21-8) as internal standard. In short, a known quantity of glycopolymer was dissolved in MilliQ to form a concentrated stock solution. Amount of glycan incorporated in the glycopolymer was determined by adding 20% of the sample to 400 µL D2O in a NMR tube. Subsequently, a determined concentration of TMSP was added to the NMR tube until the ratio between the integration of a known peak of the glycan on the polymer and the peak of the internal standard was between 0.25 and 4. Through the ratio between the total weight of the glycopolymer and the weight of the incorporated glycan, average glycan loading on the glycopolymers was determined. The concentration of the glycopolymer stock solution was defined as the concentration glycan incorporated on the polymer as to correct for valency differences between glycopolymers. All samples were diluted with MilliQ to 0.1 mM based on incorporated glycan concentration for use in biological assays.

**General protocols for HILIC-HPLC purification**

Semi-preparative HILIC-HPLC was applied on a Shimadzu (LC-20AT, SIL-20A, CBM-20A, SPD-20A, FRC-10A) LC-ESI-IT-TOF with a XBridge HILIC column, 5 µm, 10 x 250 mm at a flow rate of 3.6 mL/min, injection volume of 100 µL (10-20 mg/mL), with 0.2% of the flow is diverted to the ESI-MS detector using a splitter. The purification was performed using 10% 10 mM $NH_4HCO_3$ in MeCN (buffer B) and MeCN in 80% 10 mM $NH_4HCO_3$ (buffer A).

General conditions for a linear gradient were used as the eluent:

Linear glycan **1-23, S1-S6**

| Time (min) | A (%) | B (%) |
|---|---|---|
| 0 | 10 | 90 |
| 90 | 50 | 50 |

#### 3) Analytical Data

##### NMR nomenclature

NMR data was obtained at room temperature on a 600 MHz instrument from Bruker. The chemical shift δ is given in parts per million (ppm) and refers to tetramethyl silane and the residual solvent peak [$^{1}$H-NMR: δ($D_2O$) =4.79 ppm]. NMR data is given as follows: $^{1}$H-NMR: chemical shift (multiplicity, coupling constants, relative integral, functional group); $^{13}$C data are extracted from HSQC spectra and given as follows: chemical shift. Multiplicity is defined as follows: s = singlet; d = doublet; t = triplet; m= multiplet. Signals were assigned by numbering the monosaccharide units starting at the reducing end of the oligosaccharide. The assignment was perfromed by using 2D-NMR spectra (COSY, HSQC, TOCSY, NOESY). The yield/concentration of the final products was determined by NMR spectroscopy using n-propanol as an internal standard. High resolution masses were measured on an Agilent 6560 Ion Mobility Q-TOF LC-MS system.

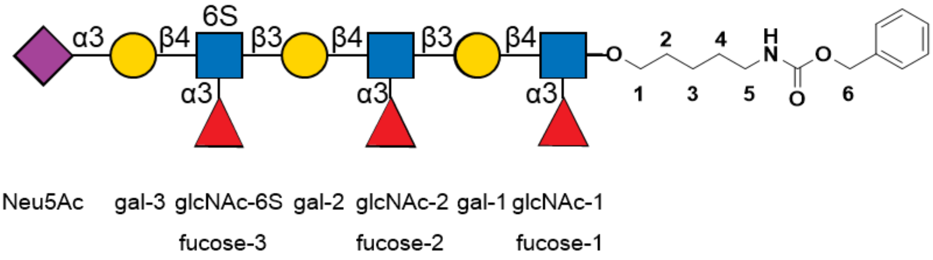

and

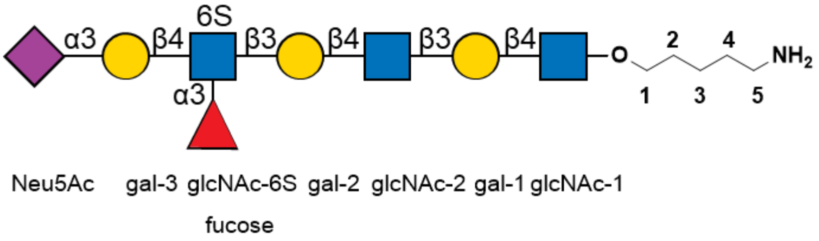

and

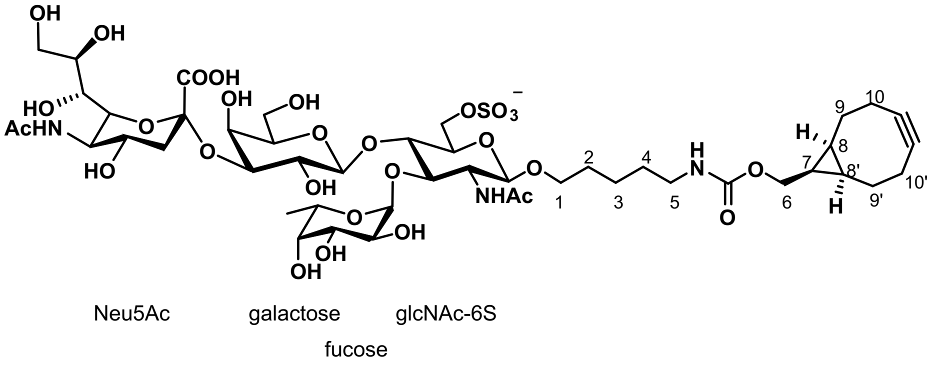

**Determination of α2,3-sialylation 4 and α2,6-sialylation 4' mixture ratio of PmST1 M144D sialylation**

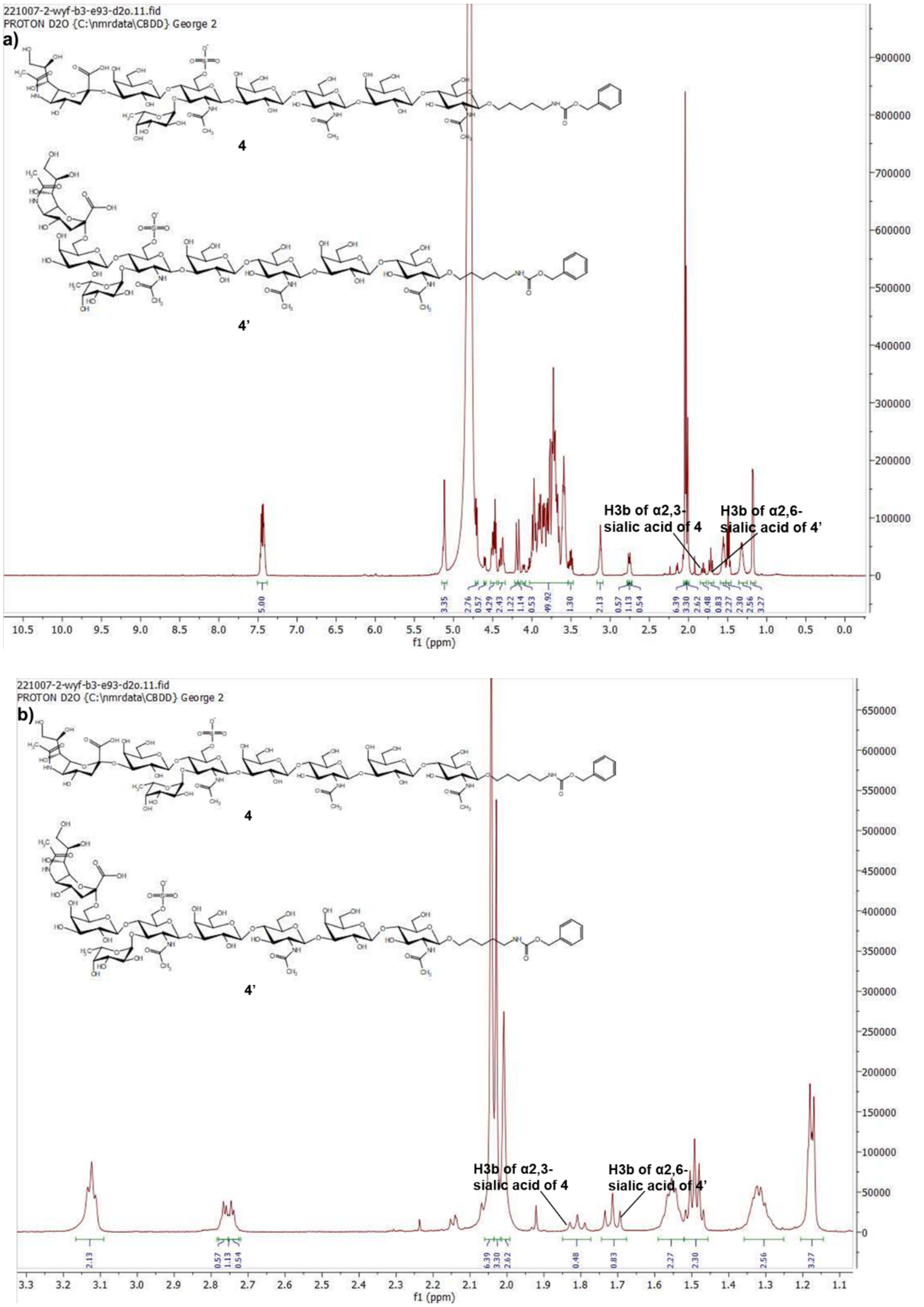

**Figure S1.** a) 600 MHz 1D $^1$H NMR of **4** and **4'**, recorded at 298K in $D_2O$. b) $^1$H NMR spectra (δ 1.1 to 3.3) of **4** and **4'**. When comparing the H3b of α2,3-Neu5Ac of **4** to H3b of α2,6-Neu5Ac of **4'**, the **4**:**4'** ratio is 1:1.7.

#### 4) Experimental Procedures and Analysis

##### Compound 15

**15** was prepared from known starting material[8] (2.0 mg, 2.9 µmol) using the general procedure for installation of α2,3Neu5Ac using ST3Gal4. After P6 purification, **15** was obtained as a white solid (2.5 mg, 88%).

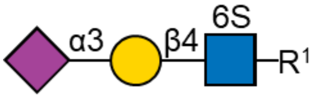

$^{1}$H (600 MHz, $D_2O$): δ (ppm)

| | H-1 | H-2 | H-3 | H-4 | H-5 | H-6 | H-7 | H-8 | H-9 | NHAc |
|---|---|---|---|---|---|---|---|---|---|---|
| GlcNAc-6S | 4.53 (d, J = 8.2 Hz, 1H) | 3.73 | n/a | 3.79 | 3.80 | 4.45 – 4.39 (m, 1H), 4.34 – 4.28 (m, 1H) | - | - | - | 2.06 – 1.98 (m, 6H) |
| Galactose | 4.61 (d, J = 7.9 Hz, 1H) | 3.57 | 4.13 (dd, J = 9.9, 3.2 Hz, 1H) | 3.98 (d, J = 3.2 Hz, 1H) | n/a | n/a | - | - | - | - |
| Neu5Ac | - | - | 2.76 (dd, J = 12.4, 4.7 Hz, 1H), 1.82 (t, J = 12.1 Hz, 1H) | 3.70 | 3.87 | n/a | n/a | 3.93 | 3.90, 3.66 | 2.06 – 1.98 (m, 6H) |

$^{13}$C (150 MHz, $D_2O$): δ (ppm)

| | C-1 | C-2 | C-3 | C-4 | C-5 | C-6 | C-7 | C-8 | C-9 | NHAc |
|---|---|---|---|---|---|---|---|---|---|---|
| GlcNAc-6S | 101.17 | 55.12 | n/a | 77.41 | 72.63 | 66.48 | - | - | - | 22.22 |
| Galactose | 102.21 | 69.48 | 75.56 | 67.65 | n/a | n/a | - | - | - | - |
| Neu5Ac | n/a | n/a | 39.68 | n/a | 51.78 | n/a | n/a | 71.61 | 62.73 | 22.22 |

| Linker | 1 | 2 | 3 | 4 | 5 | 6 |
|---|---|---|---|---|---|---|
| H | 3.87, 3.58 | 1.55 (q, J = 6.9 Hz, 2H) | 1.36 – 1.27 (m, 2H) | 1.49 (p, J = 7.3 Hz, 2H) | 3.15 – 3.09 (m, 2H) | 5.12 (s, 2H) |
| C | 70.41 | 28.29 | 22.46 | 28.47 | 40.56 | 66.93 |

HRMS (ESI-MS): m/z calculated for $C_{38}H_{58}N_3O_{24}S$ [M-H]$^-$: 972.3136; found: 972.3124.

**Compound 16**

**16** was prepared from known starting material[8] (4.0 mg, 3.8 μmol) using the general procedure for installation of α2,3Neu5Ac using ST3Gal4. After P6 purification, **16** was obtained as a white solid (4.7 mg, 92%).

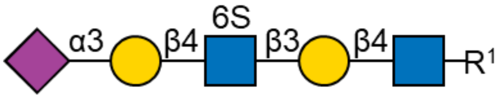

$^1$H (600 MHz, $D_2O$): δ (ppm)

| | H-1 | H-2 | H-3 | H-4 | H-5 | H-6 | H-7 | H-8 | H-9 | NHAc |
|---|---|---|---|---|---|---|---|---|---|---|
| GlcNAc | 4.51 (d, J = 7.5 Hz, 1H) | 3.72 | 3.70 | 3.69 | 3.58 | 3.99, 3.83 | - | - | - | 2.06 – 1.98 (m, 9H) |
| Galactose-1 | 4.47 (d, J = 7.8 Hz, 1H) | 3.60 | 3.74 | 4.20 (d, *J* = 3.2 Hz, 1H) | n/a | 3.76 (2H) | - | - | - | - |
| GlcNAc-6S | 4.72 (d, J = 8.5 Hz, 1H) | 3.85 | 3.74 | 3.81 | 3.82 | 4.41 (d, J = 10.9 Hz, 1H), 4.32 (d, J = 11.0 Hz, 1H) | - | - | - | 2.06 – 1.98 (m, 9H) |
| Galactose-2 | 4.61 (d, J = 7.9 Hz, 1H) | 3.58 | 4.14 (dd, J = 9.8, 3.1 Hz, 1H) | 3.98 | n/a | n/a | - | - | - | - |
| Neu5Ac | - | - | 2.76 (dd, J = 12.5, 4.6 Hz, 1H), 1.88 – 1.74 (m, 2H) | 3.69 | 3.86 | n/a | n/a | 3.92 | 3.90, 3.66 | 2.06 – 1.98 (m, 9H) |

$^{13}$C (150 MHz, $D_2O$): δ (ppm)

| | C-1 | C-2 | C-3 | C-4 | C-5 | C-6 | C-7 | C-8 | C-9 | NHAc |
|---|---|---|---|---|---|---|---|---|---|---|
| GlcNAc | 101.09 | 55.11 | 72.57 | 78.74 | 74.96 | 60.24 | - | - | - | 22.18 |
| Galactose-1 | 102.97 | 70.17 | 82.47 | 68.30 | n/a | 61.31 | - | - | - | - |
| GlcNAc- | 102.97 | 55.26 | 72.29 | 77.67 | 72.85 | 66.57 | - | - | - | 22.18 |

| | | | | | | | | | | |
|---|---|---|---|---|---|---|---|---|---|---|
| 6S | | | | | | | | | | |
| Galactose-2 | 102.30 | 69.73 | 75.56 | 67.60 | n/a | n/a | - | - | - | - |
| Neu5Ac | n/a | n/a | 39.45 | n/a | 51.96 | n/a | n/a | 71.66 | 62.64 | 22.18 |

| Linker | 1 | 2 | 3 | 4 | 5 | 6 |
|---|---|---|---|---|---|---|
| H | 3.88, 3.57 | 1.59 – 1.52 (m, 2H) | 1.36 – 1.28 (m, 2H) | 1.49 (p, J = 7.3 Hz, 2H) | 3.12 (t, J = 6.8 Hz, 2H) | 5.12 (s, 2H) |
| C | 70.41 | 28.40 | 22.37 | 28.59 | 40.43 | 66.83 |

HRMS (ESI-MS): m/z calculated for $C_{52}H_{80}N_{4}O_{34}S$ $[M-2H]^{2-}$: 668.2193; found: 668.2123.

**Compound 1**

**1** was prepared from **15** (2.0 mg, 2.0 μmol) using the general procedure for installation of α1,3Fuc using FUT6. After P6 and HILIC HPLC purification, **1** was obtained as a white solid (2.2 mg, 96%). NMR and HRMS data is reported with previous reference.[7]

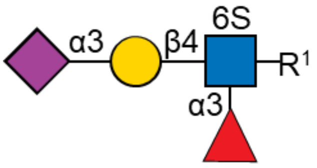

**Compound 2**

**2** was prepared from **16** (2.0 mg, 1.5 μmol) using the general procedure for installation of α1,3Fuc using FUT6. After P6 and HILIC HPLC purification, **2** was obtained as a white solid (2.3 mg, 94%). NMR and HRMS data is reported with previous reference.[7]

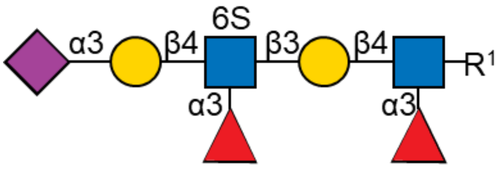

**Compound 3**

**3** was prepared from **17** (2.6 mg, 1.5 μmol) using the general procedure for installation of α1,3Fuc using FUT6. After P6 and HILIC HPLC purification, **3** was obtained as a white solid (2.2 mg, 68%). NMR and HRMS data is reported with previous reference.[7]

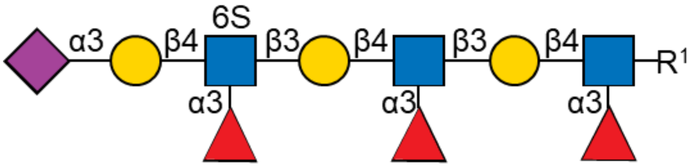

**Compound S1**

**S1** was prepared from **17** (1.0 mg, 0.6 μmol) using the installation of α1,3 Fuc using FUT9 in rate control. After purification, **S1** was obtained as a white solid (0.8 mg, 78%).

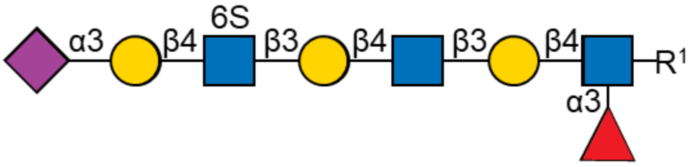

$^1$H (600 MHz, $D_2O$): δ (ppm)

| | H-1 | H-2 | H-3 | H-4 | H-5 | H-6 | H-7 | H-8 | H-9 | NHAc |
|---|---|---|---|---|---|---|---|---|---|---|
| GlcNAc-1 | 4.52 (d, J = 8.0 Hz, 1H) | 3.87 | n/a | n/a | 3.58 | 3.97, 3.84 | - | - | - | 2.05 – 1.98 (m, 12H) |
| Galactose-1 | 4.44 (d, J = 7.7 Hz, 1H) | 3.52 | 3.71 | 4.10 | n/a | 3.71 (2H) | - | - | - | - |
| GlcNAc-2 | 4.70 | 3.79 | 3.73 | 3.73 | 3.59 | 3.97, 3.84 | - | - | - | 2.05 – 1.98 (m, 12H) |
| Galactose-2 | 4.48 (d, J = 8.0 Hz, 1H) | 3.61 | 3.73 | 4.20 (d, J = 3.3 Hz, 1H) | n/a | 3.75 (2H) | - | - | - | - |
| GlcNAc-6S | 4.72 | 3.84 | 3.74 | 3.80 | 3.82 | 4.41 (d, J = 11.0 Hz, 1H), 4.34 – 4.29 (m, 1H) | - | - | - | 2.05 – 1.98 (m, 12H) |
| Galactose-3 | 4.60 (d, J = 7.8 Hz, 1H) | 3.57 | 4.13 (dd, J = 10.0, 3.2 Hz, 1H) | 3.97 (d, J = 3.6 Hz, 1H) | n/a | n/a | - | - | - | - |
| Fucose | 5.09 (d, J = 4.0 Hz, 1H) | 3.70 | n/a | n/a | 4.81 | 1.16 (d, J = 6.6 Hz, 3H) | - | - | - | - |

| | | | | | | | | | | |
|---|---|---|---|---|---|---|---|---|---|---|
| Neu5Ac | - | - | 2.76 (dd, J = 12.4, 4.7 Hz, 1H), 1.81 (t, J = 12.1 Hz, 1H) | 3.69 | 3.85 | n/a | n/a | 3.91 | 3.89, 3.65 | 2.05 – 1.98 (m, 12H) |

$^{13}C$ (150 MHz, $D_2O$): δ (ppm)

| | C-1 | C-2 | C-3 | C-4 | C-5 | C-6 | H-7 | H-8 | H-9 | NHAc |
|---|---|---|---|---|---|---|---|---|---|---|
| GlcNAc-1 | 100.85 | 55.89 | n/a | n/a | 75.14 | 59.88 | - | - | - | |
| Galactose-1 | 101.87 | 70.54 | 81.87 | 68.28 | n/a | 61.72 | - | - | - | - |
| GlcNAc-2 | 102.65 | 55.19 | 72.19 | 78.51 | 74.99 | 59.88 | - | - | - | 22.22 |
| Galactose-2 | 102.96 | 70.01 | 82.59 | 68.30 | n/a | 61.04 | - | - | - | - |
| GlcNAc-6S | 102.92 | 55.19 | 72.19 | 77.44 | 72.79 | 66.55 | - | - | - | 22.22 |
| Galactose-3 | 102.26 | 69.40 | 75.42 | 67.54 | n/a | n/a | - | - | - | - |
| Fucose | 98.72 | n/a | n/a | n/a | 66.63 | 15.30 | - | - | - | - |
| Neu5Ac | n/a | n/a | 39.66 | n/a | 51.70 | n/a | n/a | 71.64 | 62.56 | 22.22 |

| Linker | 1 | 2 | 3 | 4 | 5 | 6 |
|---|---|---|---|---|---|---|
| H | 3.87, 3.57 | 1.58 – 1.52 (m, 2H) | 1.35 – 1.27 (m, 2H) | 1.49 (p, J = 7.3 Hz, 2H) | 3.12 (t, J = 6.9 Hz, 2H) | 5.11 (s, 2H) |
| C | 70.60 | 28.40 | 22.45 | 28.51 | 40.48 | 66.87 |

HRMS (ESI-MS): m/z calculated for $C_{72}H_{113}N_5O_{48}S$ $[M-2H]^{2-}$: 923.8143; found: 923.8344.

##### Compound S5

**S5** was prepared from **S4**[7] (1.0 mg, 0.4 μmol) using the general procedure for installation of α1,3Fuc using FUT6. After P6 purification, **S5** was obtained as a white solid (0.9 mg, 86%).

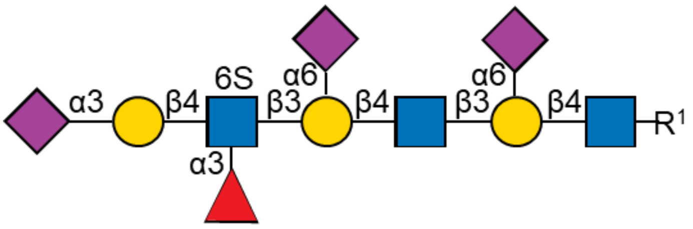

$^1H$ (600 MHz, $D_2O$): δ (ppm)

| | H-1 | H-2 | H-3 | H-4 | H-5 | H-6 | H-7 | H-8 | H-9 | NHAc |
|---|---|---|---|---|---|---|---|---|---|---|

| | | | | | | | | | | |
|---|---|---|---|---|---|---|---|---|---|---|
| GlcNAc-1 | 4.54 (d, J = 7.3 Hz, 1H) | 3.73 | n/a | n/a | 3.59 | 3.99, 3.82 | - | - | - | 2.08 – 1.99 (m, 18H) |
| Galactose-1 | 4.44 | 3.60 | 3.73 | 4.17 | n/a | 4.01, 3.56 | - | - | - | - |
| GlcNAc-2 | 4.72 (d, J = 7.9 Hz, 1H) | 3.81 | n/a | n/a | n/a | 3.97, 3.85 | - | - | - | 2.08 – 1.99 (m, 18H) |
| Galactose-2 | 4.44 | 3.60 | 3.73 | 4.17 | n/a | 4.01, 3.56 | - | - | - | - |
| GlcNAc-6S | 4.74 | 3.97 | 3.91 | 4.03 | 3.80 | 4.41 – 4.31 (m, 2H) | - | - | - | 2.08 – 1.99 (m, 18H) |
| Galactose-3 | 4.62 (d, J = 7.8 Hz, 1H) | 3.52 | 4.11 (dd, J = 9.8, 3.1 Hz, 1H) | 3.97 | n/a | n/a | - | - | - | - |
| Fucose | 5.13 | 3.69 | n/a | n/a | 4.83 | 1.18 (d, *J* = 6.4 Hz, 3H) | - | - | - | - |
| Neu5Ac-1 | - | - | 2.70 – 2.63 (m, 2H), 1.78 – 1.66 (m, 2H) | 3.68 | 3.82 | n/a | n/a | 3.91 | 3.89, 3.66 | 2.08 – 1.99 (m, 18H) |
| Neu5Ac-2 | - | - | 2.70 – 2.63 (m, 2H), 1.78 – 1.66 (m, 2H) | 3.68 | 3.82 | n/a | n/a | 3.91 | 3.89, 3.66 | 2.08 – 1.99 (m, 18H) |
| Neu5Ac-3 | - | - | 2.76 (dd, J = 12.4, 4.6 Hz, 1H), 1.82 (t, J = 12.2 Hz, 1H) | 3.70 | 3.87 | n/a | n/a | 3.91 | 3.89, 3.66 | 2.08 – 1.99 (m, 18H) |

$^{13}C$ (150 MHz, $D_2O$): δ (ppm)

| | C-1 | C-2 | C-3 | C-4 | C-5 | C-6 | C-7 | C-8 | C-9 | NHAc |
|---|---|---|---|---|---|---|---|---|---|---|
| GlcNAc-1 | 100.90 | 54.89 | n/a | n/a | 74.59 | 60.30 | - | - | - | 22.20 |
| Galactose-1 | 103.56 | 69.86 | 82.58 | 68.06 | n/a | 63.52 | - | - | - | - |
| GlcNAc-2 | 102.86 | 54.87 | n/a | n/a | n/a | 60.19 | - | - | - | 22.20 |
| Galactose-2 | 103.56 | 69.86 | 82.58 | 68.06 | n/a | 63.52 | - | - | - | - |
| GlcNAc-6S | 102.56 | 56.04 | n/a | 72.50 | 72.89 | 66.27 | - | - | - | 22.20 |
| Galactose-3 | 101.09 | 69.63 | 75.65 | 67.27 | n/a | n/a | - | - | - | - |
| Fucose | 98.58 | n/a | n/a | n/a | 66.68 | 15.30 | - | - | - | - |
| Neu5Ac-1 | n/a | n/a | 40.04 | n/a | 51.93 | n/a | n/a | 71.63 | 62.78 | 22.20 |
| Neu5Ac-2 | n/a | n/a | 40.04 | n/a | 51.93 | n/a | n/a | 71.63 | 62.78 | 22.20 |
| Neu5Ac-3 | n/a | n/a | 39.79 | n/a | 51.89 | n/a | n/a | 71.63 | 62.78 | 22.20 |

| Linker | 1 | 2 | 3 | 4 | 5 | 6 |
|---|---|---|---|---|---|---|
| H | 3.88,3.58 | 1.61 – 1.52 (m, 2H) | 1.37 – 1.24 (m, 2H) | 1.50 (p, J = 7.2 Hz, 2H) | 3.13 (t, J = 6.8 Hz, 2H) | 5.12 (s, 2H) |
| C | 70.22 | 28.28 | 22.44 | 28.49 | 40.43 | 66.77 |

HRMS (ESI-MS): m/z calculated for $C_{94}H_{146}N_7O_{64}S$ $[M-3H]^{3-}$: 809.9385; found: 809.9263.

**Compound 6**

**6** was prepared from **5**[13] (6.4 mg, 6.5 µmol) using the general procedure for installation of β1,3GlcNAc using Hpβ3GlcNAcT. After P6 purification, **6** was obtained as a white solid (6.5 mg, 84%).

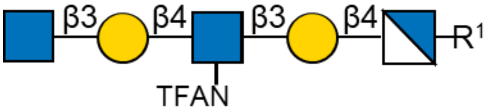

$^{1}H$ (600 MHz, $D_2O$): δ (ppm)

| | H-1 | H-2 | H-3 | H-4 | H-5 | H-6 | NHAc |
|---|---|---|---|---|---|---|---|
| $GlcNH_2$ | 4.75 (d, *J* = 8.6 Hz, 1H) | 3.04 (dd, *J* = 10.6, 8.5 Hz, 1H) | 3.87 | n/a | n/a | n/a | - |
| Galactose-1 | 4.44 (d, *J* = 7.9 | 3.60 | 3.74 | 4.19 (d, *J* = 3.3 Hz, | n/a | 3.78 (4H) | - |

| | | | | | | | |
|---|---|---|---|---|---|---|---|
| | Hz, 1H) | | | 1H) | | | |
| GlcNHTFA | 4.83 (d, *J* = 8.2 Hz, 1H) | 3.90 | n/a | n/a | n/a | n/a | - |
| Galactose-2 | 4.49 (d, *J* = 7.8 Hz, 1H) | 3.60 | 3.74 | 4.17 (d, *J* = 3.3 Hz, 1H) | n/a | 3.78 (4H) | - |
| GlcNAc | 4.69 (d, *J* = 8.5 Hz, 1H) | 3.78 | n/a | n/a | 3.46 | 3.90, 3.79 | 2.05 (s, 3H) |

$^{13}C$ (150 MHz, $D_2O$): δ (ppm)

| | C-1 | C-2 | C-3 | C-4 | C-5 | C-6 | NHAc |
|---|---|---|---|---|---|---|---|
| $GlcNH_2$ | 98.50 | 55.45 | n/a | n/a | n/a | n/a | - |
| Galactose-1 | 103.11 | 70.00 | 82.36 | 68.21 | n/a | 61.06 | - |
| GlcNHTFA | 102.01 | 55.84 | n/a | n/a | n/a | n/a | - |
| Galactose-2 | 102.96 | 70.00 | 82.36 | 68.41 | n/a | 61.06 | - |
| GlcNAc | 102.94 | 55.65 | n/a | n/a | n/a | 60.51 | 22.18 |

| Linker | 1 | 2 | 3 | 4 | 5 | 6 |
|---|---|---|---|---|---|---|
| H | 3.92, 3.68 | 1.68 – 1.60 (m, 2H) | 1.42 – 1.32 (m, 2H) | 1.57 – 1.48 (m, 2H) | 3.16 – 3.11 (m, 2H) | 5.12 (s, 2H) |
| C | 70.68 | 28.31 | 22.43 | 28.65 | 40.36 | 66.79 |

HRMS (ESI-MS): m/z calculated for $C_{47}H_{74}F_3N_4O_{27}$ $[M+H]^+$: 1183.4488; found: 1183.4637.

**Compound 7**

**7** was prepared from **6** (6.5 mg, 5.5 µmol) using the general procedure for 6-O-sulfate installation of terminal GlcNAc using CHST2. After P6 purification, **7** was obtained as a white solid (4.9 mg, 71%).

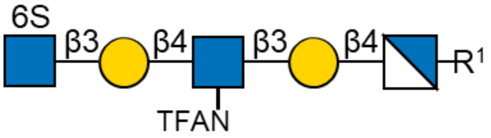

$^1H$ (600 MHz, $D_2O$): δ (ppm)

| | H-1 | H-2 | H-3 | H-4 | H-5 | H-6 | NHAc |
|---|---|---|---|---|---|---|---|
| $GlcNH_2$ | 4.71 | 3.00 | 3.82 | n/a | n/a | 3.97, | - |

| | | | | | | | |
|---|---|---|---|---|---|---|---|
| | | (t, $J$ = 9.5 Hz, 1H) | | | | 3.83 | |
| Galactose-1 | 4.44 (d, $J$ = 7.9 Hz, 1H) | 3.60 | 3.74 | 4.19 (d, $J$ = 3.3 Hz, 1H) | n/a | 3.77 (4H) | - |
| GlcNHTFA | 4.84 (d, $J$ = 8.0 Hz, 1H) | 3.91 | n/a | n/a | n/a | n/a | - |
| Galactose-2 | 4.49 (d, $J$ = 7.9 Hz, 1H) | 3.60 | 3.74 | 4.20 (d, $J$ = 3.2 Hz, 1H) | n/a | 3.77 (4H) | - |
| GlcNAc-6S | 4.70 | 3.81 | 3.59 | 3.53 | 3.68 | 4.35 (d, $J$ = 11.0 Hz, 1H), 4.23 (dd, $J$ = 11.2, 5.8 Hz, 1H) | 2.05 (s, 3H) |

$^{13}$C (150 MHz, $D_2O$): δ (ppm)

| | C-1 | C-2 | C-3 | C-4 | C-5 | C-6 | NHAc |
|---|---|---|---|---|---|---|---|
| $GlcNH_2$ | 98.99 | 55.54 | n/a | n/a | n/a | 59.90 | - |
| Galactose-1 | 103.11 | 70.00 | 82.43 | 68.34 | n/a | 61.06 | - |
| GlcNHTFA | 102.01 | 55.73 | n/a | n/a | n/a | n/a | - |
| Galactose-2 | 102.91 | 70.00 | 82.43 | 68.34 | n/a | 61.06 | - |
| GlcNAc-6S | 102.88 | 55.65 | 73.64 | 69.74 | 73.73 | 67.23 | 22.18 |

| Linker | 1 | 2 | 3 | 4 | 5 | 6 |
|---|---|---|---|---|---|---|
| H | 3.92, 3.68 | 1.68 – 1.60 (m, 2H) | 1.42 – 1.32 (m, 2H) | 1.57 – 1.48 (m, 2H) | 3.14 (t, $J$ = 6.8 Hz, 2H) | 5.12 (s, 2H) |
| C | 70.68 | 28.31 | 22.23 | 28.65 | 40.36 | 66.79 |

HRMS (ESI-MS): m/z calculated for $C_{47}H_{72}F_3N_4O_{30}S$ $[M-H]^-$: 1261.3909; found: 1261.3280.

**Compound 8**

**8** was prepared from **7** (4.9 mg, 3.9 μmol) using the general procedure for installation of β1,4Gal using B4GalT4. After P6 purification, **8** was obtained as a white solid (4.7 mg, 85%).

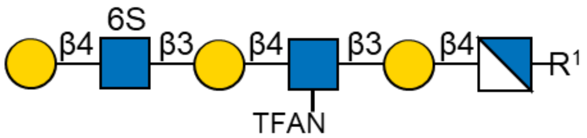

$^{1}H$ (600 MHz, $D_2O$): δ (ppm)

| | H-1 | H-2 | H-3 | H-4 | H-5 | H-6 | NHAc |
|---|---|---|---|---|---|---|---|
| $GlcNH_2$ | 4.75 | 3.04 | 3.84 | n/a | n/a | 3.98, 3.83 | - |
| Galactose-1 | 4.45 (d, *J* = 7.9 Hz, 1H) | 3.60 | 3.74 | 4.19 (d, *J* = 3.4 Hz, 1H) | n/a | 3.77 (6H) | - |
| GlcNHTFA | 4.84 (d, *J* = 7.9 Hz, 1H) | 3.90 | n/a | n/a | n/a | n/a | - |
| Galactose-2 | 4.49 (d, *J* = 7.9 Hz, 1H) | 3.61 | 3.74 | 4.20 (d, *J* = 4.0 Hz, 1H) | n/a | 3.77 (6H) | - |
| GlcNAc-6S | 4.74 | 3.85 | n/a | n/a | 3.83 | 4.41 (d, *J* = 10.8 Hz, 1H), 4.32 (dd, *J* = 11.1, 4.2 Hz, 1H) | 2.05 (s, 3H) |
| Galactose-3 | 4.53 (d, *J* = 7.8 Hz, 1H) | 3.55 | 3.70 | 3.94 | n/a | 3.77 (6H) | - |

$^{13}C$ (150 MHz, $D_2O$): δ (ppm)

| | C-1 | C-2 | C-3 | C-4 | C-5 | C-6 | NHAc |
|---|---|---|---|---|---|---|---|
| $GlcNH_2$ | 98.70 | 55.42 | n/a | n/a | n/a | 59.93 | - |
| Galactose-1 | 103.00 | 70.00 | 82.43 | 68.34 | n/a | 61.18 | - |
| GlcNHTFA | 102.17 | 55.63 | n/a | n/a | n/a | n/a | - |
| Galactose-2 | 103.00 | 70.00 | 82.43 | 68.34 | n/a | 61.18 | - |
| GlcNAc-6S | 102.81 | 55.17 | n/a | n/a | 72.52 | 66.53 | 22.18 |

| | | | | | | | |
|---|---|---|---|---|---|---|---|
| Galactose-3 | 102.69 | 71.13 | 72.54 | 68.72 | n/a | 61.18 | - |

| Linker | 1 | 2 | 3 | 4 | 5 | 6 |
|---|---|---|---|---|---|---|
| H | 3.92, 3.68 | 1.68 – 1.60 (m, 2H) | 1.42 – 1.32 (m, 2H) | 1.57 – 1.48 (m, 2H) | 3.14 (t, *J* = 6.8 Hz, 2H) | 5.12 (s, 2H) |
| C | 70.68 | 28.31 | 22.23 | 28.65 | 40.36 | 66.79 |

HRMS (ESI-MS): m/z calculated for $C_{53}H_{82}F_3N_4O_{35}S$ $[M-H]^-$: 1423.4437; found: 1423.3723.

##### Compound 9

**9** was prepared from **8** (4.7 mg, 3.3 μmol) using the general procedure for installation of α2,3Neu5Ac using ST3Gal4. After P6 purification, **9** was obtained as a white solid (4.3 mg, 77%).

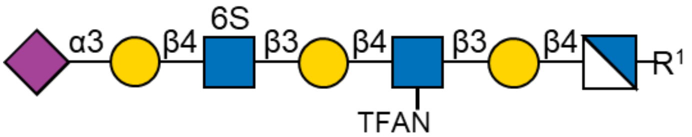

$^1$H (600 MHz, $D_2O$): δ (ppm)

| | H-1 | H-2 | H-3 | H-4 | H-5 | H-6 | H-7 | H-8 | H-9 | NHAc |
|---|---|---|---|---|---|---|---|---|---|---|
| $GlcNH_2$ | 4.40 | 2.65 (t, *J* = 9.1 Hz, 1H) | 3.53 (t, *J* = 9.4 Hz, 1H) | n/a | n/a | 3.98, 3.88 | - | - | - | - |
| Galactose-1 | 4.43 (d, *J* = 7.8 Hz, 1H) | 3.59 | 3.74 | 4.20 | n/a | 3.77 (6H) | - | - | - | - |
| GlcNHTFA | 4.84 (d, *J* = 8.6 Hz, 1H) | 3.90 | n/a | n/a | n/a | n/a | - | - | - | - |
| Galactose-2 | 4.49 (d, *J* = 7.8 Hz, 1H) | 3.60 | 3.74 | 4.21 | n/a | 3.77 (6H) | - | - | - | - |
| GlcNAc-6S | 4.72 (d, *J* = 7.5 Hz, | 3.85 | n/a | n/a | 3.83 | 4.45 – 4.42 (m, 1H), 4.33 | - | - | - | 2.06 – 2.01 (m, 6H) |

| | | | | | | | | | | |
|---|---|---|---|---|---|---|---|---|---|---|
| | 1H) | | | | | (d, *J* = 10.7 Hz, 1H) | | | | |
| Galactose-3 | 4.61 (d, *J* = 7.9 Hz, 1H) | 3.57 | 4.13 (dd, *J* = 10.4, 2.5 Hz, 1H) | 3.98 (d, *J* = 3.4 Hz, 1H) | n/a | 3.77 (6H) | - | - | - | - |
| Nue5Ac | - | - | 2.76 (dd, *J* = 12.5, 4.7 Hz, 1H), 1.81 (t, *J* = 12.2 Hz, 1H) | 3.68 | 3.86 | n/a | n/a | n/a | 3.91, 3.66 | 2.06 – 2.01 (m, 6H) |

$^{13}$C (150 MHz, $D_2O$): δ (ppm)

| | C-1 | C-2 | C-3 | C-4 | C-5 | C-6 | C-7 | C-8 | C-9 | NHAc |
|---|---|---|---|---|---|---|---|---|---|---|
| $GlcNH_2$ | 102.65 | 56.24 | 74.14 | n/a | n/a | 59.95 | - | - | - | - |
| Galactose-1 | 102.97 | 70.00 | 82.43 | 68.34 | n/a | 61.18 | - | - | - | - |
| GlcNHTFA | 102.06 | 55.63 | n/a | n/a | n/a | n/a | - | - | - | - |
| Galactose-2 | 102.85 | 70.00 | 82.43 | 68.34 | n/a | 61.18 | - | - | - | - |
| GlcNAc-6S | 102.81 | 55.17 | n/a | n/a | 72.52 | 66.74 | - | - | - | 22.18 |
| Galactose-3 | 102.26 | 69.63 | 75.41 | 67.65 | n/a | 61.18 | - | - | - | - |
| Nue5Ac | n/a | n/a | 39.62 | n/a | 51.88 | n/a | n/a | n/a | 62.73 | 22.18 |

| Linker | 1 | 2 | 3 | 4 | 5 | 6 |
|---|---|---|---|---|---|---|
| H | 3.92, 3.68 | 1.68 – 1.60 (m, 2H) | 1.42 – 1.32 (m, 2H) | 1.57 – 1.48 (m, 2H) | 3.14 (t, *J* = 6.2 Hz, 2H) | 5.12 (s, 2H) |
| C | 70.46 | 28.31 | 22.23 | 28.65 | 40.36 | 66.79 |

HRMS (ESI-MS): m/z calculated for $C_{64}H_{98}F_3N_5O_{43}S$ $[M-2H]^{2-}$: 856.7659; found: 856.7254.

**Compound 10**

The GlcNTFA moiety of **9** (4.3 mg, 2.5 μmol) was converted to $GlcNH_2$ by dissolving the substrate in $H_2O$ to a final concentration of 10 mM in 0.25 mL total volume. The pH of the solution was adjusted to 10 using 25% $NH_4OH$ aqueous solution. The reaction mixture was incubated overnight at 37 °C with gentle shaking. Progress of the reaction was monitored by LC-MS and once complete the solvent was removed by lyophilization. Then the residue was applicated to P6 size-exclusion column chromatography using $NH_4HCO_3$ buffer (50 mM) as eluent (1 mL/min), providing the desired product **10** as a white solid (3.9 mg, 97%).

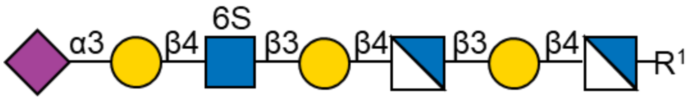

$^1$H (600 MHz, $D_2O$): δ (ppm)

| | H-1 | H-2 | H-3 | H-4 | H-5 | H-6 | H-7 | H-8 | H-9 | NHAc |
|---|---|---|---|---|---|---|---|---|---|---|
| $GlcNH_2$-1 | 4.48 (d, *J* = 7.6 Hz, 1H) | 2.76 | 3.61 | n/a | n/a | 3.98, 3.88 | - | - | - | - |
| Galactose-1 | 4.46 (d, *J* = 7.9 Hz, 1H) | 3.63 | 3.73 | 4.20 | n/a | 3.76 (6H) | - | - | - | - |
| $GlcNH_2$-2 | 4.77 | 2.92 | 3.70 | n/a | n/a | 3.98, 3.88 | - | - | - | - |
| Galactose-2 | 4.51 (d, *J* = 8.0 Hz, 1H) | 3.72 | 3.85 | 4.20 | n/a | 3.76 (6H) | - | - | - | - |
| GlcNAc-6S | 4.72 (d, *J* = 8.2 Hz, 1H) | 3.84 | n/a | n/a | 3.83 | 4.42 (d, *J* = 10.8 Hz, 1H), 4.32 (d, *J* = 11.0 Hz, 1H) | - | - | - | 2.06 – 2.01 (m, 6H) |
| Galactose-3 | 4.61 (d, *J* = 7.8 Hz, 1H) | 3.57 | 4.13 (dd, *J* = 9.7, 1.8 Hz, 1H) | 3.98 | n/a | 3.76 (6H) | - | - | - | - |
| Nue5Ac | - | - | 2.76 (dd, *J* = 12.4, 4.8 Hz, 1H), | 3.70 | 3.86 | n/a | n/a | n/a | 3.91, 3.66 | 2.06 – 2.01 (m, 6H) |

| | | | |
|---|---|---|---|
| | | | 1.81 (t, *J* = 12.1 Hz, 1H) |

$^{13}$C (150 MHz, $D_2O$): δ (ppm)

| | C-1 | C-2 | C-3 | C-4 | C-5 | C-6 | C-7 | C-8 | C-9 | NHAc |
|---|---|---|---|---|---|---|---|---|---|---|
| $GlcNH_2$-1 | 102.65 | 56.20 | n/a | n/a | n/a | 60.12 | - | - | - | - |
| Galactose-1 | 102.97 | n/a | 82.53 | 68.28 | n/a | 61.24 | - | - | - | - |
| $GlcNH_2$-2 | 102.61 | 56.20 | n/a | n/a | n/a | 60.12 | - | - | - | - |
| Galactose-2 | 102.76 | n/a | 82.23 | 68.28 | n/a | 61.24 | - | - | - | - |
| GlcNAc-6S | 102.81 | 55.27 | n/a | n/a | 72.52 | 66.48 | - | - | - | 22.18 |
| Galactose-3 | 102.26 | 69.58 | 75.49 | 67.53 | n/a | 61.24 | - | - | - | - |
| Nue5Ac | n/a | n/a | 39.62 | n/a | 51.88 | n/a | n/a | n/a | 62.53 | 22.18 |

| Linker | 1 | 2 | 3 | 4 | 5 | 6 |
|---|---|---|---|---|---|---|
| H | 3.91, 3.65 | 1.68 – 1.60 (m, 2H) | 1.42 – 1.32 (m, 2H) | 1.57 – 1.48 (m, 2H) | 3.14 (t, *J* = 6.6 Hz, 2H) | 5.12 (s, 2H) |
| C | 70.46 | 28.31 | 22.23 | 28.65 | 40.36 | 66.79 |

HRMS (ESI-MS): m/z calculated for $C_{62}H_{99}N_5O_{42}S$ $[M-2H]^{2-}$: 808.7748; found: 808.7229.

**Compound 11**

**11** was prepared from **10** (1.9 mg, 1.2 μmol) using the general procedure for installation of α1,3Fuc using FUT6. After P6 purification, **11** was obtained as a white solid (1.9 mg, 92%).

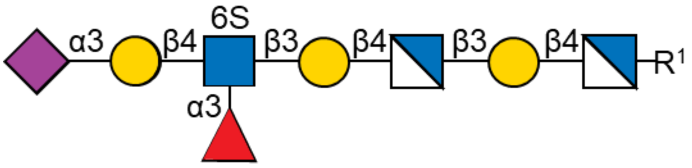

$^{1}$H (600 MHz, $D_2O$): δ (ppm)

| | H-1 | H-2 | H-3 | H-4 | H-5 | H-6 | H-7 | H-8 | H-9 | NHAc |
|---|---|---|---|---|---|---|---|---|---|---|
| $GlcNH_2$-1 | 4.61 | 2.89 | 3.74 | n/a | n/a | 3.97, 3.84 | - | - | - | - |
| Galactose-1 | 4.46 (d, *J* = 7.9 Hz, 1H) | 3.61 | 3.72 | 4.21 | n/a | 3.78 (4H) | - | - | - | - |
| $GlcNH_2$-2 | 4.91 | 3.07 | 3.82 | n/a | n/a | 3.97, 3.84 | - | - | - | - |

| | | | | | | | | | | |
|---|---|---|---|---|---|---|---|---|---|---|
| Galactose-2 | 4.52 (d, *J* = 7.9 Hz, 1H) | 3.73 | 3.88 | 4.21 | n/a | 3.78 (4H) | - | - | - | - |
| GlcNAc-6S | 4.75 (d, *J* = 8.4 Hz, 1H) | 3.99 | 3.91 | 4.03 | 3.80 | 4.39 – 4.34 (m, 2H) | - | - | - | 2.05 – 2.01 (m, 6H) |
| Galactose-3 | 4.59 (d, *J* = 7.9 Hz, 1H) | 3.52 | 4.11 (dd, *J* = 9.9, 3.2 Hz, 1H) | 3.96 | n/a | 3.71 (2H) | - | - | - | - |
| Fucose | 5.13 (d, *J* = 4.1 Hz, 1H) | 3.69 | n/a | n/a | 4.82 | 1.18 (d, J = 6.6 Hz, 3H) | - | - | - | - |
| Nue5Ac | - | - | 2.76 (dd, J = 12.4, 4.7 Hz, 1H), 1.81 (t, J = 12.2 Hz, 1H) | 3.69 | 3.86 | n/a | n/a | n/a | 3.90, 3.66 | 2.05 – 2.01 (m, 6H) |

$^{13}C$ (150 MHz, $D_2O$): δ (ppm)

| | C-1 | C-2 | C-3 | C-4 | C-5 | C-6 | H-7 | H-8 | H-9 | NHAc |
|---|---|---|---|---|---|---|---|---|---|---|
| $GlcNH_2$-1 | 101.43 | 55.41 | n/a | n/a | n/a | 60.12 | - | - | - | - |
| Galactose-1 | 103.19 | n/a | 82.54 | 68.31 | n/a | 61.26 | - | - | - | - |
| $GlcNH_2$-2 | n/a | 55.93 | n/a | n/a | n/a | 60.12 | - | - | - | - |
| Galactose-2 | 102.70 | n/a | 82.05 | 68.31 | n/a | 61.26 | - | - | - | - |
| GlcNAc-6S | 102.61 | 55.64 | n/a | 72.78 | 72.39 | 66.24 | - | - | - | 22.19 |
| Galactose-3 | 101.41 | 69.36 | 75.28 | 67.34 | n/a | 61.55 | - | - | - | - |
| Fucose | 98.40 | n/a | n/a | n/a | 66.63 | 15.36 | - | - | - | - |
| Nue5Ac | n/a | n/a | 39.80 | n/a | 51.61 | n/a | n/a | n/a | 62.79 | 22.19 |

| Linker | 1 | 2 | 3 | 4 | 5 | 6 |
|---|---|---|---|---|---|---|
| H | 3.87, 3.66 | 1.68 – 1.60 (m, 2H) | 1.42 – 1.32 (m, 2H) | 1.57 – 1.48 (m, 2H) | 3.13 (t, J = 6.8 Hz, 2H) | 5.12 (s, 2H) |
| C | 70.60 | 28.67 | 22.25 | 28.75 | 40.31 | 66.79 |

HRMS (ESI-MS): m/z calculated for $C_{68}H_{109}N_5O_{46}S$ $[M-2H]^{2-}$: 881.8037; found: 881.7367.

**Compound 4**

Amine free glycan **11** (0.7 mg, 0.37 µmol) were mixed with of AcOSu (2.4 eq, 8.9 µL, 0.89 µmol from 100 mM stock solution) and 8 eq of DIPEA (8.0 eq, 0.5 µL, 2.96 µmol) in 300 uL DMSO. The reactions were stirred at room temperature and monitored by LC-MS until full acetylation was observed. In the event starting amine was detected after overnight incubation, additional AcOSu (2 eq, 7.4 µL, 0.74 µmol from 100 mM stock solution) was added until complete conversion was observed after another 18h. After the starting materials were consumed, the reaction was lyophilized and the residue was applicated to P6 size-exclusion column chromatography using $NH_4HCO_3$ buffer (50 mM) as eluent (1 mL/min), providing the desired product **4** as a white solid (0.7 mg, 95%).

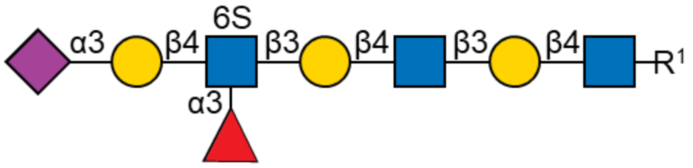

$^1$H (600 MHz, $D_2O$): δ (ppm)

| | H-1 | H-2 | H-3 | H-4 | H-5 | H-6 | H-7 | H-8 | H-9 | NHAc |
|---|---|---|---|---|---|---|---|---|---|---|
| GlcNAc-1 | 4.51 (d, J = 7.7 Hz, 1H) | 3.71 | 3.69 | 3.69 | 3.58 | 3.97, 3.84 | - | - | - | 2.06 – 1.99 (m, 12H) |
| Galactose-1 | 4.46 | 3.59 | 3.72 | 4.16 (d, *J* = 3.2 Hz, 1H) | n/a | 3.76 (4H) | - | - | - | - |
| GlcNAc-2 | 4.71 (d, J = 8.5 Hz, 1H) | 3.81 | n/a | n/a | 3.59 | 3.97, 3.84 | - | - | - | 2.06 – 1.99 (m, 12H) |
| Galactose-2 | 4.48 | 3.59 | 3.72 | 4.20 (d, *J* = 3.2 Hz, | n/a | 3.76 (4H) | - | - | - | - |

| | | | | | | | | | | |
|---|---|---|---|---|---|---|---|---|---|---|
| | | | | 1H) | | | | | | |
| GlcNAc-6S | 4.74 | 4.00 | 3.91 | 4.03 | 3.80 | 4.40 – 4.34 (m, 2H) | - | - | - | 2.06 – 1.99 (m, 12H) |
| Galactose-3 | 4.60 (d, J = 7.8 Hz, 1H) | 3.52 | 4.10 | 3.97 | n/a | 3.71 (2H) | - | - | - | - |
| Fucose | 5.13 (d, J = 4.4 Hz, 1H) | 3.69 | n/a | n/a | 4.82 | 1.18 (d, J = 6.5 Hz, 3H) | - | - | - | - |
| Neu5Ac | - | - | 2.76 (dd, J = 12.7, 4.6 Hz, 1H), 1.81 (t, J = 12.1 Hz, 1H) | 3.69 | 3.86 | n/a | n/a | n/a | 3.90, 3.66 | 2.06 – 1.99 (m, 12H) |

$^{13}C$ (150 MHz, $D_2O$): δ (ppm)

| | C-1 | C-2 | C-3 | C-4 | C-5 | C-6 | H-7 | H-8 | H-9 | NHAc |
|---|---|---|---|---|---|---|---|---|---|---|
| GlcNAc-1 | 101.09 | 55.16 | 72.59 | 78.46 | 75.14 | 60.12 | - | - | - | 22.29 |
| Galactose-1 | 102.91 | 69.66 | 82.31 | 68.31 | n/a | 61.32 | - | - | - | - |
| GlcNAc-2 | 102.71 | 55.14 | n/a | n/a | 74.99 | 60.12 | - | - | - | 22.29 |
| Galactose-2 | 102.96 | 69.66 | 82.31 | 68.31 | n/a | 61.32 | - | - | - | - |
| GlcNAc-6S | 102.61 | 55.92 | n/a | 72.91 | 72.39 | 66.09 | - | - | - | 22.29 |
| Galactose-3 | 101.34 | 69.73 | 75.52 | 67.54 | n/a | 61.71 | - | - | - | - |
| Fucose | 98.47 | n/a | n/a | n/a | 66.78 | 15.36 | - | - | - | - |
| Neu5Ac | n/a | n/a | 39.80 | n/a | 51.79 | n/a | n/a | n/a | 62.67 | 22.29 |

| Linker | 1 | 2 | 3 | 4 | 5 | 6 |
|---|---|---|---|---|---|---|
| H | 3.87, 3.57 | 1.58 – 1.52 (m, 2H) | 1.35 – 1.27 (m, 2H) | 1.49 (p, J = 7.3 Hz, 2H) | 3.12 (t, J = 6.9 Hz, 2H) | 5.12 (s, 2H) |
| C | 70.60 | 28.40 | 22.45 | 28.51 | 40.48 | 66.87 |

HRMS (ESI-MS): m/z calculated for $C_{72}H_{113}N_5O_{48}S$ $[M-2H]^{2-}$: 923.8143; found: 923.7939.

**Compound 13**

**13** was prepared from **12** (4.0 mg, 2.0 µmol) using the general procedure for installation of α1,3Fuc using FUT6. After P6 and HILIC HPLC purification, **13** was obtained as a white solid (4.1 mg, 95%).

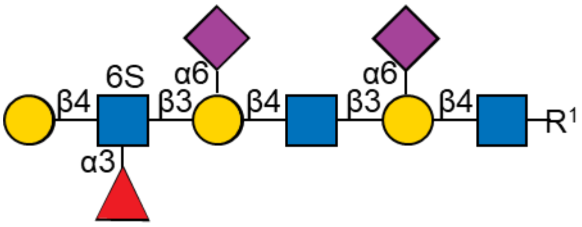

$^1$H (600 MHz, $D_2O$): δ (ppm)

| | H-1 | H-2 | H-3 | H-4 | H-5 | H-6 | H-7 | H-8 | H-9 | NHAc |
|---|---|---|---|---|---|---|---|---|---|---|
| GlcNAc-1 | 4.54 | 3.73 | n/a | n/a | n/a | 3.99, 3.82 | - | - | - | 2.11 – 1.97 (m, 15H) |
| Galactose-1 | 4.44 | 3.61 | 3.73 | 4.17 | n/a | 4.00, 3.55 | - | - | - | - |
| GlcNAc-2 | 4.72 (d, J = 7.7 Hz, 1H) | 3.81 | n/a | n/a | n/a | 3.97, 3.85 | - | - | - | 2.11 – 1.97 (m, 15H) |
| Galactose-2 | 4.44 | 3.61 | 3.73 | 4.17 | n/a | 4.00, 3.55 | - | - | - | - |
| GlcNAc-6S | 4.75 (d, J = 8.2 Hz, 1H) | 3.97 | 3.91 | 4.02 | 3.80 | 4.40 – 4.32 (m, 2H) | - | - | - | 2.11 – 1.97 (m, 15H) |
| Galactose-3 | 4.55 | 3.49 (dd, J = 9.8, 7.8 Hz, 1H) | 3.68 | 3.92 | n/a | 3.73 (2H) | - | - | - | - |
| Fucose | 5.13 (d, J = 4.2 Hz, 1H) | 3.70 | n/a | n/a | 4.84 (d, J = 6.9 Hz, 1H) | 1.19 (d, J = 6.5 Hz, 3H) | - | - | - | - |
| Neu5Ac-1 | - | - | 2.69 – 2.61 (m, 2H), 1.77 – | 3.69 | 3.81 | n/a | 3.57 | 3.90 | 3.88, 3.65 | 2.11 – 1.97 (m, 15H) |

| | | | | | | | | | | |
|---|---|---|---|---|---|---|---|---|---|---|
| | | | 1.67 (m, 2H) | | | | | | | |
| Neu5Ac-2 | - | - | 2.69 – 2.61 (m, 2H), 1.77 – 1.67 (m, 2H) | 3.69 | 3.81 | n/a | 3.57 | 3.90 | 3.88, 3.65 | 2.11 – 1.97 (m, 15H) |

$^{13}$C (150 MHz, $D_2O$): δ (ppm)

| | C-1 | C-2 | C-3 | C-4 | C-5 | C-6 | C-7 | C-8 | C-9 | NHAc |
|---|---|---|---|---|---|---|---|---|---|---|
| GlcNAc-1 | 100.90 | 54.89 | n/a | n/a | n/a | 60.34 | - | - | - | 22.20 |
| Galactose-1 | 103.56 | 69.59 | 82.38 | 68.06 | n/a | 63.26 | - | - | - | - |
| GlcNAc-2 | 102.83 | 54.91 | n/a | n/a | n/a | 60.34 | - | - | - | 22.20 |
| Galactose-2 | 103.56 | 69.59 | 82.38 | 68.06 | n/a | 63.26 | - | - | - | - |
| GlcNAc-6S | 102.68 | 55.89 | n/a | 72.72 | 72.72 | 66.12 | - | - | - | 22.20 |
| Galactose-3 | 101.63 | 71.23 | 72.57 | 68.67 | n/a | 61.49 | - | - | - | - |
| Fucose | 98.36 | n/a | n/a | n/a | 66.59 | 15.31 | - | - | - | - |
| Neu5Ac-1 | n/a | n/a | 40.12 | n/a | 51.89 | n/a | 68.68 | 71.72 | 62.77 | 22.20 |
| Neu5Ac-2 | n/a | n/a | 40.12 | n/a | 51.89 | n/a | 68.68 | 71.72 | 62.77 | 22.20 |

| Linker | 1 | 2 | 3 | 4 | 5 | 6 |
|---|---|---|---|---|---|---|
| H | 3.88,3.58 | 1.61 – 1.52 (m, 2H) | 1.37 – 1.24 (m, 2H) | 1.50 (p, J = 7.3 Hz, 2H) | 3.13 (t, J = 6.8 Hz, 2H) | 5.12 (s, 2H) |
| C | 70.22 | 28.25 | 22.38 | | 40.37 | 66.73 |

HRMS (ESI-MS): m/z calculated for $C_{83}H_{130}N_6O_{56}S$ $[M-2H]^{2-}$: 1069.3620; found: 1069.3497.

**Compound 14**

**14** was prepared from **13** (4.1 mg, 1.9 μmol) using the general procedure for the removal of Neu5Ac using *C. perfringens* neuraminidase. After P6 purification, **14** was obtained as a white solid (3.0 mg, quant).

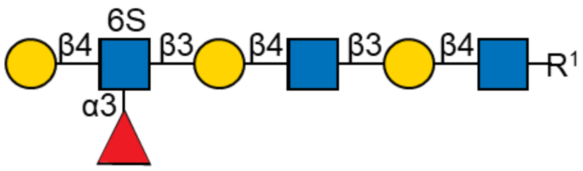

$^{1}$H (600 MHz, $D_2O$): δ (ppm)

| | H-1 | H-2 | H-3 | H-4 | H-5 | H-6 | NHAc |
|---|---|---|---|---|---|---|---|

| | | | | | | | |
|---|---|---|---|---|---|---|---|
| GlcNAc-1 | 4.51 | 3.72 | 3.68 | 3.69 | 3.58 | 3.97, 3.83 | 2.09 – 1.98 (m, 9H) |
| Galactose-1 | 4.46 | 3.60 | 3.72 | 4.16 (d, J = 3.3 Hz, 1H) | 3.72 | 3.76 | - |
| GlcNAc-2 | 4.71 (d, J = 8.2 Hz, 1H) | 3.81 | 3.73 | 3.73 | 3.58 | 3.97, 3.84 | 2.09 – 1.98 (m, 9H) |
| Galactose-2 | 4.48 | 3.60 | 3.72 | 4.19 (d, J = 3.2 Hz, 1H) | 3.72 | 3.76 | - |
| GlcNAc-6S | 4.75 | 4.00 | 3.91 | 4.01 | 3.81 | 4.39 – 4.35 (m, 2H) | 2.09 – 1.98 (m, 9H) |
| Galactose-3 | 4.53 (d, J = 8.1 Hz, 1H) | 3.50 (t, J = 8.9 Hz, 1H) | 3.68 | 3.92 | n/a | 3.73 | - |
| Fucose | 5.14 (d, J = 4.0 Hz, 1H) | 3.70 | n/a | n/a | 4.84 | 1.19 (d, J = 6.6 Hz, 3H) | - |

$^{13}$C (150 MHz, $D_2O$): δ (ppm)

| | C-1 | C-2 | C-3 | C-4 | C-5 | C-6 | NHAc |
|---|---|---|---|---|---|---|---|
| GlcNAc-1 | 101.12 | 55.18 | 72.69 | 78.67 | 74.87 | 60.09 | 22.23 |
| Galactose-1 | 103.16 | 70.02 | 82.56 | 68.39 | 75.18 | 61.33 | - |
| GlcNAc-2 | 102.98 | 55.10 | 72.23 | 78.22 | 74.87 | 60.09 | 22.23 |
| Galactose-2 | 103.03 | 70.02 | 82.56 | 68.35 | 75.18 | 61.33 | - |
| GlcNAc-6S | 102.71 | 56.01 | n/a | 73.12 | 72.61 | 66.44 | 22.23 |
| Galactose-3 | 101.59 | 71.21 | 72.76 | 68.93 | n/a | 61.30 | - |
| Fucose | 98.69 | n/a | n/a | n/a | 66.99 | 15.44 | - |

| Linker | 1 | 2 | 3 | 4 | 5 | 6 |
|---|---|---|---|---|---|---|
| H | 3.88, 3.55 | 1.59 – 1.52 (m, 2H) | 1.35 – 1.26 (m, 2H) | 1.49 (p, J = 7.3 Hz, 2H) | 3.17 – 3.10 (m, 2H) | 5.12 (s, 2H) |
| C | 70.76 | 28.24 | 22.56 | 28.53 | 40.59 | 66.85 |

HRMS (ESI-MS): m/z calculated for $C_{61}H_{96}N_4O_{40}S$ [M-2H]$^{2-}$: 778.2666; found: 778.2627.

**Compound 4**

**4** was prepared from **14** (2.0 mg, 1.3 μmol) using the general procedure for the installation of α2,3 Neu5Ac using Pd2,3ST. During the reaction, tiny α2,6-sialylation side product **4'** was observed (**4**:**4'**>10:1). After P6 and HILIC HPLC purification, **4** was obtained as a white solid (1.5 mg, 65%). NMR data agrees with the above reported data for compound **4**.

**Compound G**

**G** was prepared from **1** (2.2 mg, 2.0 μmol) using the general procedure for Cbz deprotection using $Pd(OH)_2$ reduction. After purification, **G** was obtained as a white solid (1.5 mg, 78%).

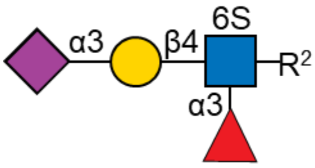

$^{1}H$ (600 MHz, $D_2O$): δ (ppm)

| | H-1 | H-2 | H-3 | H-4 | H-5 | H-6 | H-7 | H-8 | H-9 | NHAc |
|---|---|---|---|---|---|---|---|---|---|---|
| GlcNAc-6S | 4.58 (d, J = 8.0 Hz, 1H) | 3.91 | n/a | 4.02 (t, J = 9.2 Hz, 1H) | 3.81 | 4.41 – 4.37 (m, 2H) | - | - | - | 2.09 – 1.99 (m, 6H) |
| Galactose | 4.61 (d, J = 7.8 Hz, 1H) | 3.52 (dd, J = 9.8, 7.8 Hz, 1H) | 4.11 (dd, J = 9.8, 3.2 Hz, 1H) | 3.96 (d, J = 3.5 Hz, 1H) | n/a | 3.70 | - | - | - | - |
| Fucose | 5.11 (d, J = 4.0 Hz, 1H) | 3.70 | n/a | n/a | 4.82 | 1.18 (d, J = 6.6 Hz, 3H) | - | - | - | - |
| Neu5Ac | - | - | 2.76 (dd, J = 12.4, 4.6 Hz, 1H), 1.81 (t, J = 12.2 Hz, 1H) | 3.69 | 3.86 | n/a | n/a | 3.93 | 3.90, 3.66 | 2.09 – 1.99 (m, 6H) |

$^{13}C$ (150 MHz, $D_2O$): δ (ppm)

| | C-1 | C-2 | C-3 | C-4 | C-5 | C-6 | C-7 | C-8 | C-9 | NHAc |
|---|---|---|---|---|---|---|---|---|---|---|
| GlcNAc-6S | 101.06 | 55.65 | n/a | 72.88 | 72.54 | 66.04 | - | - | - | 22.23 |
| Galactose | 101.37 | 69.43 | 75.60 | 67.43 | n/a | 61.54 | - | - | - | - |
| Fucose | 98.52 | n/a | n/a | n/a | 66.88 | 15.33 | - | - | - | - |
| Neu5Ac | n/a | n/a | 39.83 | n/a | 51.90 | n/a | n/a | 71.64 | 62.63 | 22.23 |

| Linker | 1 | 2 | 3 | 4 | 5 |
|---|---|---|---|---|---|
| H | 3.88, 3.64 | 1.62 (p, J = 7.1 Hz, 2H) | 1.48 – 1.36 (m, 2H) | 1.69 (p, J = 7.7 Hz, 2H) | 3.00 (t, J = 7.6 Hz, 2H) |

| | | | | | |
|---|---|---|---|---|---|
| C | 70.36 | 28.32 | 22.10 | 26.52 | 39.40 |

HRMS (ESI-MS): m/z calculated for $C_{36}H_{61}N_3O_{26}S$ $[M-2H]^{2-}$: 491.6637; found: 491.6746.

**Compound N**

**N** was prepared from **2** (1.2 mg, 0.7 μmol) using the general procedure for Cbz deprotection using $Pd(OH)_2$ reduction. After purification, **N** was obtained as a white solid (0.9 mg, 82%).

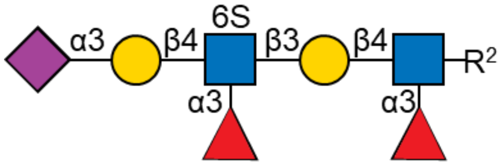

$^1H$ (600 MHz, $D_2O$): δ (ppm)

| | H-1 | H-2 | H-3 | H-4 | H-5 | H-6 | H-7 | H-8 | H-9 | NHAc |
|---|---|---|---|---|---|---|---|---|---|---|
| GlcNAc | 4.54 (d, J = 8.2 Hz, 1H) | 3.90 | n/a | n/a | 3.58 | 4.00, 3.85 | - | - | - | 2.06 – 2.01 (m, 9H) |
| Galactose-1 | 4.44 (d, J = 7.8 Hz, 1H) | 3.51 | 3.71 | 4.12 (d, J = 3.4 Hz, 1H) | n/a | 3.71 | - | - | - | - |
| GlcNAc-6S | 4.73 (d, J = 8.4 Hz, 1H) | 3.99 | 3.91 | 4.02 | 3.81 | 4.40 – 4.32 (m, 2H) | - | - | - | 2.06 – 2.01 (m, 9H) |
| Galactose-2 | 4.60 (d, J = 7.8 Hz, 1H) | 3.53 | 4.10 (dd, J = 9.9, 3.2 Hz, 1H) | 3.96 (d, J = 3.2 Hz, 1H) | n/a | 3.71 | - | - | - | - |
| Fucose-1 | 5.10 (d, J = 4.0 Hz, 1H) | 3.69 | n/a | n/a | 4.82 | 1.15 (d, J = 6.6 Hz, 3H) | | | | - |
| Fucose-2 | 5.13 (d, J = 4.0 Hz, 1H) | 3.69 | n/a | n/a | 4.82 | 1.18 (d, J = 6.6 Hz, 3H) | - | - | - | - |
| Neu5Ac | - | - | 2.76 (dd, J = 12.4, 4.6 Hz, 1H), | 3.68 | 3.86 | n/a | n/a | 3.93 | 3.90, 3.65 | 2.06 – 2.01 (m, 9H) |

| | | | |
|---|---|---|---|
| | | | 1.80 (t, J = 12.1 Hz, 1H) |

$^{13}C$ (150 MHz, $D_2O$): δ (ppm)

| | C-1 | C-2 | C-3 | C-4 | C-5 | C-6 | C-7 | C-8 | C-9 | NHAc |
|---|---|---|---|---|---|---|---|---|---|---|
| GlcNAc | 101.04 | 55.89 | n/a | n/a | 75.10 | 59.88 | - | - | - | 22.30 |
| Galactose-1 | 101.84 | 70.16 | n/a | 68.24 | n/a | 61.68 | - | - | - | - |
| GlcNAc-6S | 102.51 | 55.89 | n/a | 72.71 | 72.42 | 66.11 | - | - | - | 22.30 |
| Galactose-2 | 101.33 | 69.50 | 75.55 | 67.35 | n/a | 61.68 | - | - | - | - |
| Fucose-1 | 98.74 | n/a | n/a | n/a | 66.76 | 15.37 | | | | - |
| Fucose-2 | 98.54 | n/a | n/a | n/a | 66.76 | 15.37 | - | - | - | - |
| Neu5Ac | n/a | n/a | 39.75 | n/a | 51.74 | n/a | n/a | 71.54 | 62.64 | 22.30 |

| Linker | 1 | 2 | 3 | 4 | 5 |
|---|---|---|---|---|---|
| H | 3.90, 3.60 | 1.63 – 1.54 (m, 2H) | 1.43 – 1.36 (m, 2H) | 1.67 (p, J = 7.7 Hz, 2H) | 3.01 – 2.93 (m, 2H) |
| C | 70.26 | 28.14 | 22.21 | 26.41 | 39.33 |

HRMS (ESI-MS): m/z calculated for $C_{56}H_{94}N_4O_{40}S$ $[M-2H]^{2-}$: 747.2588; found: 747.2692.

**Compound S6**

**S6** was prepared from **S1** (0.4 mg, 0.2 μmol) using the general procedure for Cbz deprotection using $Pd(OH)_2$ reduction. After purification, **S6** was obtained as a white solid (0.25 mg, 67%).

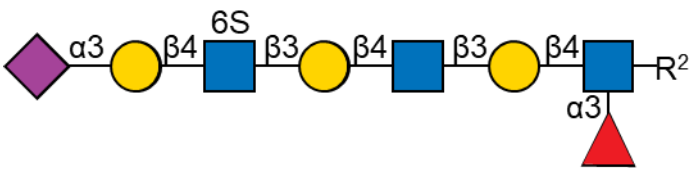

$^1H$ (600 MHz, $D_2O$): δ (ppm)

| | H-1 | H-2 | H-3 | H-4 | H-5 | H-6 | H-7 | H-8 | H-9 | NHAc |
|---|---|---|---|---|---|---|---|---|---|---|
| GlcNAc-1 | 4.54 (d, J = 8.2 Hz, 1H) | 3.90 | n/a | n/a | 3.59 | 3.98, 3.85 | - | - | - | 2.09 – 1.98 (m, 12H) |
| Galactose-1 | 4.45 (d, J = 7.7 Hz, 1H) | 3.52 | 3.73 | 4.11 (d, J = 3.3 Hz, 1H) | n/a | 3.71 (2H) | - | - | - | - |

| | | | | | | | | | | |
|---|---|---|---|---|---|---|---|---|---|---|
| GlcNAc-2 | 4.71 | 3.80 | 3.74 | 3.74 | 3.59 | 3.98, 3.85 | - | - | - | 2.09 – 1.98 (m, 12H) |
| Galactose-2 | 4.48 (d, J = 7.8 Hz, 1H) | 3.59 | 3.73 | 4.20 (d, J = 3.2 Hz, 1H) | n/a | 3.76 (2H) | - | - | - | - |
| GlcNAc-6S | 4.72 | 3.85 | 3.74 | 3.81 | 3.81 | 4.43 – 4.39 (m, 1H), 4.32 (d, J = 10.7 Hz, 1H) | - | - | - | 2.09 – 1.98 (m, 12H) |
| Galactose-3 | 4.61 (d, J = 7.9 Hz, 1H) | 3.56 | 4.13 (dd, J = 9.6, 2.5 Hz, 1H) | 3.97 | n/a | n/a | - | - | - | - |
| Fucose | 5.11 (d, J = 4.1 Hz, 1H) | 3.70 | - | - | 4.82 | 1.16 (d, J = 6.6 Hz, 3H) | - | - | - | - |
| Neu5Ac | - | - | 2.76 (dd, J = 12.4, 4.7 Hz, 1H), 1.81 (t, J = 12.2 Hz, 1H) | 3.67 | 3.86 | n/a | n/a | 3.92 | 3.90, 3.66 | 2.09 – 1.98 (m, 12H) |

$^{13}$C (150 MHz, $D_2O$): δ (ppm)

| | C-1 | C-2 | C-3 | C-4 | C-5 | C-6 | H-7 | H-8 | H-9 | NHAc |
|---|---|---|---|---|---|---|---|---|---|---|
| GlcNAc-1 | 101.19 | 55.84 | n/a | n/a | 75.06 | 59.84 | - | - | - | 22.26 |
| Galactose-1 | 101.92 | 70.83 | 81.93 | 68.25 | n/a | 61.52 | - | - | - | - |
| GlcNAc-2 | 102.70 | 55.18 | 72.31 | 78.43 | 75.06 | 59.84 | - | - | - | 22.26 |
| Galactose-2 | 103.13 | 69.60 | 82.30 | 68.35 | n/a | 61.30 | - | - | - | - |
| GlcNAc-6S | 103.23 | 55.44 | 72.37 | 77.79 | 72.66 | 66.64 | - | - | - | 22.26 |
| Galactose- | 102.34 | 69.59 | 75.52 | 67.60 | n/a | n/a | - | - | - | - |

| 3 | | | | | | | | | | |
|---|---|---|---|---|---|---|---|---|---|---|
| Fucose | 98.81 | n/a | n/a | n/a | 66.85 | 15.48 | - | - | - | - |
| Neu5Ac | n/a | n/a | 39.60 | n/a | 51.77 | n/a | n/a | 71.96 | 62.88 | 22.26 |

| Linker | 1 | 2 | 3 | 4 | 5 |
|---|---|---|---|---|---|
| H | 3.91, 3.61 | 1.60 (p, J = 6.6 Hz, 2H) | 1.46 – 1.37 (m, 2H) | 1.68 (p, J = 7.7 Hz, 2H) | 3.02 – 2.97 (m, 2H) |
| C | 70.41 | 28.24 | 22.29 | 26.44 | 39.43 |

HRMS (ESI-MS): m/z calculated for $C_{64}H_{107}N_5O_{46}S$ $[M-2H]^{2-}$: 856.7959; found: 856.8520.

**Compound X**

**X** was prepared from **3** (0.6 mg, 0.3 μmol) using the general procedure for Cbz deprotection using $Pd(OH)_2$ reduction. After purification, **X** was obtained as a white solid (0.3 mg, 56%).

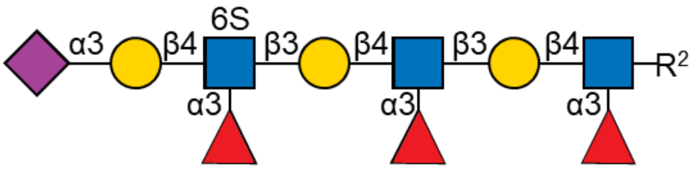

$^1H$ (600 MHz, $D_2O$): δ (ppm)

| | H-1 | H-2 | H-3 | H-4 | H-5 | H-6 | H-7 | H-8 | H-9 | NHAc |
|---|---|---|---|---|---|---|---|---|---|---|
| GlcNAc-1 | 4.54 (d, J = 8.2 Hz, 1H) | 3.90 | n/a | n/a | 3.58 | 3.99, 3.86 | - | - | - | 2.06 – 2.01 (m, 12H) |
| Galactose-1 | 4.44 | 3.51 | 3.72 | 4.11 | n/a | 3.71 (4H) | - | - | - | - |
| GlcNAc-2 | 4.72 | 3.97 | n/a | n/a | 3.59 | 3.96, 3.86 | - | - | - | 2.06 – 2.01 (m, 12H) |
| Galactose-2 | 4.46 | 3.53 | 3.72 | 4.13 (d, J = 3.3 Hz, 1H) | n/a | 3.71 (4H) | - | - | - | - |
| GlcNAc-6S | 4.74 | 4.00 | 3.91 | 4.03 | 3.80 | 4.41 – 4.34 (m, 2H) | - | - | - | 2.06 – 2.01 (m, 12H) |
| Galactose-3 | 4.61 (d, J = 7.8 Hz, 1H) | 3.53 | 4.10 | 3.96 | n/a | n/a | - | - | - | - |

| | | | | | | | | | | |
|---|---|---|---|---|---|---|---|---|---|---|
| Fucose-1 | 5.11 (d, J = 4.0 Hz, 1H) | 3.70 | n/a | n/a | 4.82 | 1.16 | - | - | - | - |
| Fucose-2 | 5.13 | 3.70 | n/a | n/a | 4.82 | 1.16 | - | - | - | - |
| Fucose-3 | 5.14 | 3.70 | n/a | n/a | 4.82 | 1.18 (d, J = 6.7 Hz, 3H) | - | - | - | - |
| Neu5Ac | - | - | 2.76 (dd, J = 12.3, 4.6 Hz, 1H), 1.81 (t, J = 12.1 Hz, 1H) | 3.70 | 3.86 | n/a | n/a | 3.93 | 3.90, 3.65 | 2.06 – 2.01 (m, 12H) |

$^{13}C$ (150 MHz, $D_2O$): δ (ppm)

| | C-1 | C-2 | C-3 | C-4 | C-5 | C-6 | H-7 | H-8 | H-9 | NHAc |
|---|---|---|---|---|---|---|---|---|---|---|
| GlcNAc-1 | 101.20 | 55.96 | n/a | n/a | 75.10 | 59.87 | - | - | - | 22.33 |
| Galactose-1 | 101.67 | 70.64 | 82.03 | 68.38 | n/a | 61.58 | - | - | - | - |
| GlcNAc-2 | 102.51 | 56.08 | n/a | n/a | 75.27 | 59.87 | - | - | - | 22.33 |
| Galactose-2 | 102.05 | 70.37 | 82.03 | 68.50 | n/a | 61.58 | - | - | - | - |
| GlcNAc-6S | 102.68 | 56.08 | n/a | 72.87 | 72.39 | 66.23 | - | - | - | 22.33 |
| Galactose-3 | 101.31 | 70.37 | 75.67 | 67.61 | n/a | n/a | - | - | - | - |
| Fucose-1 | 98.72 | n/a | n/a | n/a | 66.82 | 15.39 | - | - | - | - |
| Fucose-2 | 98.67 | n/a | n/a | n/a | 66.82 | 15.39 | - | - | - | - |
| Fucose-3 | 98.41 | n/a | n/a | n/a | 66.82 | 15.39 | - | - | - | - |
| Neu5Ac | n/a | n/a | 39.83 | n/a | 51.85 | n/a | n/a | 71.70 | 62.68 | 22.33 |

| Linker | 1 | 2 | 3 | 4 | 5 |
|---|---|---|---|---|---|
| H | 3.91, 3.61 | 1.60 (p, J = 6.6 Hz, 2H) | 1.44 – 1.37 (m, 2H) | 1.68 (p, J = 7.7 Hz, 2H) | 3.02 – 2.97 (m, 2H) |
| C | 70.37 | 28.29 | 22.33 | 26.55 | 39.46 |

HRMS (ESI-MS): m/z calculated for $C_{76}H_{127}N_5O_{54}S$ $[M-2H]^{2-}$: 1002.8538; found: 1002.8325.

**Compound S**

**S** was prepared from **14** (0.6 mg, 0.4 μmol) using the general procedure for Cbz deprotection using $Pd(OH)_2$ reduction. After purification, **S** was obtained as a white solid (0.4 mg, 73%).

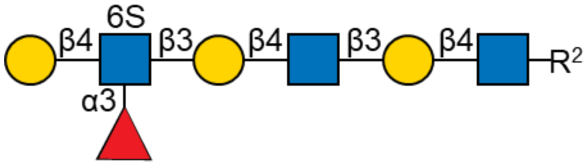

$^{1}H$ (600 MHz, $D_2O$): δ (ppm)

| | H-1 | H-2 | H-3 | H-4 | H-5 | H-6 | NHAc |
|---|---|---|---|---|---|---|---|
| GlcNAc-1 | 4.53 | 3.73 | 3.70 | 3.69 | 3.58 | 3.99, 3.82 | 2.06 – 2.02 (m, 9H) |
| Galactose-1 | 4.46 | 3.59 | 3.72 | 4.17 (d, J = 3.2 Hz, 1H) | 3.72 | 3.76 | - |
| GlcNAc-2 | 4.71 | 3.81 | 3.73 | 3.73 | 3.58 | 3.97, 3.84 | 2.06 – 2.02 (m, 9H) |
| Galactose-2 | 4.48 | 3.59 | 3.72 | 4.19 (d, J = 3.2 Hz, 1H) | 3.72 | 3.76 | - |
| GlcNAc-6S | 4.75 | 3.99 | 3.91 | 4.01 | 3.81 | 4.39 – 4.35 (m, 2H) | 2.06 – 2.02 (m, 9H) |
| Galactose-3 | 4.53 | 3.50 | 3.68 | 3.92 | n/a | 3.74 | - |
| Fucose | 5.14 (d, J = 4.0 Hz, 1H) | 3.70 | n/a | n/a | 4.84 | 1.19 (d, J = 6.6 Hz, 3H) | - |

$^{13}C$ (150 MHz, $D_2O$): δ (ppm)

| | C-1 | C-2 | C-3 | C-4 | C-5 | C-6 | NHAc |
|---|---|---|---|---|---|---|---|
| GlcNAc-1 | 101.12 | 55.18 | 72.64 | 78.32 | 74.77 | 60.22 | 22.23 |
| Galactose-1 | 102.84 | 70.02 | 82.41 | 68.39 | 75.18 | 61.23 | - |
| GlcNAc-2 | 102.70 | 55.16 | 72.23 | 78.22 | 74.87 | 60.09 | 22.23 |
| Galactose-2 | 102.84 | 70.02 | 82.41 | 68.35 | 75.18 | 61.23 | - |
| GlcNAc-6S | 102.49 | 56.01 | n/a | 73.12 | 72.61 | 66.44 | 22.23 |
| Galactose-3 | 101.97 | 71.22 | 72.76 | 68.54 | n/a | 61.30 | - |
| Fucose | 98.55 | n/a | n/a | n/a | 66.71 | 15.47 | - |

| Linker | 1 | 2 | 3 | 4 | 5 |
|---|---|---|---|---|---|
| H | 3.91, 3.61 | 1.61 (p, J = | 1.44 – 1.37 | 1.68 (p, J = | 3.02 – 2.97 |

| | | | | | |
|---|---|---|---|---|---|
| | | 6.7 Hz, 2H) | (m, 2H) | 7.7 Hz, 2H) | (m, 2H) |
| C | 70.37 | 28.29 | 22.33 | 26.55 | 39.46 |

HRMS (ESI-MS): m/z calculated for $C_{53}H_{90}N_4O_{38}S$ $[M-2H]^{2-}$: 711.2482; found: 711.2476.

**Compound Y**

**Y** was prepared from **4** (5.0 mg, 2.7 μmol) using the general procedure for Cbz deprotection using $Pd(OH)_2$ reduction. After purification, **Y** was obtained as a white solid (3.6 mg, 78%).

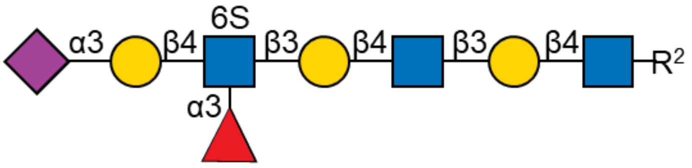

$^1H$ (600 MHz, $D_2O$): δ (ppm)

| | H-1 | H-2 | H-3 | H-4 | H-5 | H-6 | H-7 | H-8 | H-9 | NHAc |
|---|---|---|---|---|---|---|---|---|---|---|
| GlcNAc-1 | 4.53 (d, *J* = 7.5 Hz, 1H) | 3.72 | 3.69 | 3.69 | 3.58 | 3.97, 3.83 | - | - | - | 2.06 – 2.01 (m, 12H) |
| Galactose-1 | 4.47 | 3.60 | 3.72 | 4.17 (d, *J* = 3.2 Hz, 1H) | n/a | 3.76 (4H) | - | - | - | - |
| GlcNAc-2 | 4.71 (d, *J* = 8.5 Hz, 1H) | 3.81 | n/a | n/a | 3.58 | 3.97, 3.83 | - | - | - | 2.06 – 2.01 (m, 12H) |
| Galactose-2 | 4.48 | 3.60 | 3.72 | 4.20 (d, J = 3.2 Hz, 1H) | n/a | 3.76 (4H) | - | - | - | - |
| GlcNAc-6S | 4.73 | 4.00 | 3.91 | 4.02 | 3.80 | 4.40 – 4.35 (m, 2H) | - | - | - | 2.06 – 2.01 (m, 12H) |
| Galactose-3 | 4.60 (d, J = 7.7 Hz, 1H) | 3.52 | 4.10 | 3.98 | n/a | 3.71 (2H) | - | - | - | - |
| Fucose | 5.13 (d, *J* | 3.68 | n/a | n/a | 4.82 | 1.18 (d, *J* = | - | - | - | - |

| | | | | | | | | | | |
|---|---|---|---|---|---|---|---|---|---|---|
| | = 4.1 Hz, 1H) | | | | | 6.5 Hz, 3H) | | | | |
| Neu5Ac | - | - | 2.76 (dd, *J* = 12.3, 4.6 Hz, 1H), 1.81 (t, J = 12.1 Hz, 1H) | 3.69 | 3.86 | n/a | n/a | n/a | 3.90, 3.65 | 2.06 – 2.01 (m, 12H) |

$^{13}$C (150 MHz, $D_2O$): δ (ppm)

| | C-1 | C-2 | C-3 | C-4 | C-5 | C-6 | H-7 | H-8 | H-9 | NHAc |
|---|---|---|---|---|---|---|---|---|---|---|
| GlcNAc-1 | 101.16 | 55.15 | 72.80 | 78.34 | 74.90 | 59.97 | - | - | - | 22.13 |
| Galactose-1 | 103.05 | 69.64 | 82.38 | 68.28 | n/a | 61.08 | - | - | - | - |
| GlcNAc-2 | 102.70 | 55.23 | n/a | n/a | 74.90 | 59.97 | - | - | - | 22.13 |
| Galactose-2 | 102.88 | 69.64 | 82.38 | 68.20 | n/a | 61.08 | - | - | - | - |
| GlcNAc-6S | 102.58 | 56.07 | n/a | 73.05 | 72.94 | 66.17 | - | - | - | 22.13 |
| Galactose-3 | 101.40 | 69.58 | 75.63 | 67.48 | n/a | 61.42 | - | - | - | - |
| Fucose | 98.57 | n/a | n/a | n/a | 66.77 | 15.49 | - | - | - | - |
| Neu5Ac | n/a | n/a | 39.76 | n/a | 51.85 | n/a | n/a | n/a | 62.60 | 22.13 |

| Linker | 1 | 2 | 3 | 4 | 5 |
|---|---|---|---|---|---|
| H | 3.91, 3.62 | 1.61 (p, *J* = 6.8 Hz, 2H) | 1.44 – 1.37 (m, 2H) | 1.68 (p, J = 7.7 Hz, 2H) | 3.00 (t, *J* = 7.7 Hz, 2H) |
| C | 70.75 | 28.22 | 22.33 | 26.55 | 39.24 |

HRMS (ESI-MS): m/z calculated for $C_{64}H_{107}N_5O_{46}S$ $[M-2H]^{2-}$: 856.7959; found: 856.7989.

##### Compound 21

**21** was prepared from **G** (0.5 mg, 0.5 µmol) using the general procedure for synthesis of BCN linked glycans. After purification, **21** was obtained as a white solid (0.6 mg, 79%).

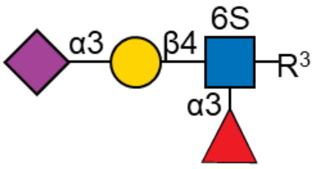

$^{1}H$ (600 MHz, $D_2O$): δ (ppm)

| | H-1 | H-2 | H-3 | H-4 | H-5 | H-6 | H-7 | H-8 | H-9 | NHAc |
|---|---|---|---|---|---|---|---|---|---|---|
| GlcNAc-6S | 4.57 (d, *J* = 7.8 Hz, 1H) | 3.90 | n/a | 4.02 (t, *J* = 9.0 Hz, 1H) | 3.81 | 4.40 – 4.35 (m, 2H) | - | - | - | 2.06 – 2.01 (m, 6H) |
| Galactose | 4.62 (d, *J* = 7.8 Hz, 1H) | 3.52 (t, *J* = 8.8 Hz, 1H) | 4.11 | 3.96 | n/a | 3.71 | - | - | - | - |
| Fucose | 5.11 (d, J = 4.1 Hz, 1H) | 3.70 | n/a | n/a | 4.82 | 1.18 (d, J = 6.6 Hz, 3H) | - | - | - | - |
| Neu5Ac | - | - | 2.76 (dd, J = 12.5, 4.6 Hz, 1H), 1.81 (t, J = 12.1 Hz, 1H) | 3.69 | 3.86 | n/a | n/a | 3.94 | 3.90, 3.66 | 2.06 – 2.01 (m, 6H) |

$^{13}C$ (150 MHz, $D_2O$): δ (ppm)

| | C-1 | C-2 | C-3 | C-4 | C-5 | C-6 | C-7 | C-8 | C-9 | NHAc |
|---|---|---|---|---|---|---|---|---|---|---|
| GlcNAc-6S | 100.90 | 55.74 | n/a | 72.68 | 72.54 | 65.93 | - | - | - | 22.23 |
| Galactose | 101.21 | 69.45 | 75.50 | n/a | n/a | 61.54 | - | - | - | - |
| Fucose | 98.57 | n/a | n/a | n/a | 66.80 | 15.34 | - | - | - | - |
| Neu5Ac | n/a | n/a | 39.83 | n/a | 51.78 | n/a | n/a | 71.54 | 62.61 | 22.23 |

| Linker | 1 | 2 | 3 | 4 | 5 |
|---|---|---|---|---|---|
| H | 3.89, 3.59 | 1.60 – 1.56 (m, 2H) | 1.37 – 1.31 (m, 2H) | 1.54 – 1.47 (m, 2H) | 3.12 (t, *J* = 6.2 Hz, 2H) |
| C | 70.48 | 28.40 | 22.64 | 28.62 | 40.56 |

| BCN | 6 | 7 | 8,8’ | 9,9’ | 10,10’ |
|---|---|---|---|---|---|
| H | 4.19 (d, *J* = 8.4 Hz, 2H) | 1.41 (t, J = 8.7 Hz, 1H) | 1.00 (t, *J* = 9.6 Hz, 1H) | 2.26 (2H), 1.62 (2H) | 2.36 – 2.21 (m, 4H) |
| C | 63.81 | 17.01 | 19.63 | 28.59 | 20.77 |

HRMS (ESI-MS): m/z calculated for $C_{47}H_{73}N_3O_{28}S$ $[M-2H]^{2-}$: 579.7056; found: 579.6722.

**Compound 23**

**23** was prepared from **X** (2.0 mg, 1.0 μmol) using the general procedure for synthesis of BCN linked glycans. After purification, **23** was obtained as a white solid (1.0 mg, 46%).

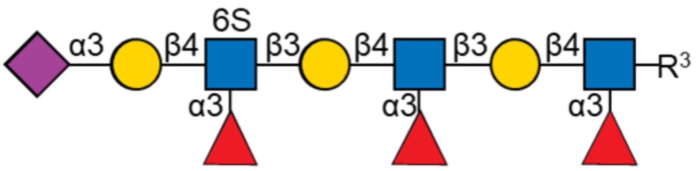

$^1$H (600 MHz, $D_2O$): δ (ppm)

| | H-1 | H-2 | H-3 | H-4 | H-5 | H-6 | H-7 | H-8 | H-9 | NHAc |
|---|---|---|---|---|---|---|---|---|---|---|
| GlcNAc-1 | 4.53 (d, *J* = 7.9 Hz, 1H) | 3.88 | n/a | n/a | 3.58 | 3.98, 3.85 | - | - | - | 2.06 – 2.01 (m, 12H) |
| Galactose-1 | 4.44 | 3.51 | 3.71 | 4.10 (d, *J* = 3.3 Hz, 1H) | n/a | 3.71 (4H) | - | - | - | - |
| GlcNAc-2 | 4.71 | 3.97 | n/a | n/a | 3.59 | 3.98, 3.85 | - | - | - | 2.06 – 2.01 (m, 12H) |
| Galactose-2 | 4.46 | 3.53 | 3.71 | 4.12 | n/a | 3.71 (4H) | - | - | - | - |
| GlcNAc-6S | 4.74 | 4.00 | 3.91 | 4.03 | 3.80 | 4.40 – 4.34 (m, 2H) | - | - | - | 2.06 – 2.01 (m, 12H) |
| Galactose-3 | 4.60 (d, *J* = 7.8 Hz, 1H) | 3.53 | 4.10 | 3.96 | n/a | n/a | - | - | - | - |
| Fucose-1 | 5.10 (d, *J* = 3.9 Hz, 1H) | 3.70 | n/a | n/a | 4.82 | 1.15 (d, *J* = 6.5 Hz, 6H) | - | - | - | - |
| Fucose-2 | 5.13 | 3.70 | n/a | n/a | 4.82 | 1.15 (d, *J* = 6.5 Hz, 6H) | - | - | - | - |
| Fucose-3 | 5.14 | 3.70 | n/a | n/a | 4.82 | 1.18 (d, J = 6.7 Hz, | - | - | - | - |

| | | | | | | | | | | |
|---|---|---|---|---|---|---|---|---|---|---|
| | | | | | | 3H) | | | | |
| Neu5Ac | - | - | 2.76 (d, *J* = 12.1 Hz, 1H), 1.81 (t, *J* = 12.0 Hz, 1H) | 3.70 | 3.86 | n/a | n/a | 3.93 | 3.90, 3.65 | 2.06 – 2.01 (m, 12H) |

$^{13}$C (150 MHz, $D_2O$): δ (ppm)

| | C-1 | C-2 | C-3 | C-4 | C-5 | C-6 | H-7 | H-8 | H-9 | NHAc |
|---|---|---|---|---|---|---|---|---|---|---|
| GlcNAc-1 | 101.01 | 55.96 | n/a | n/a | 75.10 | 59.87 | - | - | - | 22.26 |
| Galactose-1 | 101.67 | 70.14 | 82.03 | 68.19 | n/a | 61.58 | - | - | - | - |
| GlcNAc-2 | 102.23 | 56.08 | n/a | n/a | 75.27 | 59.87 | - | - | - | 22.26 |
| Galactose-2 | 102.05 | 70.35 | 82.03 | 68.16 | n/a | 61.58 | - | - | - | - |
| GlcNAc-6S | 102.68 | 56.08 | n/a | 72.87 | 72.39 | 66.03 | - | - | - | 22.26 |
| Galactose-3 | 101.21 | 70.37 | 75.44 | 67.61 | n/a | n/a | - | - | - | - |
| Fucose-1 | 98.72 | n/a | n/a | n/a | 67.02 | 15.53 | - | - | - | - |
| Fucose-2 | 98.67 | n/a | n/a | n/a | 67.02 | 15.53 | - | - | - | - |
| Fucose-3 | 98.41 | n/a | n/a | n/a | 67.02 | 15.53 | - | - | - | - |
| Neu5Ac | n/a | n/a | 39.85 | n/a | 51.97 | n/a | n/a | 71.70 | 62.50 | 22.26 |

| Linker | 1 | 2 | 3 | 4 | 5 |
|---|---|---|---|---|---|
| H | 3.89, 3.58 | 1.60 – 1.56 (m, 2H) | 1.37 – 1.31 (m, 2H) | 1.54 – 1.47 (m, 2H) | 3.12 (t, *J* = 6.8 Hz, 2H) |
| C | 70.92 | 28.40 | 22.64 | 28.62 | 40.43 |

| BCN | 6 | 7 | 8,8’ | 9,9’ | 10,10’ |
|---|---|---|---|---|---|
| H | 4.19 (d, *J* = 8.3 Hz, 2H) | 1.40 (t, *J* = 8.8 Hz, 1H) | 0.99 (t, *J* = 10.0 Hz, 2H) | 2.26 (2H), 1.62 (2H) | 2.36 – 2.21 (m, 4H) |
| C | 63.81 | 16.81 | 19.63 | 28.59 | 20.62 |

HRMS (ESI-MS): m/z calculated for $C_{87}H_{139}N_5O_{56}S$ $[M-2H]^{2-}$: 1090.8957; found: 1090.8874.

**Compound 22**

**22** was prepared from **Y** (1.8 mg, 0.1 μmol) using the general procedure for synthesis of BCN linked glycans. After purification, **22** was obtained as a white solid (1.1 mg, 58%).

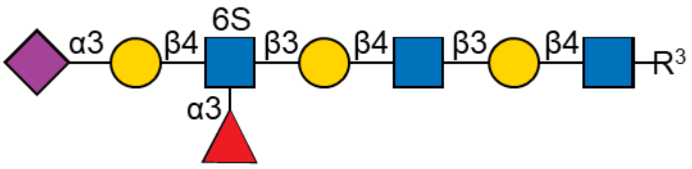

$^{1}$H (600 MHz, $D_2O$): δ (ppm)

| | H-1 | H-2 | H-3 | H-4 | H-5 | H-6 | H-7 | H-8 | H-9 | NHAc |
|---|---|---|---|---|---|---|---|---|---|---|
| GlcNAc-1 | 4.53 (d, *J* = 7.6 Hz, 1H) | 3.72 | 3.69 | 3.69 | 3.58 | 3.97, 3.83 | - | - | - | 2.06 – 2.01 (m, 12H) |
| Galactose-1 | 4.47 | 3.60 | 3.72 | 4.16 (d, *J* = 3.2 Hz, 1H) | n/a | 3.76 (4H) | - | - | - | - |
| GlcNAc-2 | 4.71 | 3.81 | n/a | n/a | 3.58 | 3.97, 3.83 | - | - | - | 2.06 – 2.01 (m, 12H) |
| Galactose-2 | 4.48 | 3.60 | 3.72 | 4.19 | n/a | 3.76 (4H) | - | - | - | - |
| GlcNAc-6S | 4.73 | 3.99 | 3.91 | 4.03 (t, *J* = 9.5 Hz, 1H) | 3.80 | 4.40 – 4.35 (m, 2H) | - | - | - | 2.06 – 2.01 (m, 12H) |
| Galactose-3 | 4.60 (d, *J* = 7.8 Hz, 1H) | 3.52 | 4.11 (dd, *J* = 9.7, 3.2 Hz, 1H) | 3.96 | n/a | 3.71 (2H) | - | - | - | - |
| Fucose | 5.13 (d, *J* = 4.0 Hz, 1H) | 3.68 | n/a | n/a | 4.82 | 1.18 (d, *J* = 6.6 Hz, 3H) | - | - | - | - |
| Neu5Ac | - | - | 2.76 (dd, *J* = 12.5, 4.6 Hz, 1H), 1.81 (t, J = 12.1 | 3.69 | 3.86 | n/a | n/a | n/a | 3.90, 3.65 | 2.06 – 2.01 (m, 12H) |

| | | | |
|---|---|---|---|
| | | | Hz, 1H) |

$^{13}C$ (150 MHz, $D_2O$): δ (ppm)

| | C-1 | C-2 | C-3 | C-4 | C-5 | C-6 | H-7 | H-8 | H-9 | NHAc |
|---|---|---|---|---|---|---|---|---|---|---|
| GlcNAc-1 | 101.16 | 55.05 | 72.80 | 78.34 | 74.90 | 60.17 | - | - | - | 22.13 |
| Galactose-1 | 103.05 | 69.64 | 82.45 | 68.31 | n/a | 61.08 | - | - | - | - |
| GlcNAc-2 | 102.70 | 55.05 | n/a | n/a | 74.90 | 60.17 | - | - | - | 22.13 |
| Galactose-2 | 102.88 | 69.64 | 82.45 | 68.26 | n/a | 61.08 | - | - | - | - |
| GlcNAc-6S | 102.58 | 55.87 | n/a | 72.69 | 72.94 | 66.17 | - | - | - | 22.13 |
| Galactose-3 | 101.30 | 69.74 | 75.51 | 67.48 | n/a | 61.84 | - | - | - | - |
| Fucose | 98.50 | n/a | n/a | n/a | 66.77 | 15.39 | - | - | - | - |
| Neu5Ac | n/a | n/a | 39.83 | n/a | 51.85 | n/a | n/a | n/a | 62.60 | 22.13 |

| Linker | 1 | 2 | 3 | 4 | 5 |
|---|---|---|---|---|---|
| H | 3.89, 3.58 | 1.60 – 1.56 (m, 2H) | 1.37 – 1.31 (m, 2H) | 1.54 – 1.47 (m, 2H) | 3.13 (t, *J* = 5.6 Hz, 2H) |
| C | 70.55 | 28.40 | 22.64 | 28.69 | 40.43 |

| BCN | 6 | 7 | 8,8’ | 9,9’ | 10,10’ |
|---|---|---|---|---|---|
| H | 4.21 – 4.18 (m, 2H) | 1.41 (t, J = 8.6 Hz, 1H) | 1.00 (t, *J* = 10.0 Hz, 2H) | 2.26 (2H), 1.62 (2H) | 2.36 – 2.21 (m, 4H) |
| C | 63.73 | 17.03 | 19.53 | 28.59 | 20.74 |

HRMS (ESI-MS): m/z calculated for $C_{75}H_{119}N_5O_{48}S$ $[M-2H]^{2-}$: 944.8378; found: 944.7797.

**Compound 19**

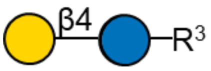

**19** was prepared (0.8 mg, 1.9 µmol) using the general procedure for synthesis of BCN linked glycans. After purification, **19** was obtained as a white solid (0.6 mg, 1.0 µmol, 53%).

$^{1}$**H-NMR:** (600 MHz,$D_2O$) δ 4.39 (d, *J* = 8.1 Hz, 1H, H-1 Glc), 4.37 (d, *J* = 7.8 Hz, 1H, H-1 Gal), 4.10 (d, *J* = 8.3 Hz, 2H, C(O)OC$\underline{H}_2$ ), 3.90 (dd, *J* = 12.4, 2.2 Hz, 1H, H-6a Glc), 3.87 – 3.81 (m, 2H, H-4 Gal, OC$\underline{H}_a$($CH_2$)$_4$), 3.75 – 3.67 (m, 3H , H-6b Glc, H-6a Gal, H-6b Gal), 3.67 – 3.63 (m, 1H, H-5 Gal), 3.62 – 3.54 (m, 4H, OC$\underline{H}_b$($CH_2$)$_4$, H-3 Gal, H-3 Glc, H-4 Glc), 3.52 – 3.49 (m, 1H, H-5 Glc), 3.47 (dd, *J* = 9.9, 7.8 Hz, 1H, H-2 Gal), 3.23 (t, *J* = 8.5 Hz, 1H, H-2 Glc), 3.05 (t, *J* = 6.8 Hz, 2H,

$O(CH_2)_4C\underline{H_2}$), 2.29 – 2.11 (m, 6H, 2x $C\underline{H}_aC{\equiv}C$, 2x $C\underline{H}_aCH_2C{\equiv}C$, 2x $C\underline{H}_bC{\equiv}C$), 1.62 – 1.53 (m, 2H, $OCH_2C\underline{H_2}(CH_2)_3$), 1.51 (d, *J* = 11.1 Hz, 2H, $C\underline{H}_bCH_2C{\equiv}C$), 1.45 (t, *J* = 7.5 Hz, 2H, $O(CH_2)C\underline{H_2}$), 1.32 (m, 3H, 2x $OCH_2C\underline{H}$ (BCN), $O(CH_2)_2C\underline{H_2}(CH_2)$), 0.91 (t, *J* = 9.8 Hz, 2H, 2x $OCH_2CHC\underline{H}$ (BCN)).
**$^{13}$C-NMR:** (150 MHz, $D_2O$) δ 158.8 (NC(O)O), 102.9 (C-1 Gal), 102.1 (C-1 Glc), 100.2 ($\underline{C}{\equiv}\underline{C}$), 78.4 (C-4 Glc), 75.3 (C-5 Gal), 74.8 (C-5 Glc), 74.4 (C-3 Glc), 72.8 (C-2 Glc), 72.5 (C-3 Gal), 70.9 (C-2 Gal), 70.4 ($O\underline{C}H_2(CH_2)_4$), 68.5 (C-4 Gal), 63.6 ($C(O)O\underline{C}H_2$), 61.0 (C-6 Gal), 60.1 (C-6 Glc), 40.2 ($(CH_2)_4\underline{C}H_2$), 28.6 ($O(CH_2)_4\underline{C}H_2$, 2x $\underline{C}H_2CH_2C{\equiv}C$), 28.4 ($OCH_2\underline{C}H_2(CH_2)_3$), 22.3 ($O(CH_2)_2\underline{C}H_2(CH_2)_2$), 20.8 (2x $\underline{C}H_2C{\equiv}C$), 19.8 (2x $OCH_2CH\underline{C}H$ (BCN)), 17.2 ($OCH_2\underline{C}H$ (BCN)).
HRMS (ESI-MS): m/z calculated for $C_{28}H_{46}NO_{13}$, $[M+H]^+$ 604.2964; measured 604.2960.

**Compound 20**

**20** was prepared (3.5 mg, 4.9 µmol) using the general procedure for synthesis of BCN linked glycans. After purification, **20** was obtained as a white solid (1.9 mg, 2.1 µmol, 44%).

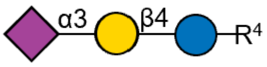

$^1$H (600 MHz, $D_2O$): δ (ppm)

| | H-1 | H-2 | H-3 | H-4 | H-5 | H-6 | H-7 | H-8 | H-9 | NHAc |
|---|---|---|---|---|---|---|---|---|---|---|
| Glucose | 4.46 (d, *J* = 7.8 Hz, 1H) | 3.49 | 4.03(dd, *J* = 9.9,3.1 Hz, 1H) | 3.89 | 3.63 | 3.57, 3.78 | - | - | - | - |
| Galactose | 4.39(d, *J* = 8.0 Hz, 1H) | 3.22 | 3.57 | 3.58 | 3.50 | 3.90, 3.73 | - | - | - | - |
| Neu5Ac | - | - | 2.68(dd, *J* = 12.4,4.6 Hz, 1H), 1.72(t, *J* = 12.1 Hz, 1H) | 3.62 | 3.77 | 3.57 | 3.52 | 3.81 | 3.67 | 1.95 (s, 3H) |

$^{13}$C (150 MHz, $D_2O$): δ (ppm)

| | C-1 | C-2 | C-3 | C-4 | C-5 | C-6 | C-7 | C-8 | C-9 | NHAc |
|---|---|---|---|---|---|---|---|---|---|---|
| Glucose | 102.72 | 69.36 | 75.48 | 67.43 | 75.16 | 62.59 | - | - | - | - |
| Galactose | 102.07 | 72.91 | 75.16 | 78.22 | 74.84 | 60.17 | - | - | - | - |
| Neu5Ac | n/a | n/a | 39.71 | 68.39 | 51.63 | 72.91 | 68.07 | 71.78 | 60.98 | 22.13 |

| Linker | 1 | 2 | 3 | 4 | 5 |
|---|---|---|---|---|---|
| H | 3.84, 3.61 | 1.57 (m, 2H) | 1.32 (m, 2H) | 1.57(m, 2H) | 3.06 (t, *J* = 6.7 |

| | | | | | |
|---|---|---|---|---|---|
| | | | | | Hz, 2H) |
| C | 70.49 | 28.43 | 22.31 | 28.43 | 40.35 |

| BCN | 6 | 7 | 8,8’ | 9,9’ | 10,10’ |
|---|---|---|---|---|---|
| H | 4.12 (d, *J* = 8.3 Hz, 2H) | 1.33 (m, 1H) | 0.91 (t, *J* = 10.5 Hz, 1H) | 2.19(2H), 1.57 (2H) | 2.22 – 2.15 (m, 4H) |
| C | 63.72 | 16.99 | 19.57 | 28.59 | 20.69 |

HRMS (ESI-MS): m/z calculated for $C_{39}H_{62}N_2O_{21}$ $[M-H]^-$: 893.3767; found: 893.3773.

#### Compound 26

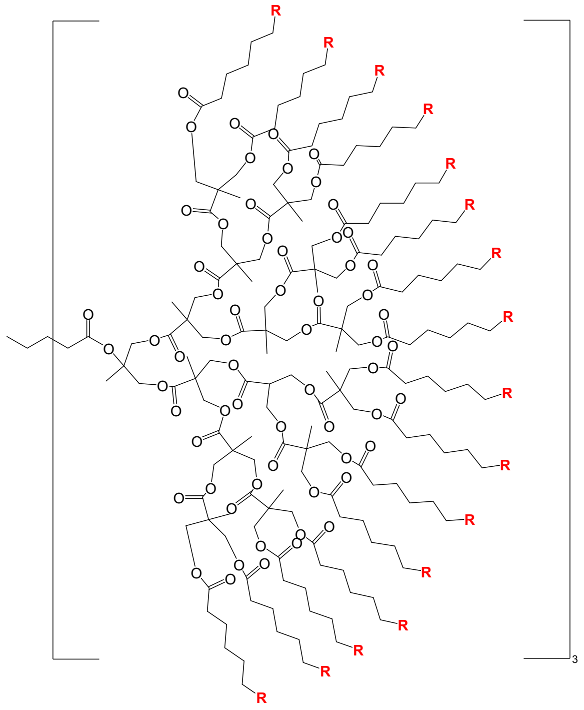

R= 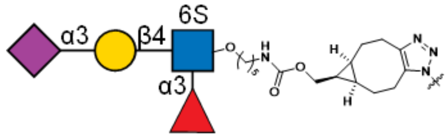

**26** was prepared using **21** (0.48 mg, 0.39 µmol) using the general procedure for synthesis of polymeric glycans. After purification, **26** was obtained as a white solid and glycan loading was determined to be 37% using the general procedure for glycopolymer characterization.

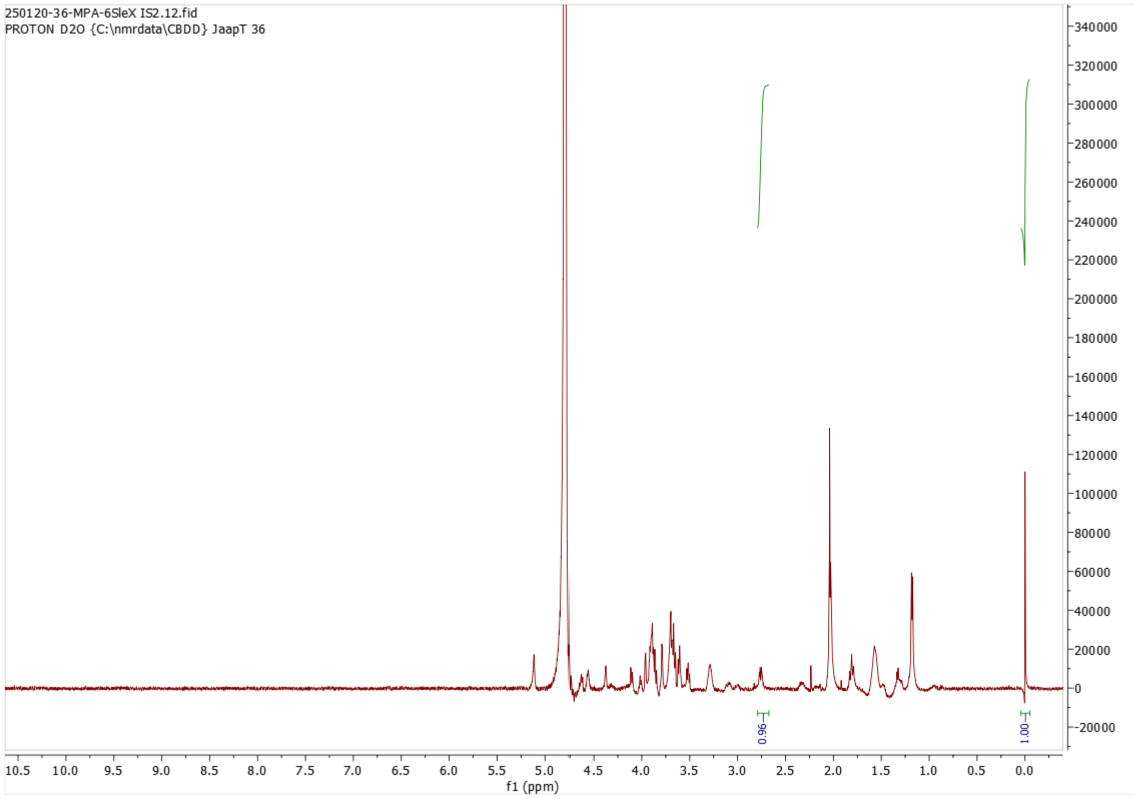

$^{1}$H NMR of **26**; 600MHz; $D_2O$

**Compound 28**

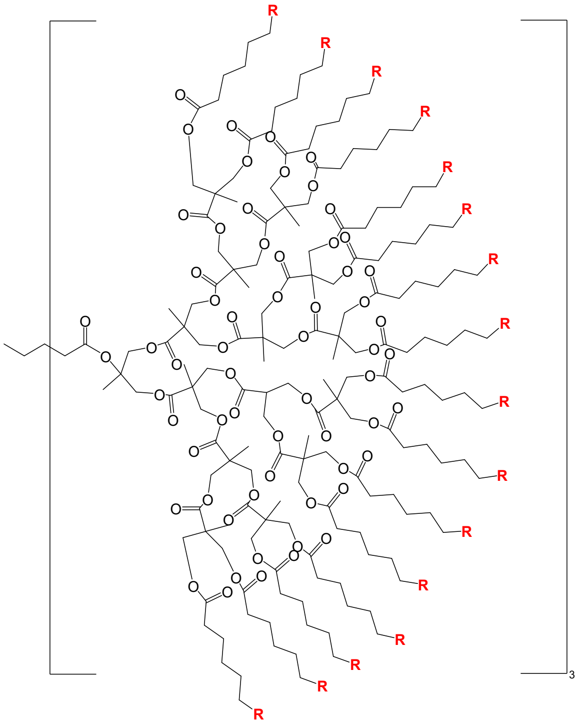

R= 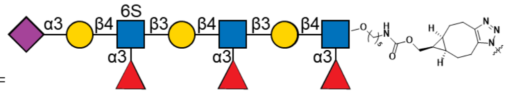

**28** was prepared using **23** (1.4 mg, 0.67 μmol) using the general procedure for synthesis of polymeric glycans. After purification, **28** was obtained as a white solid and glycan loading was determined to be 53% using the general procedure for glycopolymer characterization.

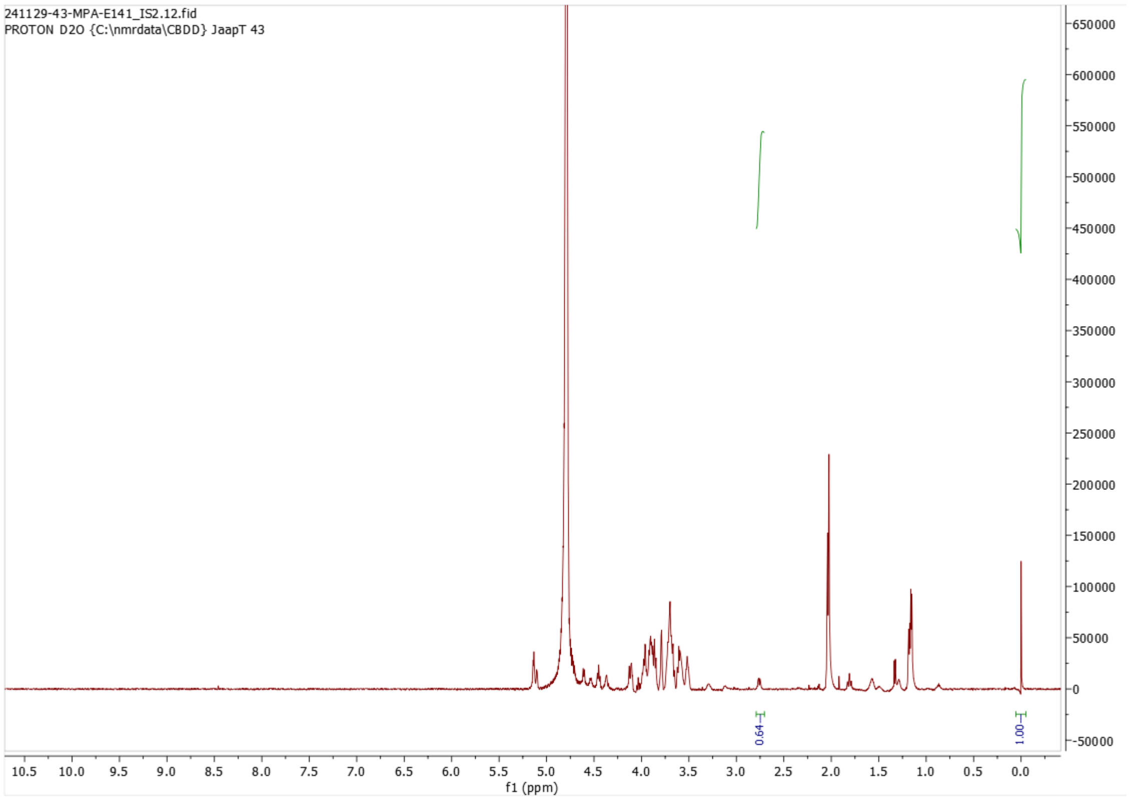

[1]H NMR of **28**; 600MHz; $D_2O$

**Compound 27**

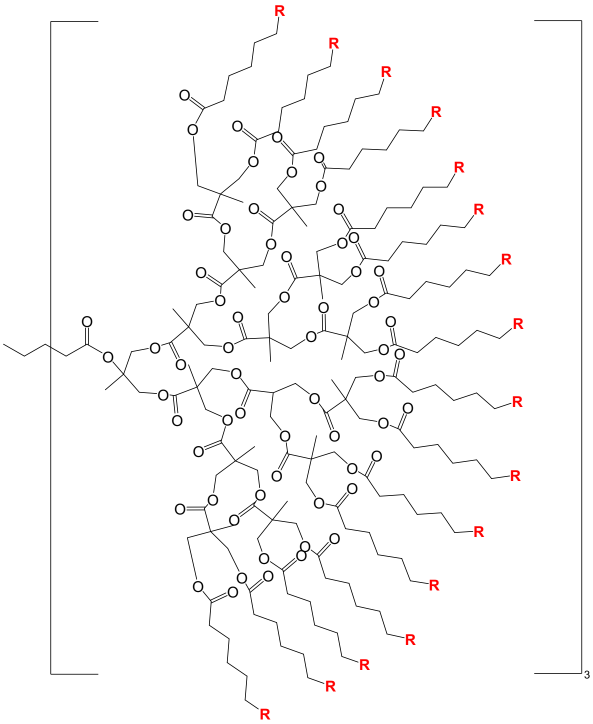

R=

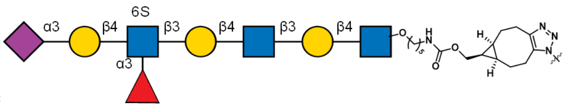

**27** was prepared using **22** (0.49 mg, 0.26 μmol) using the general procedure for synthesis of polymeric glycans. After purification, **27** was obtained as a white solid and glycan loading was determined to be 58% using the general procedure for glycopolymer characterization.

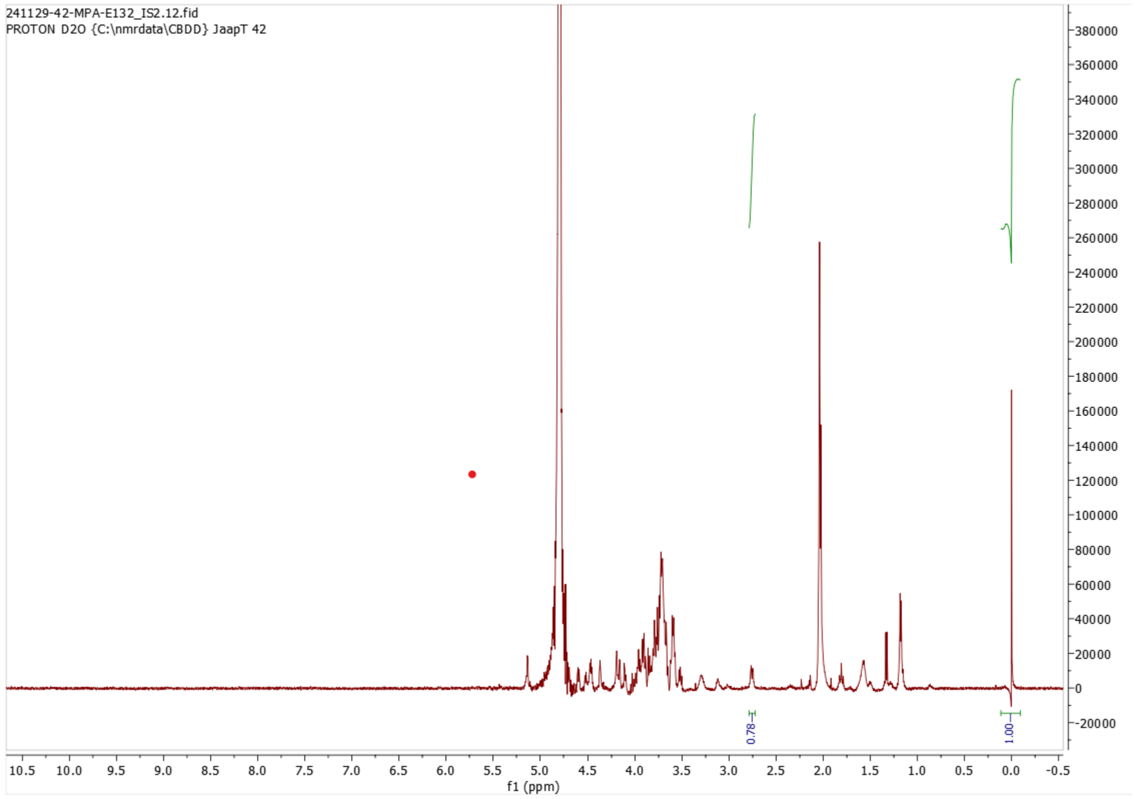

[1]H NMR of **27**; 600MHz; $D_2O$

**Compound 25**

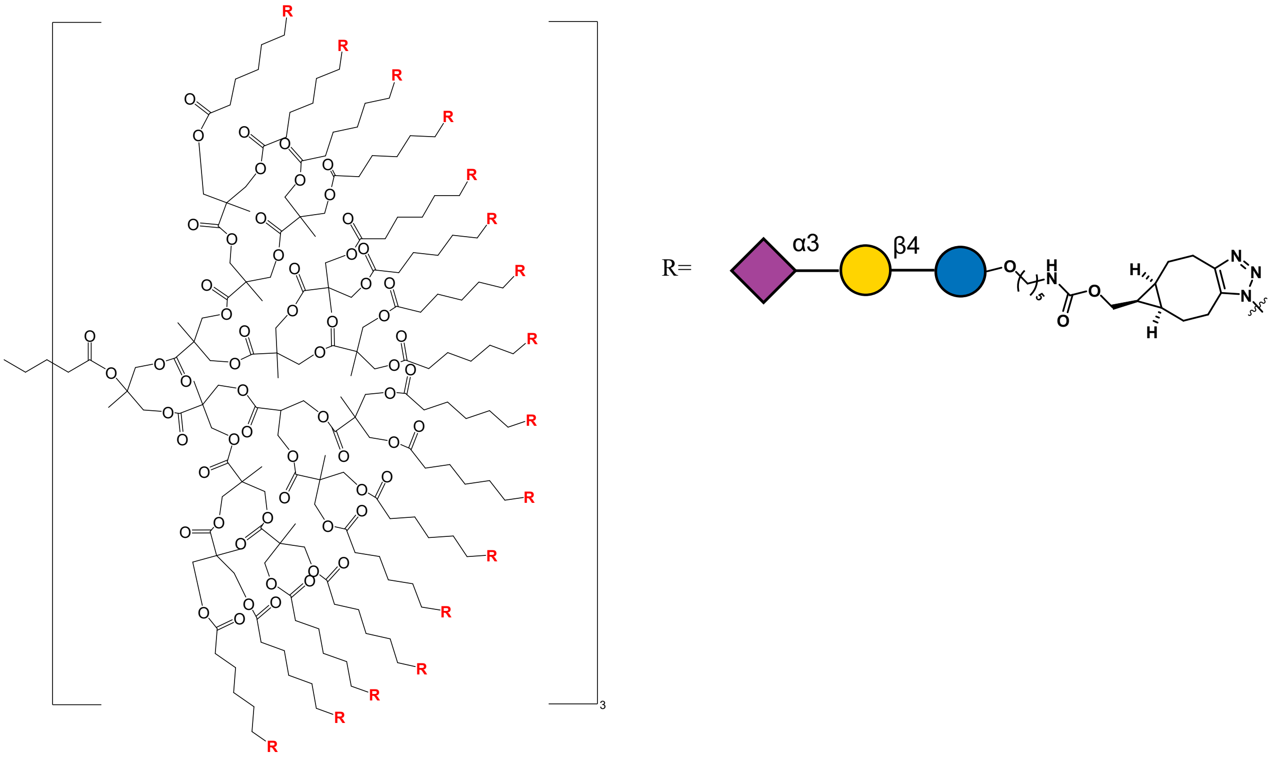

**25** was prepared using **20** (1.4 mg, 1.55 µmol) using the general procedure for synthesis of polymeric glycans. After purification, **25** was obtained as a white solid and glycan loading was determined to be 48% using the general procedure for glycopolymer characterization.

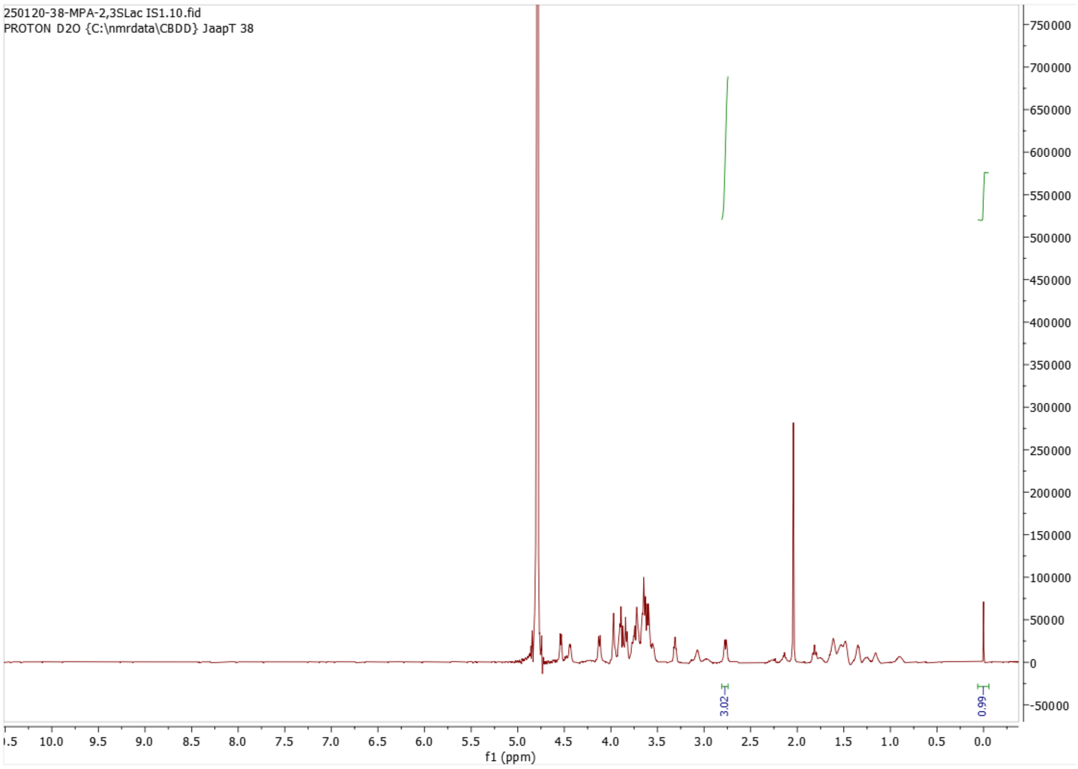

$^{1}$H NMR of **25**; 600MHz; $D_2O$

**Compound 24**

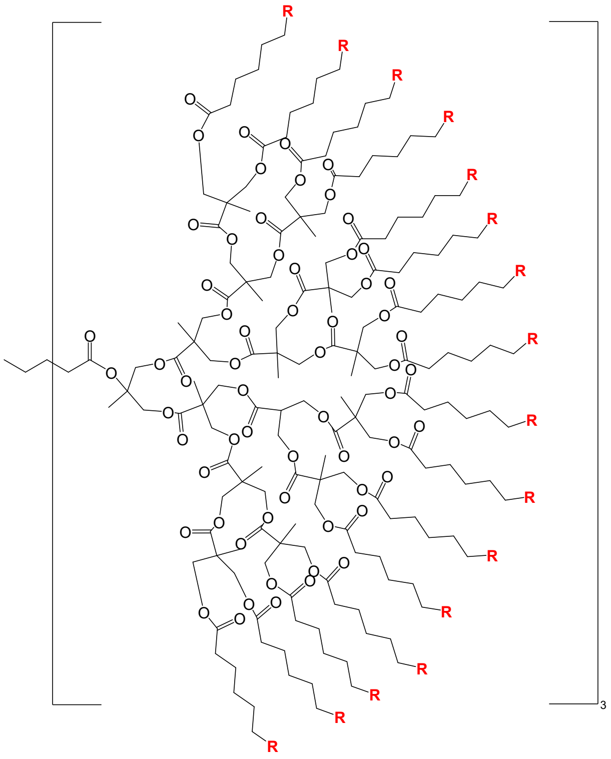

R=

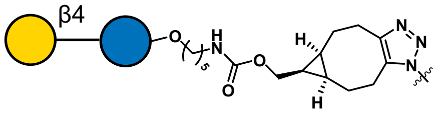

**24** was prepared using **19** (0.4 mg, 0.67 μmol) using the general procedure for synthesis of polymeric glycans. After purification, **24** was obtained as a white solid and glycan loading was determined to be 51% using the general procedure for glycopolymer characterization.

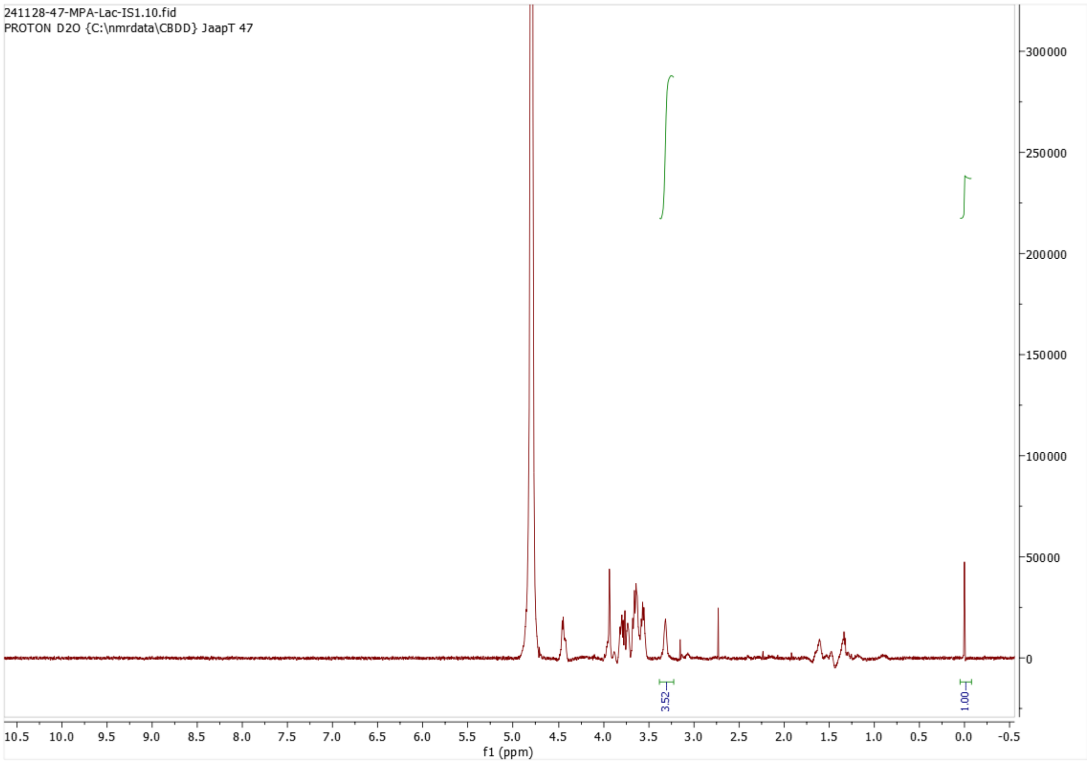

$^{1}H$ NMR of **24**; 600MHz; $D_2O$

#### 5) Microarray and Nanoparticles

##### Expression plasmid generation

A plasmid encoding recombinant MERS-CoV spike protein NTD (GenBank: KC164505; AA 18-351) was produced via Gibson assembly using cDNAs encoding codon-optimized open reading frames of the full-length MERS-CoV spike. The MERS-CoV NTD was cloned into a pcDNA5 expression vector in frame with a C-terminal human IgG1 Fc, a tobacco etch virus (TEV) cleavage site (ENLYFQG), a 6xHis-tag, and a Strep-tag (WSHPQFEK; IBA, Germany).[14] The pA-LS nanoparticle (NP) expression vector was created as described previously.[15-16] In short, domain B of protein A (pA; GenBank: M18264.1, AA 212-270) of *Staphylococcus aureus* was N-terminally fused to a Strep-tag (WSHPQFEK) and C-terminally to a Gly-Ser linker and 6,7-dimethyl-8-ribityllumazine synthase (LS; GenBank: AAC06489.1, AA 1-154) of *Aquifex aeolicus*. The pA-LS sequence was constructed in a pUC57 plasmid by GenScript USA, Inc. and later ligated into the pCD5 expression vector via NheI/NotI restriction sites. pcDNA5-Siglec-9-Fc was a gift from Dr. Matthew Macauley (University of Alberta, Edmonton, AB, Canada).[14]

##### Protein expression and purification

Proteins were expressed by transfecting HEK293S GnTI$^-$ cells with polyethyleneimine hydrochloride (PEI-HCl) as previously described.[16] Briefly, cells were grown to 60% confluency in Dulbecco's Modified Eagle Medium (DMEM; Gibco) supplemented with 10% fetal calf serum (FCS; Sigma), 25 units/mL of penicillin and 0.025 mg/mL streptomycin (Sigma). Expression vectors were incubated with PEI-HCl in a 1:8 ratio (μg DNA/μg PEI) and DMEM for 20 min before addition to the cells. At 6 h post-transfection, the medium was replaced with 293 SFM II medium (Gibco) supplemented with Primatone 3.0 g/L (Kerry), bicarbonate 3.6 g/L, glucose 2.0 g/L, valproic acid 0.4 g/L, glutaMAX 1% (Gibco), and DMSO 1.5%. Supernatants were collected after 5 days of incubation at 37 °C and 5% $CO_2$. Protein expression was analyzed with SDS-PAGE and subsequent Western blotting using StrepMAB-Classic HRP 1:3000 (2-1509-001, IBA). Proteins were purified using Strep-Tactin Sepharose beads (2-1201-002, IBA) according to the manufacturers protocol. Proteins were checked on Coomassie blue-stained SDS-PAGE gel.

##### Glycan microarray

All compounds were printed on NHS-ester activated glass slides (NEXTERION Slide H, Schott Inc.) using a Scienion sciFLEXARRAYER S3 non-contact microarray equipped with a Scienion PDC80 nozzle (Scienion Inc.).[17] Individual compounds were dissolved in sodium phosphate buffer (250 mM, pH 8.5) at a concentration of 100 µM and were printed in replicates of 6 with spot volume ~400 pL, at 20 °C and 50% humidity. Each slide has 24 subarrays in a 3x8 layout. After printing, slides were incubated in a humidity chamber for 8 h and then blocked for 1 h with a 50 mM ethanolamine in Tris buffer (pH 9.0, 100 mM) at 50 °C. Blocked slides were rinsed with DI water (50 mL), spun dry, and kept in a desiccator at room temperature for future use.

MERS-NTD-Fc and Siglec-9-Fc proteins were conjugated at 100 µg/mL with pA-LS NP in a 1:1 molar ratio and incubated overnight at 4 ºC. All proteins were prepared in a final volume of 50 µL PBS-T. Proteins were incubated on the array surface in a humidified chamber for 90 min. Slides were washed with PBS-T (PBS + 0.1% Tween (50 mL)), PBS (50 mL), and deionized water (2x) (50 mL) before being dried by centrifugation. Upon washing, slides incubated with MERS-NTD-Fc or Siglec-9-Fc were incubated with 5 µg/mL of primary antibody StrepMAB-Classic HRP precomplexed with 2.5 µg/mL of secondary antibody goat anti-mouse IgG Alexa Fluor 555 in PBS-T for 1 h and the slides were washed again.

Viral isolates diluted in PBS-T (PBS + 0.1% Tween) to an HA titer of 128 were applied to subarrays in the presence of oseltamivir (200 nM, in 100 µL) in a humidified chamber for 1 h. Next, the microarray slide was rinsed with PBS-T (PBS + 0.1% Tween (50 mL)), PBS (50 mL), and deionized water (2x) (50 mL), and dried by centrifugation. The slide was incubated for 1 h in the presence of the CR6261 stem specific antibody (100 µL, 5 µg/mL in PBS-T) and washed as described above. A secondary goat anti-human AlexaFluor-647 antibody (100 µL, 2 µg/mL in PBS-T) was applied and the resulting slide was incubated for 1 h in a humidified chamber and then washed by the standard procedure.

After protein or virus incubations, the slides were rinsed successively with PBS-T (50 mL), PBS (50 mL), and deionized water (50 mL). Slides were dried by centrifugation after the washing steps and scanned immediately using an Innopsys Innoscan 710 microarray scanner. Various gains and PMT values were used to ensure the signals were in the linear range and to avoid saturation of the signals. Images were analyzed with Mapix software (version 8.1.0 Innopsys) and processed with an Excel macro (https://github.com/enthalpyliu/carbohydrate-microarray-processing). The average fluorescence intensity and SD were determined for each compound after removal of the highest and lowest intensities from the spot replicates to give n = 4.

Recombinant viruses that contained the HA (without a multibasic cleavage site) of interest in the background of the attenuated vaccine strain A/Puerto Rico/8/1934 (PR/8) were handled under biosafety level (BSL) 2 conditions in agreement with national regulations on genetically modified viruses (GMO-99–090 and IG 15–160_IIk). All viruses in this study are thus recombinant, but we do refer to them with the full virus name throughout the manuscript. The viruses on the array were used with an HA titer of 128.

Figure 2a in the manuscript shows a list of glycans on the microarray.

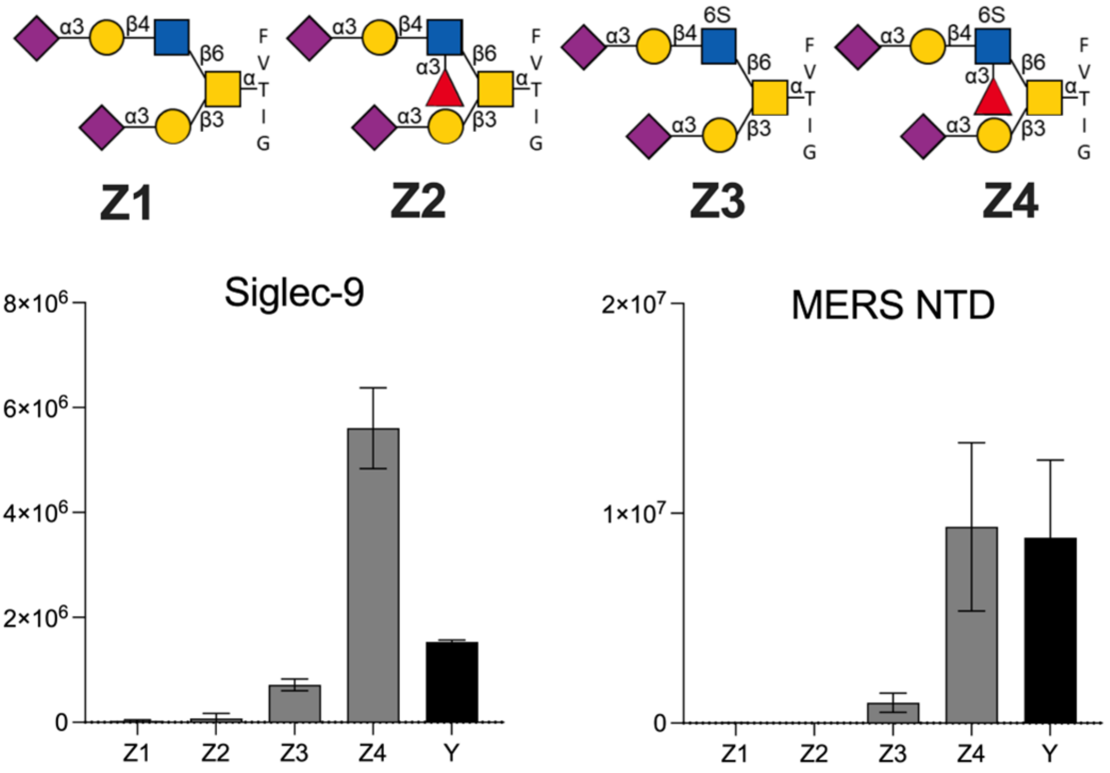

**Figure S2.** Probing the binding properties of Siglec-9 and MERS-NTD-Fc to 6-sulfo-SLe$^x$ epitope presented on O-Glycan backbone.

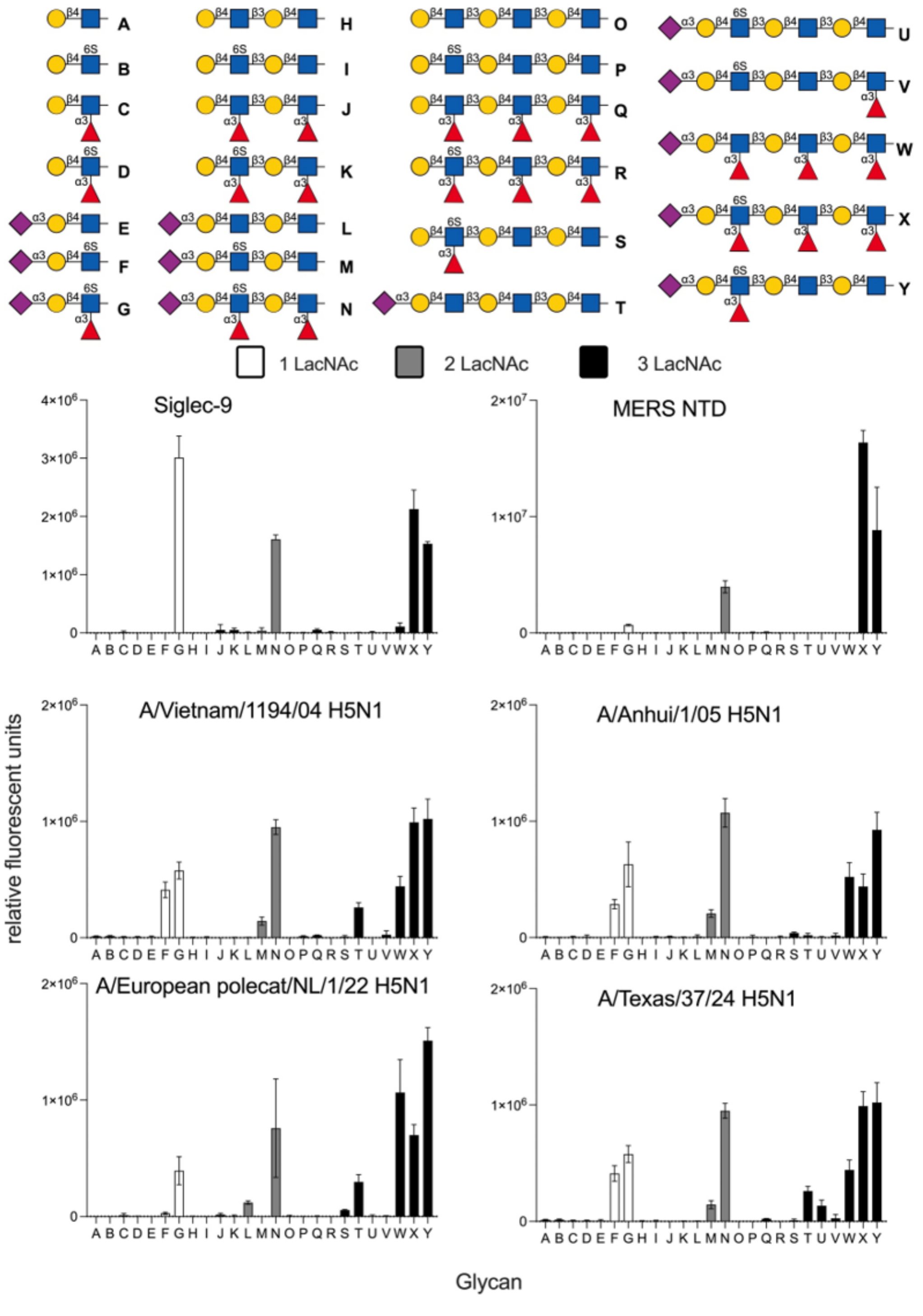

**Figure S3.** Probing different backbone length of 6-sulfo-SLe$^x$ epitope binding properties of Siglec-9, MERS-NTD and HAs of Recombinant H5N1 influenza viruses.

#### Hemagglutination assay

Hemagglutination assays were performed with 100 µL of MERS-NTD-Fc at a starting concentration of 20 µg/mL. MERS-NTD-Fc was conjugated to pA-LS NP at a 1:1 molar ratio for 30 min on ice. Conjugated protein was 2-fold serially diluted and incubated with 50 µL of

1% human erythrocytes in V-bottom 96-well plates. Hemagglutination titers were calculated as the highest dilution of MERS-NTD-Fc agglutinating human erythrocytes.

**Hemagglutination inhibition assay**

HAI assay was performed to determine whether 6-sulfo-SLe$^x$ ligands can inhibit binding of MERS-NTD-Fc to human erythrocytes. Briefly, serial 2-fold dilutions starting at 10 µM of the lactose, α2,3-sialyllactose, and 6-sulfo-SLe$^x$ ligands, all conjugated to polyglycerol-based NPs, were added onto the plate. The NP-conjugated glycans were incubated with 25 µL of 4 HA units of pA-LS-conjugated MERS-NTD-Fc for 1 h, after which 50 µL of 1% human erythrocytes were added. HAI titers were calculated as the highest dilution of NP-conjugated ligand that gave inhibition of agglutination.

An NP with lactose was taken along as negative control.

**Calculation of Minimum Inhibitory Concentration (MIC)**

| | | | | | | | | | | |
|---|---|---|---|---|---|---|---|---|---|---|
| **Well** | 1 | 2 | 3 | 4 | 5 | **6** | 7 | 8 | 9 | 10 |
| **Titer** | 2 | 4 | 8 | 16 | 32 | **64** | 128 | 256 | 512 | 1024 |
| **Molarity compound (mM)** | 0.01 | 0.005 | 0.0025 | 0.00125 | 0.000625 | **0.0003125** | 0.000156 | 7.81E-05 | 3.91E-05 | 1.95E-05 |
| **nM** | 10,000 | 5000 | 2500 | 1250 | 625 | **312.5** | 156 | 78.1 | 39.1 | 19.5 |

Basically, the starting molarity of the compounds is 0.01 mM in the first well. Therefore, by using 2-fold dilutions, the amount of compound in the 6th well is 312.5 nM. The 6$^{th}$ well is the well with an HAI titer of 64.

#### 6) Computational Approach

The 3D structure of the 6-sulfo-$SLe^x(LacNAc)_2$ ligand **4** was built using the carbohydrate builder tool from the glycam website: glycam.org.[18] The 3D structure of MERS-CoV S protein was derived from the available crystal structure (pdb code: 6Q05) of the viral protein in complex with $SLe^x$. The 3D structure of the 6-sulfo-$SLe^x$ $(LacNAc)_2$ was placed into the binding pocket of MERS-CoV S protein by molecular alignment with the $SLe^x$ of the cocrystal structure. The resulting binding pose was energy minimized using the Macromodel minimization tool of Maestro Schrödinger suite of program Maestro version 14.2.118. Three consecutive minimization stages were performed involving (1) only the ligand, (2) only the protein residues within the ligand binding site, and (3) the whole system using OPLS4 force field in water with an extended cutoff of 8.0 Å for Van der Waals, 20.0 Å for Electrostatic, and 4.0 Å for hydrogen bond interactions. A low gradient convergence threshold (0.05) in 5000 steps was applied. The obtained geometry of the complex was further fully minimized (the ligand, the protein, and water molecules) with the AMBER* force field (Figure S4). The results from the docking were validated by analysis of the ligand and complex structures. First, the ring conformation of each monosaccharide composing the ligand were scrutinized. As expected, the Galactose and the N-Acetyl-Glucosamine rings adopts the $^4C_1$ conformation, while the Fucose and the Neuraminic acid rings adopts the alternative $^1C_4$ conformation (Figure S5). Next, the f and Y dihedral angles at the glycosidic linkages were measured. f was defined as f: $O_5C_1O_xC_x$ and Y: $C_1O_xC_xC_{(x+1)}$ for all the sugars but for Neu5Ac for which f: $O_6C_2O_xC_x$ and Y: $C_2O_xC_xC_{(x+1)}$. In all the cases, the *exo*-anomeric effect was respected. The resulting values are reported in Table S1. Validation of the protein/ligand binding pose was performed by measuring interatomic distances between interacting residues. In particular, the distances between the ligand moieties and specific aminoacidic residues discussed in the main text have been measured and are reported in Figure S6. Finally, an overlap of the docking derived with the X-ray structure is reported in Figure S7. The figures were generated using the molecular graphic software PyMOL (The PyMOL Molecular Graphics System, Version 2.4 Schrödinger, LLC, http://www.pymol.org) and the Maestro Schrödinger suite of program Maestro version 14.2.118.

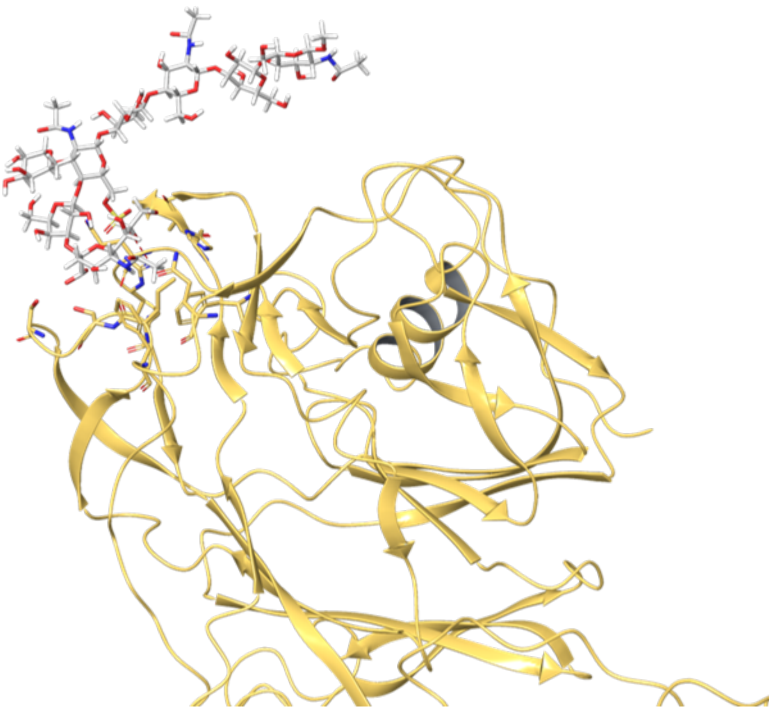

**Figure S4.** A full picture of structural basis of MERS-CoV S selectivity for the 6-sulfo-SLe$^{x}$ receptor.

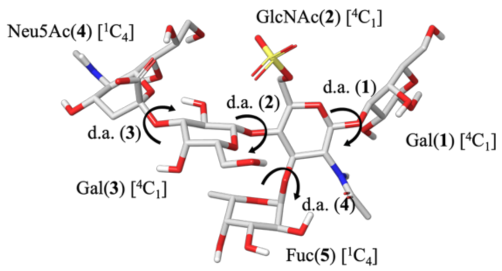

Table: Inter-residues glycosidic dihedral angles

| Glycosidic Linkage | GlcNAc(**2**)-Gal(**1**) | Gal(**3**)-GlcNAc(**2**) | Neu5Ac(**4**)-Gal(**3**) | Fuc(**5**)-GlcNAc(**2**) |
|---|---|---|---|---|
| ϕ | -73.5 | -70.0 | 68.4 | -68.8 |
| Ψ | 132.4 | -104.6 | 143.6 | 140.1 |
| | ϕ: O5C1OxCx | | Ψ: C1OxCxCx+1 | |

**Figure S5 and Table 1.** Conformation of the ligand bound in the MERS-CoV S protein binding pocket. All monosaccharide residues of the interacting pentasaccharide adopt the most stable chair conformation. The glycosidic dihedral angles are indicated as "d.a." followed by the corresponding residue number. Table S1 reports the values of the ϕ and Ψ dihedral angles for each glycosidic linkage.

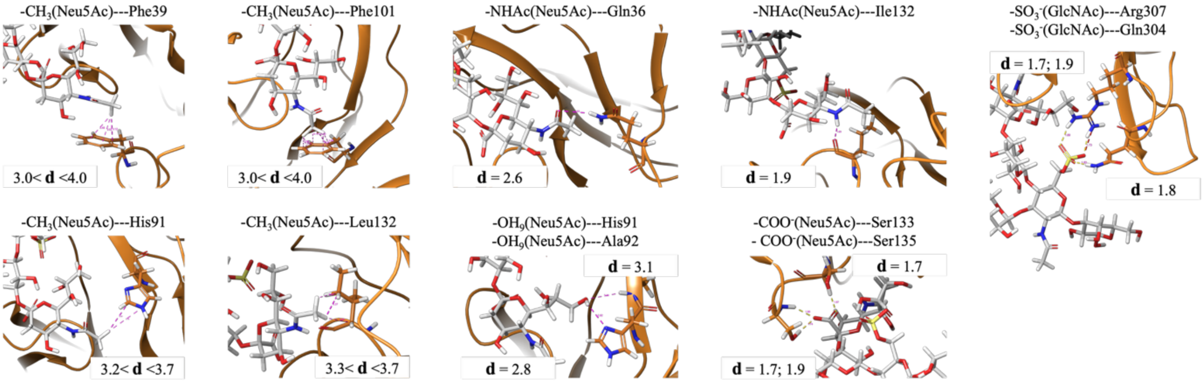

**Figure S6.** Calculated interatomic distances (**d**) between the ligand and interacting protein residues.

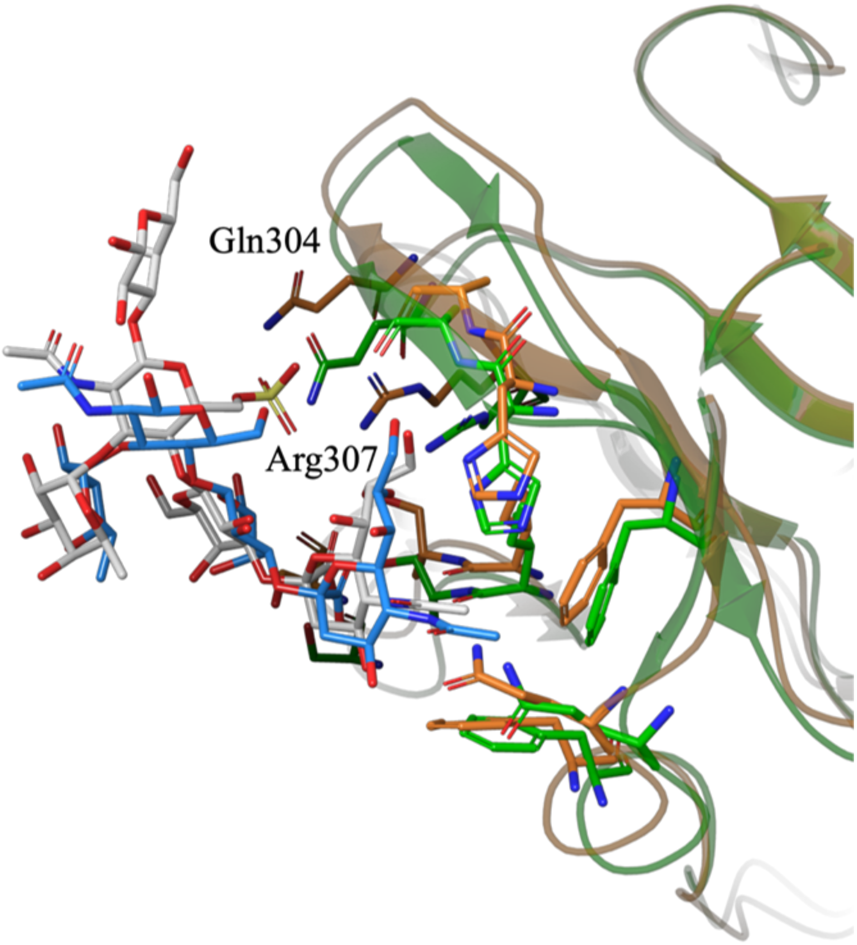

**Figure S7.** Overlay of the docking-derived structure of the MERS-CoV S protein in complex with 6-sulfo-SLe$^x$ (protein shown in orange and ligand in gray) and the crystal structure of the viral protein in complex with SLe$^x$ (protein shown in green and ligand in blue). The overall binding modes are highly similar, with the most notable difference being the orientations of Gln304 and Arg307. In the docking-derived complex, these residues adopt conformations that enable interactions with the sulfate group of 6-sulfo-SLe$^x$.

#### 7) References

1 H. Yu, S. Huang, H. Chokhawala, M. C. Sun, H. Zheng, and X. Chen "Highly Efficient Chemoenzymatic Synthesis of Naturally Occurring and Non-Natural alpha-2,6-Linked Sialosides: A *P. Damsela* alpha-2,6-Sialyltransferase with Extremely Flexible Donor-Substrate Specificity," *Angew. Chem. Int. Ed.* 45 (2006): 3938-3944. https://doi.org/10.1002/anie.200600572

2 W. J. Peng, J. Pranskevich, C. Nycholat, M. Gilbert, W. Wakarchuk, J. C. Paulson, and N. Razi "*Helicobacter Pylori* Beta 1,3-*N*-Acetylglucosaminyltransferase for Versatile Synthesis of Type 1 and Type 2 Poly-Lacnacs on *N*-Linked, *O*-Linked and I-Antigen Glycans," *Glycobiology* 22 (2012): 1453-1464. https://doi.org/Doi 10.1093/Glycob/Cws101

3 G. Sugiarto, K. Lau, J. Qu, Y. Li, S. Lim, S. Mu, J. B. Ames, A. J. Fisher, and X. Chen "A Sialyltransferase Mutant with Decreased Donor Hydrolysis and Reduced Sialidase Activities for Directly Sialylating Lewis$^{x}$," *ACS Chem. Biol.* 7 (2012): 1232-1240. https://doi.org/10.1021/cb300125k

4 Y. Li, M. Xue, X. Sheng, H. Yu, J. Zeng, V. Thon, Y. Chen, M. M. Muthana, P. G. Wang, and X. Chen "Donor Substrate Promiscuity of Bacterial β1–3-N-Acetylglucosaminyltransferases and Acceptor Substrate Flexibility of β1–4-Galactosyltransferases," *Bioorg. Med. Chem.* 24 (2016): 1696-1705. https://doi.org/https://doi.org/10.1016/j.bmc.2016.02.043

5 A. R. Prudden, L. Liu, C. J. Capicciotti, M. A. Wolfert, S. Wang, Z. Gao, L. Meng, K. W. Moremen, and G. J. Boons "Synthesis of Asymmetrical Multiantennary Human Milk Oligosaccharides," *Proc. Natl. Acad. Sci. U. S. A.* 114 (2017): 6954-6959. https://doi.org/10.1073/pnas.1701785114

6 J. B. McArthur, H. Yu, N. Tasnima, C. M. Lee, A. J. Fisher, and X. Chen "Alpha2-6-Neosialidase: A Sialyltransferase Mutant as a Sialyl Linkage-Specific Sialidase," *ACS Chem. Biol.* 13 (2018): 1228-1234. https://doi.org/10.1021/acschembio.8b00002

7 Y. Wu, G. P. Bosman, D. Chapla, C. Huang, K. W. Moremen, R. P. de Vries, and G. J. Boons "A Biomimetic Synthetic Strategy Can Provide Keratan Sulfate I and II Oligosaccharides with Diverse Fucosylation and Sulfation Patterns," *J. Am. Chem. Soc.* 146 (2024): 9230-9240. https://doi.org/10.1021/jacs.4c00363

8 Y. Wu, G. M. Vos, C. Huang, D. Chapla, A. L. M. Kimpel, K. W. Moremen, R. P. de Vries, and G. J. Boons "Exploiting Substrate Specificities of 6-*O*-Sulfotransferases to

Enzymatically Synthesize Keratan Sulfate Oligosaccharides," *JACS Au* 3 (2023): 3155-3164. https://doi.org/10.1021/jacsau.3c00488

9 W. Wang, T. Hu, P. A. Frantom, T. Zheng, B. Gerwe, D. S. Del Amo, S. Garret, R. D. Seidel, III, and P. Wu "Chemoenzymatic Synthesis of GDP-L-Fucose and the Lewis X Glycan Derivatives," *Proc. Natl. Acad. Sci. U. S. A.* 106 (2009): 16096-16101. https://doi.org/10.1073/pnas.0908248106

10 L. Liu, A. R. Prudden, C. J. Capicciotti, G. P. Bosman, J. Y. Yang, D. G. Chapla, K. W. Moremen, and G. J. Boons "Streamlining the Chemoenzymatic Synthesis of Complex *N*-Glycans by a Stop and Go Strategy," *Nat. Chem.* 11 (2019): 161-169. https://doi.org/10.1038/s41557-018-0188-3

11 L. Meng, F. Forouhar, D. Thieker, Z. Gao, A. Ramiah, H. Moniz, Y. Xiang, J. Seetharaman, S. Milaninia, M. Su, R. Bridger, L. Veillon, P. Azadi, G. Kornhaber, L. Wells, G. T. Montelione, R. J. Woods, L. Tong, and K. W. Moremen "Enzymatic Basis for *N*-glycan Sialylation: Structure of Rat Alpha2,6-Sialyltransferase (St6gal1) Reveals Conserved and Unique Features for Glycan Sialylation," *J. Biol. Chem.* 288 (2013): 34680-34698. https://doi.org/10.1074/jbc.M113.519041

12 K. W. Moremen, A. Ramiah, M. Stuart, J. Steel, L. Meng, F. Forouhar, H. A. Moniz, G. Gahlay, Z. Gao, D. Chapla, S. Wang, J. Y. Yang, P. K. Prabhakar, R. Johnson, M. D. Rosa, C. Geisler, A. V. Nairn, J. Seetharaman, S. C. Wu, L. Tong, H. J. Gilbert, J. LaBaer, and D. L. Jarvis "Expression System for Structural and Functional Studies of Human Glycosylation Enzymes," *Nat. Chem. Biol.* 14 (2018): 156-162. https://doi.org/10.1038/nchembio.2539

13 G. M. Vos, PhD thesis, Utrecht University (Utrecht University Repository), **2023**.

14 E. Rodrigues, J. Jung, H. Park, C. Loo, S. Soukhtehzari, E. N. Kitova, F. Mozaneh, G. Daskhan, E. N. Schmidt, V. Aghanya, S. Sarkar, L. Streith, C. D. St Laurent, L. Nguyen, J. P. Julien, L. J. West, K. C. Williams, J. S. Klassen, and M. S. Macauley "A Versatile Soluble Siglec Scaffold for Sensitive and Quantitative Detection of Glycan Ligands," *Nat. Commun.* 11 (2020): 5091. https://doi.org/10.1038/s41467-020-18907-6

15 W. Li, R. J. G. Hulswit, I. Widjaja, V. S. Raj, R. McBride, W. Peng, W. Widagdo, M. A. Tortorici, B. van Dieren, Y. Lang, J. W. M. van Lent, J. C. Paulson, C. A. M. de Haan, R. J. de Groot, F. J. M. van Kuppeveld, B. L. Haagmans, and B. J. Bosch "Identification of Sialic Acid-Binding Function for the Middle East Respiratory

Syndrome Coronavirus Spike Glycoprotein," *Proc. Natl. Acad. Sci. U. S. A.* 114 (2017): E8508-E8517. https://doi.org/10.1073/pnas.1712592114

16 I. Tomris, A. L. M. Kimpel, R. Liang, R. van der Woude, G. P. H. Boons, Z. Li, and R. P. de Vries "The HCoV-HKU1 N-Terminal Domain Binds a Wide Range of 9-*O*-Acetylated Sialic Acids Presented on Different Glycan Cores," *ACS Infect. Dis.* 10 (2024): 3880-3890. https://doi.org/10.1021/acsinfecdis.4c00488

17 F. Broszeit, R. J. van Beek, L. Unione, T. M. Bestebroer, D. Chapla, J. Y. Yang, K. W. Moremen, S. Herfst, R. A. M. Fouchier, R. P. de Vries, and G. J. Boons "Glycan Remodeled Erythrocytes Facilitate Antigenic Characterization of Recent a/H3N2 Influenza Viruses," *Nat. Commun.* 12 (2021): 5449. https://doi.org/10.1038/s41467-021-25713-1

18 O. C. Grant, D. Wentworth, S. G. Holmes, R. Kandel, D. Sehnal, X. Wang, Y. Xiao, P. Sheppard, T. Grelsson, A. Coulter, G. Miller, B. L. Foley, and R. J. Woods "Generating 3d Models of Carbohydrates with Glycam-Web," *bioRxiv* (2025). https://doi.org/10.1101/2025.05.08.652828

#### 8) NMR Spectra, LC-MS Data and HPLC Traces

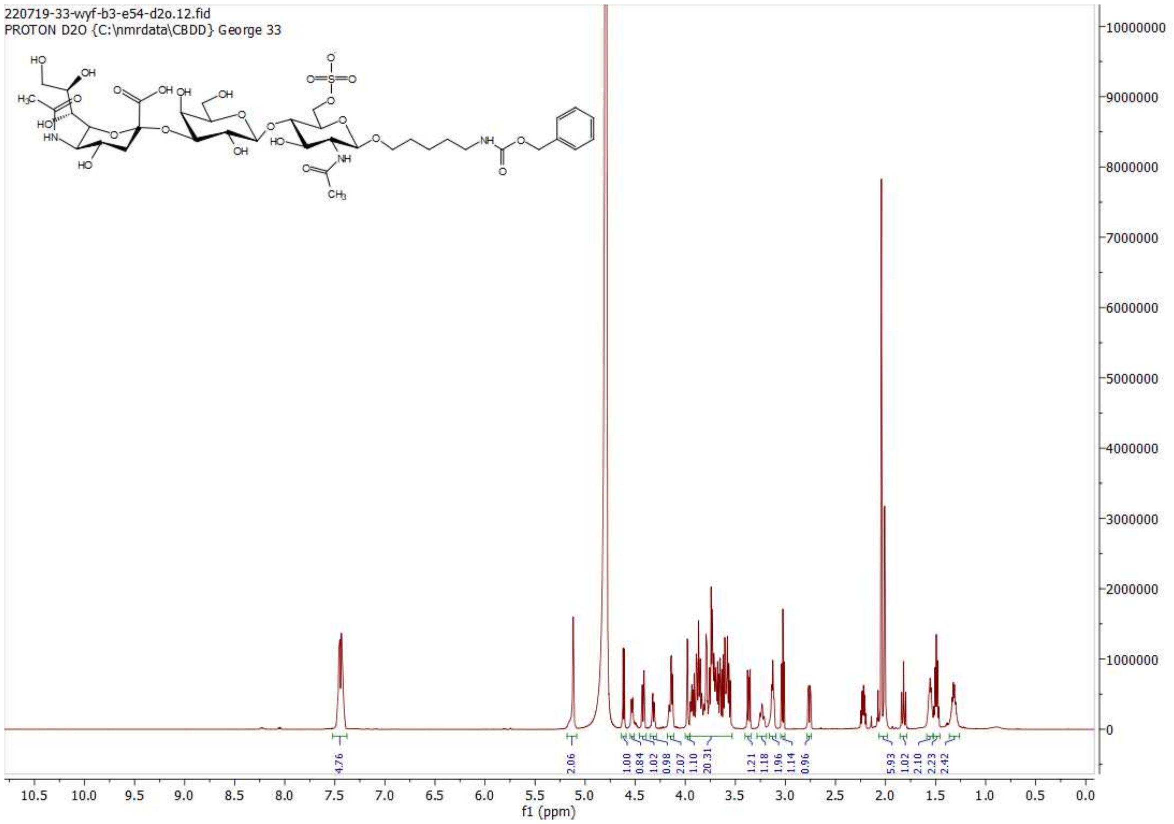

$^1$H NMR of 15; 600MHz; $D_2O$

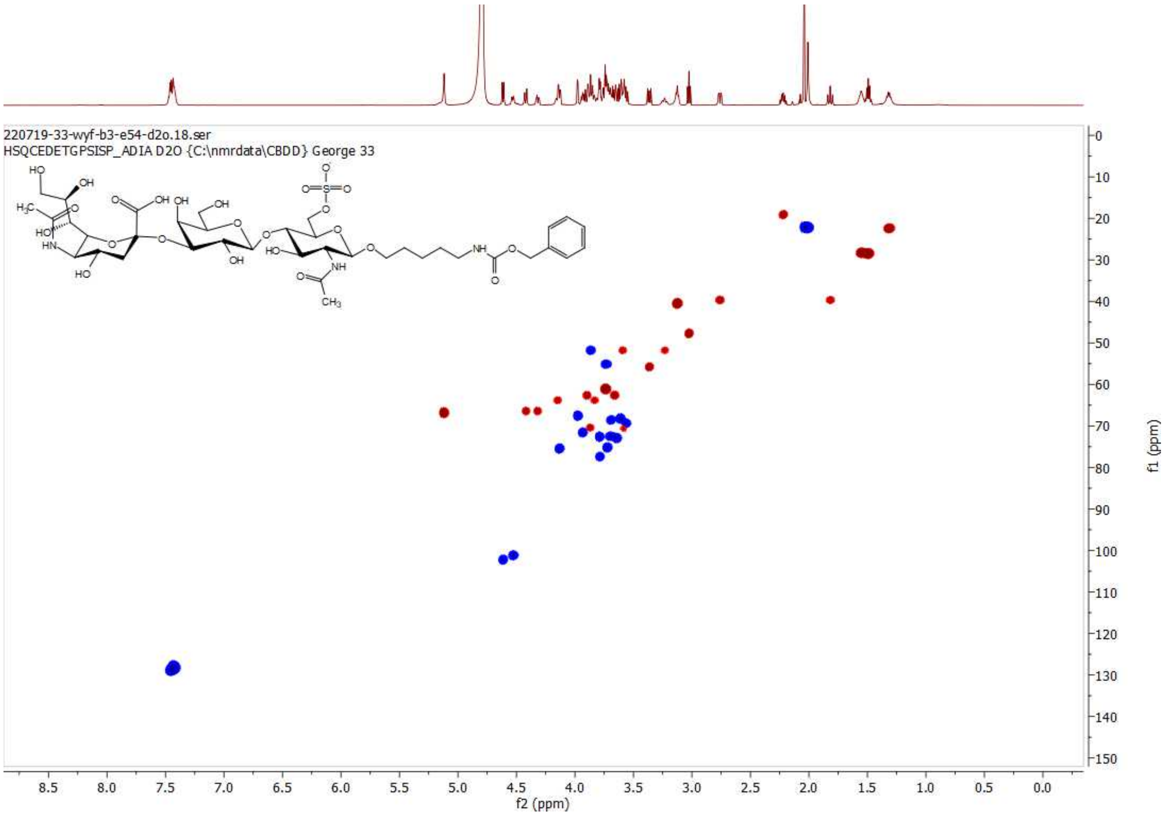

HSQC of 15; 600 MHz/150 MHz, $D_2O$

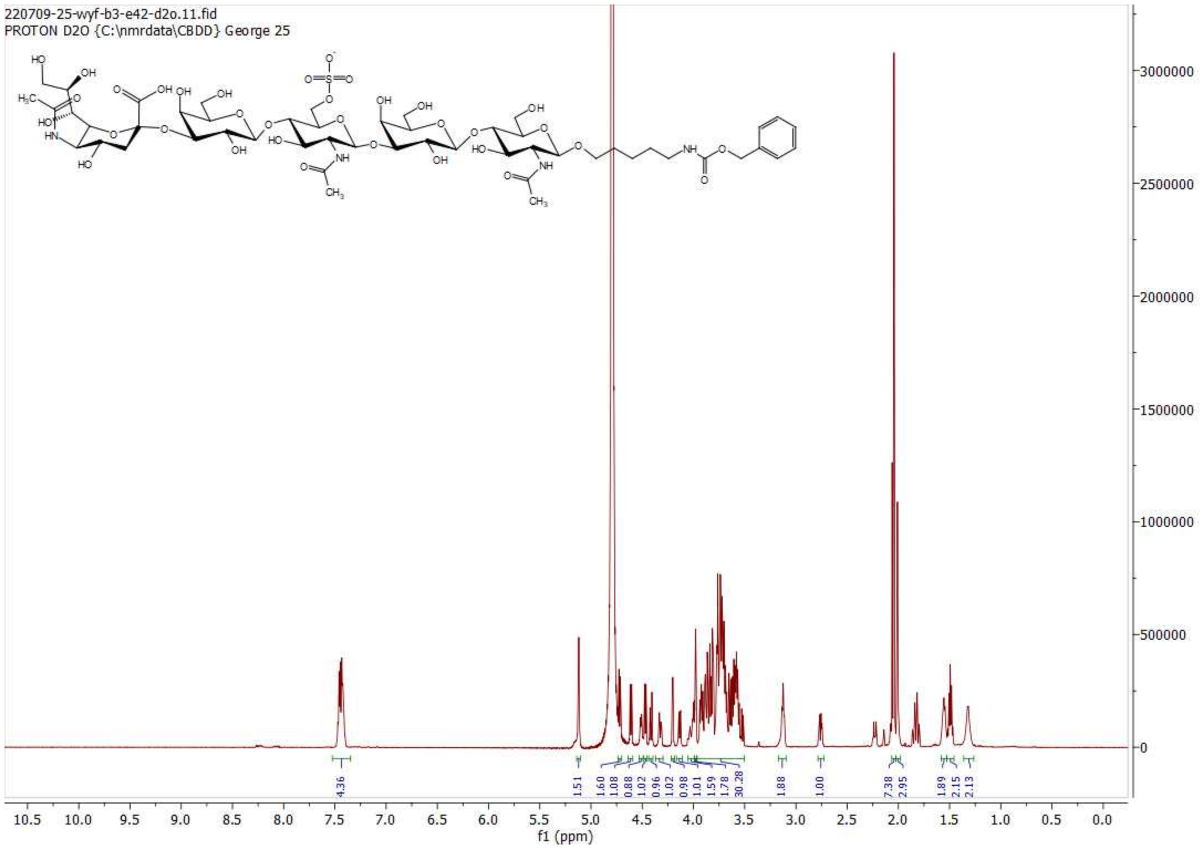

$^1$H NMR of 16; 600MHz; $D_2O$

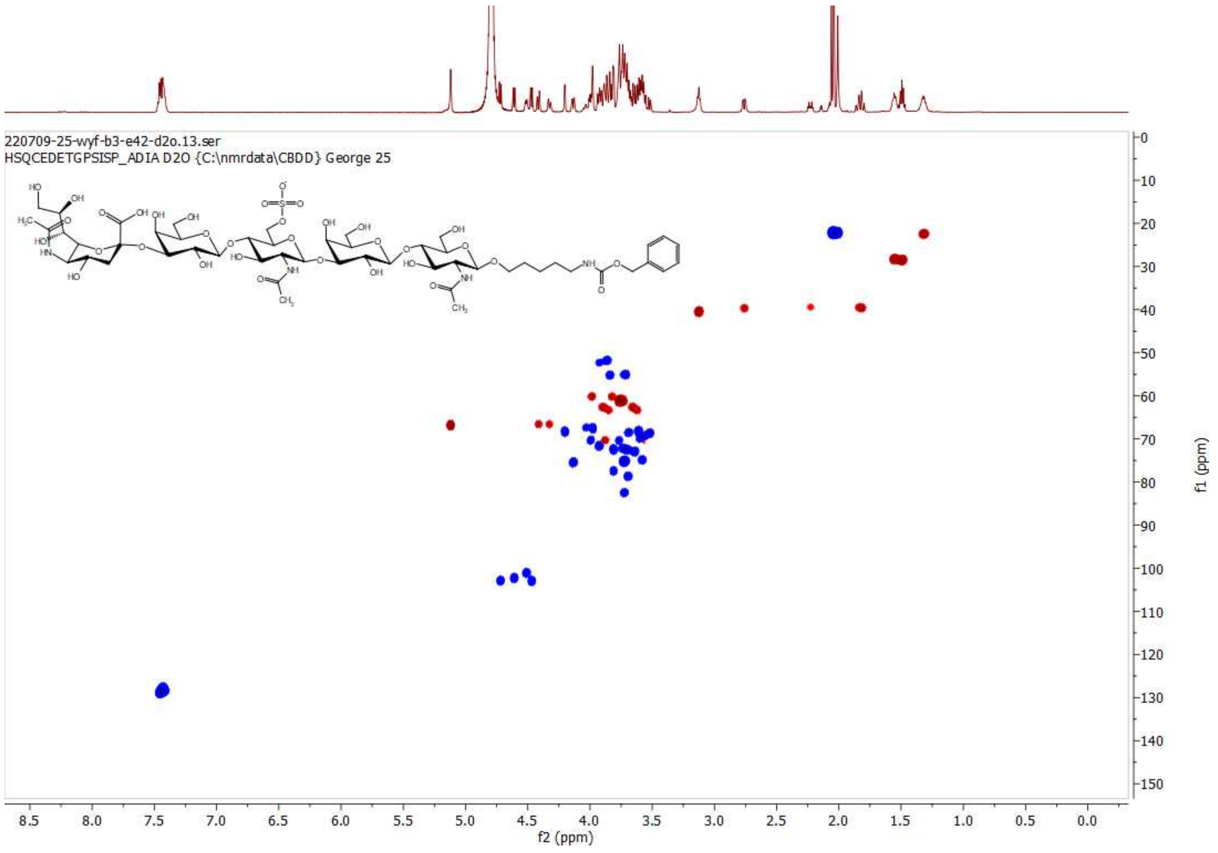

HSQC of 16; 600 MHz/150 MHz, $D_2O$

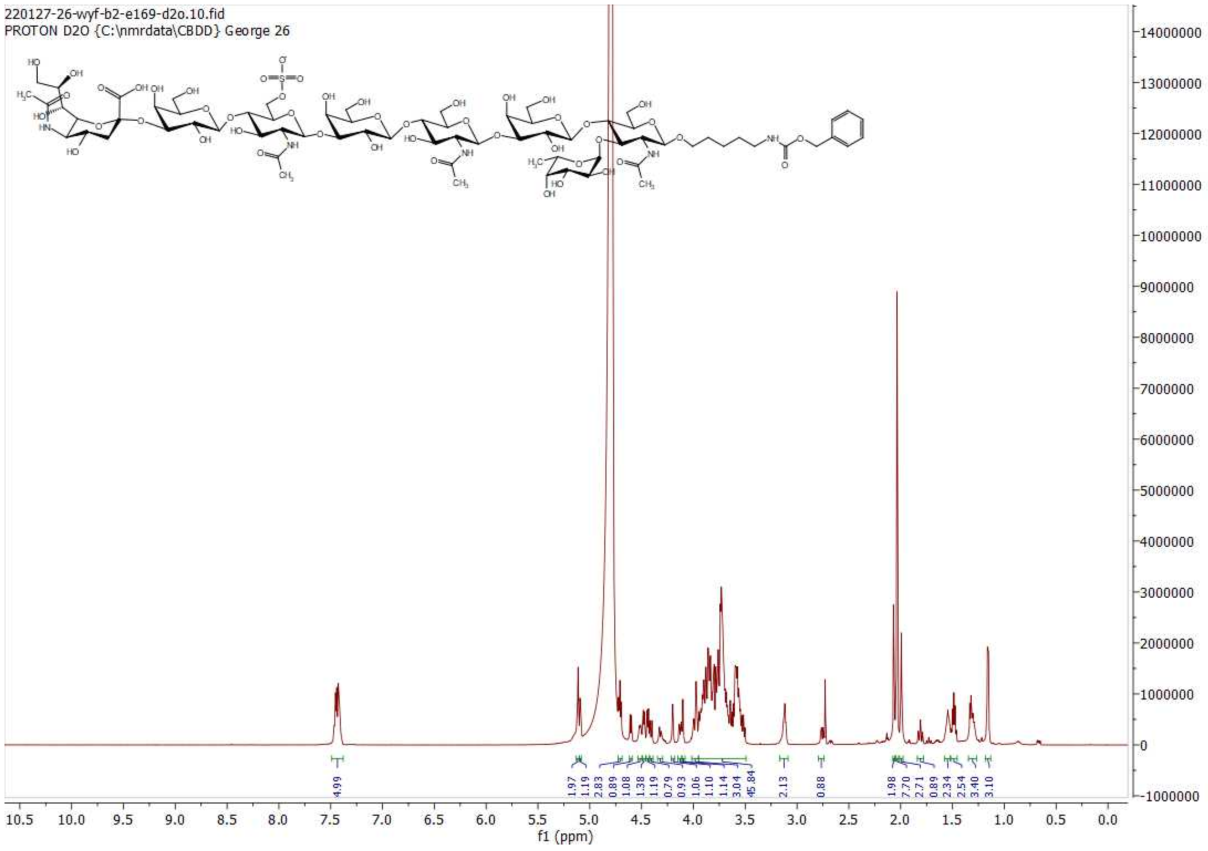

**$^{1}$H NMR of S1; 600MHz; $D_2O$**

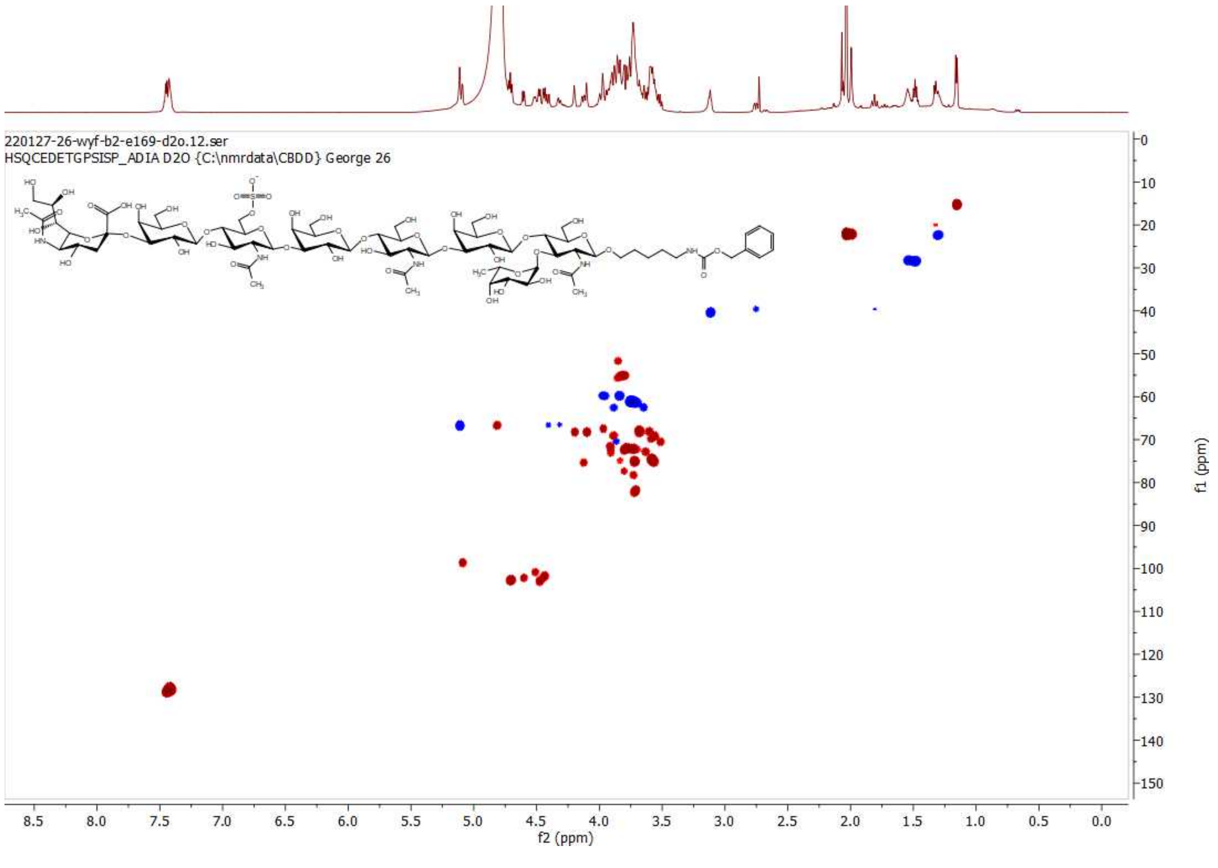

**HSQC of S1; 600 MHz/150 MHz, $D_2O$**

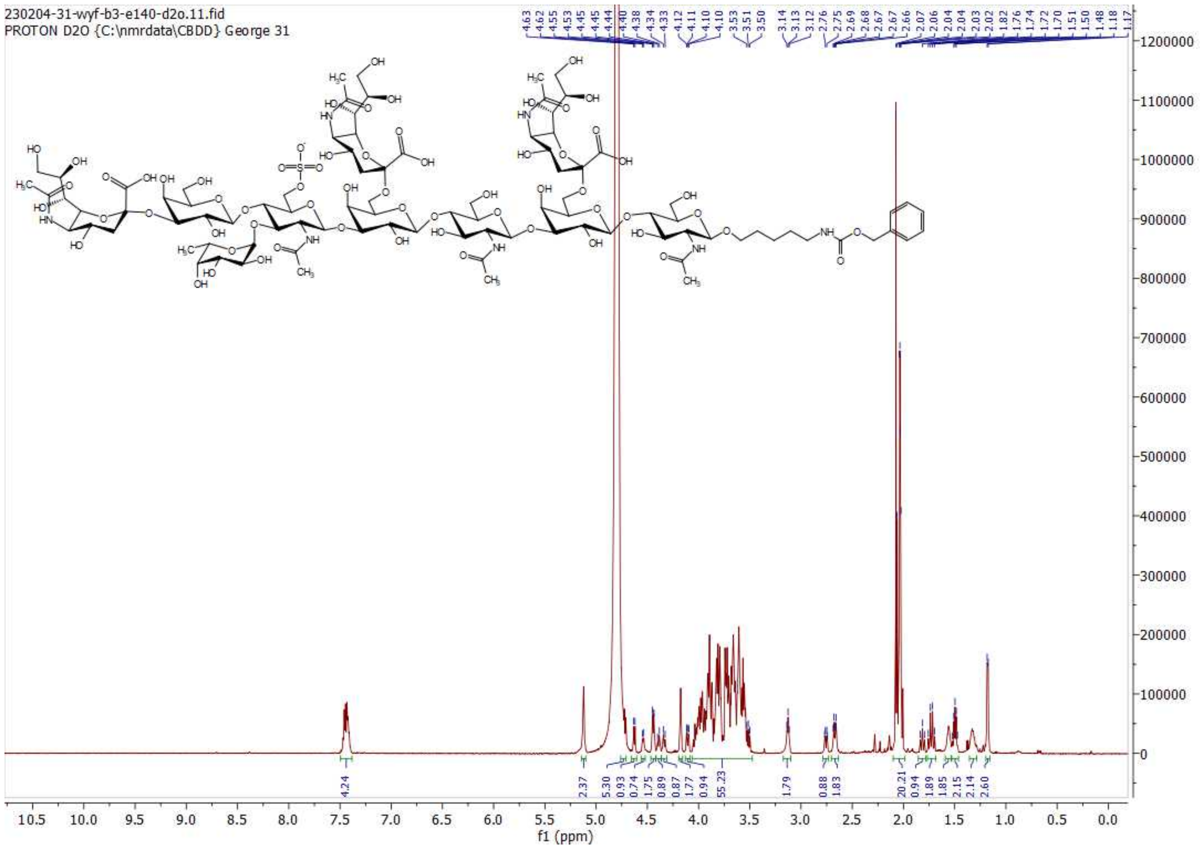

**[1]H NMR of S5; 600MHz; $D_2O$**

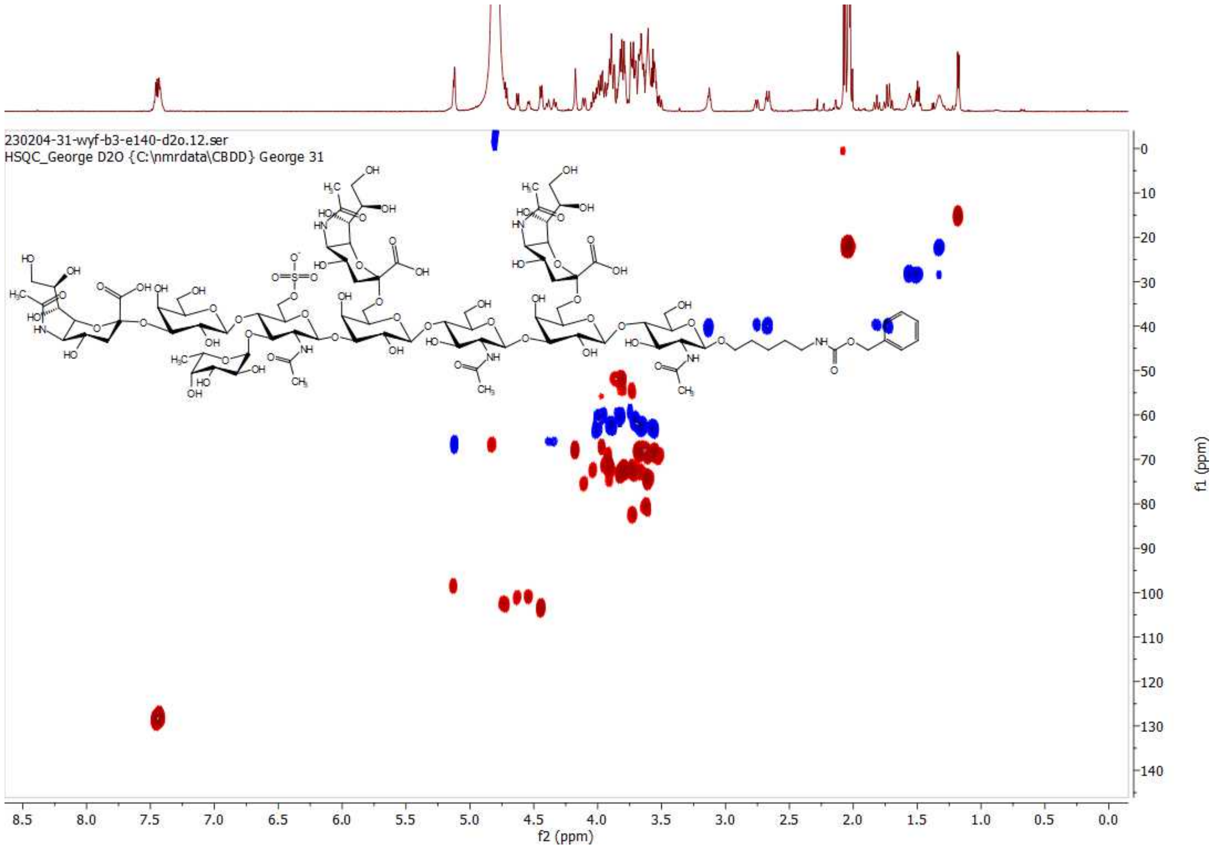

**HSQC of S5; 600 MHz/150 MHz, $D_2O$**

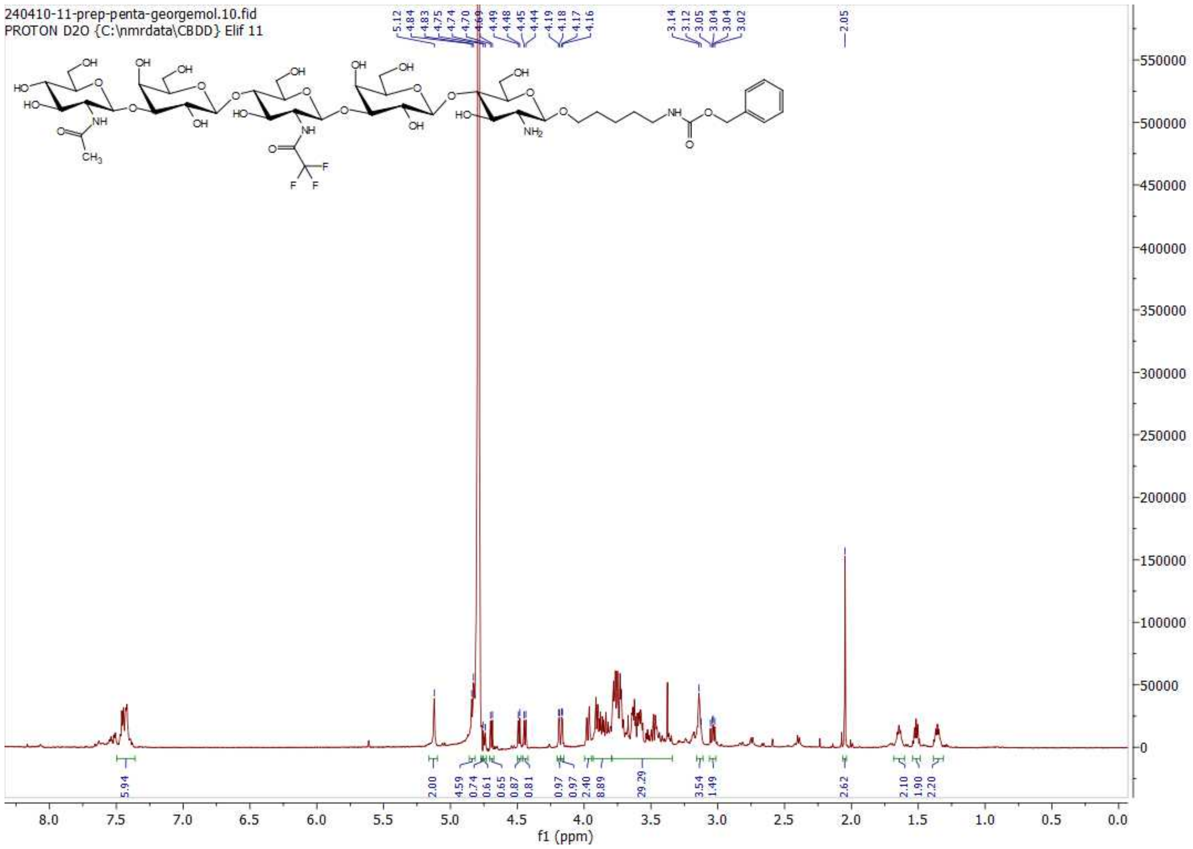

[1]H NMR of 6; 600MHz; $D_2O$

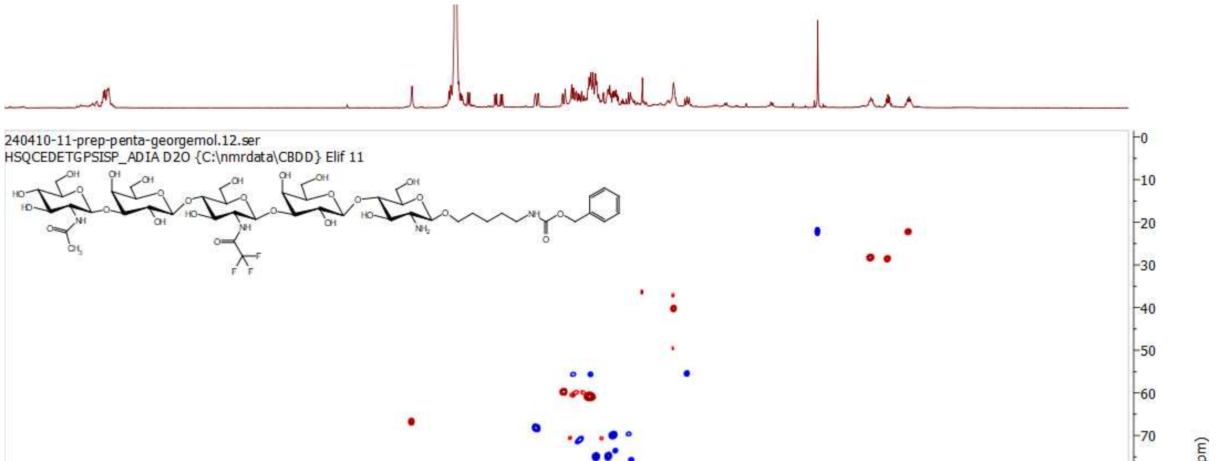

HSQC of 6; 600 MHz/150 MHz, $D_2O$

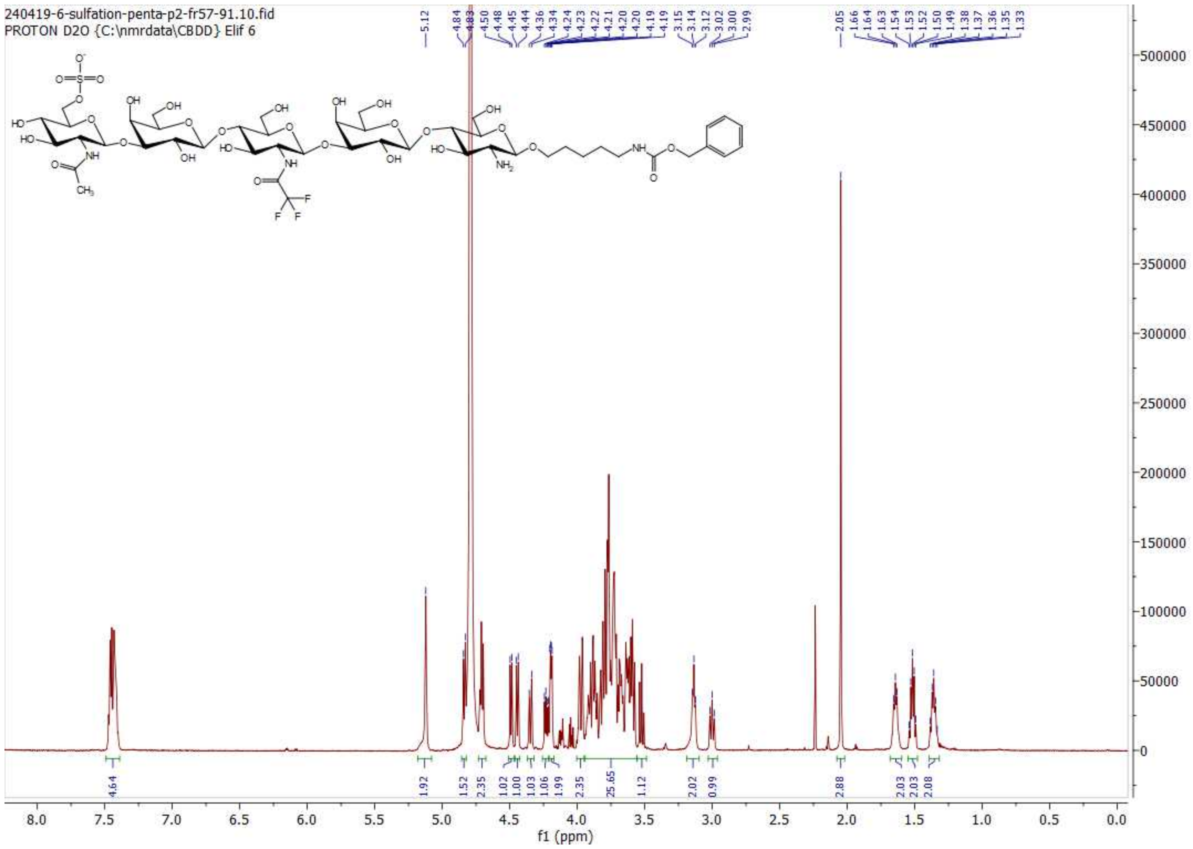

**[1]H NMR of 7; 600MHz; $D_2O$**

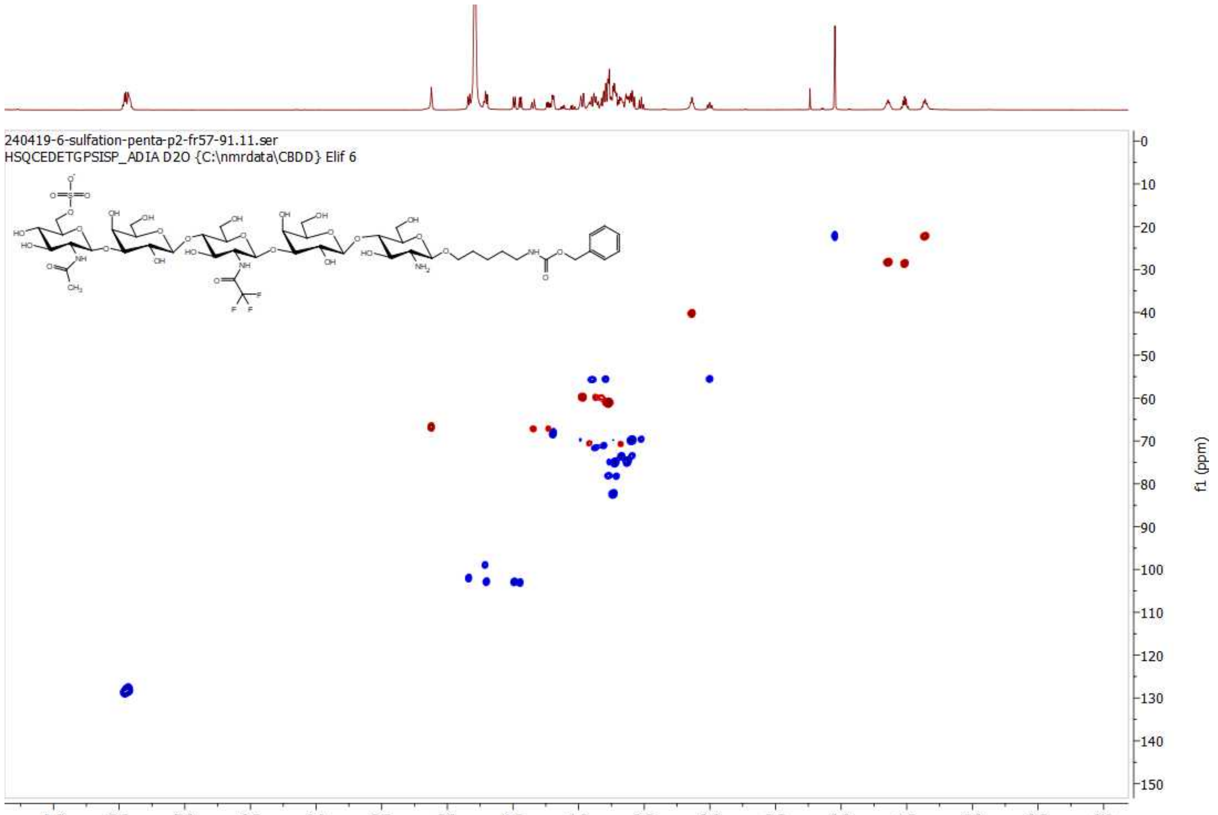

**HSQC of 7; 600 MHz/150 MHz, $D_2O$**

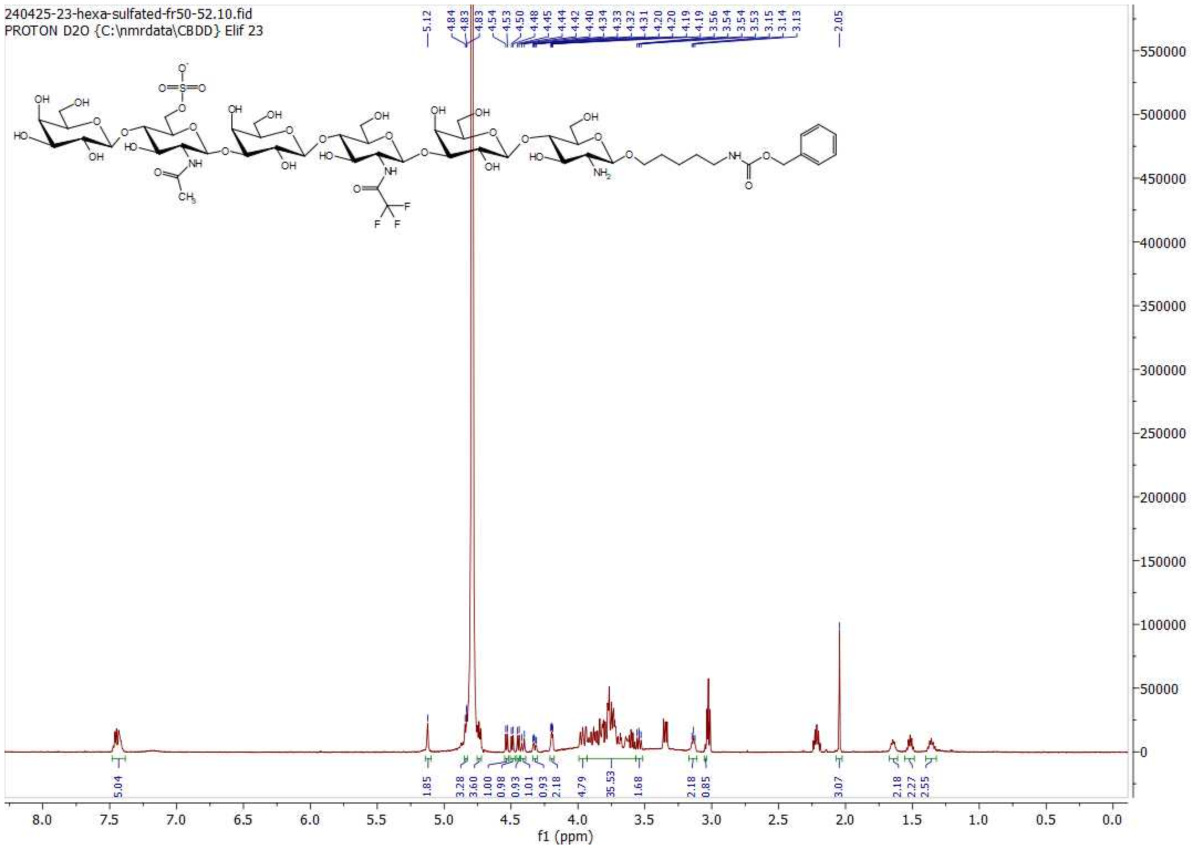

**[1]H NMR of 8; 600MHz; $D_2O$**

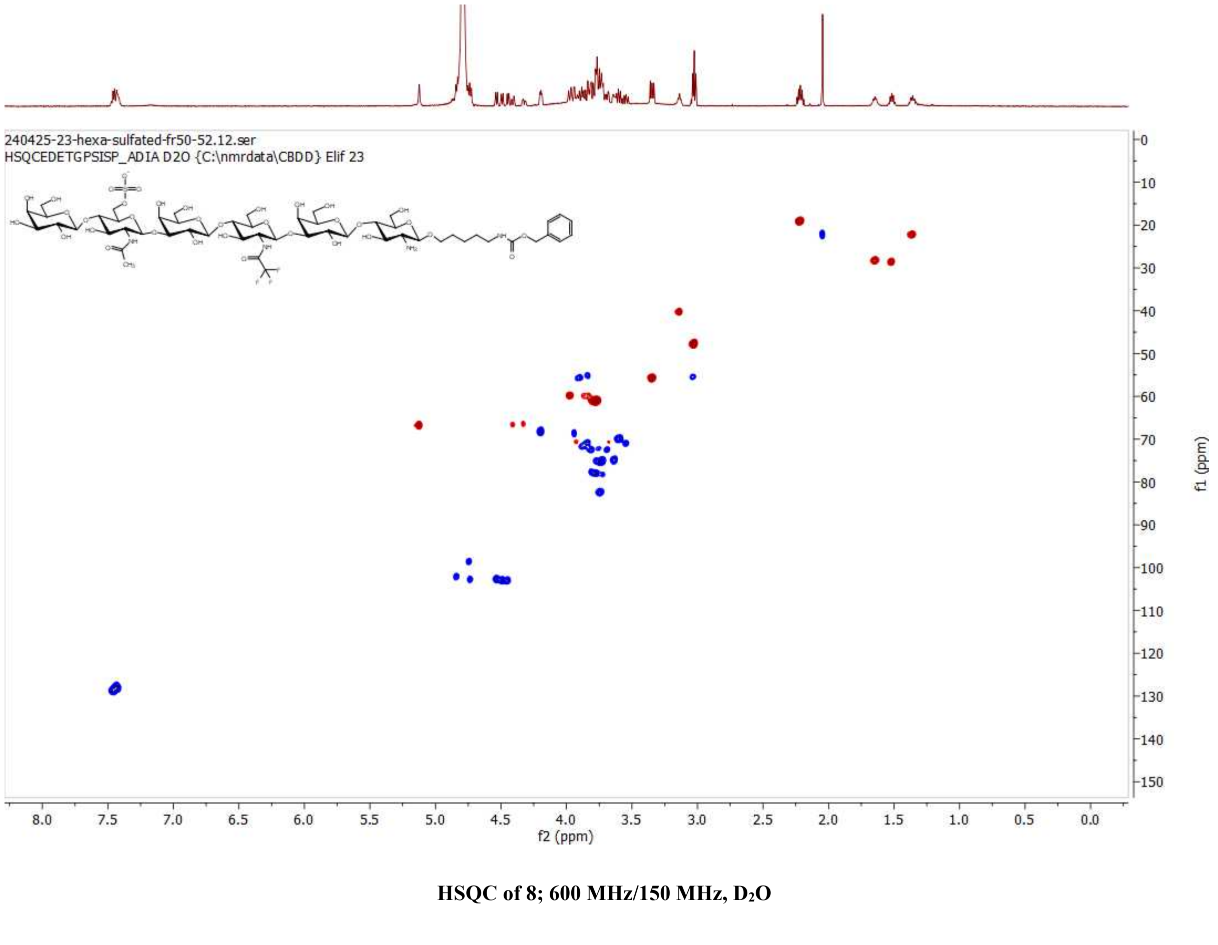

**HSQC of 8; 600 MHz/150 MHz, $D_2O$**

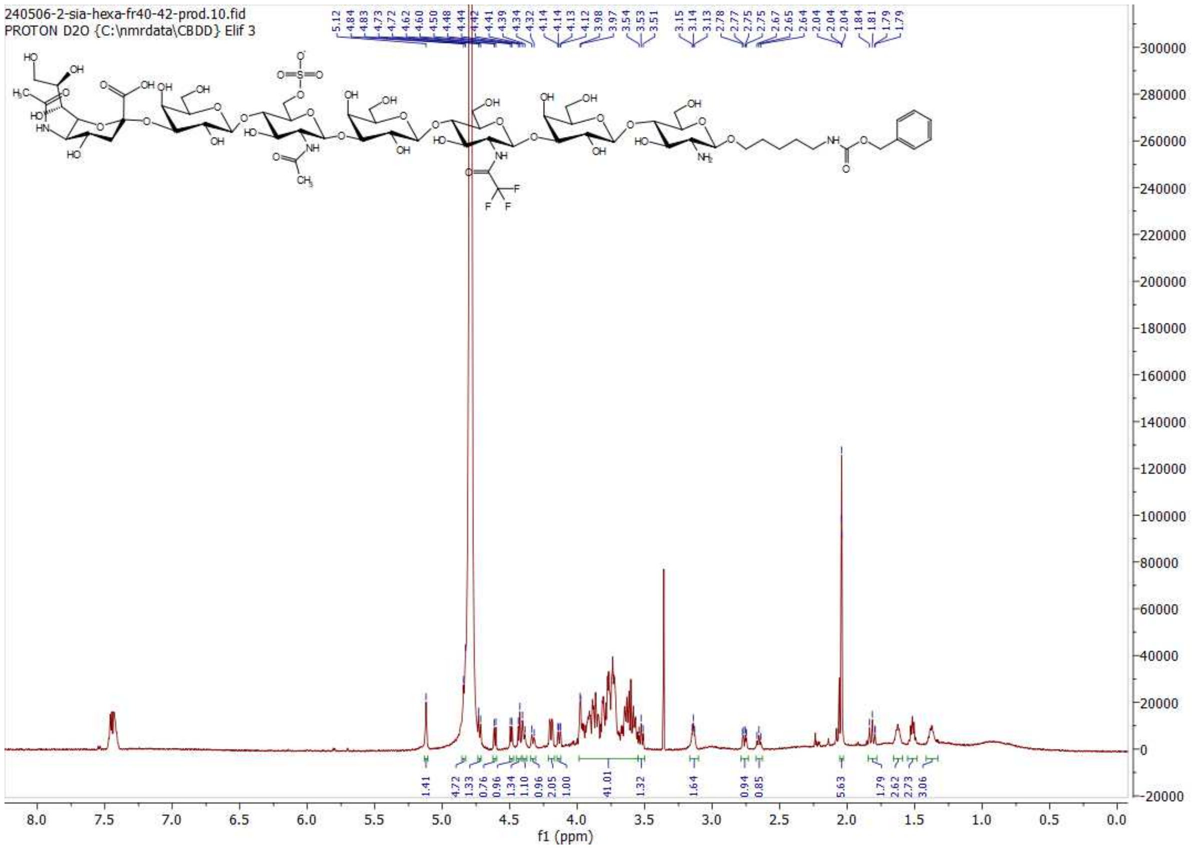

$^{1}$H NMR of 9; 600MHz; $D_2O$

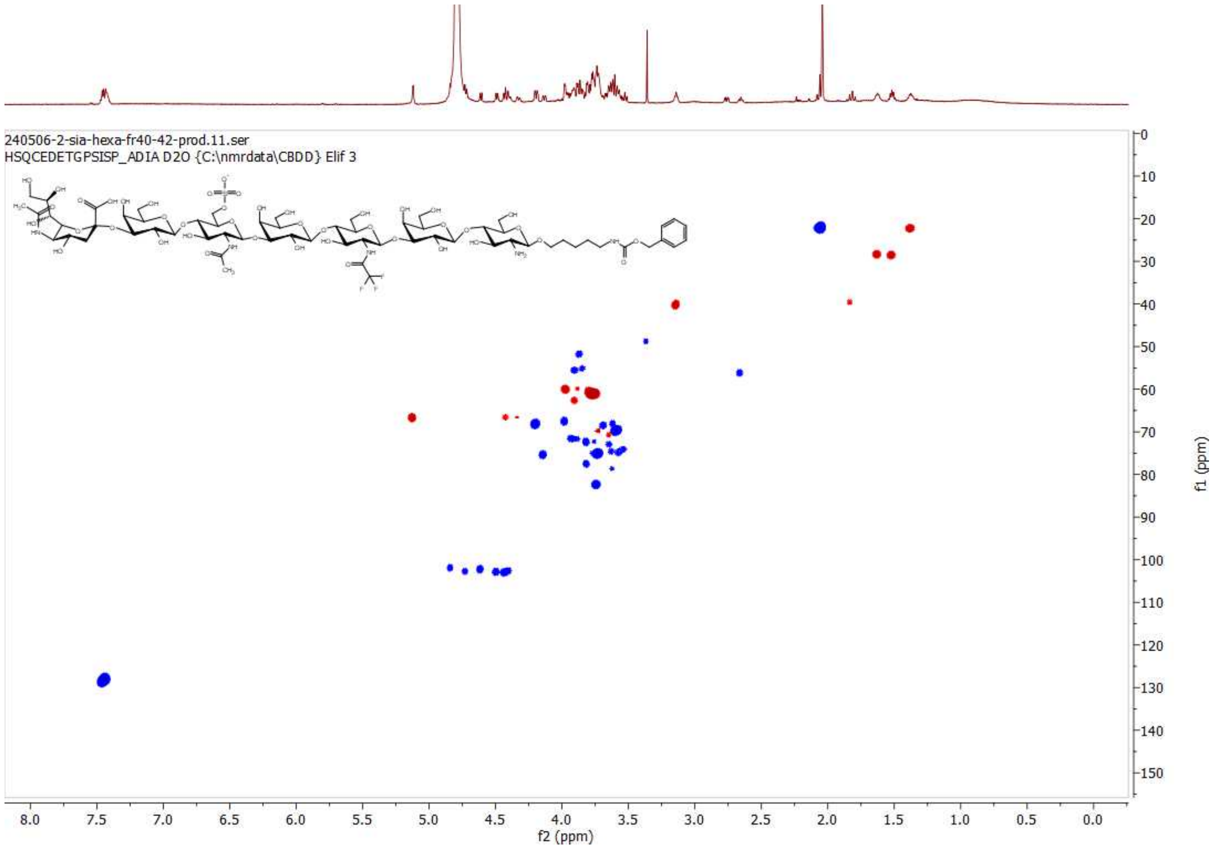

HSQC of 9; 600 MHz/150 MHz, $D_2O$

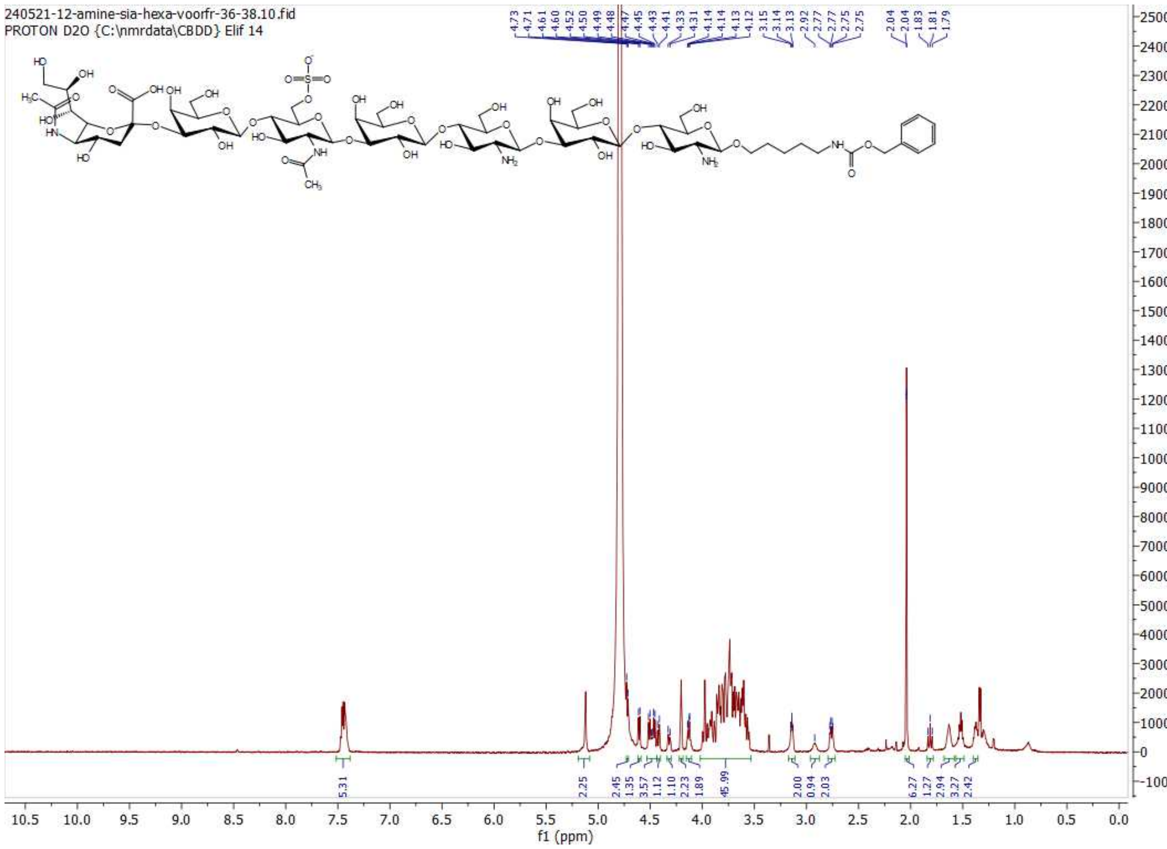

**$^{1}$H NMR of 10; 600MHz; $D_2O$**

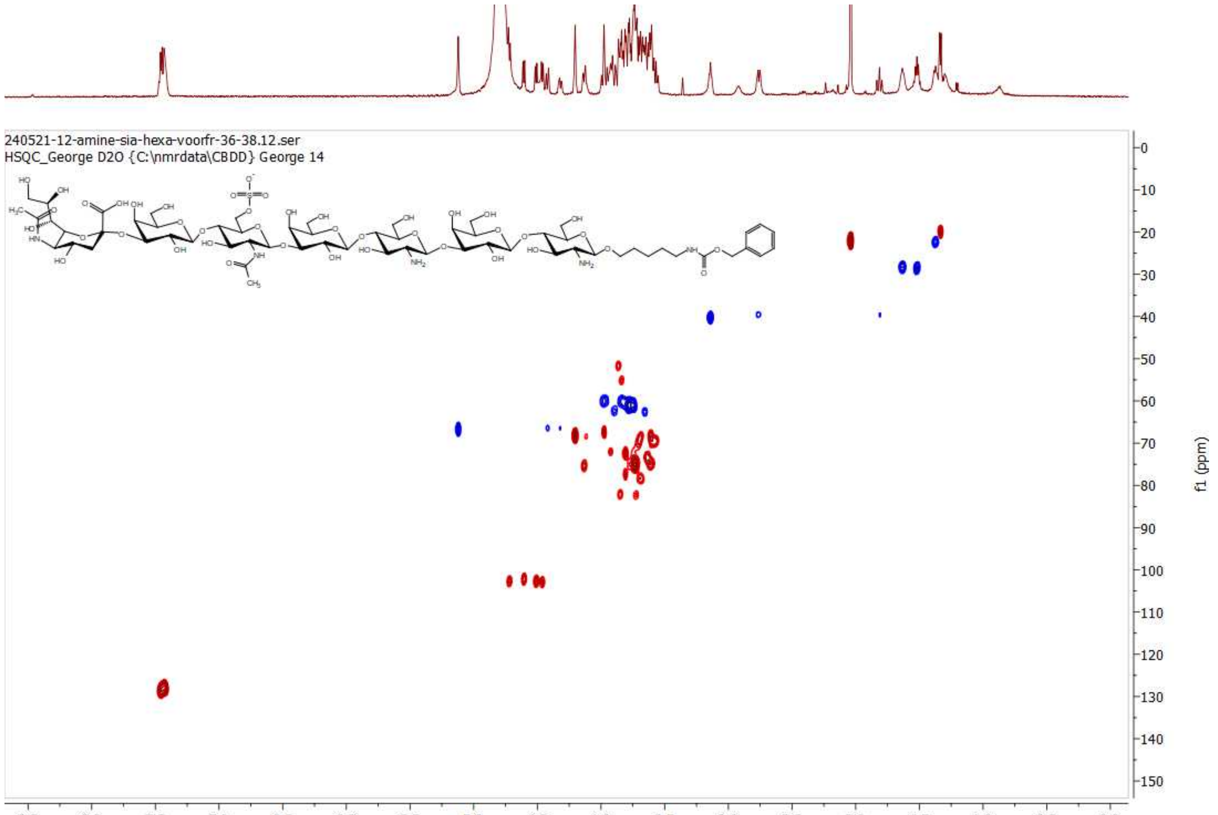

**HSQC of 10; 600 MHz/150 MHz, $D_2O$**

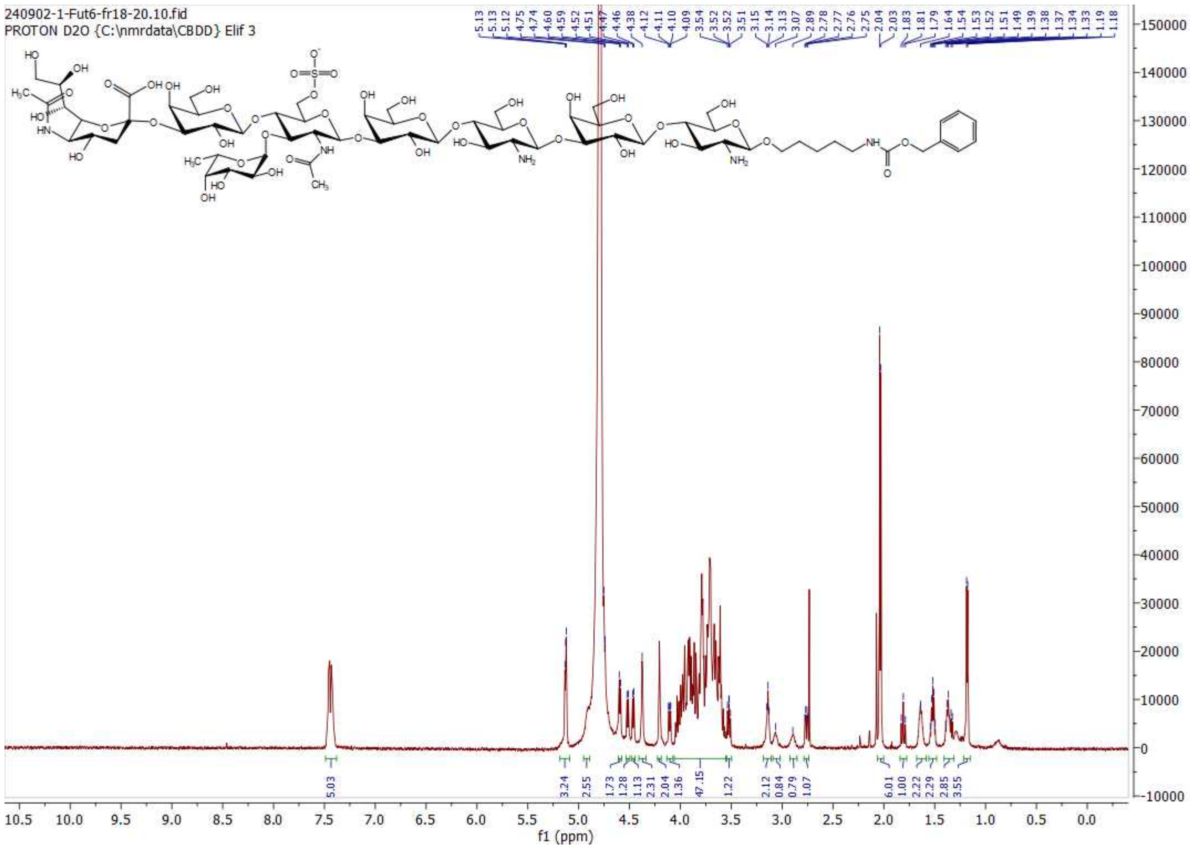

**$^{1}$H NMR of 11; 600MHz; $D_2O$**

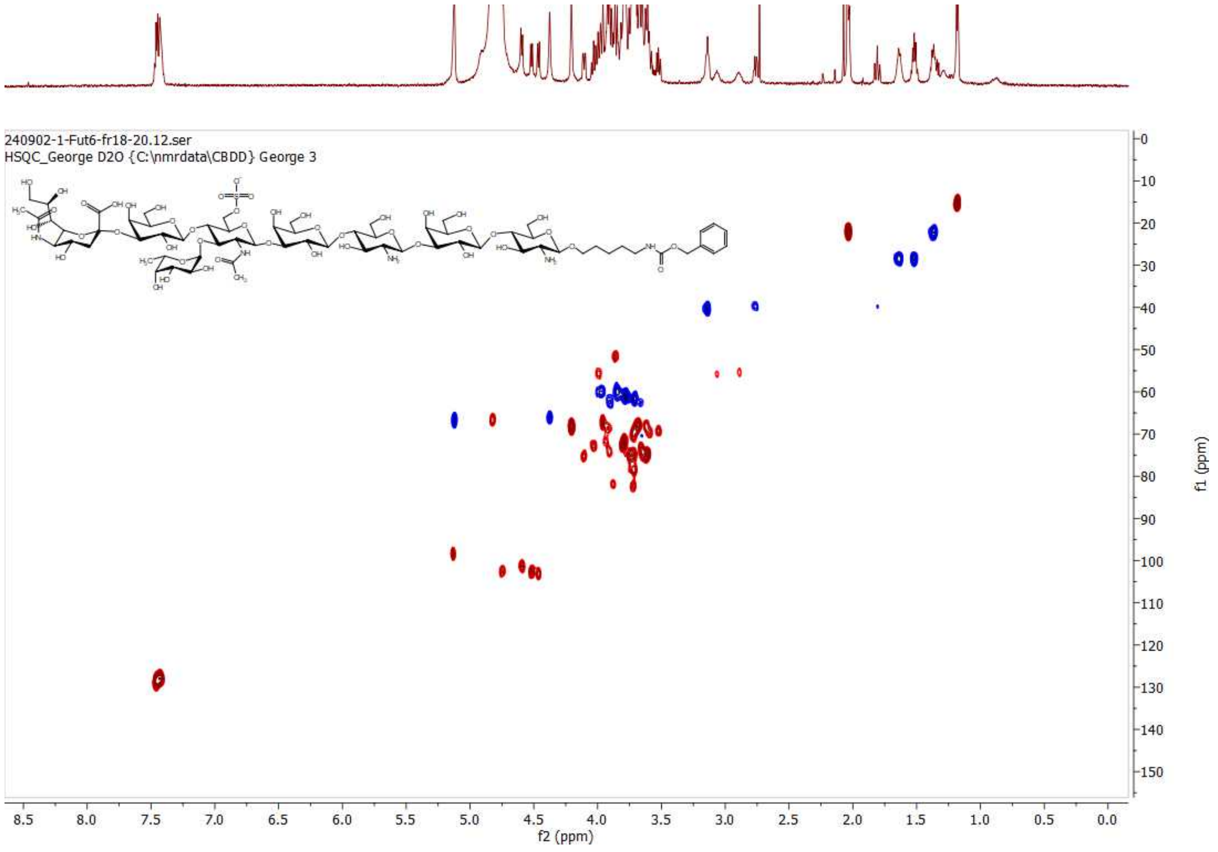

**HSQC of 11; 600 MHz/150 MHz, $D_2O$**

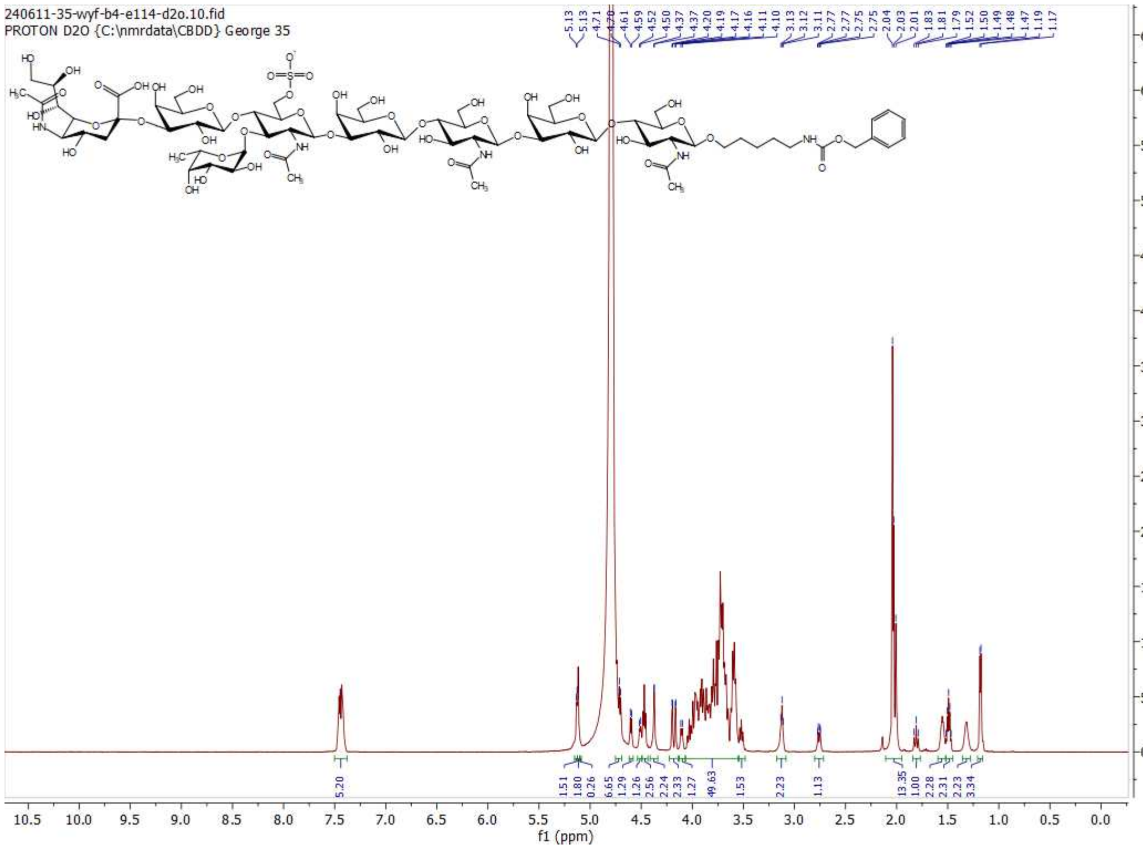

**[1]H NMR of 4; 600MHz; $D_2O$**

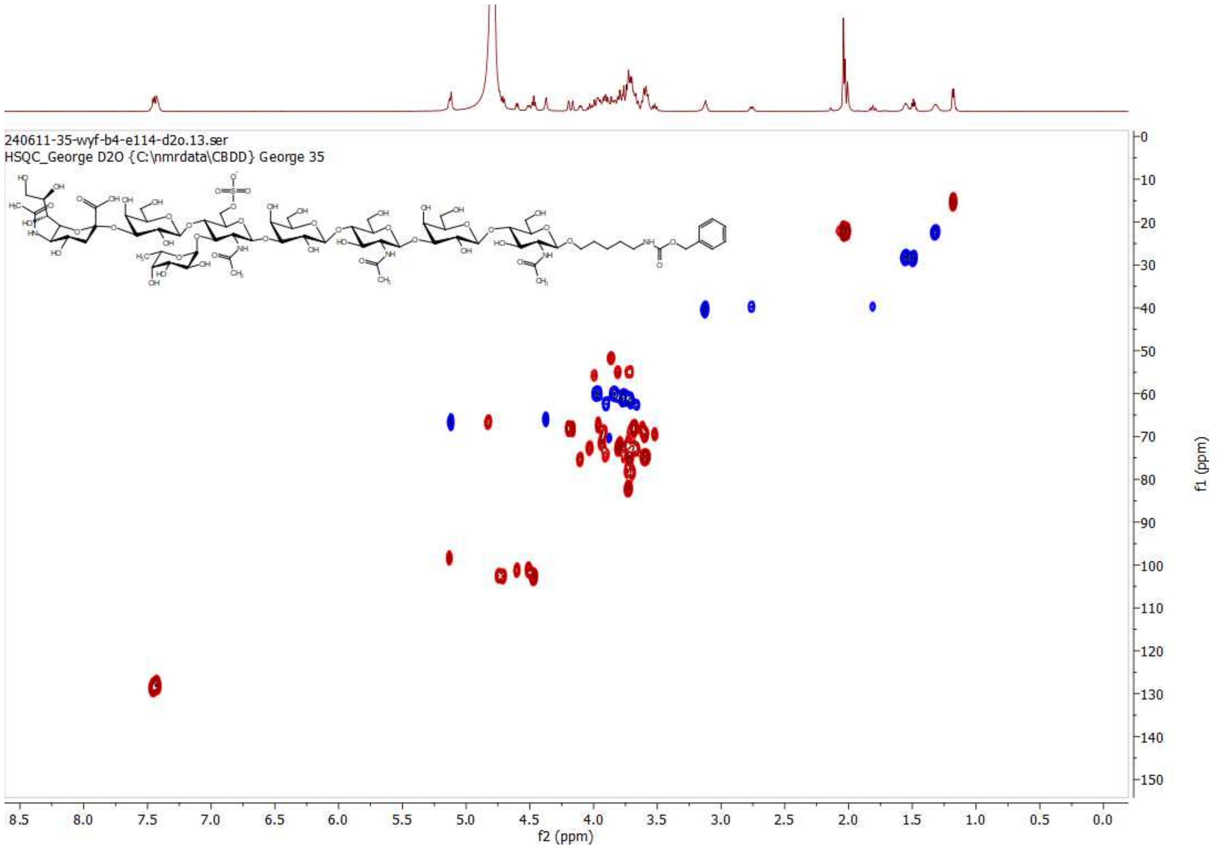

**HSQC of 4; 600 MHz/150 MHz, $D_2O$**

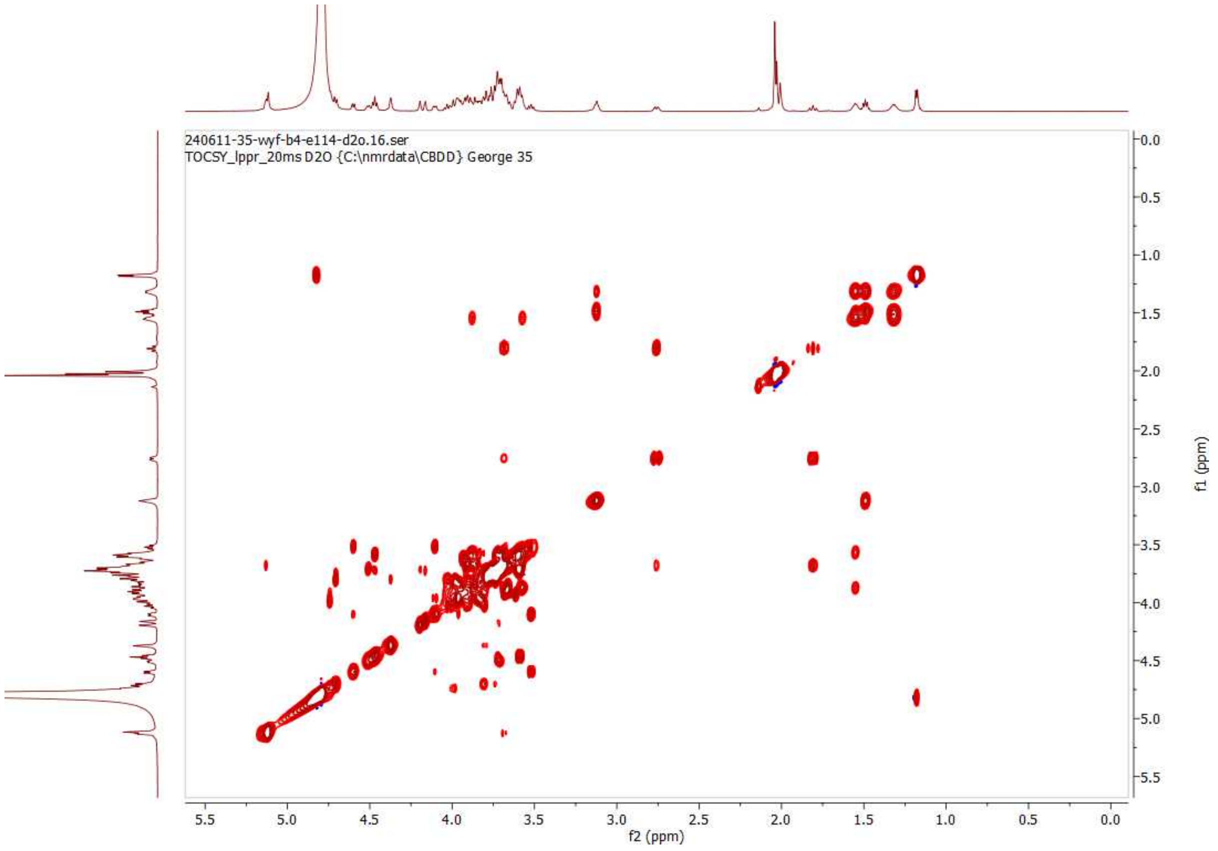

**TOCSY (20 ms) of 4; 600MHz; $D_2O$**

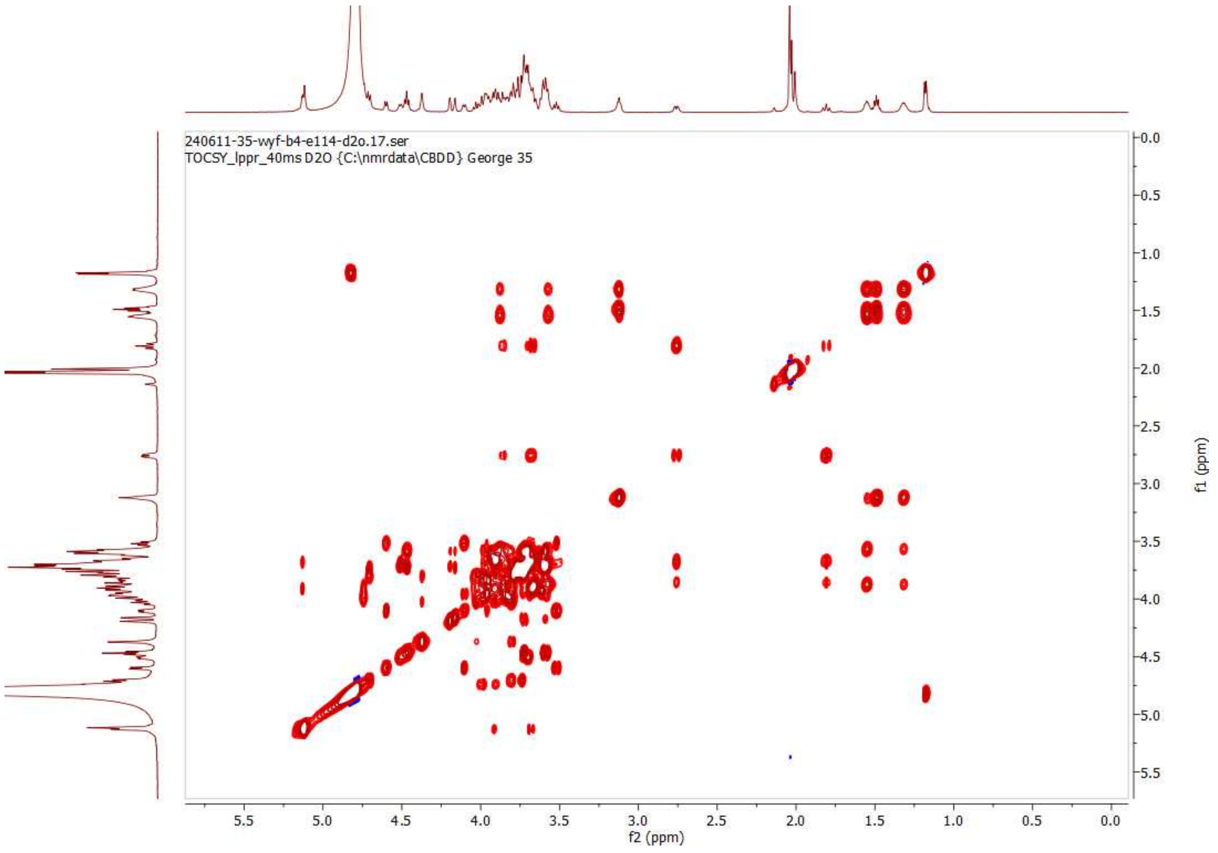

**TOCSY (40 ms) of 4; 600MHz; $D_2O$**

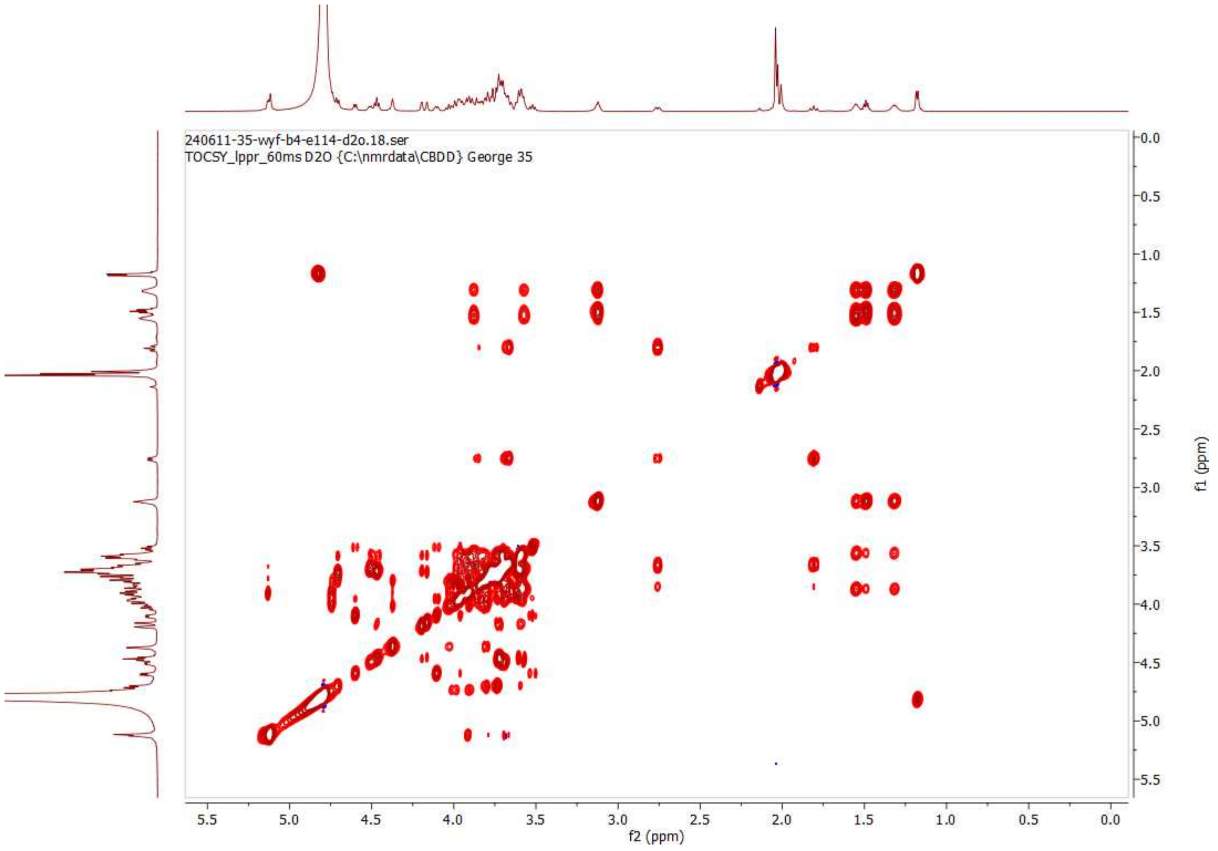

**TOCSY (60 ms) of 4; 600MHz; $D_2O$**

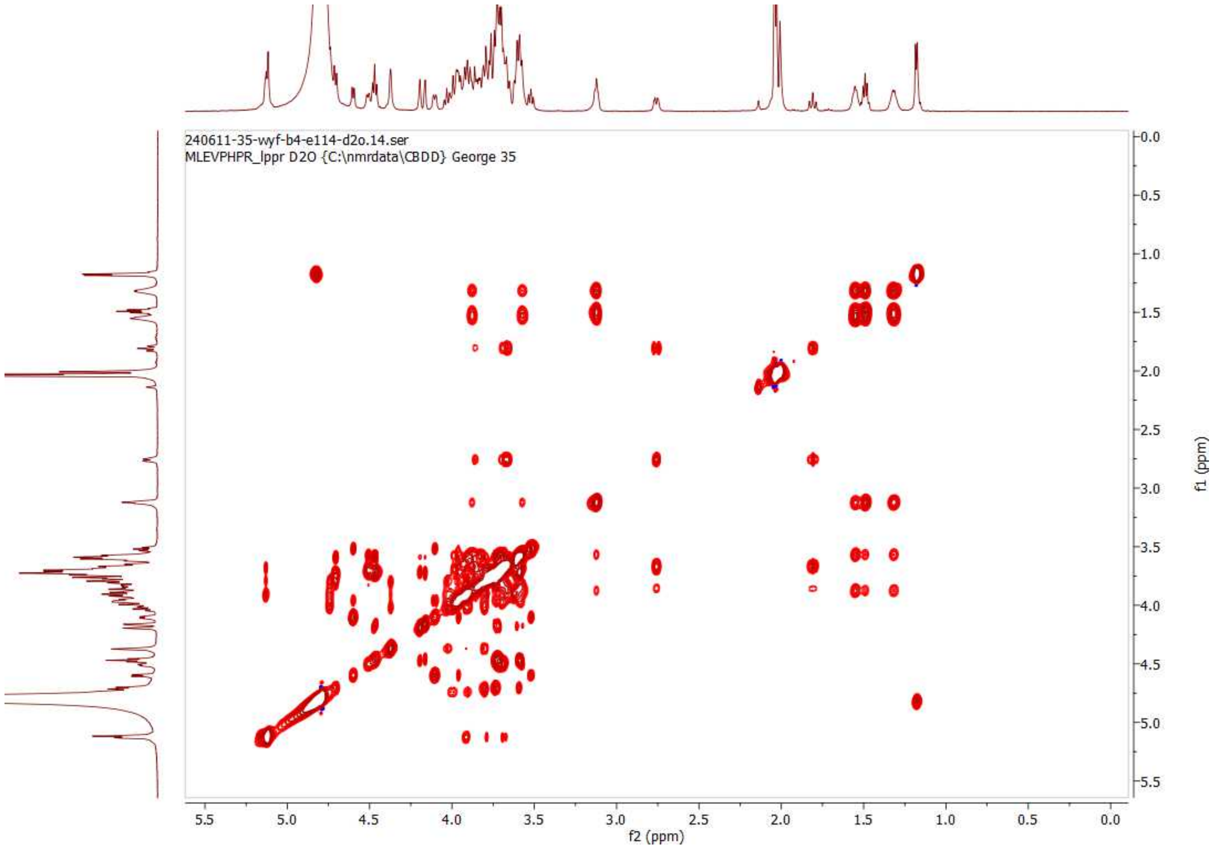

**TOCSY (80 ms) of 4; 600MHz; $D_2O$**

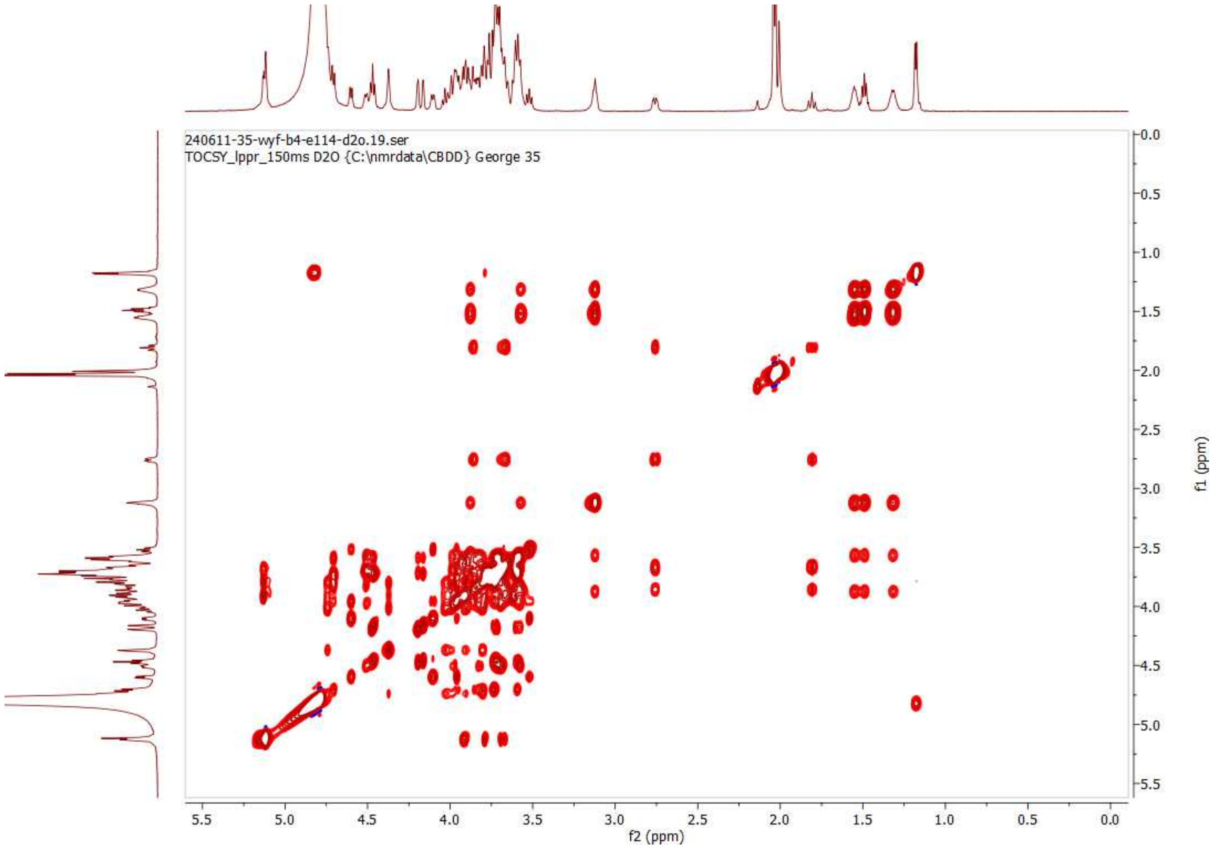

**TOCSY (150 ms) of 4; 600MHz; $D_2O$**

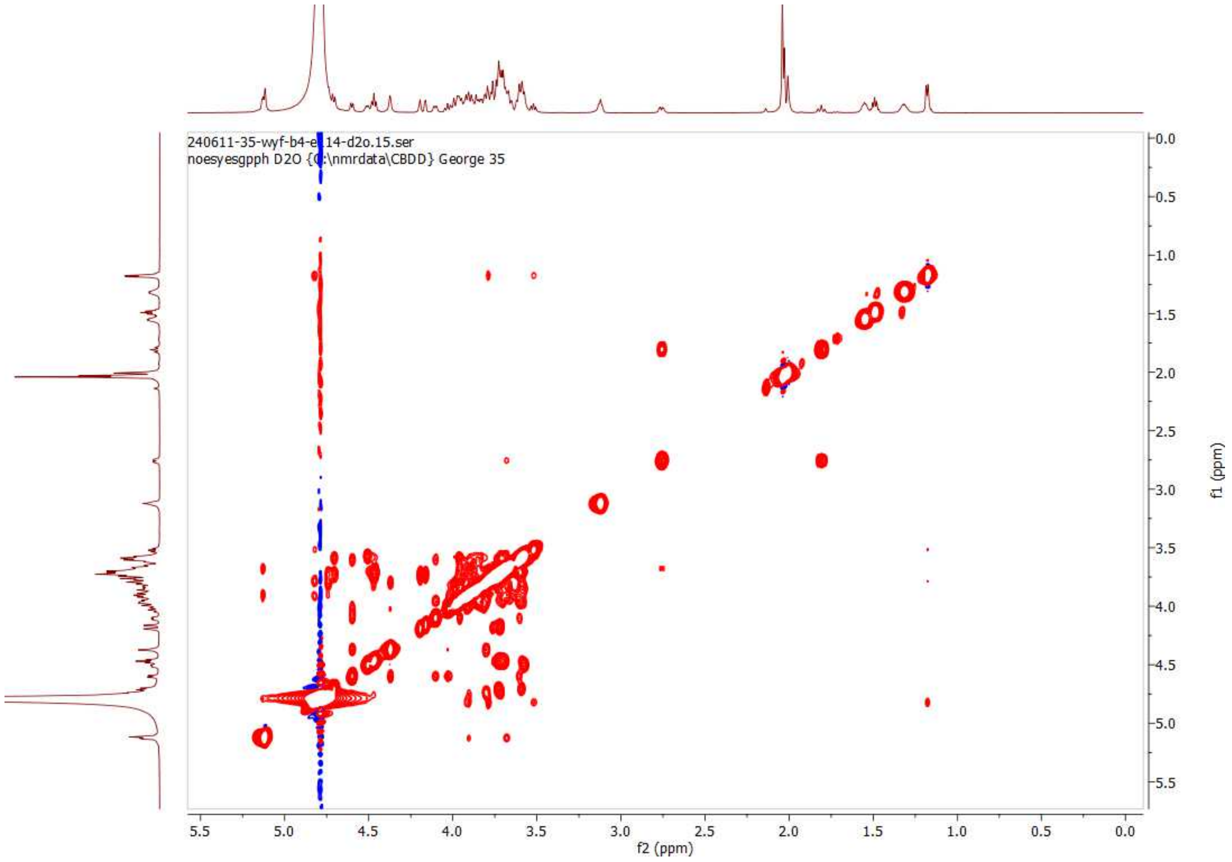

**NOESY (300 ms) of 4; 600 MHz, $D_2O$**

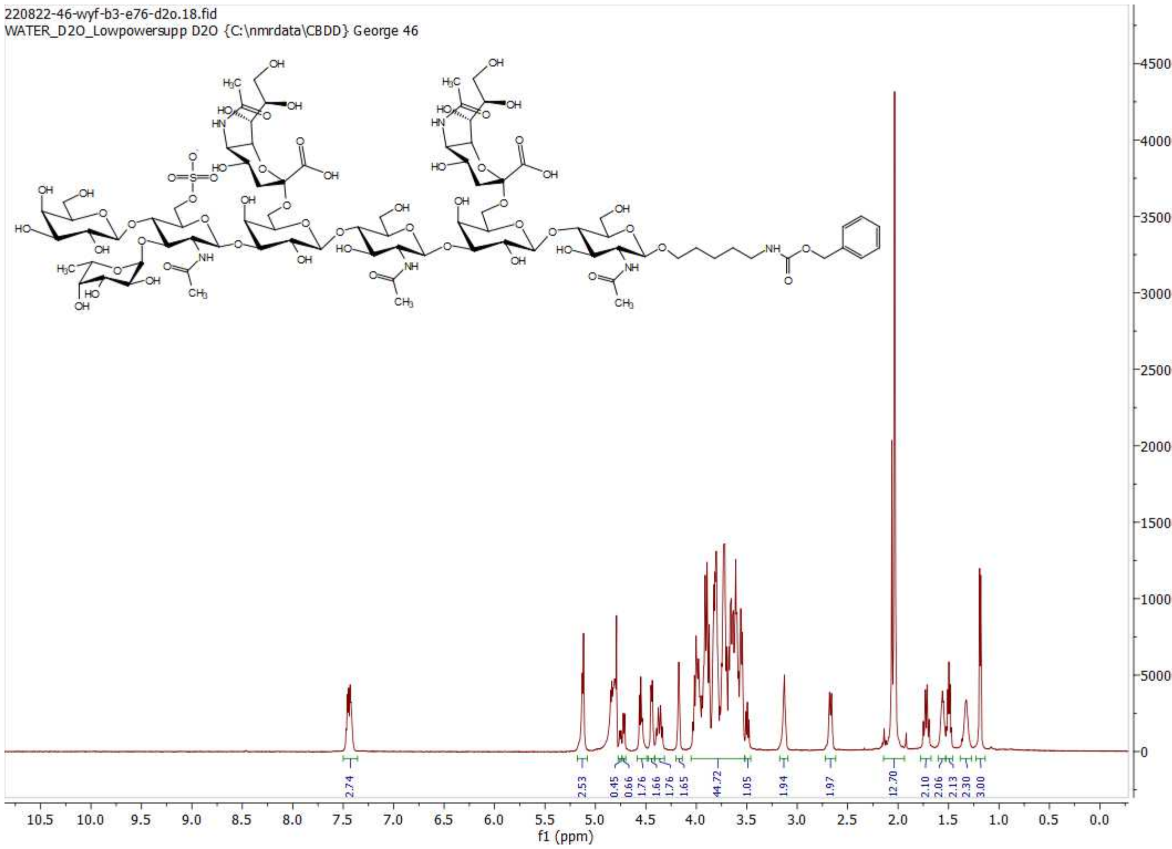

**[1]H NMR of 13; 600MHz; $D_2O$**

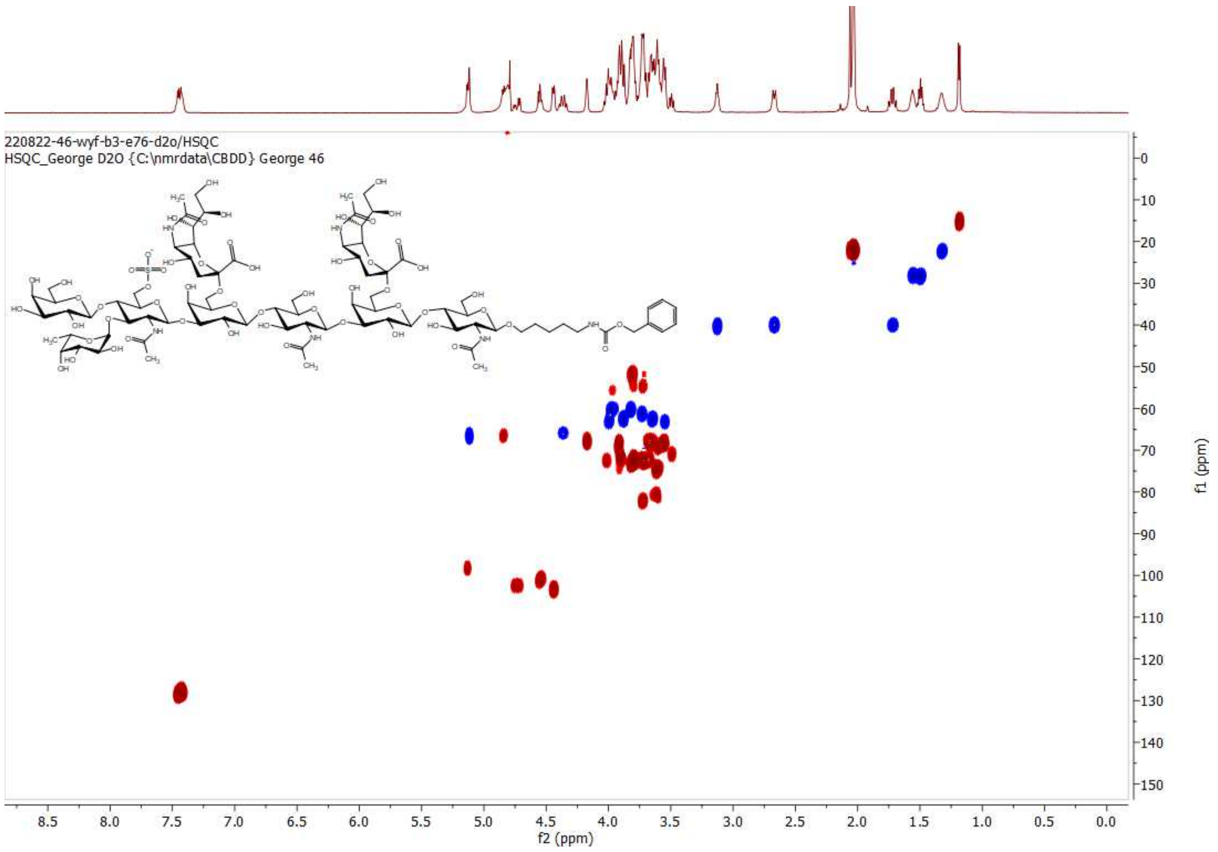

**HSQC of 13; 600 MHz/150 MHz, $D_2O$**

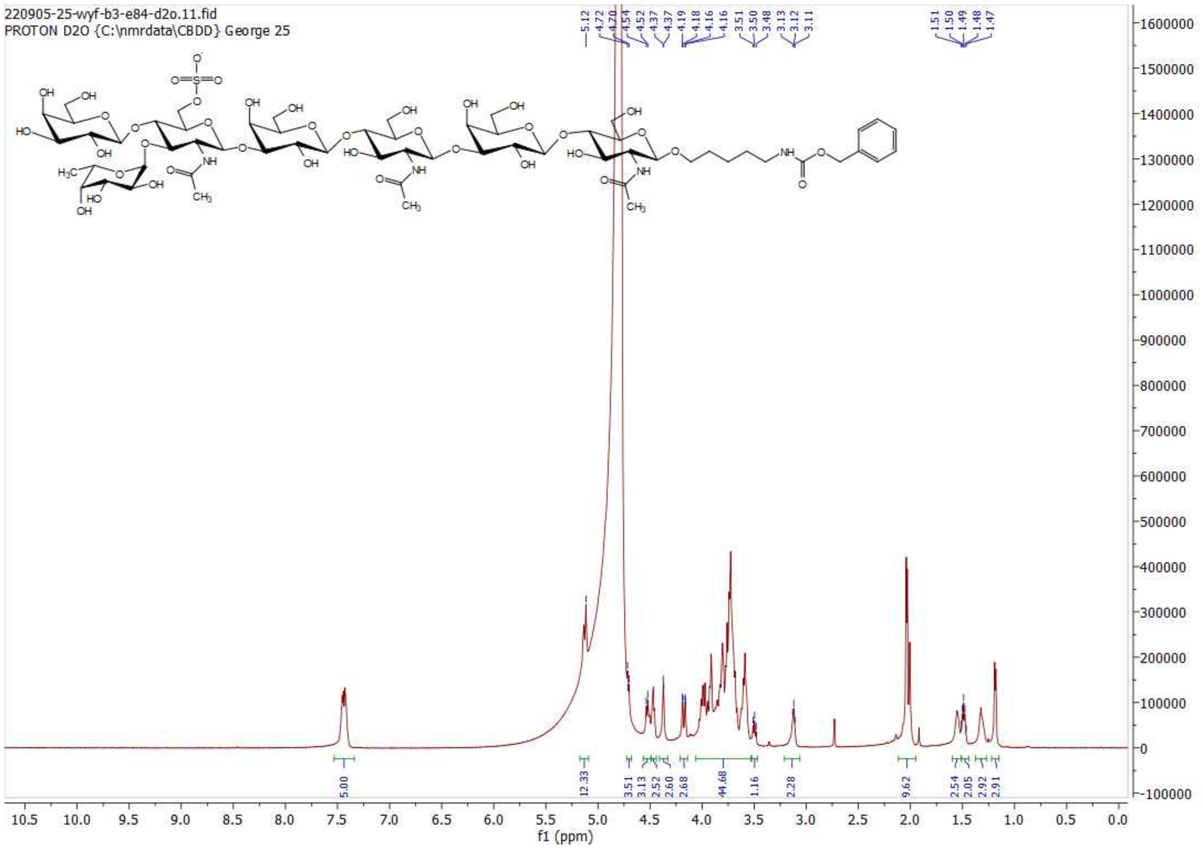

**$^1$H NMR of 14; 600MHz; $D_2O$**

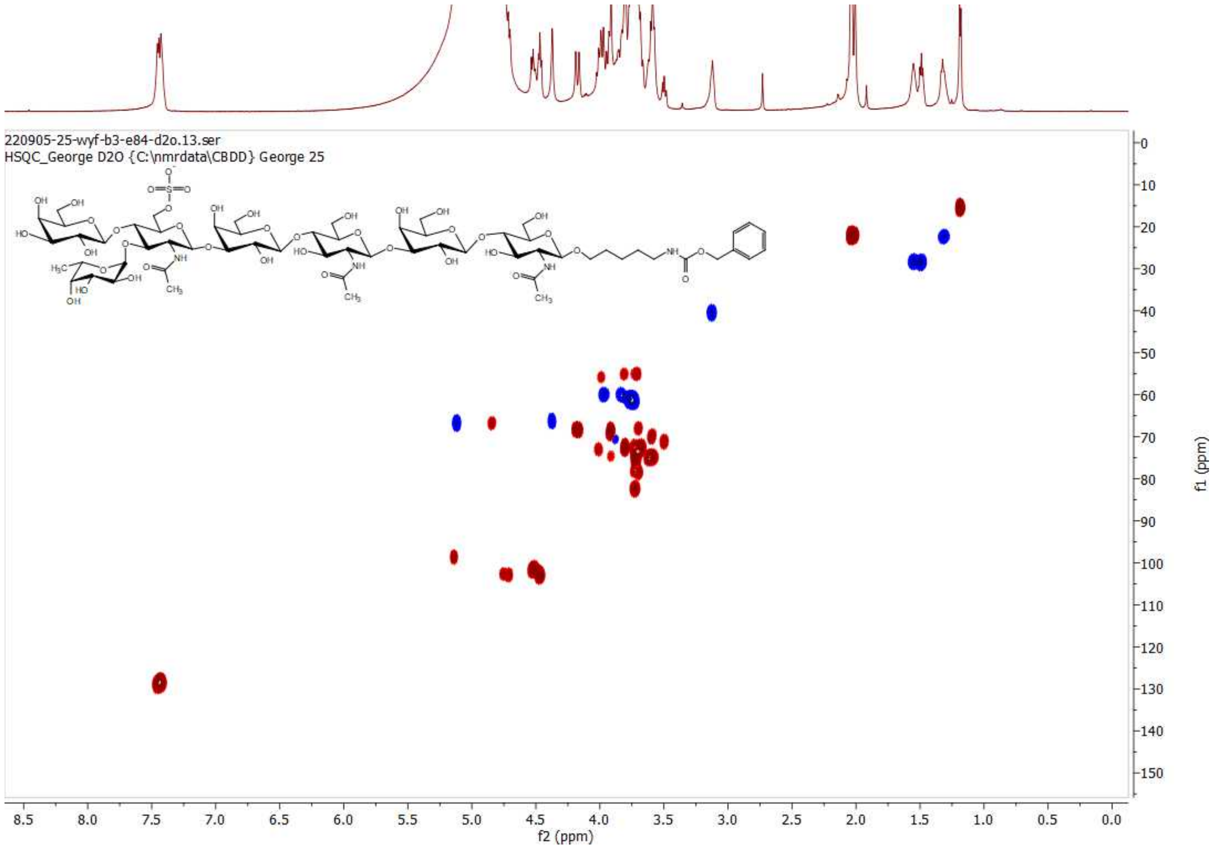

**HSQC of 14; 600 MHz/150 MHz, $D_2O$**

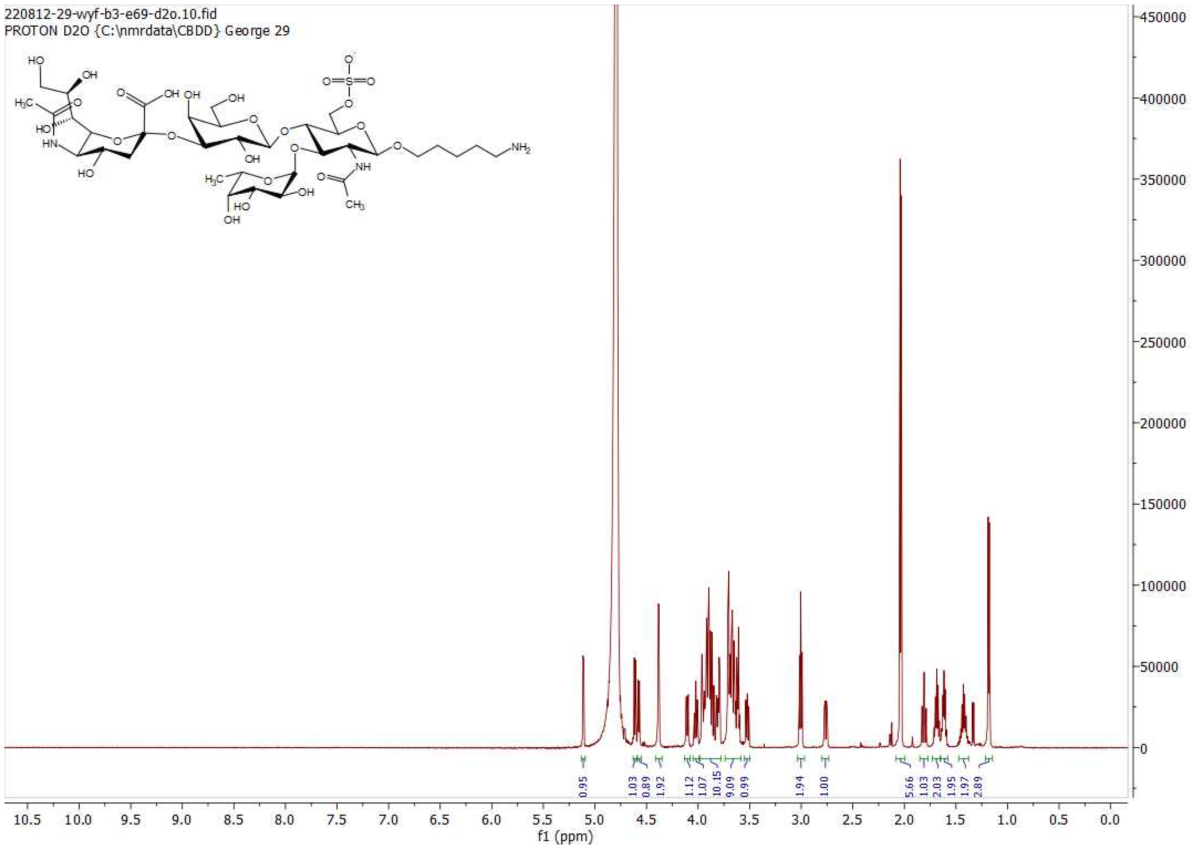

**[1]H NMR of G; 600MHz; $D_2O$**

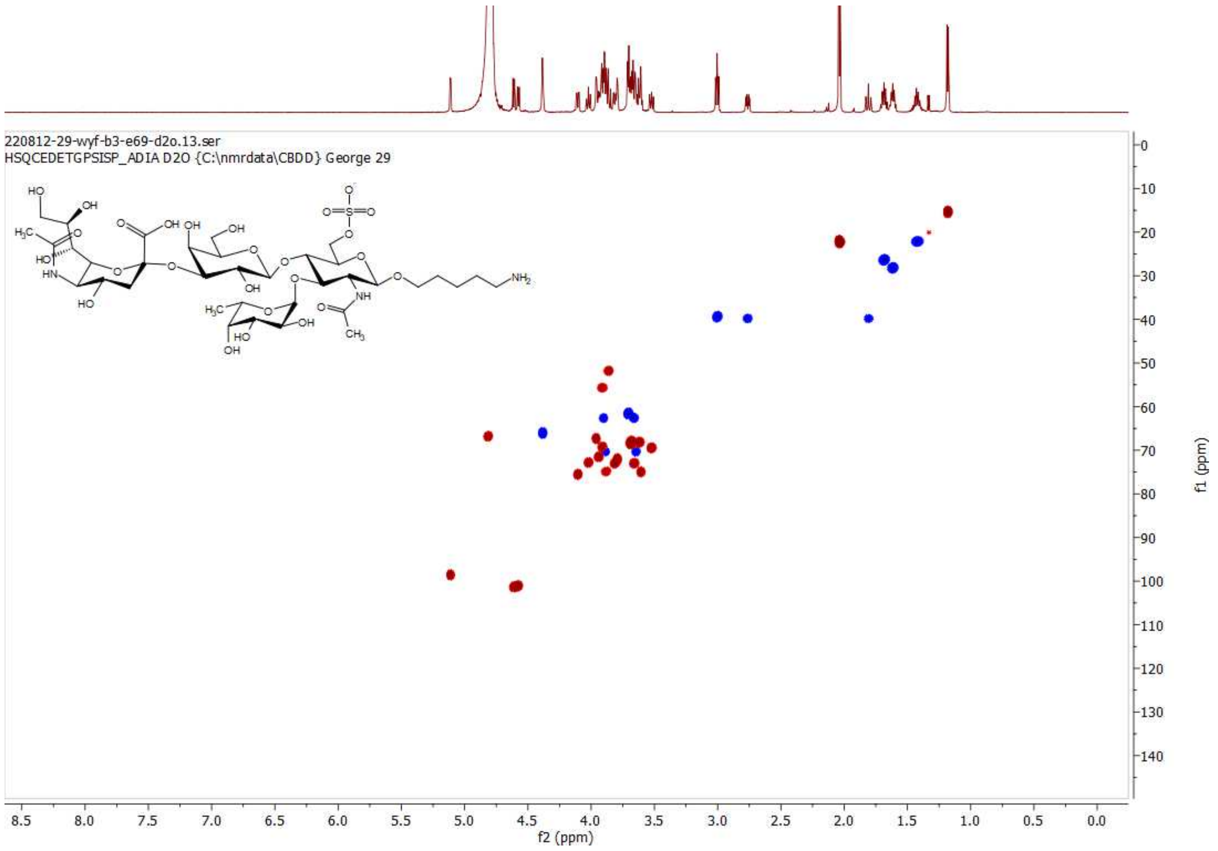

**HSQC of G; 600 MHz/150 MHz, $D_2O$**

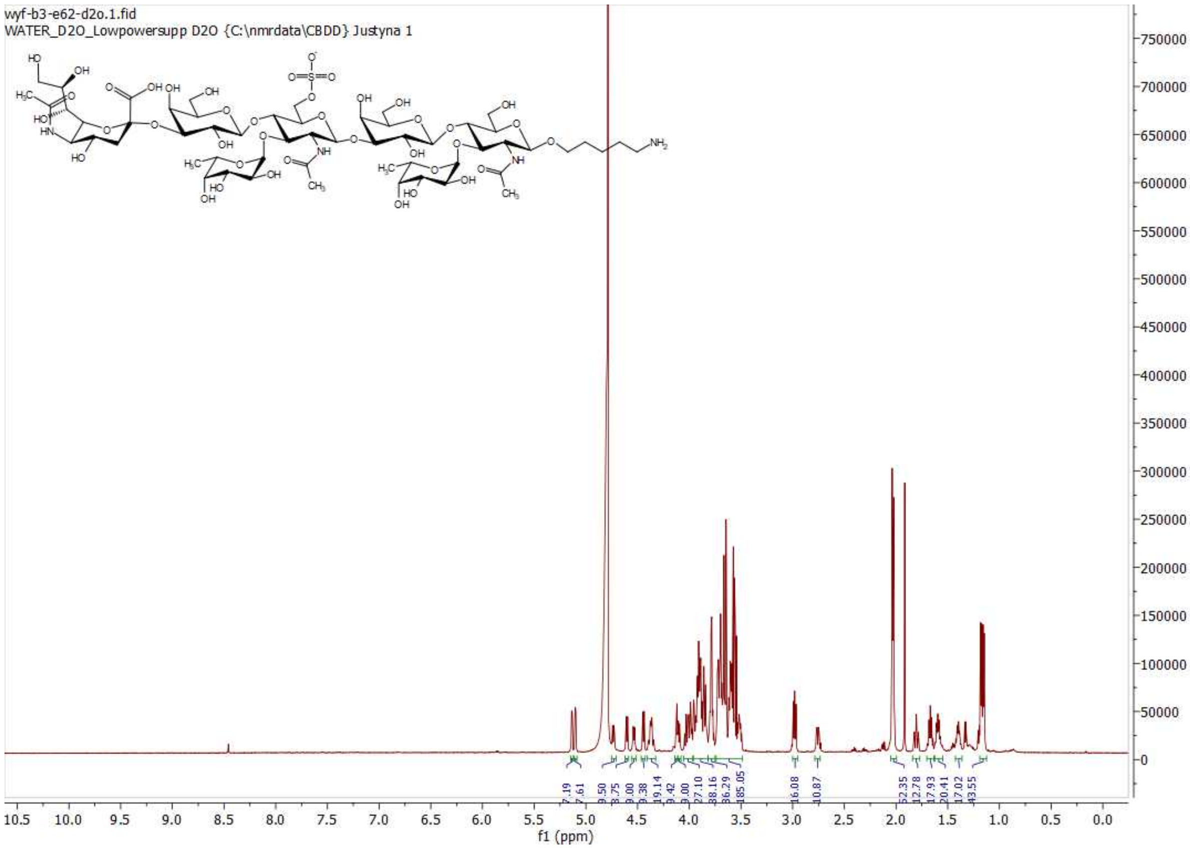

[1]H NMR of N; 600MHz; $D_2O$

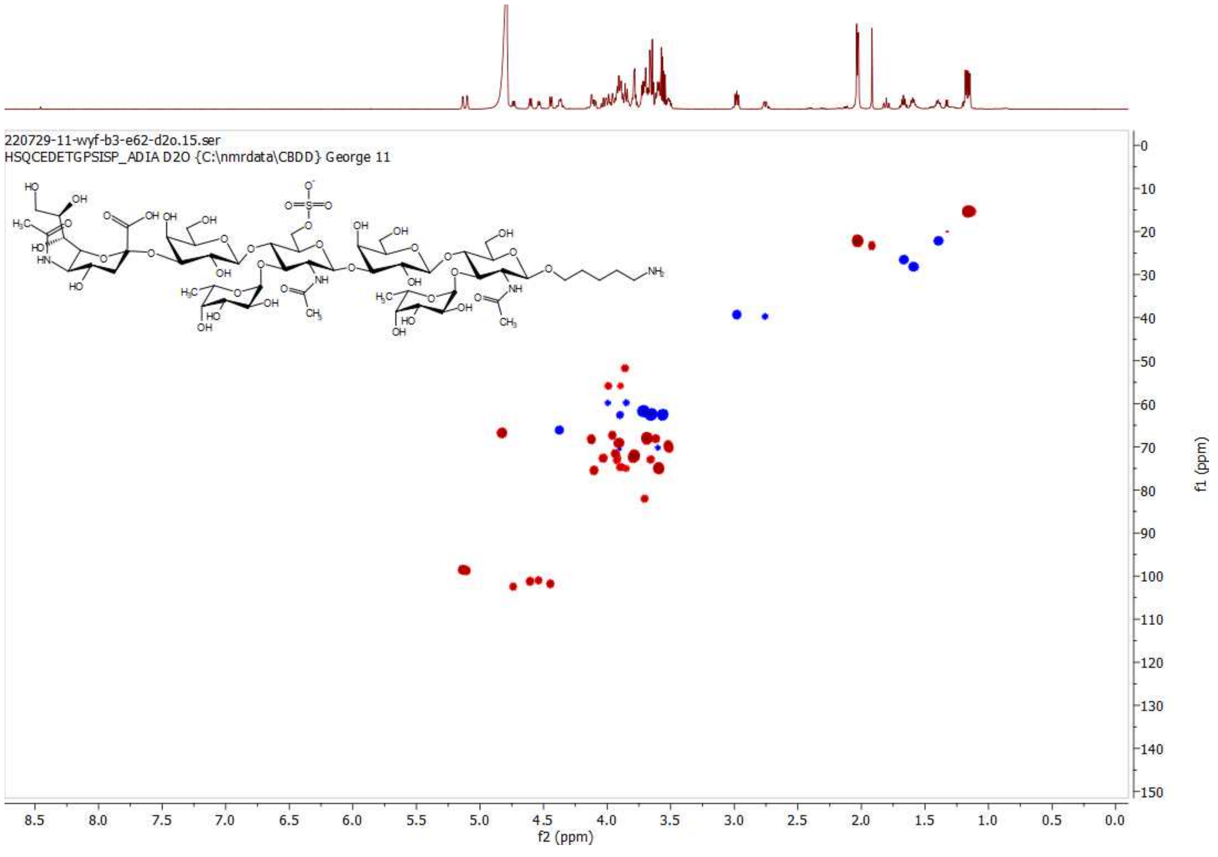

HSQC of N; 600 MHz/150 MHz, $D_2O$

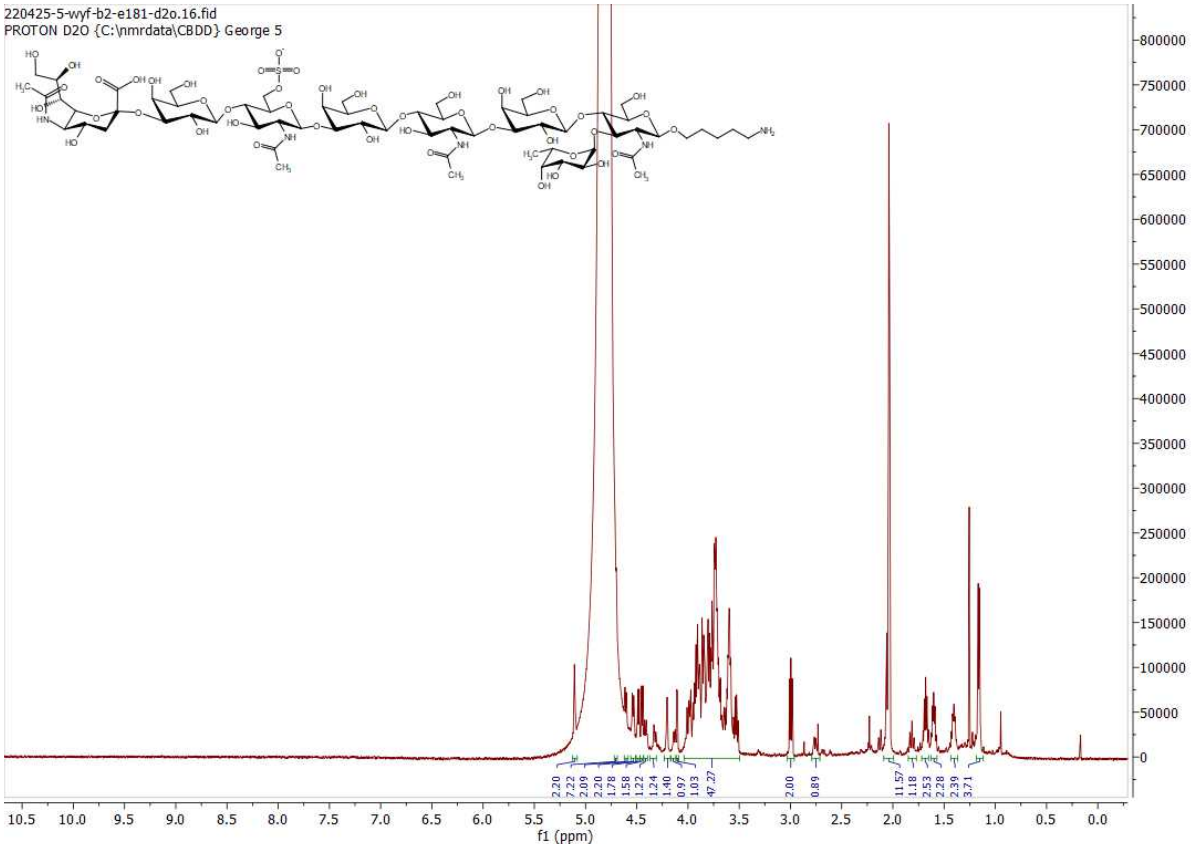

**$^{1}$H NMR of S6; 600MHz; $D_2O$**

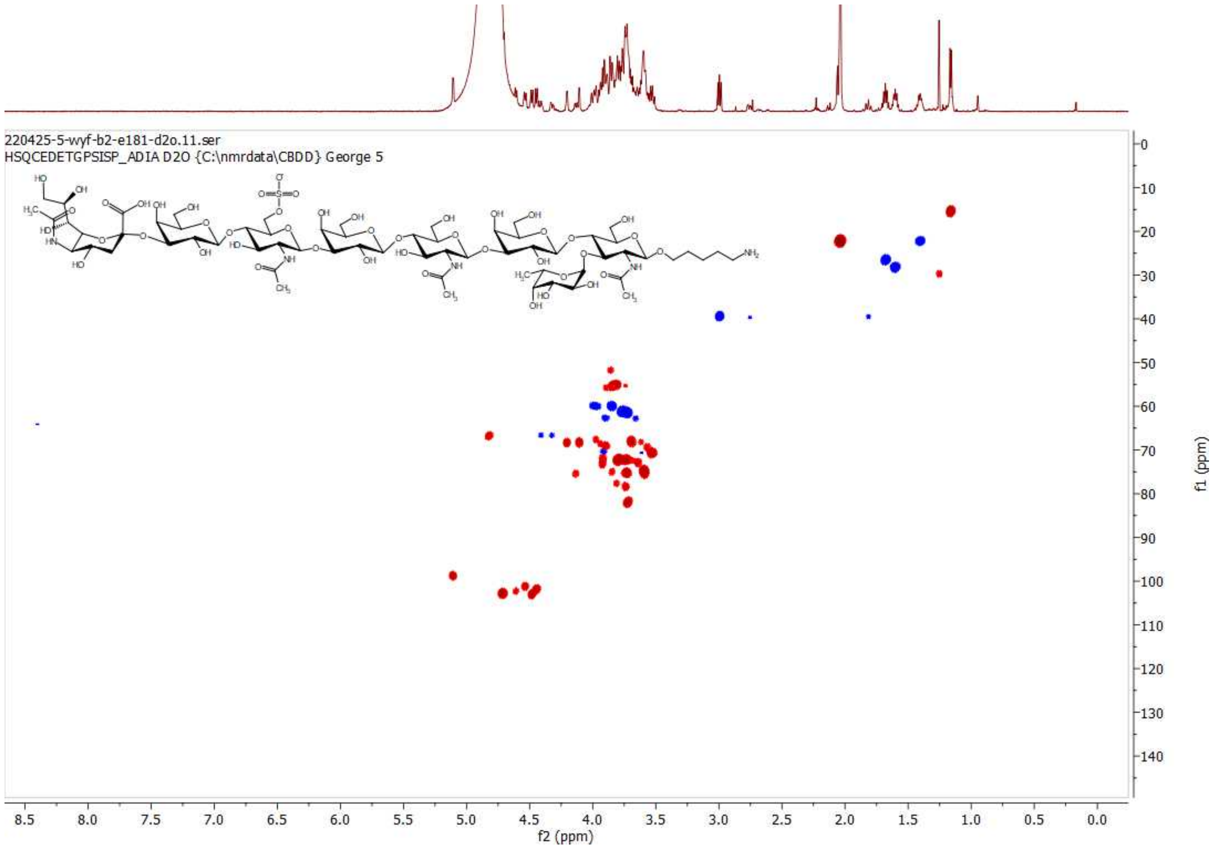

**HSQC of S6; 600 MHz/150 MHz, $D_2O$**

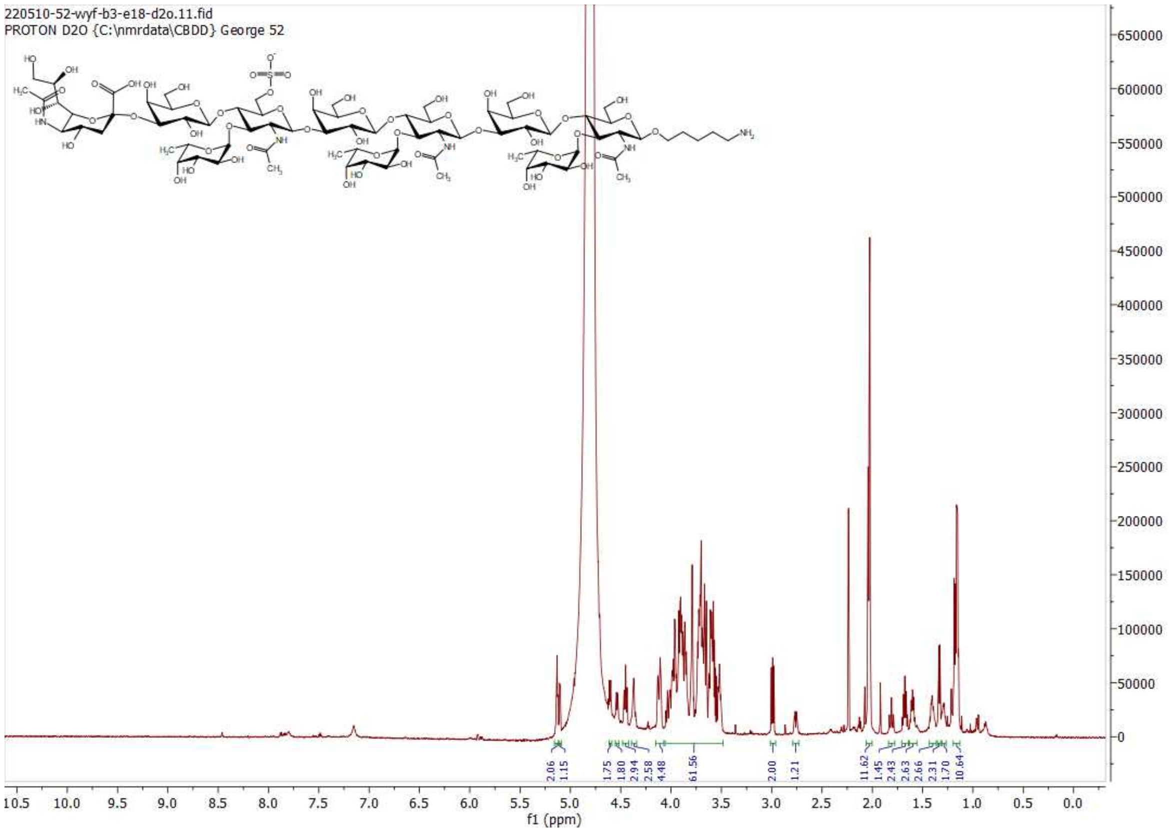

$^1$H NMR of X; 600MHz; $D_2O$

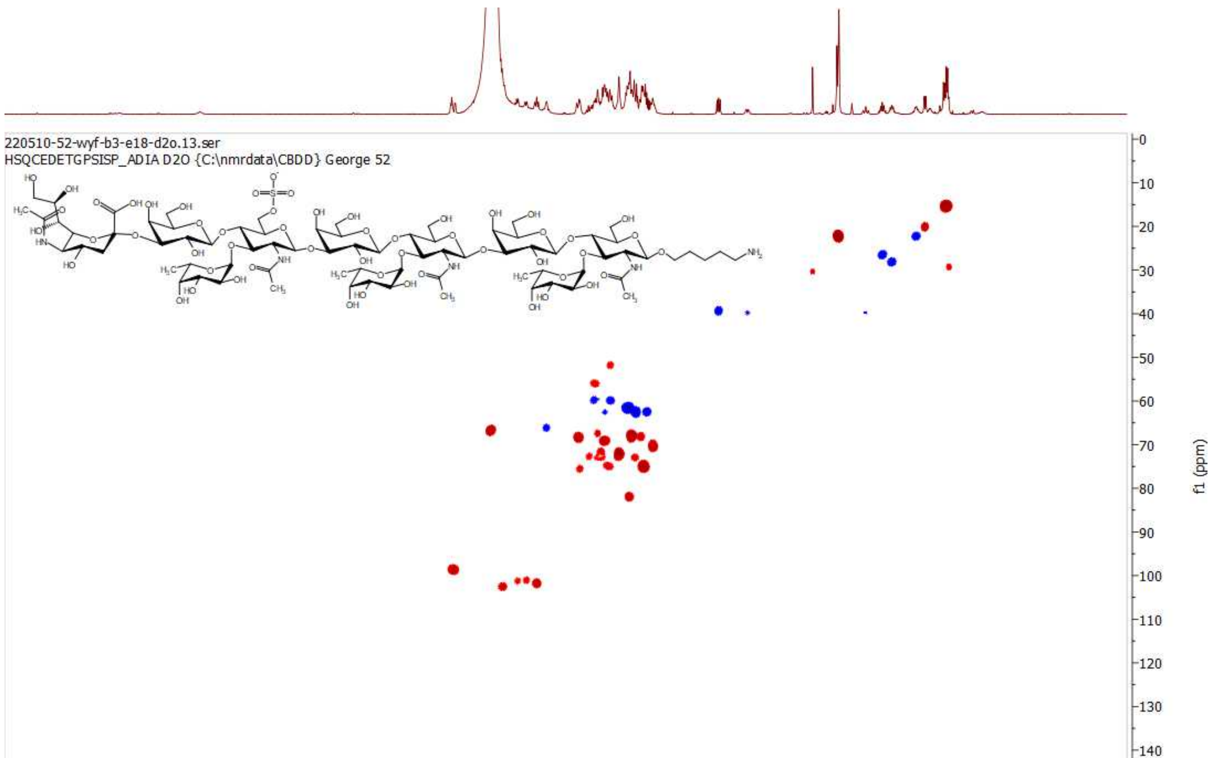

HSQC of X; 600 MHz/150 MHz, $D_2O$

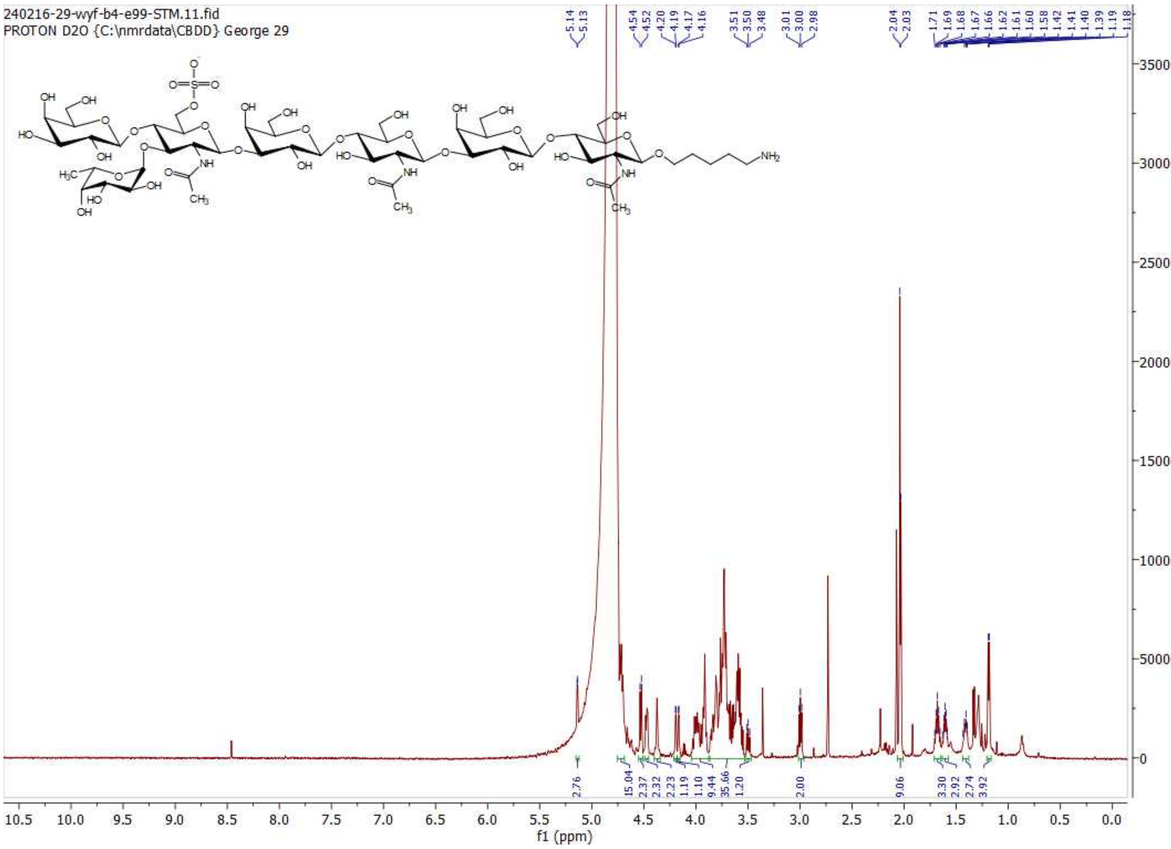

$^1$H NMR of S; 600MHz; $D_2O$

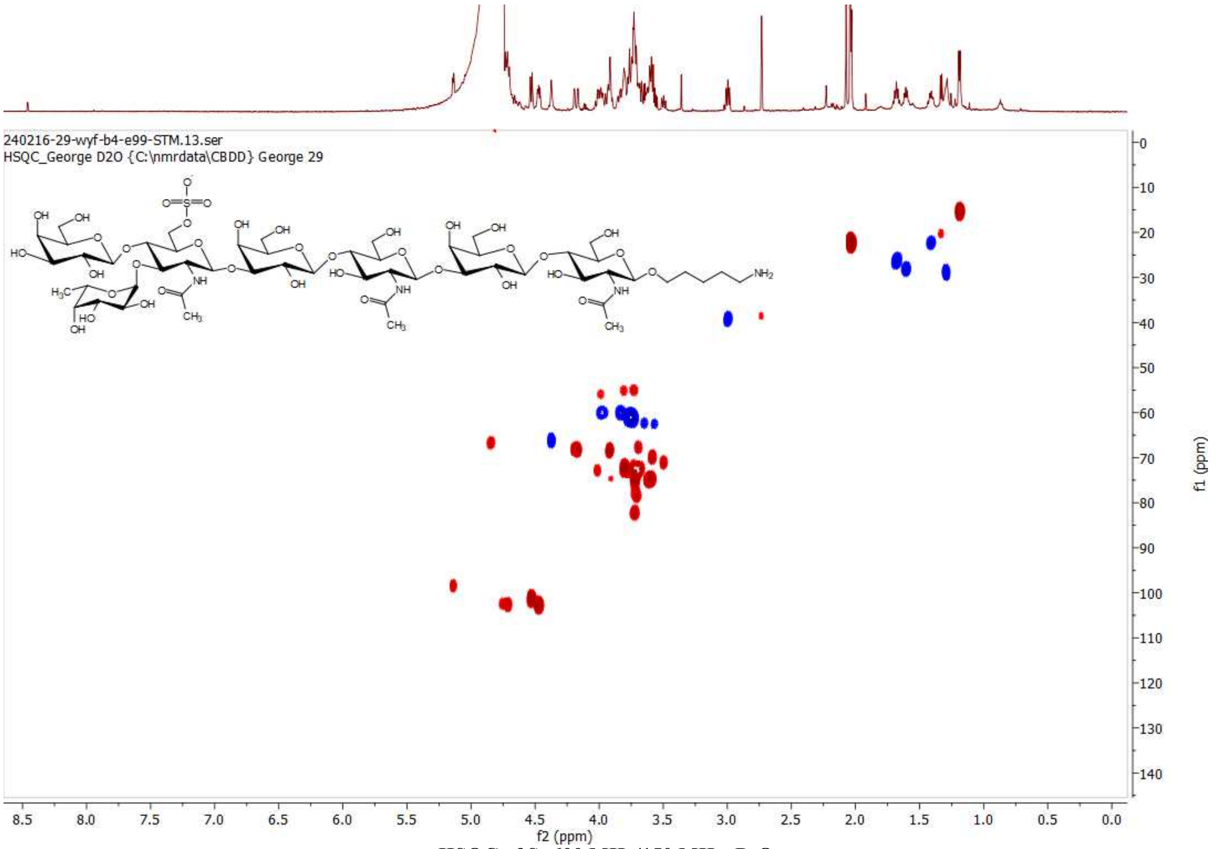

HSQC of S; 600 MHz/150 MHz, $D_2O$

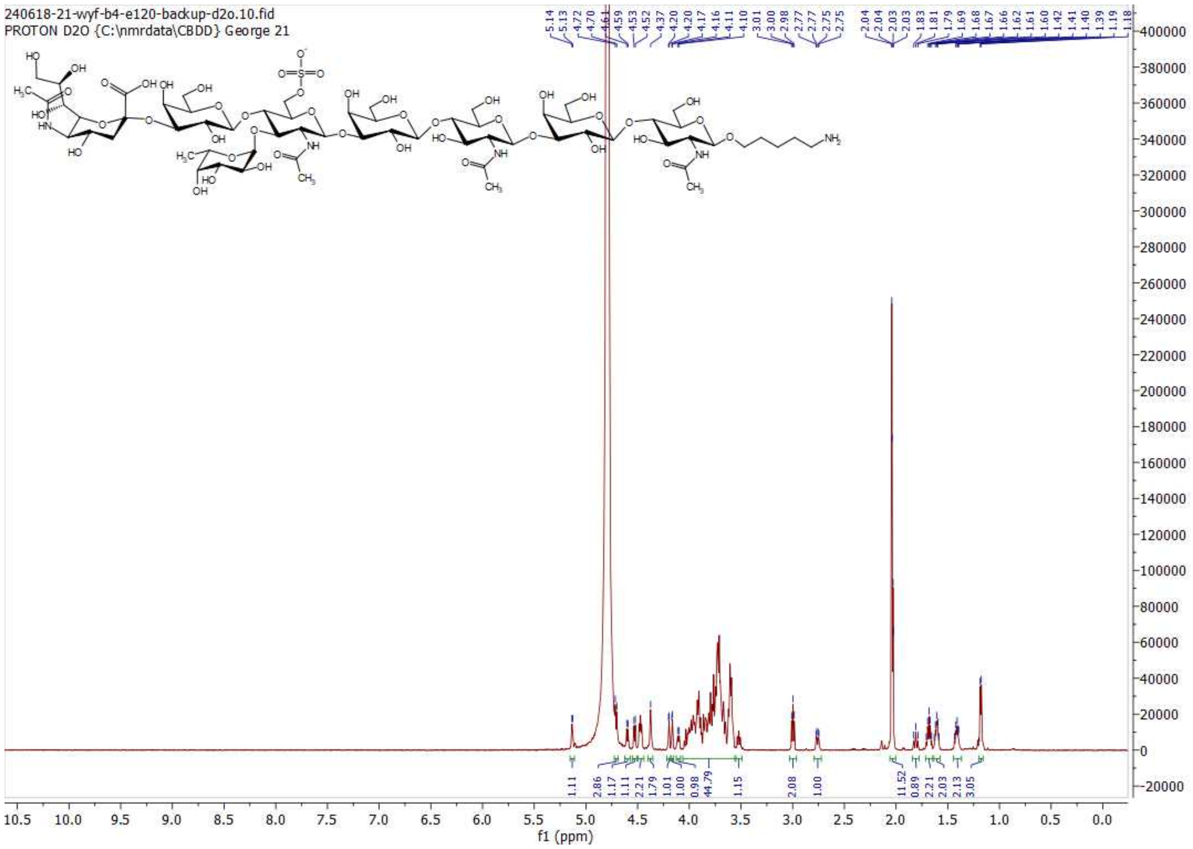

**[1]H NMR of Y; 600MHz; $D_2O$**

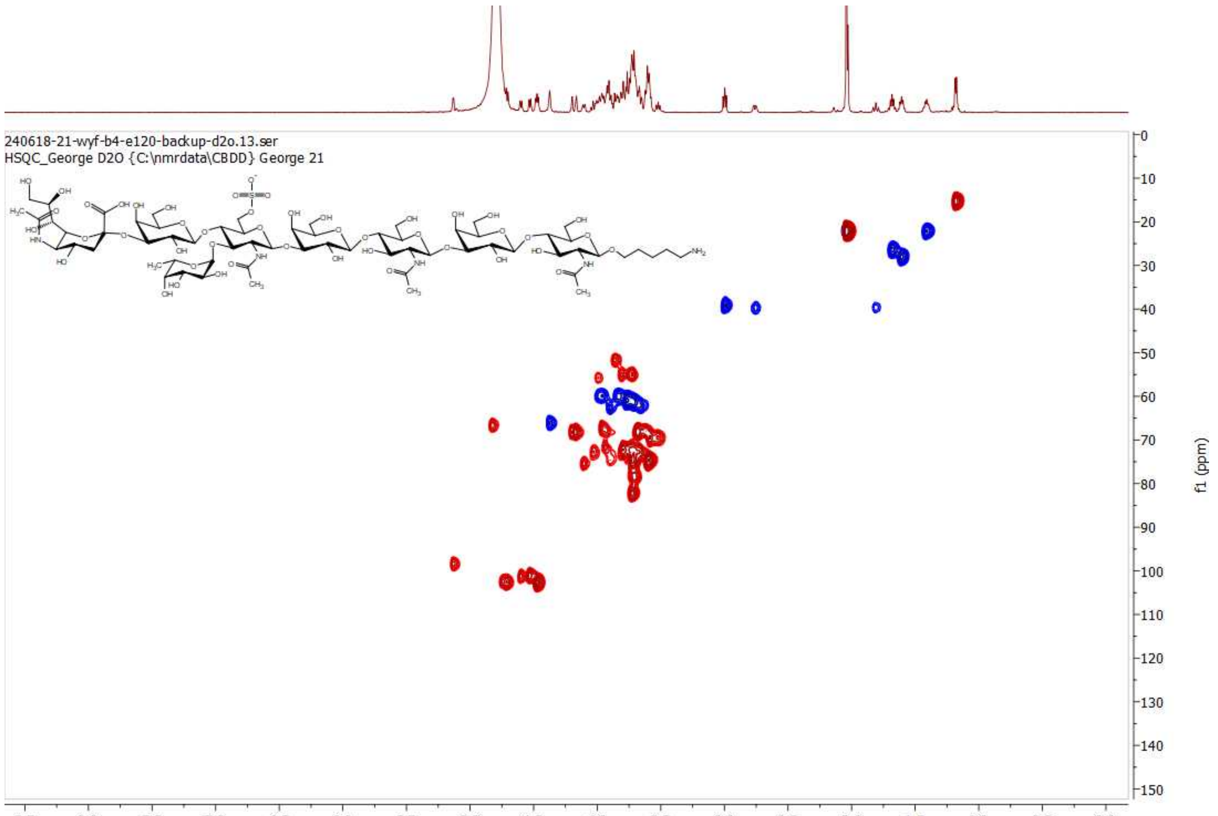

**HSQC of Y; 600 MHz/150 MHz, $D_2O$**

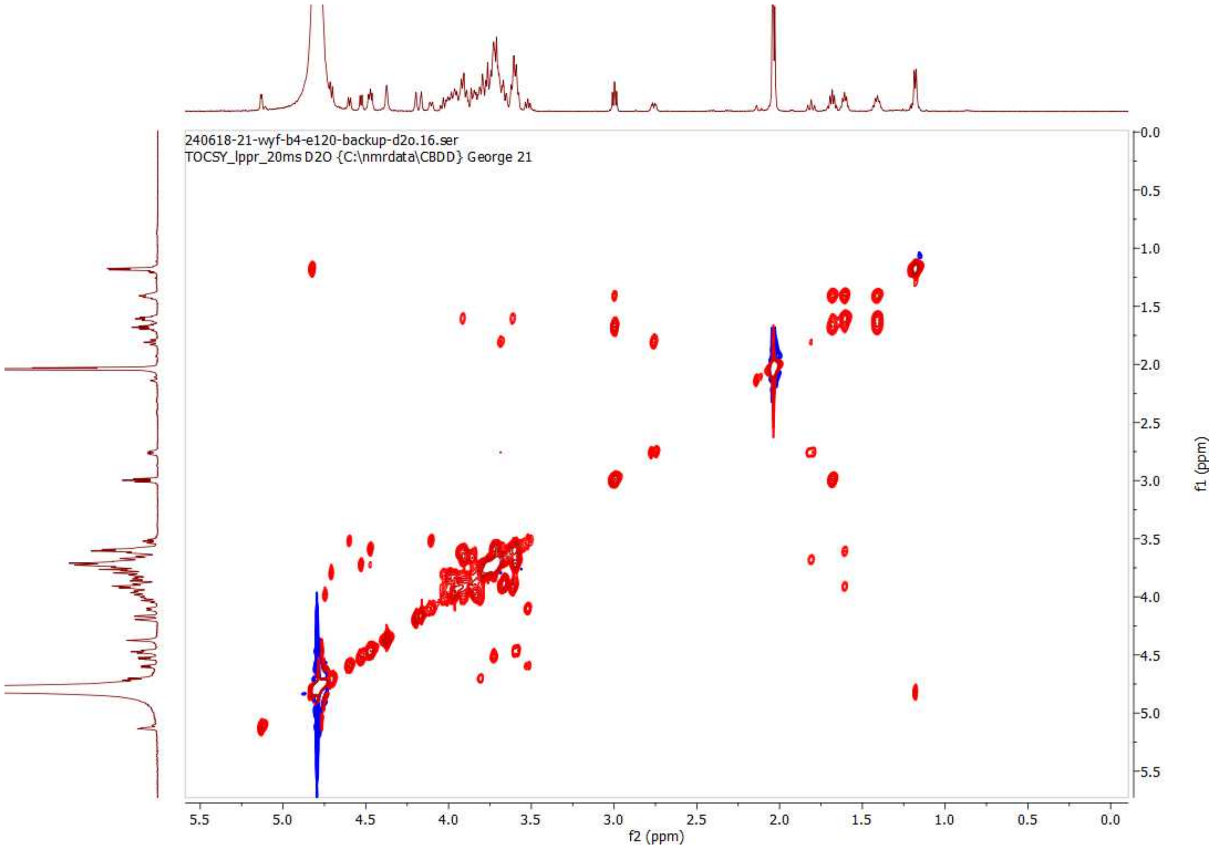

**TOCSY (20 ms) of Y; 600MHz; $D_2O$**

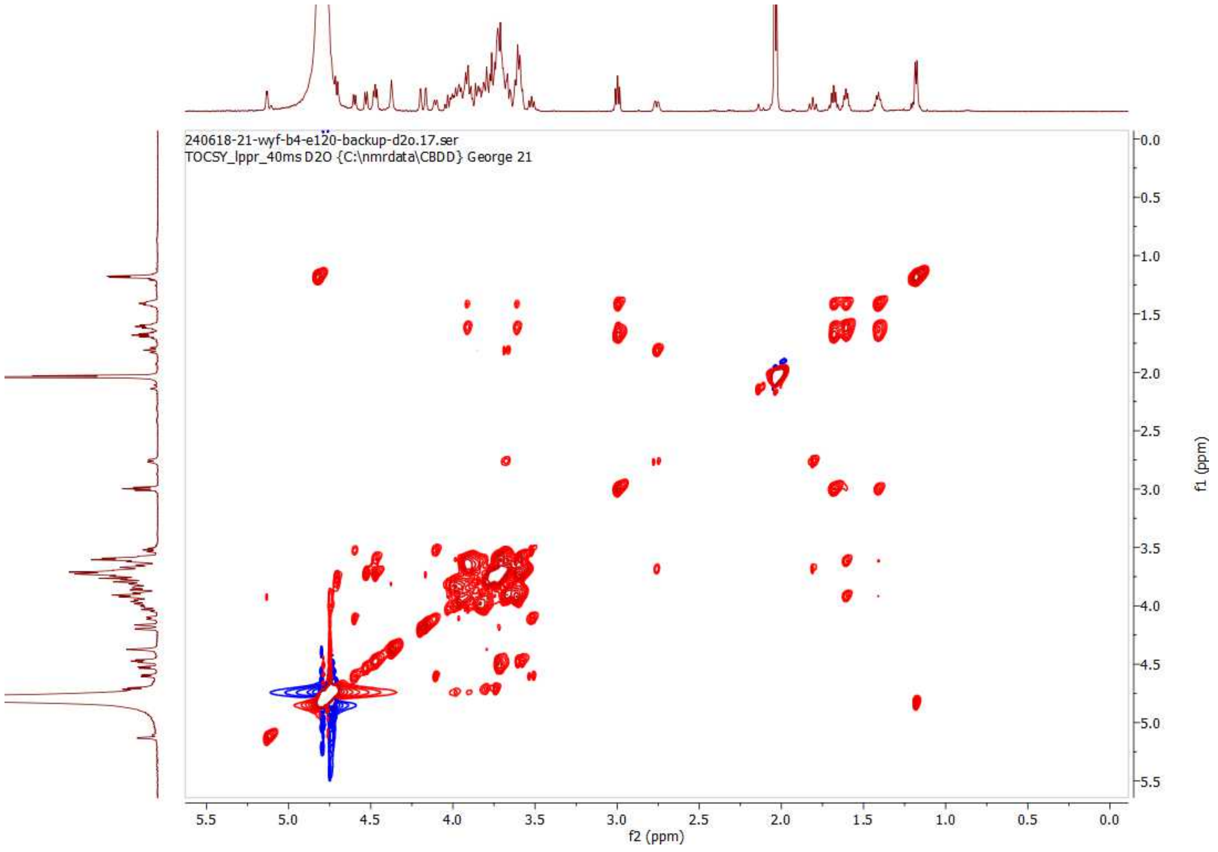

**TOCSY (40 ms) of Y; 600MHz; $D_2O$**

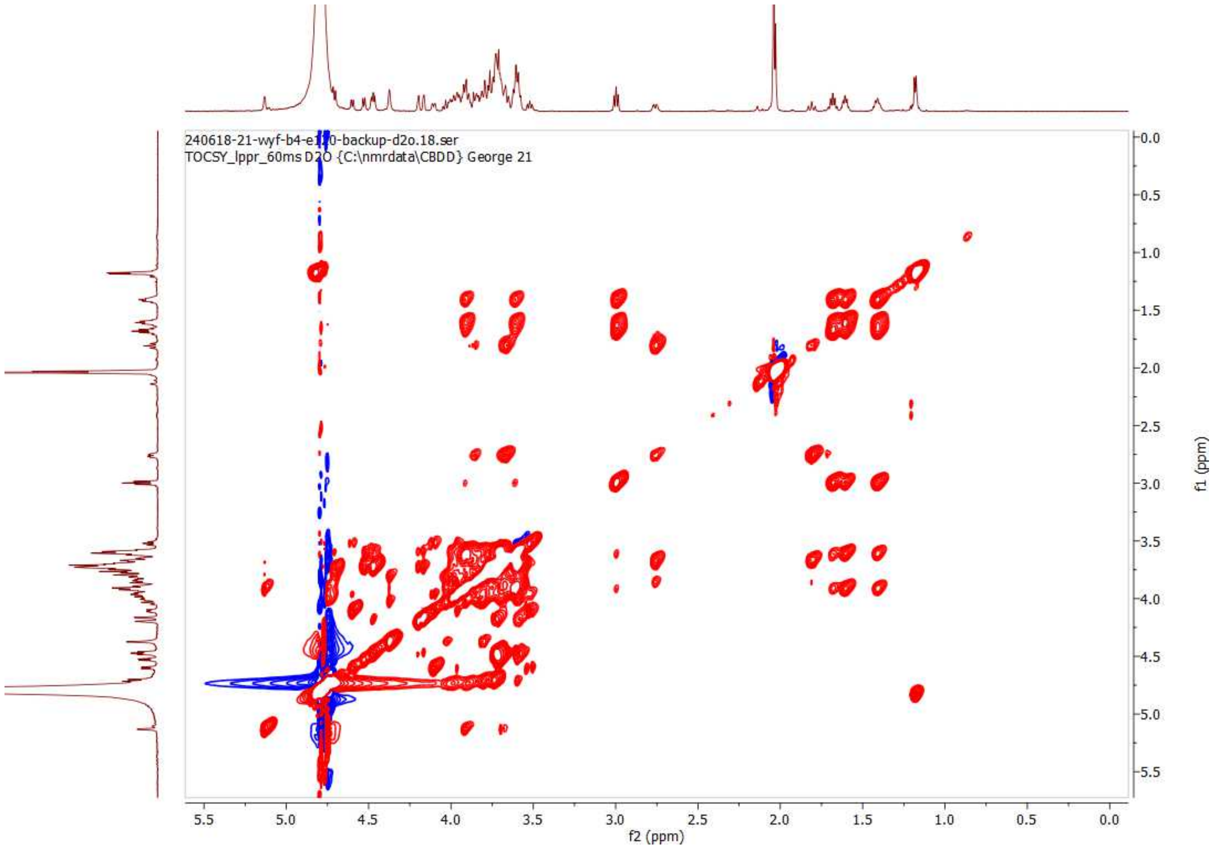

**TOCSY (60 ms) of Y; 600MHz; $D_2O$**

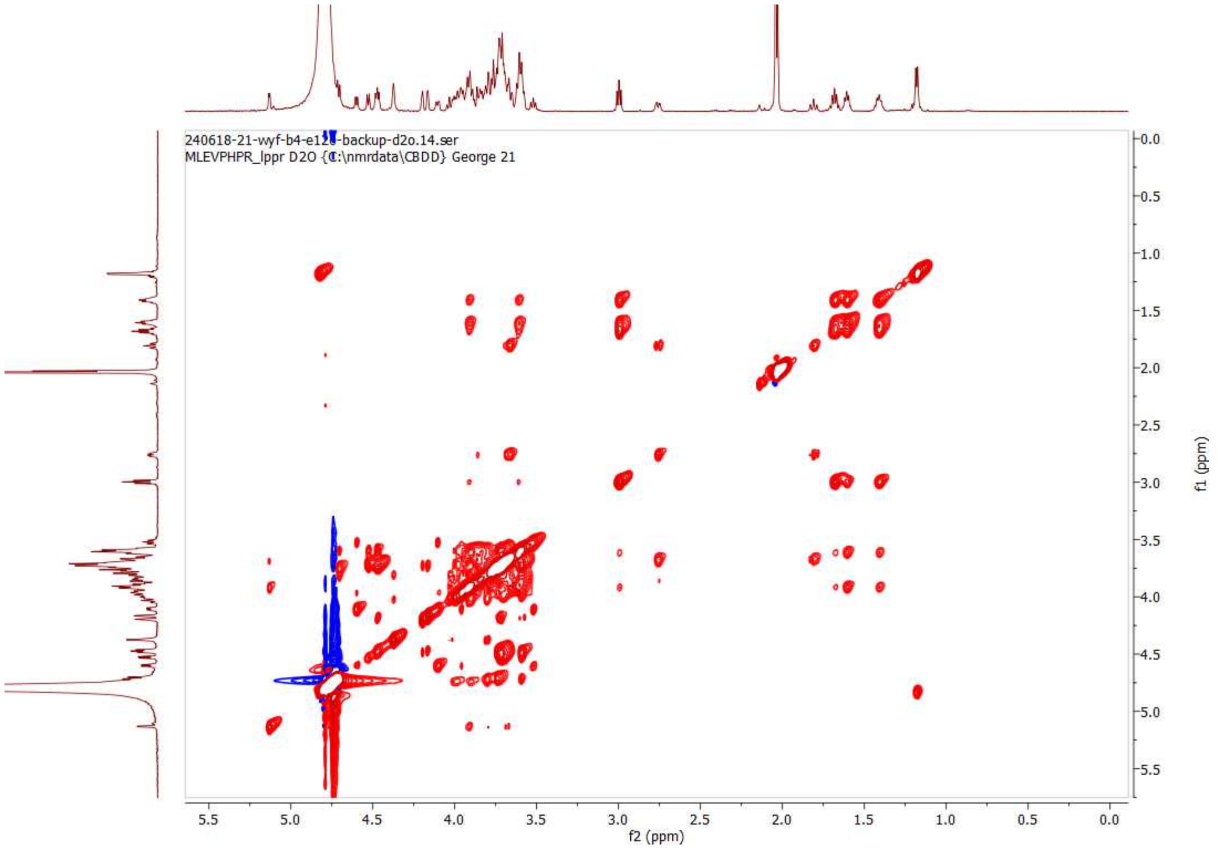

**TOCSY (80 ms) of Y; 600MHz; $D_2O$**

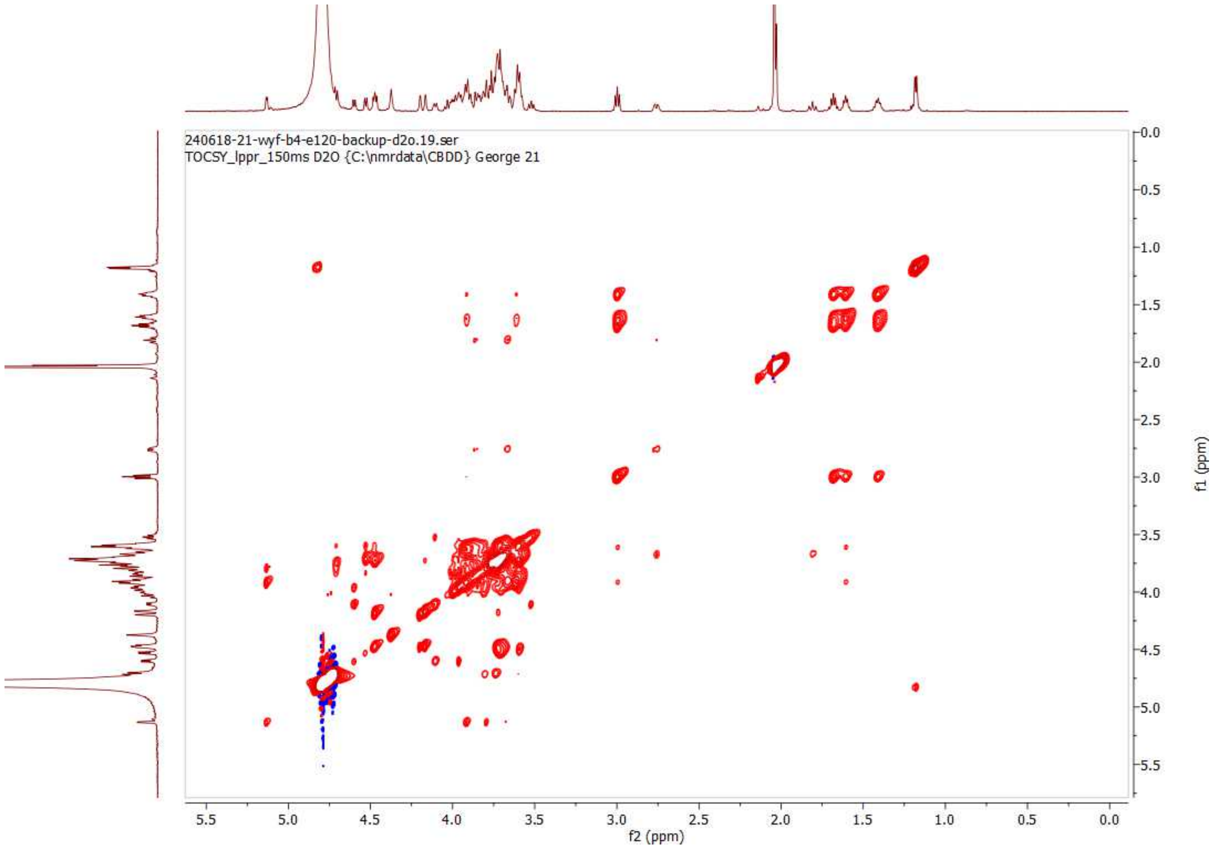

**TOCSY (150 ms) of Y; 600MHz; $D_2O$**

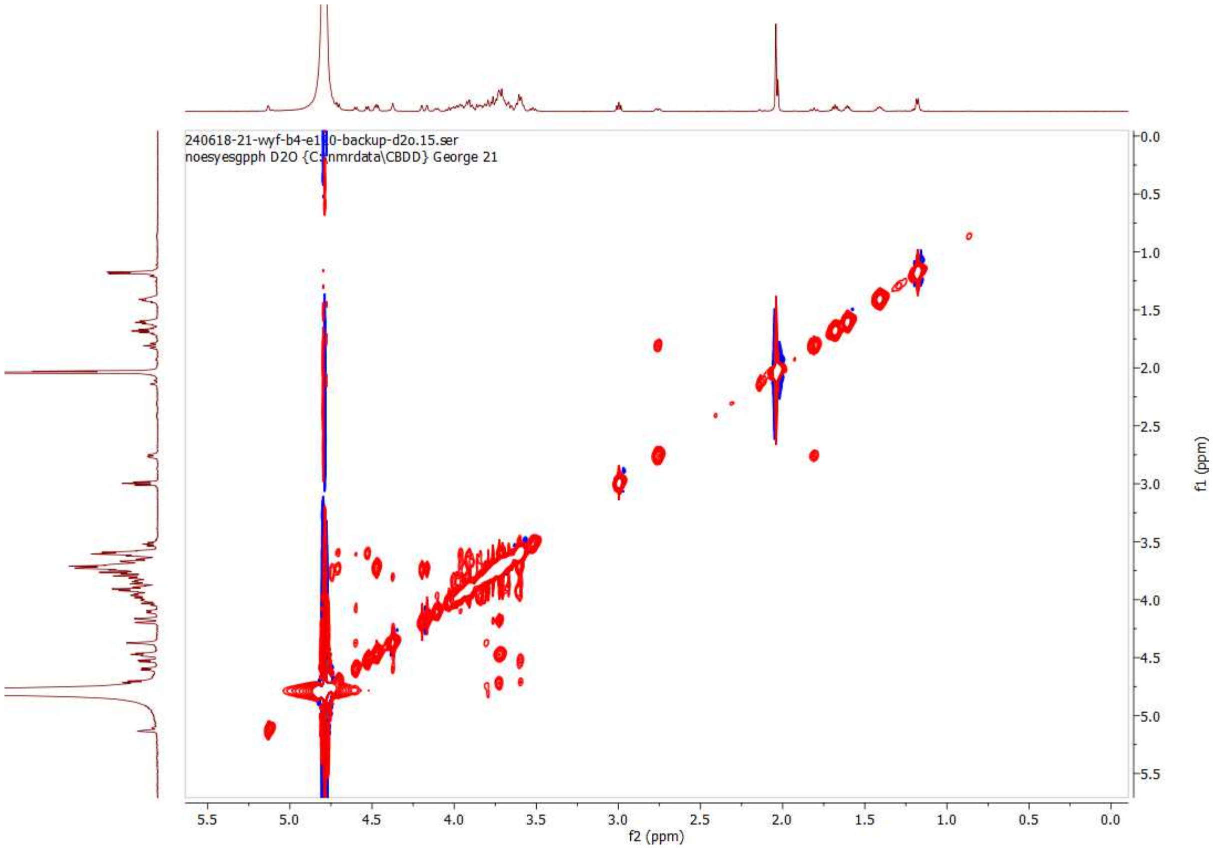

**NOESY (300 ms) of Y; 600 MHz, $D_2O$**

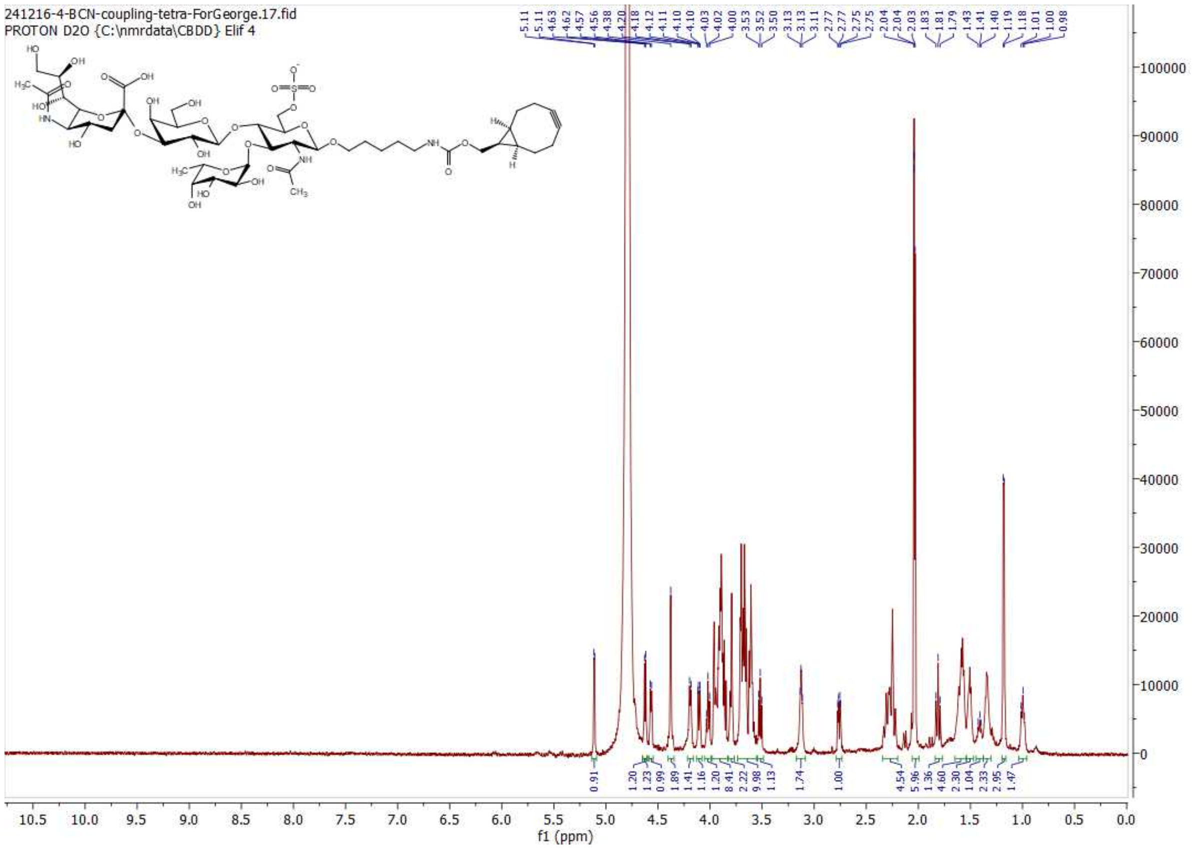

**$^{1}$H NMR of 21; 600MHz; $D_2O$**

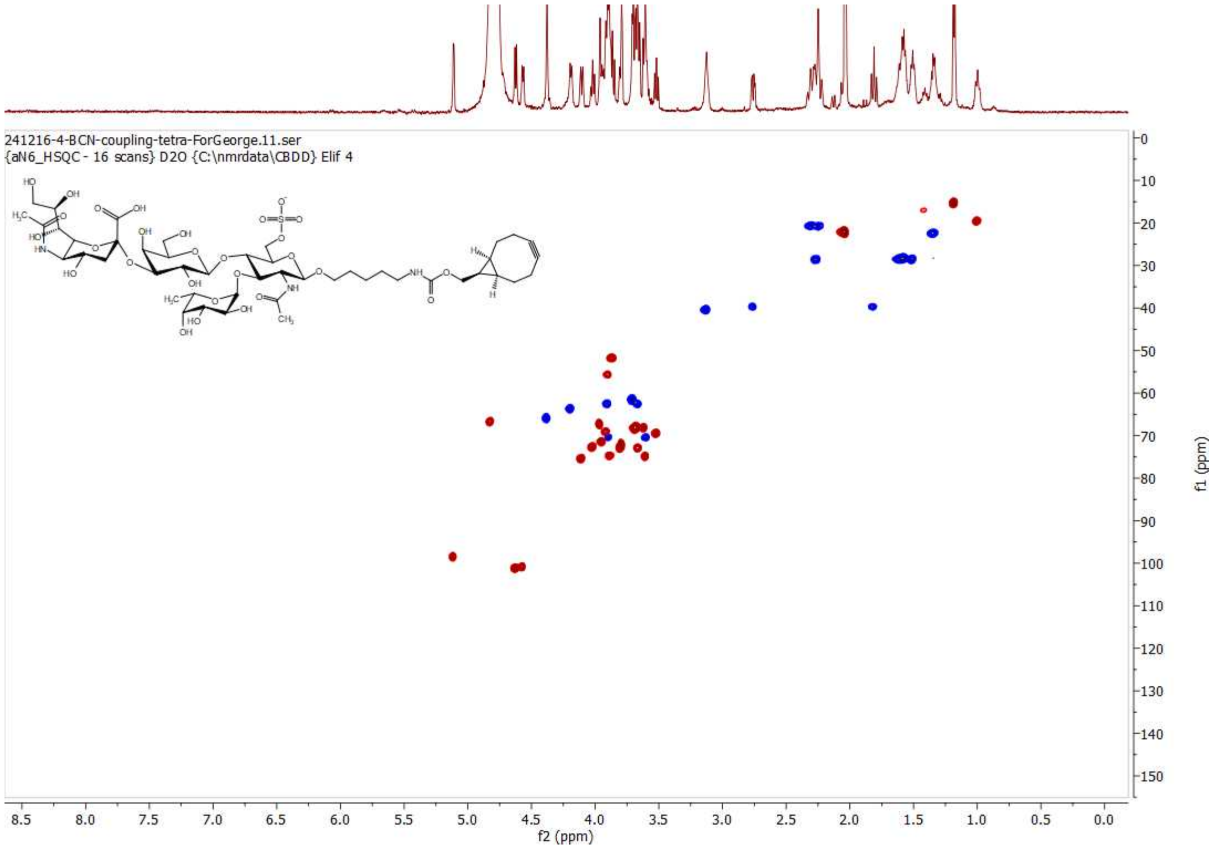

**HSQC of 21; 600 MHz/150 MHz, $D_2O$**

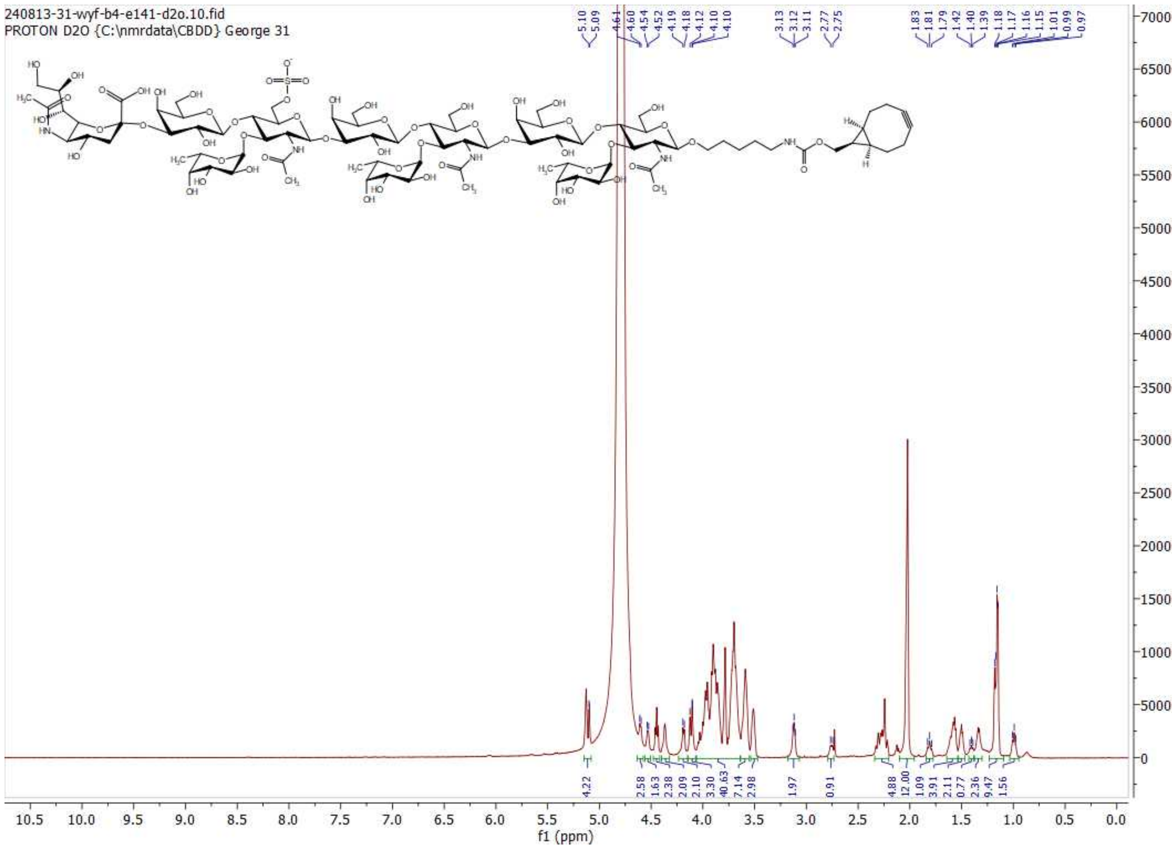

**$^{1}$H NMR of 23; 600MHz; $D_2O$**

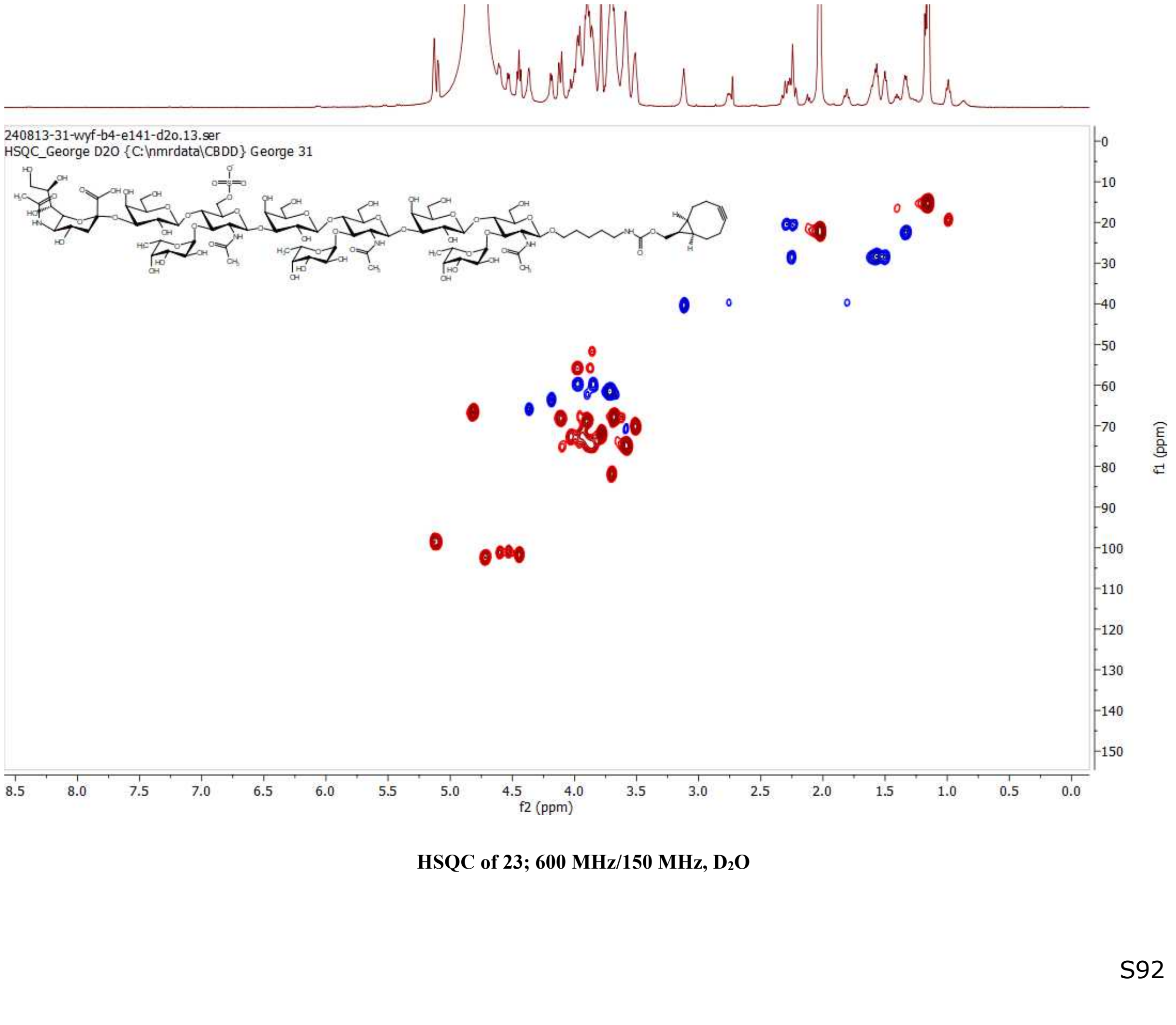

**HSQC of 23; 600 MHz/150 MHz, $D_2O$**

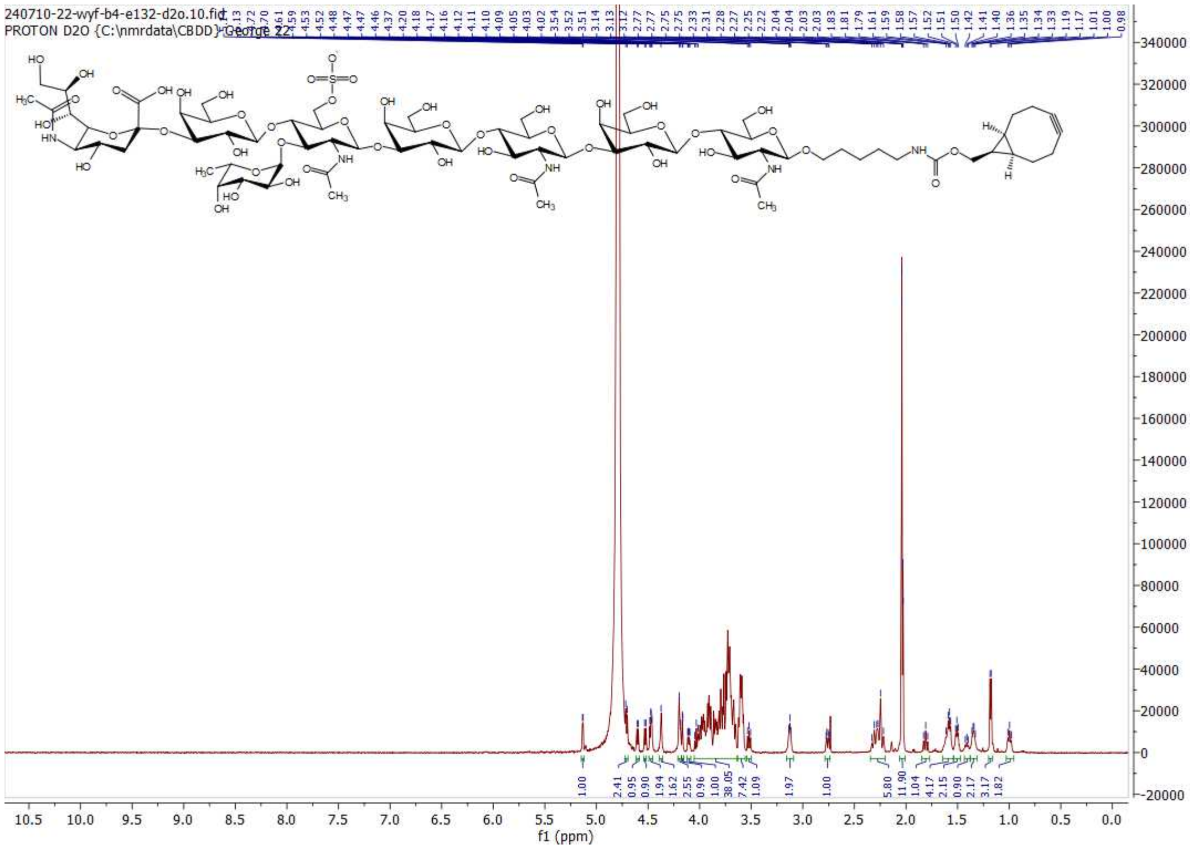

$^{1}$H NMR of 22; 600MHz; $D_2O$

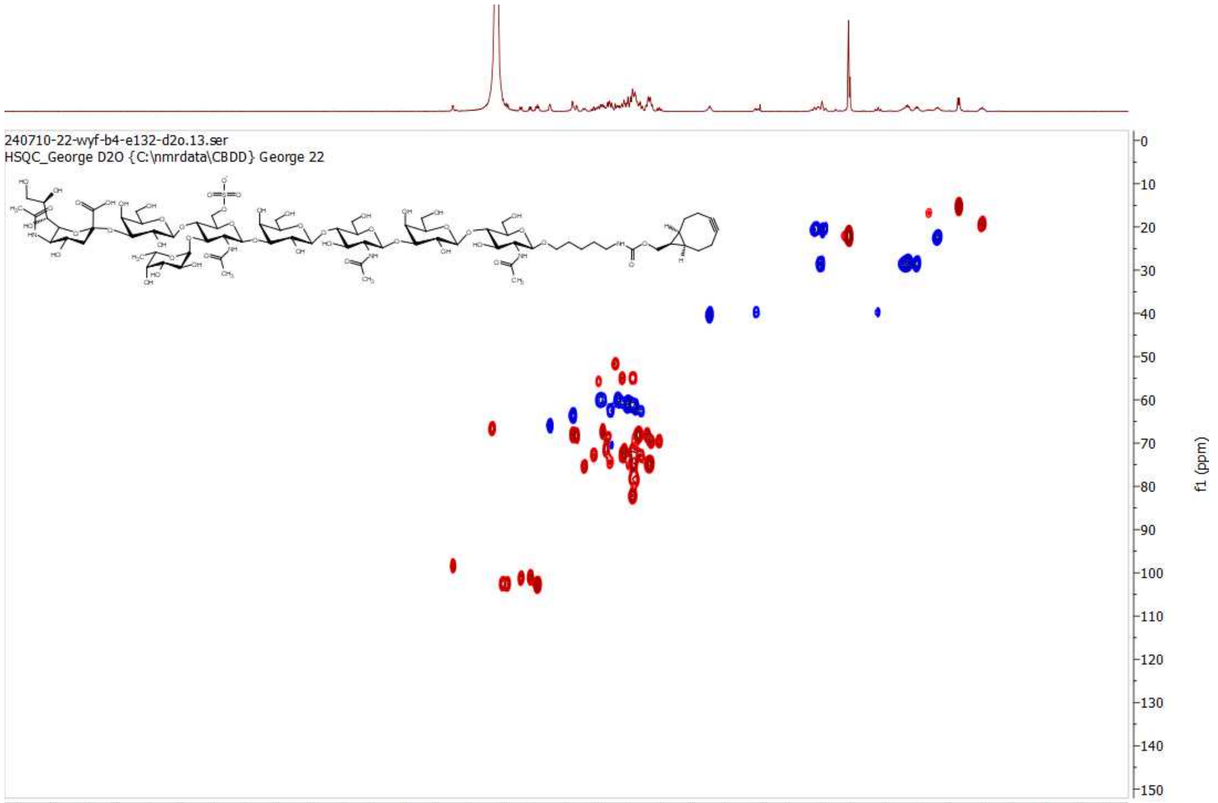

HSQC of 22; 600 MHz/150 MHz, $D_2O$

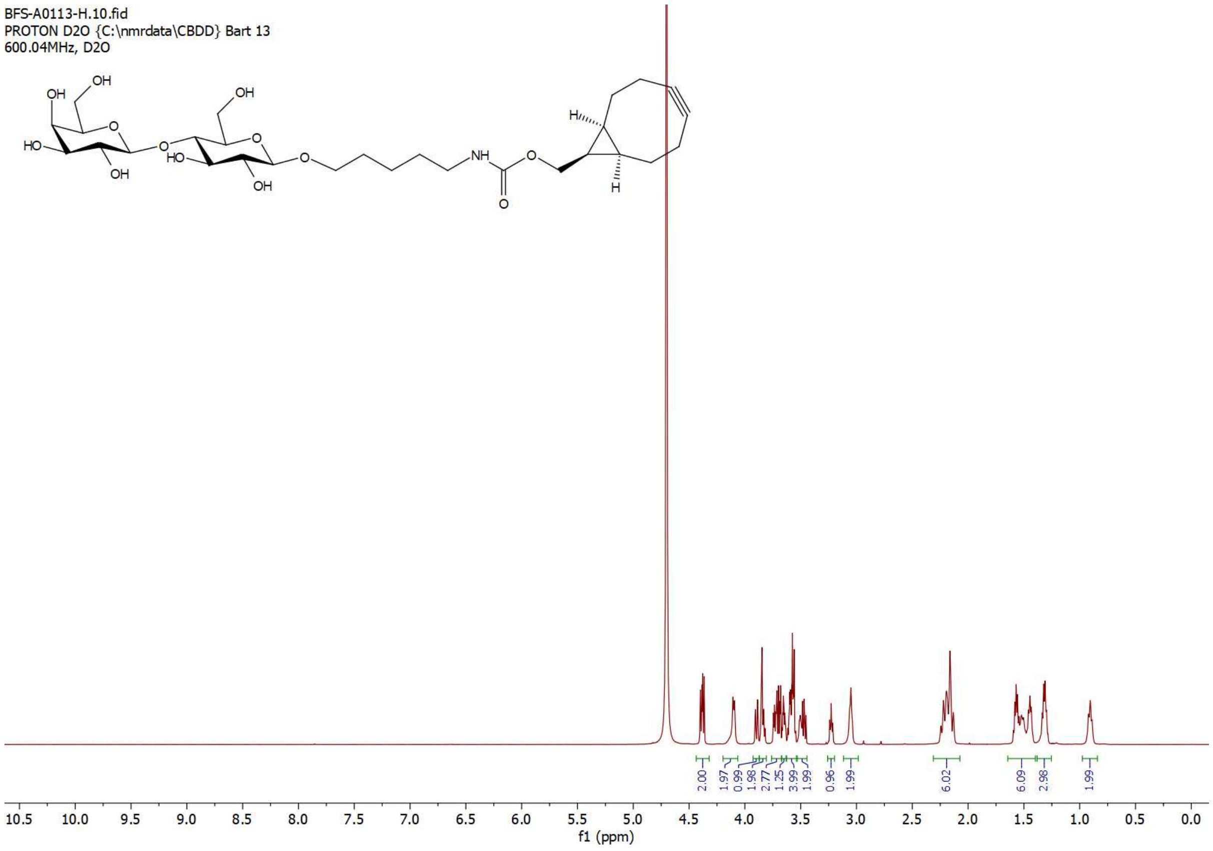

[1]H NMR of 19; 600MHz; $D_2O$

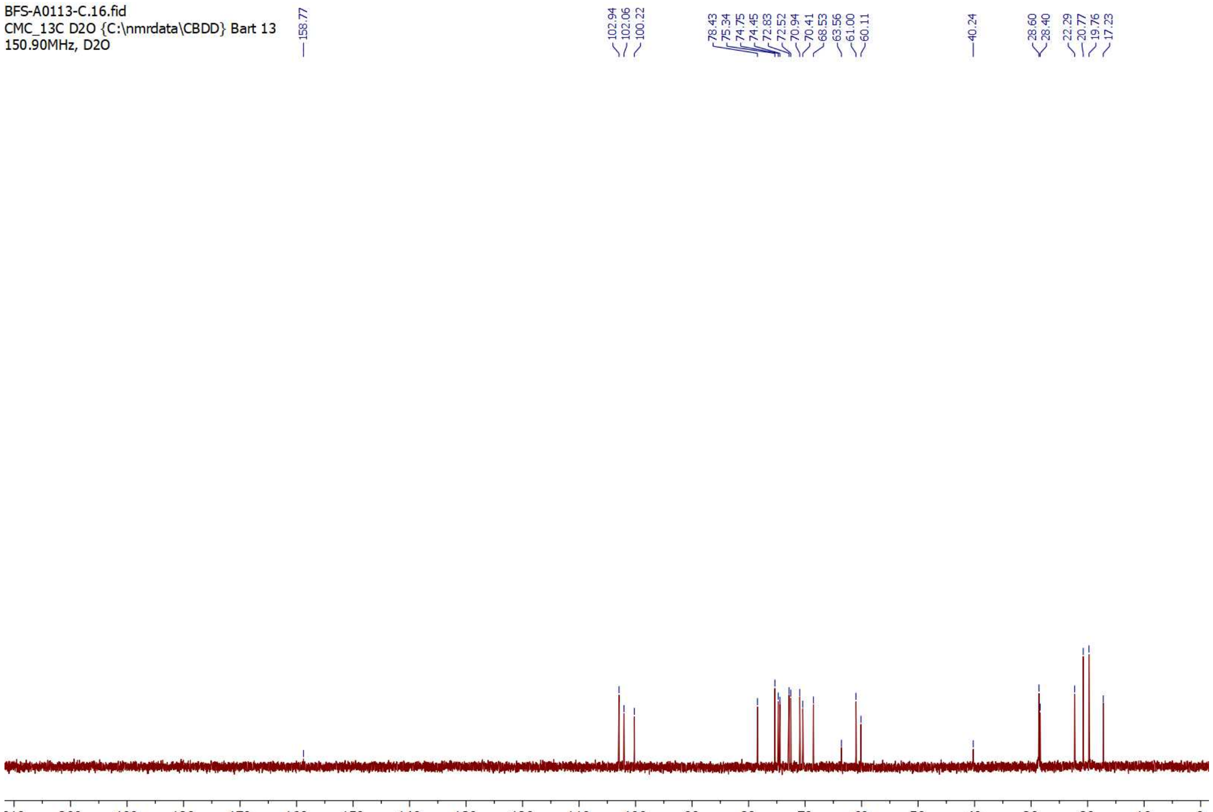

[13]C NMR of 19; 150MHz; $D_2O$

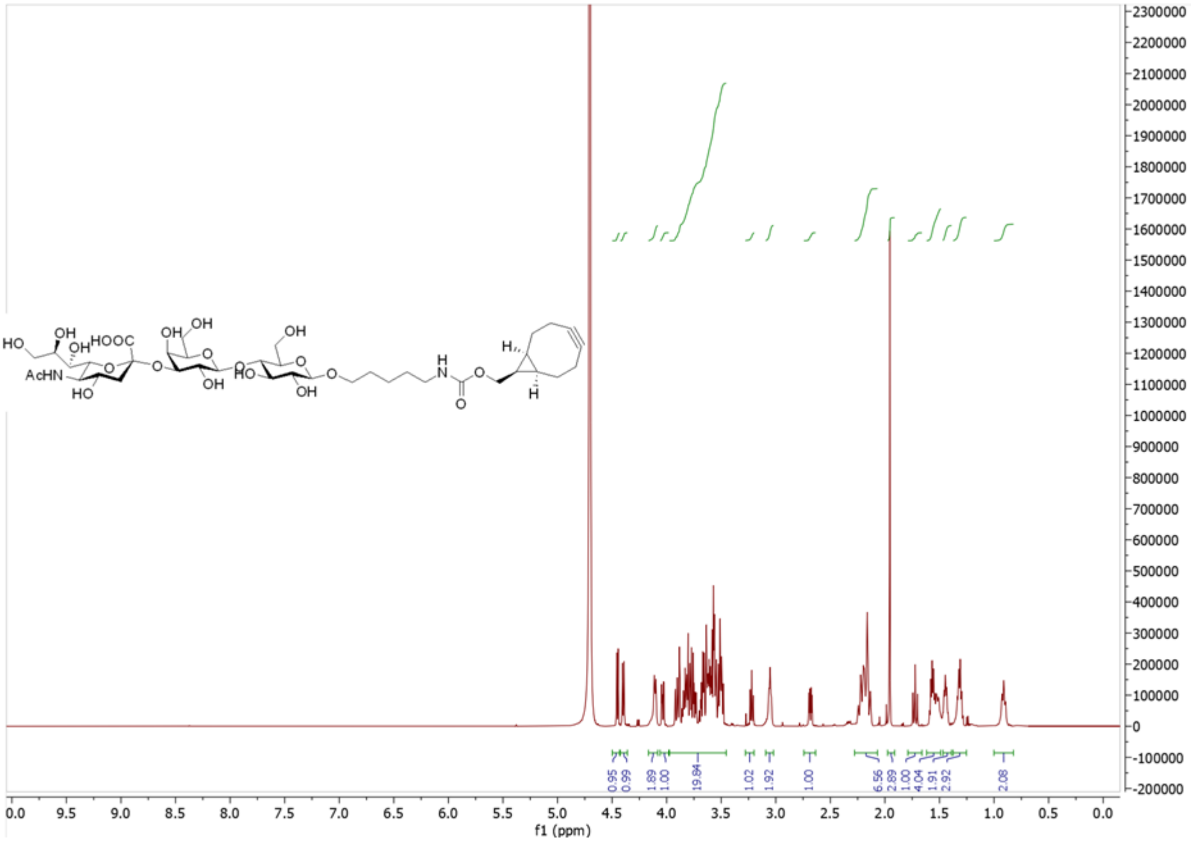

[1]H NMR of 20; 600MHz; $D_2O$

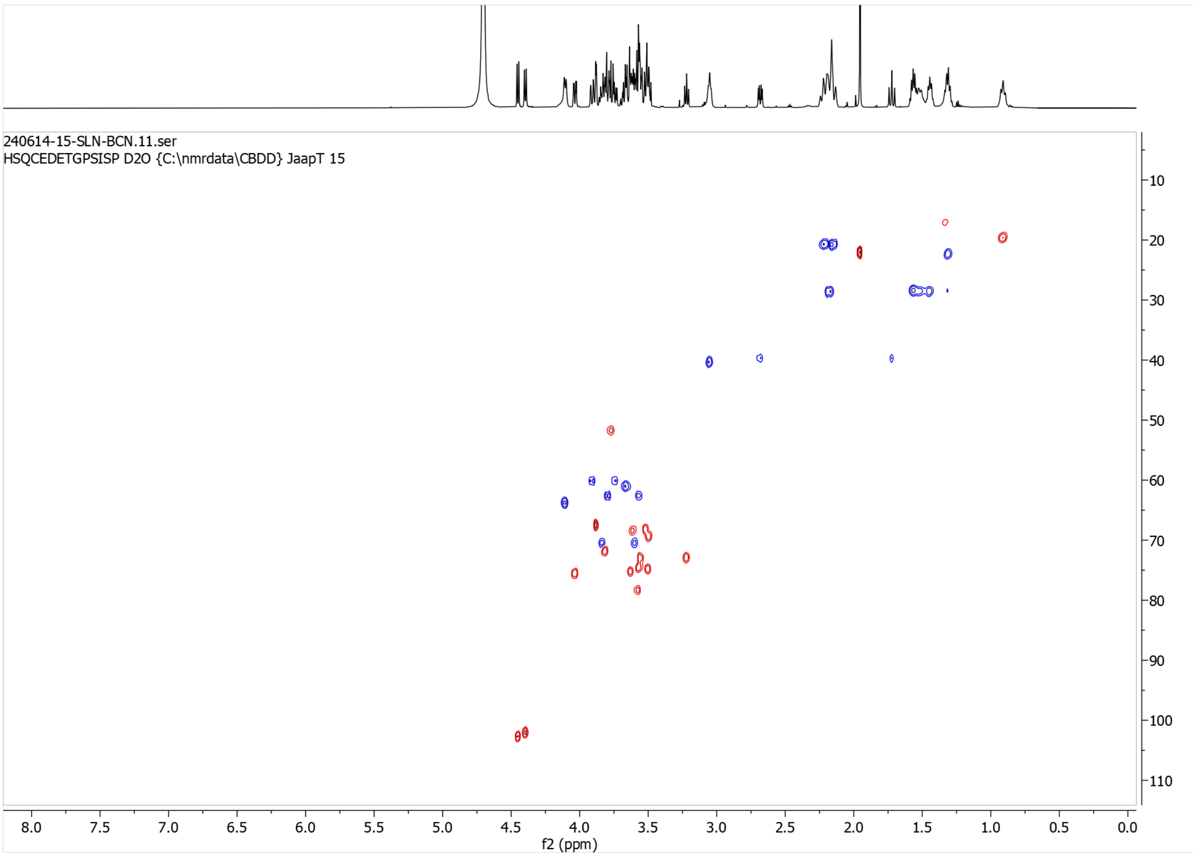

HSQC of 20; 600 MHz/150 MHz, $D_2O$

### Generic Display Report

**Analysis Info**

| | | | |
|---|---|---|---|
| Analysis Name | D:\Data\george\Compound 3 chapter 5000003.d | Acquisition Date | 7/24/2026 2:05:55 PM |
| Method | ESI_pos_300-3000mz_wide.m | Operator | wim |
| Sample Name | | Instrument | micrOTOF |
| Comment | | | |

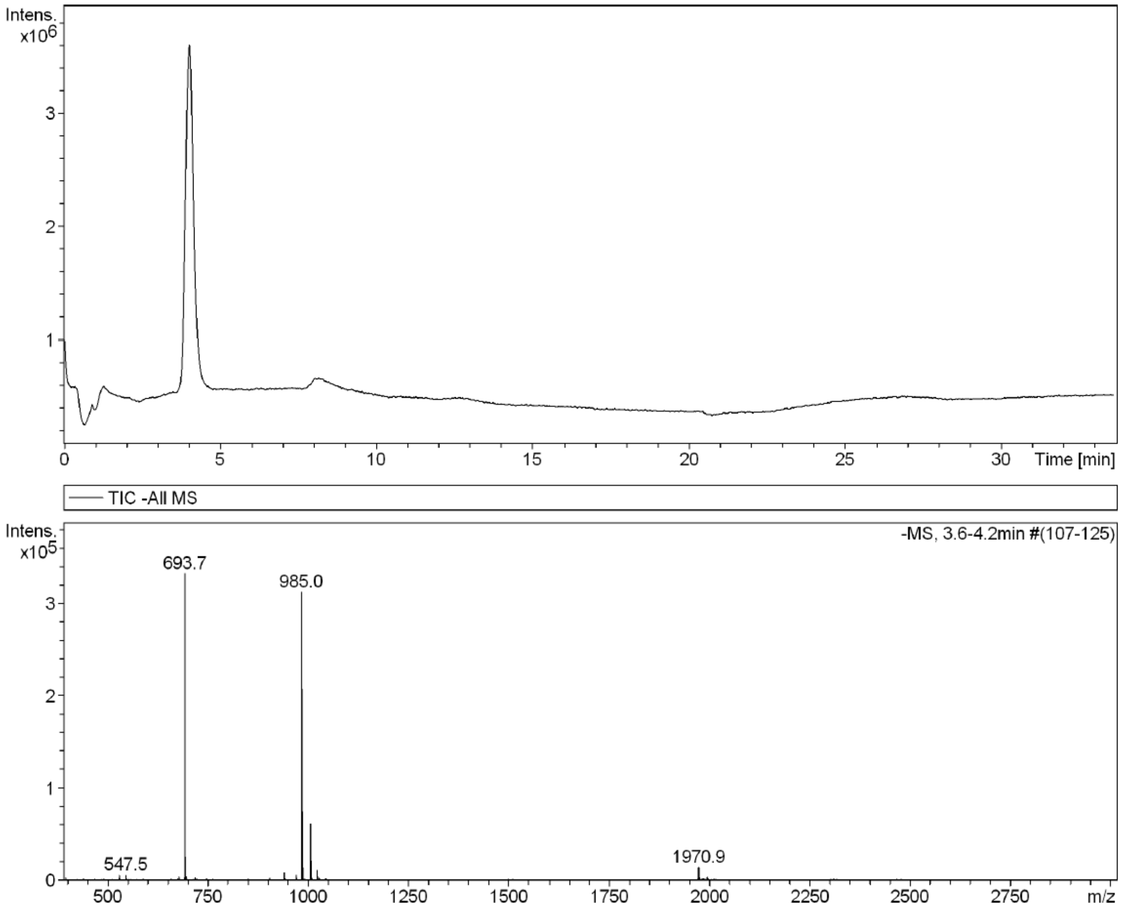

**LC-MS for purity of compound G.** Purity was determined using a SeQuant Zic HILIC guard column (20x2.1 mm) coupled to a Bruker micrOTOF-QII MS system running a gradient of 10% 10 mM $NH_4HCO_3$ in MeCN (buffer B) and MeCN in 90% 10 mM $NH_4HCO_3$ (buffer A).

#### Generic Display Report

**Analysis Info**

| | | | |
|---|---|---|---|
| Analysis Name | D:\Data\george\Compound 3 chapter 5000006.d | Acquisition Date | 7/24/2026 2:43:00 PM |
| Method | ESI_pos_300-3000mz_wide.m | Operator | wim |
| Sample Name | | Instrument | micrOTOF |
| Comment | | | |

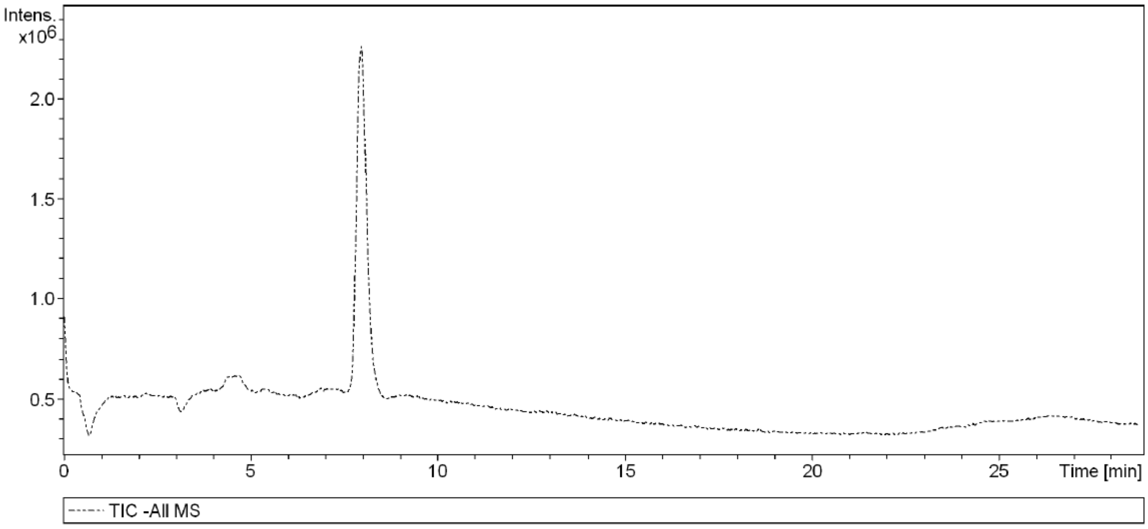

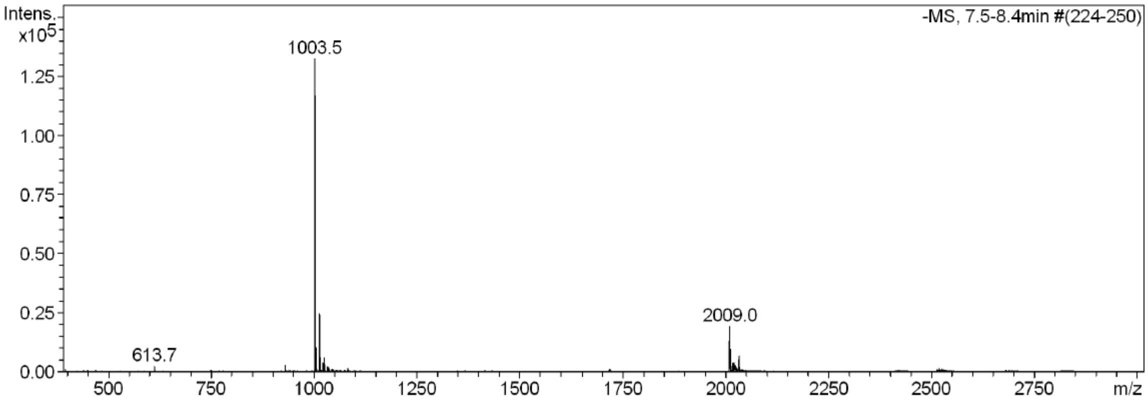

**LC-MS for purity of compound X.** Purity was determined using a SeQuant Zic HILIC guard column (20x2.1 mm) coupled to a Bruker micrOTOF-QII MS system running a gradient of 10% 10 mM $NH_4HCO_3$ in MeCN (buffer B) and MeCN in 90% 10 mM $NH_4HCO_3$ (buffer A).

#### Generic Display Report

**Analysis Info**

| | | | |
|---|---|---|---|
| Analysis Name | D:\Data\george\Compound 4 chapter 5000007.d | Acquisition Date | 7/24/2026 5:23:43 PM |
| Method | ESI_pos_300-3000mz_wide.m | Operator | wim |
| Sample Name | | Instrument | micrOTOF |
| Comment | | | |

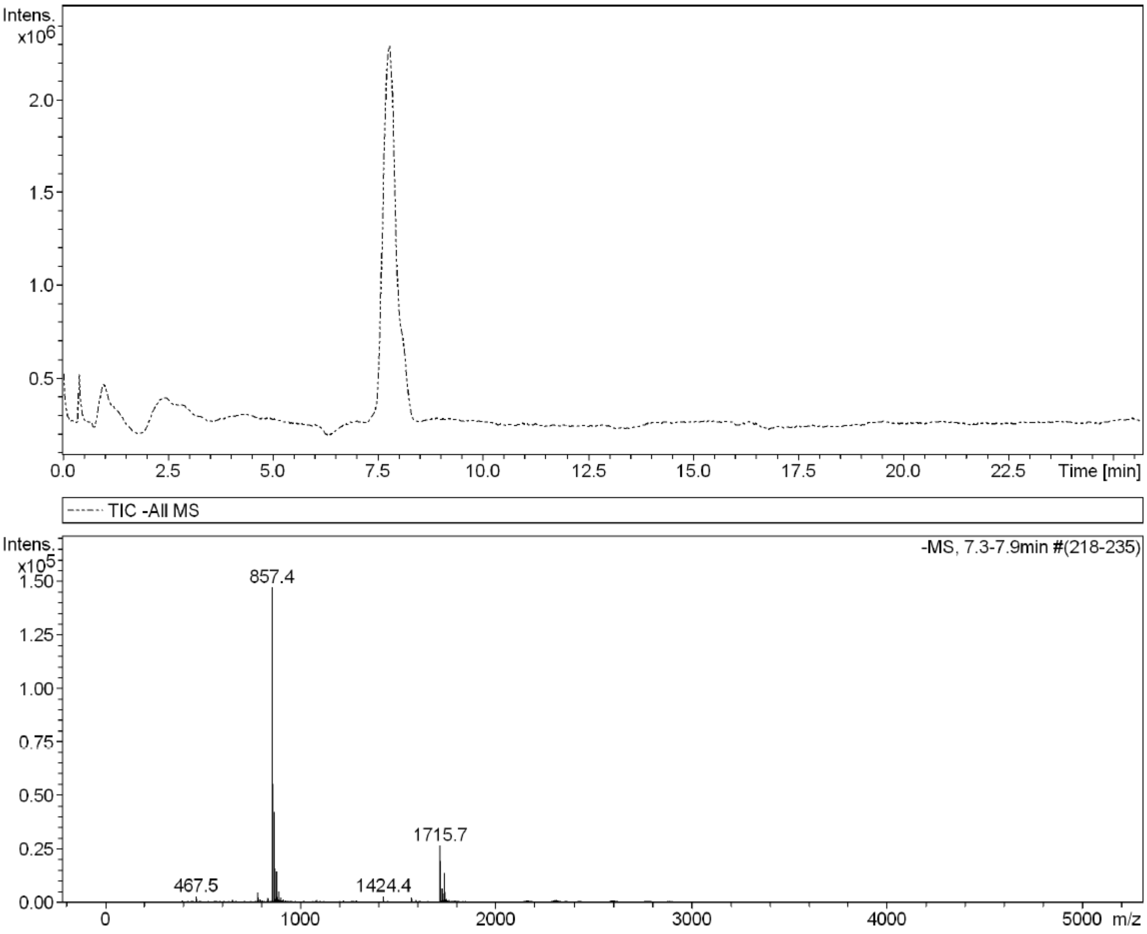

**LC-MS for purity of compound Y.** Purity was determined using a SeQuant Zic HILIC guard column (20x2.1 mm) coupled to a Bruker micrOTOF-QII MS system running a gradient of 10% 10 mM $NH_4HCO_3$ in MeCN (buffer B) and MeCN in 90% 10 mM $NH_4HCO_3$ (buffer A).

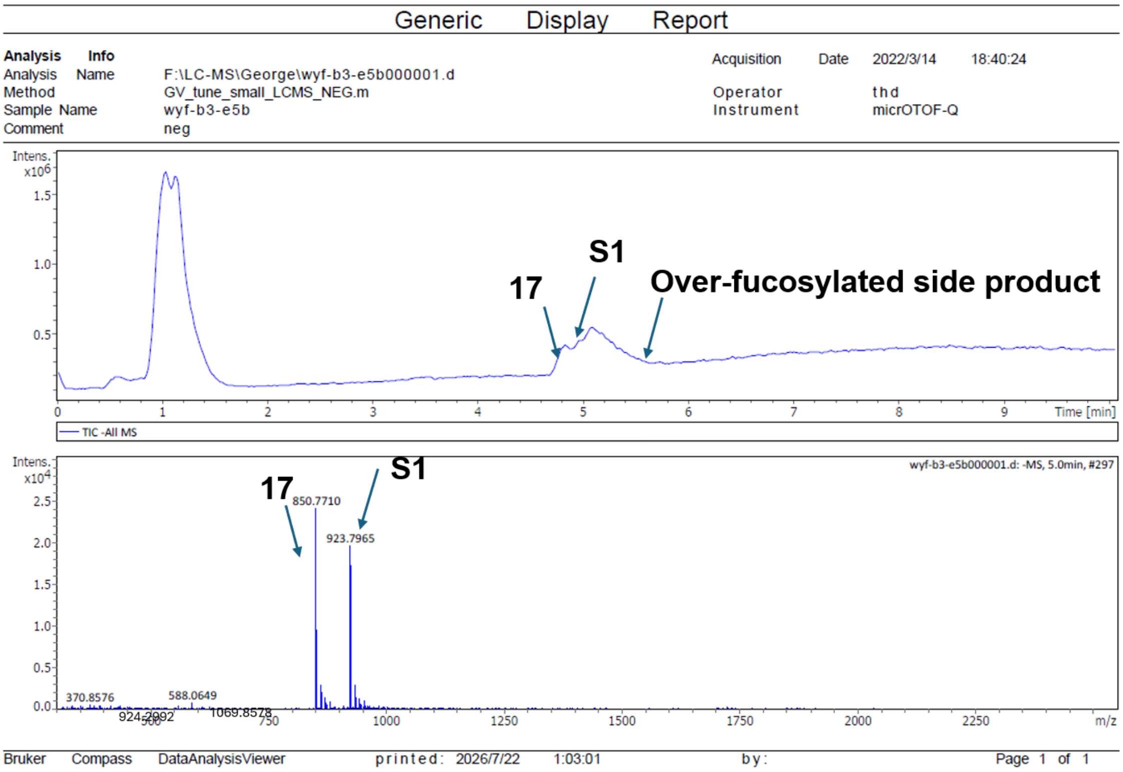

**LC-MS for Scheme S1a, FUT5 rate control.** Reaction progress for carbohydrates was monitored using a SeQuant Zic HILIC guard column (20x2.1 mm) coupled to a Bruker micrOTOF-QII MS system running a gradient of 90% MeCN:$H_2O$ to 50% MeCN:$H_2O$ over 5 min followed by isocratic 50% MeCN:$H_2O$ until product was detected. *Illustration description:* Part of crude product NMR fits **S1** standard NMR, so it is labeled as major product, however, like FUT6, FUT5 failed to maintain selective monofucosylation, with the initially formed monofucosylated product readily converted into di- and trifucosylated products. Preliminary LC–MS analysis indicated that controlling the equivalents of GDP-fucose was ineffective for improving selectivity with either FUT5 or FUT6. In contrast, FUT9 favored the formation of the monofucosylated product **S1** under controlled GDP-fucose stoichiometry. Therefore, no further investigation of GDP-fucose stoichiometry was carried out for FUT5 or FUT6.

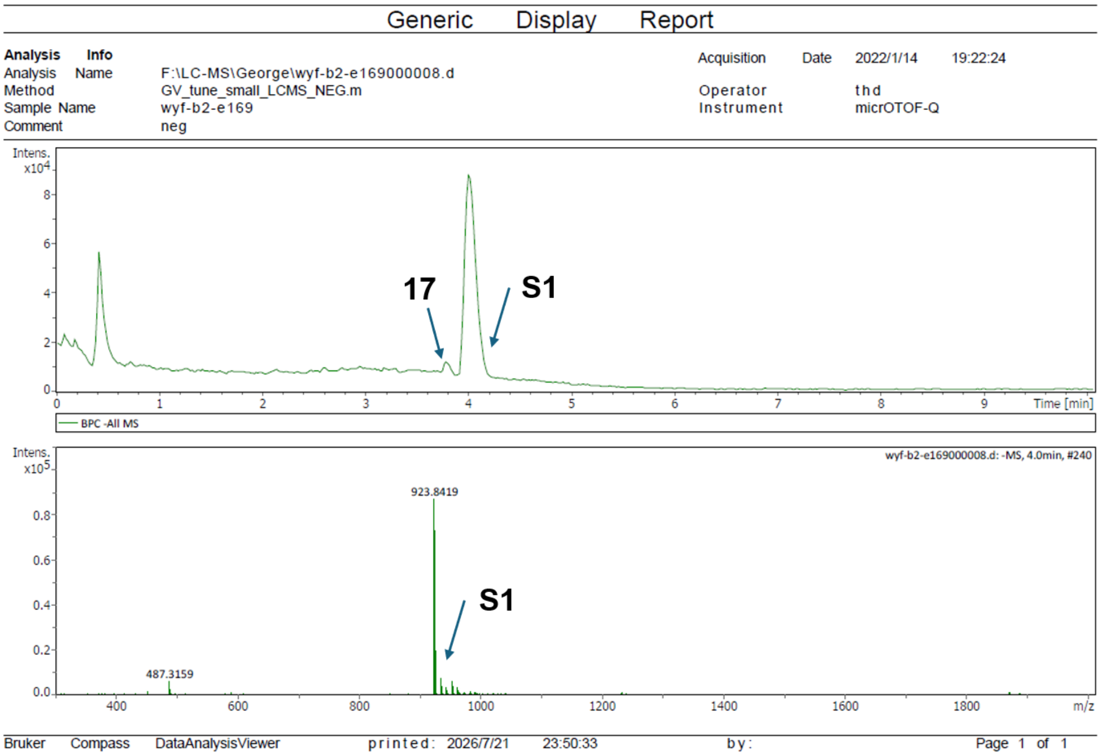

**LC-MS for Scheme S1a, FUT9 rate control.** Reaction progress for carbohydrates was monitored using a SeQuant Zic HILIC guard column (20x2.1 mm) coupled to a Bruker micrOTOF-QII MS system running a gradient of 90% MeCN:$H_2O$ to 50% MeCN:$H_2O$ over 5 min followed by isocratic 50% MeCN:$H_2O$ until product was detected. *Illustration description:* The m/z value of **17** which is 850.7710 was manually added because the corresponding peaks had very low intensity percentages, and the software did not display the full values automatically.

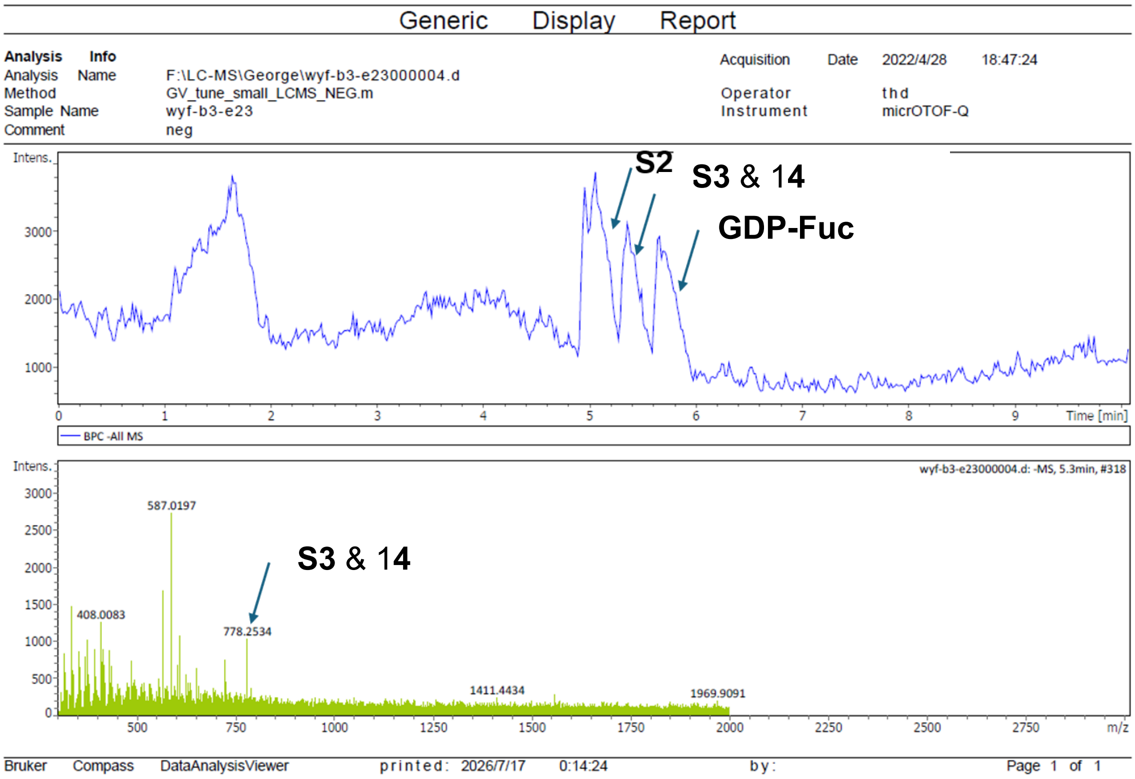

**LC-MS for Scheme S1b.** Reaction progress for carbohydrates was monitored using a SeQuant Zic HILIC guard column (20x2.1 mm) coupled to a Bruker micrOTOF-QII MS system running a gradient of 90% MeCN:$H_2O$ to 50% MeCN:$H_2O$ over 5 min followed by isocratic 50% AcCN:$H_2O$ until product was detected. *Illustration description:* compounds **S3** and **14** showed very similar retention times on the LC-MS HILIC column, resulting in a peak that was almost impossible to separate. The ratio between them was determined after SEC purification by comparing the integration of characteristic proton signals in the NMR spectrum with the corresponding signals from the established standard spectrum.

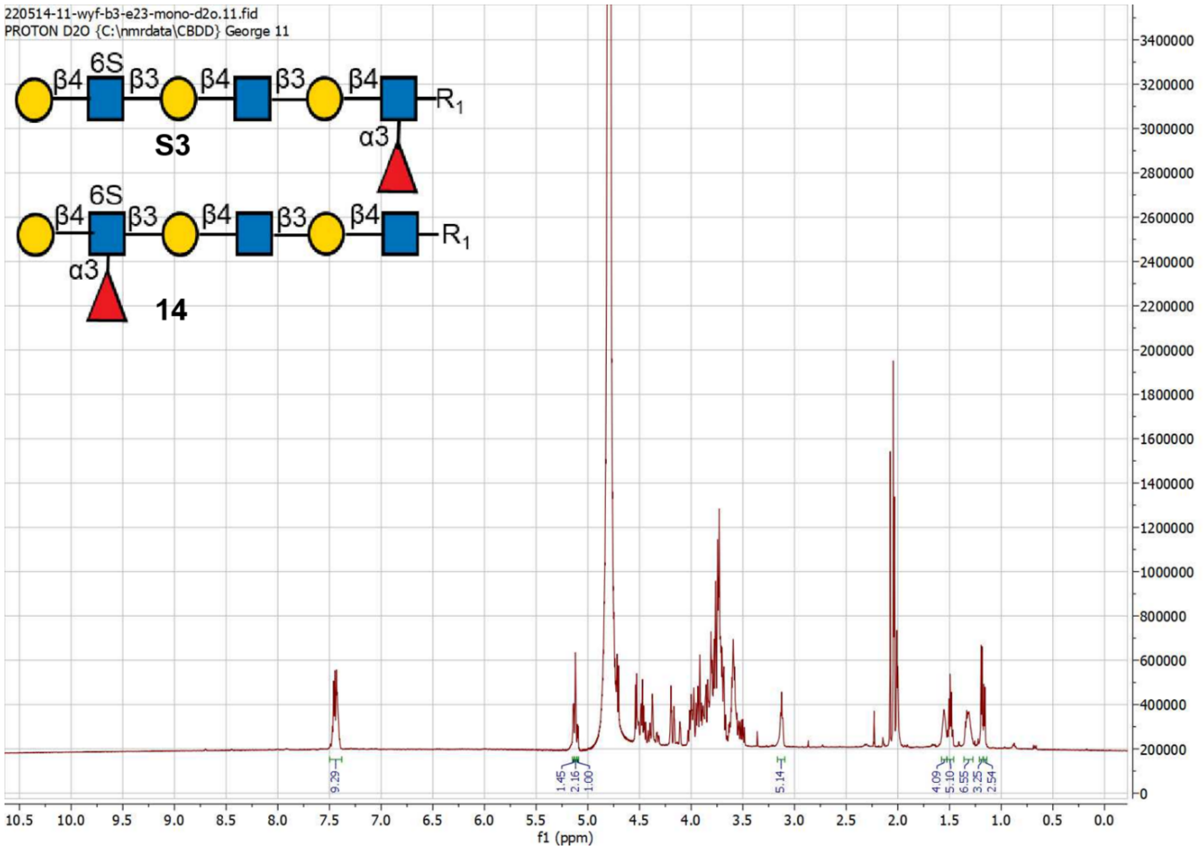

600 MHz 1D [1]H NMR of crude mixture of **S3** and **14**, recorded at 298K in $D_2O$.

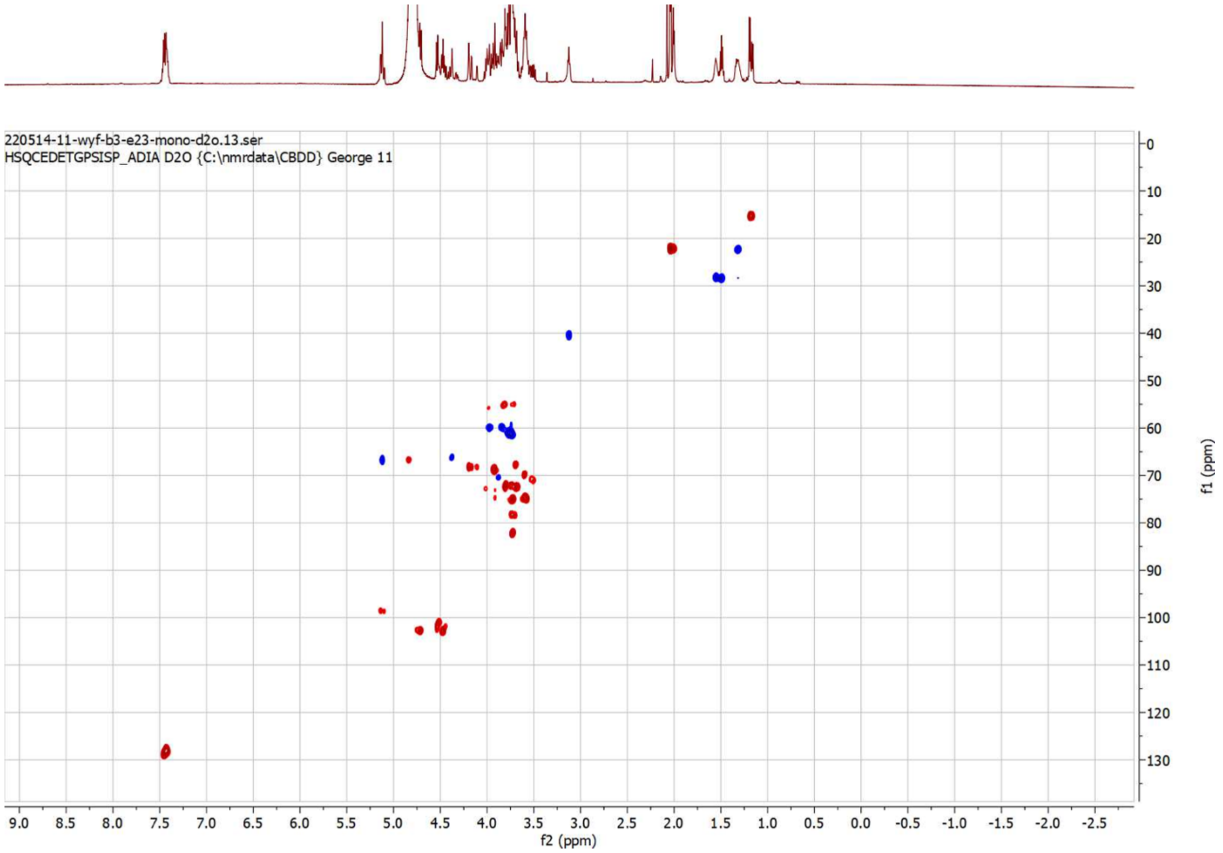

150 MHz HSQC of crude mixture of **S3** and **14**, recorded at 298K in $D_2O$.

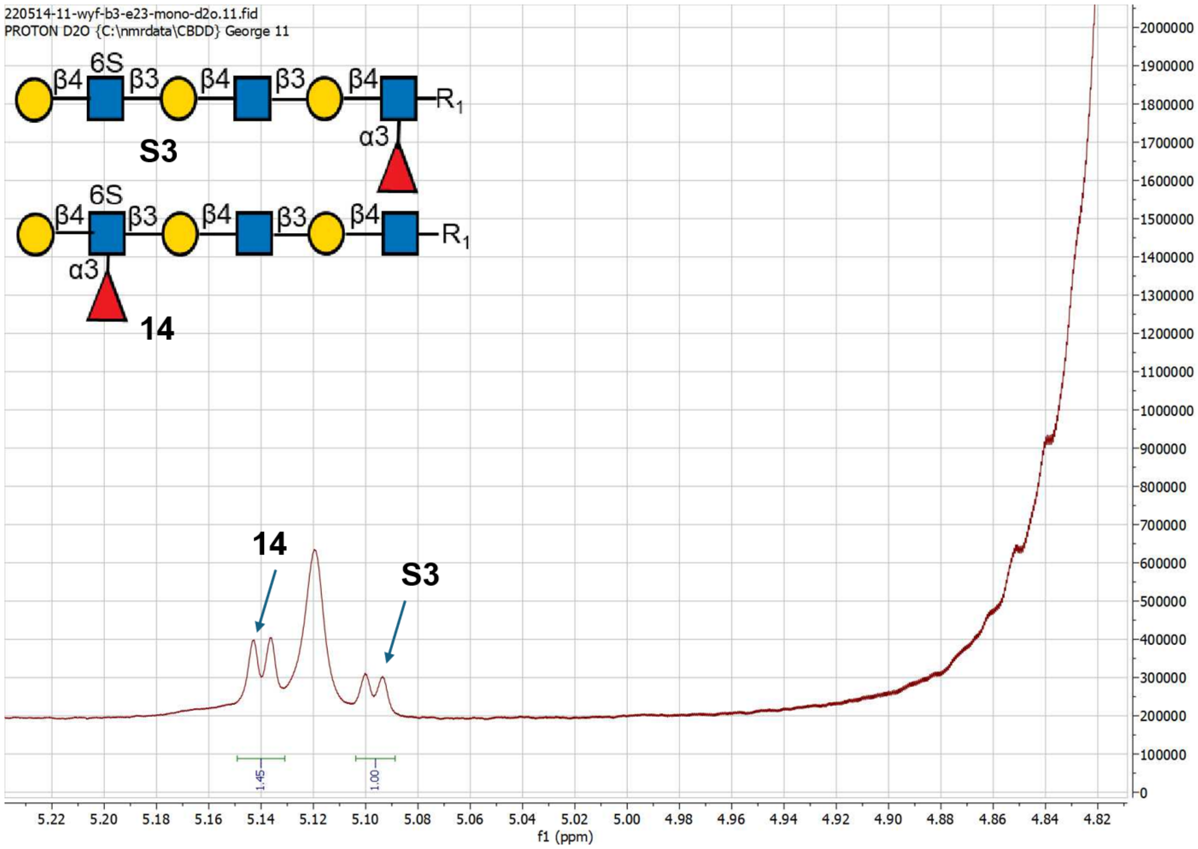

[1]H NMR spectra (δ 4.8 to 5.2) of **14** and **S3**. When comparing the H1 of Fucose of **14** to H1 of Fucose of **S3**, the **14** : **S3** ratio is 1.46:1.

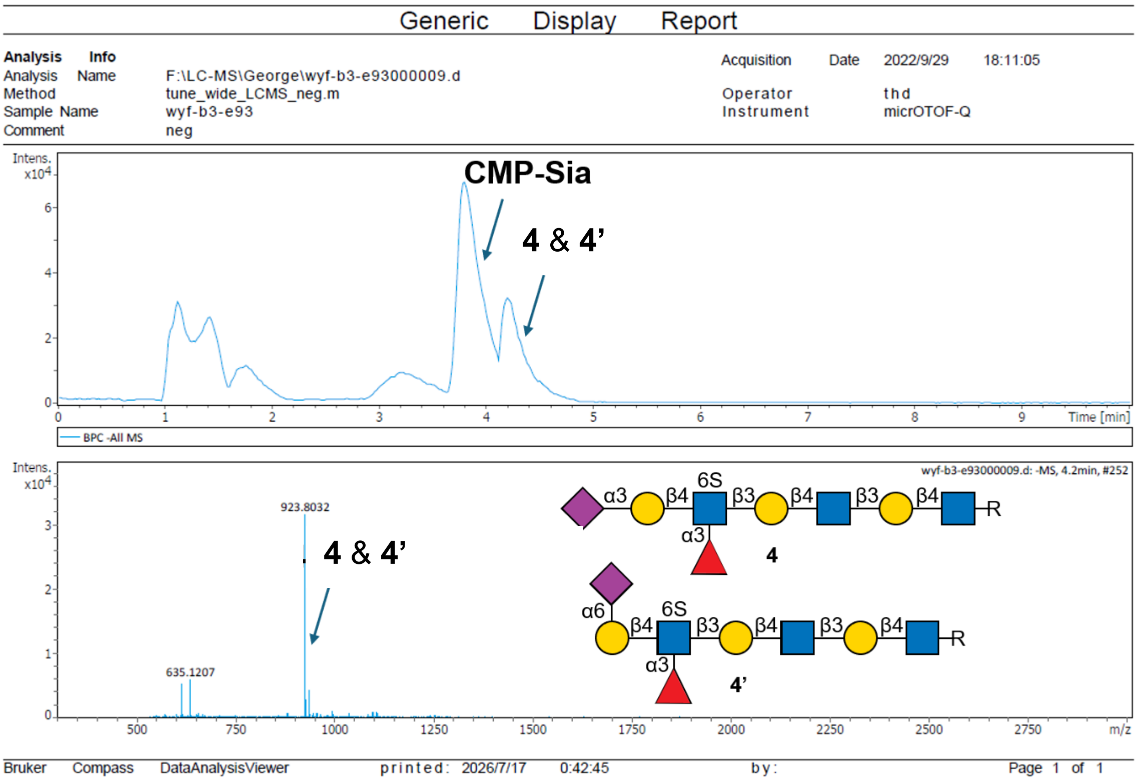

**LC-MS for Scheme S3, PmST1 M144D.** Reaction progress for carbohydrates was monitored using a SeQuant Zic HILIC guard column (20x2.1 mm) coupled to a Bruker micrOTOF-QII MS system running a gradient of 90% MeCN:$H_2O$ to 50% MeCN:$H_2O$ over 5 min followed by isocratic 50% MeCN:$H_2O$ until product was detected. *Illustration description:* Compounds **4** and **4'** showed very similar retention times on the LC-MS HILIC column, resulting in a peak that was almost impossible to separate. The ratio between them was determined after SEC purification by comparing the integration of characteristic proton signals in the NMR spectrum with the corresponding signals from the established standard spectrum (Figure S1).

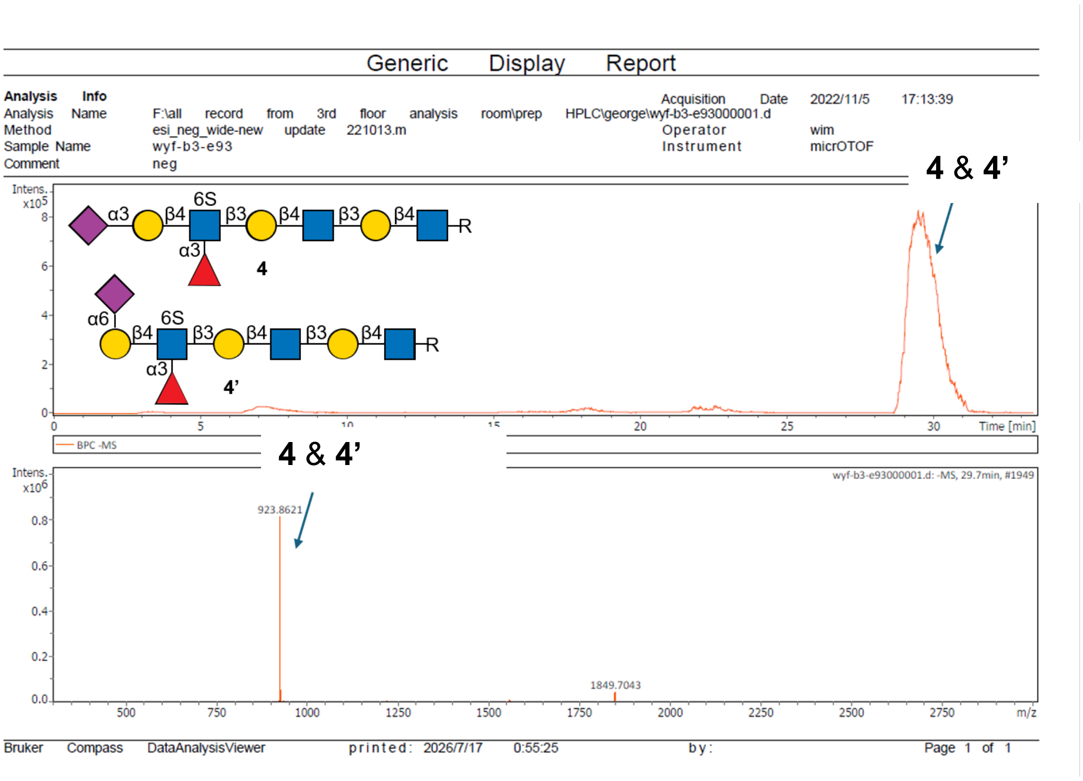

**Prep-HPLC-MS for Scheme S3, PmST1 M144D.** General protocols for HILIC-HPLC purification were used for purification**.** *Illustration description:* Because the ratio between **4** and **4′** was too close (1:1.7) and their retention time was almost identical, the two peaks completely overlapped and appeared as a single unresolved peak. Attempts using various eluent polarity ratios failed to achieve separation. Unlike the Pd2,3ST experiments, where the injected sample was the crude reaction mixture after enzyme removal by SEC, the sample analyzed here was a relatively pure mixture obtained from multiple preparative HPLC injections for product isolation. Consequently, a slight shift in retention time was observed.

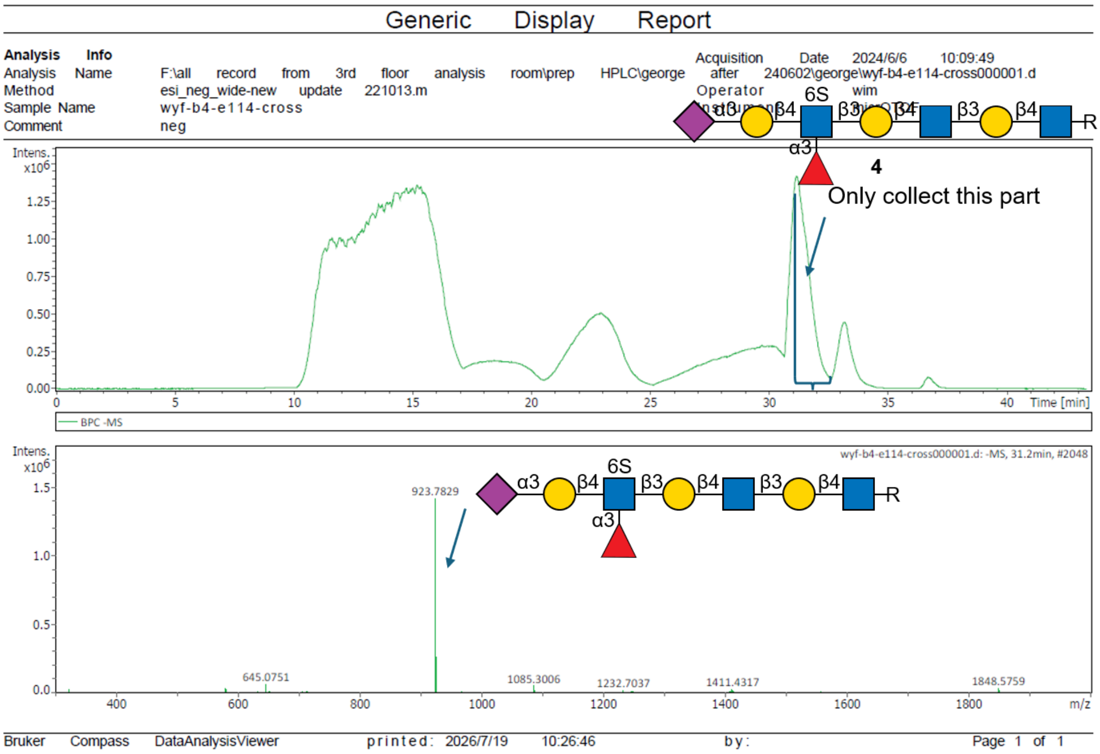

**Prep-HPLC-MS for Scheme S3, Pd2,3ST. General protocols for HILIC-HPLC purification** is used for the purification**.** *Illustration description:* Similarly, because the ratio between α2,3 and α2,6 was significantly different (>10:1), although their retention times were close, it was possible to obtain pure α2,3 by manually collecting the fractions corresponding to the most enriched peak. However, this approach was still considered suboptimal, and therefore we further developed the $NH_2$ control Fucose strategy to achieve a more reliable and efficient solution.
